# The Marburg Collection: A Golden Gate DNA Assembly Framework for Synthetic Biology Applications in *Vibrio natriegens*

**DOI:** 10.1101/2021.03.26.437105

**Authors:** Daniel Stukenberg, Tobias Hensel, Josef Hoff, Benjamin Daniel, René Inckemann, Jamie N. Tedeschi, Franziska Nousch, Georg Fritz

## Abstract

*Vibrio natriegens* is known as the world’s fastest growing organism with a doubling time of less than 10 minutes. This incredible growth speed empowers *V. natriegens* as a chassis for synthetic and molecular biology, potentially replacing *E. coli* in many applications. While first genetic parts have been built and tested for *V. natriegens*, a comprehensive toolkit containing well-characterized and standardized parts did not exist. To close this gap, we created the Marburg Collection – a highly flexible Golden Gate Assembly-based cloning toolbox optimized for the emerging chassis organism *V. natriegens*. The Marburg Collection overcomes the paradigm of plasmid construction – integrating inserts into a backbone – by enabling the *de novo* assembly of plasmids from basic genetic parts. This allows users to select the plasmid replication origin and resistance part independently, which is highly advantageous when limited knowledge about the behavior of those parts in the target organism is available. Additional design highlights of the Marburg Collection are novel connector parts, which facilitate modular circuit assembly and, optionally, the inversion of individual transcription units to reduce transcriptional crosstalk in multigene constructs. To quantitatively characterize the genetic parts contained in the Marburg Collection in *V. natriegens*, we developed a reliable microplate reader measurement workflow for reporter experiments and overcame organism-specific challenges. We think the Marburg Collection with its thoroughly characterized parts will provide a valuable resource for the growing *V. natriegens* community.

## Introduction

The gram-negative bacterium *Vibrio natriegens* was first isolated in 1958 (1) and has received increasing attention due to its extraordinary fast growth with a doubling time of less than 10 minutes (2–4). Additionally, *V. natriegens* possesses several highly useful properties including natural competence (5) and a high number of ribosomes (6), which have been utilized for protein production, both *in vivo* (7,8) and in cell-free setups (9–11). Further, *V. natriegens* is capable of exploiting a wide range of substrates, including cheap carbon sources such as sucrose and glycerol (3,12), inspiring first attempts to turn *V. natriegens* into a cell factory for the production of small molecules (5,12–14) or amino acids (3). These above-mentioned examples highlight the diverse range of capabilities of *V. natriegens*, making it a desirable model organism for various biotechnology and synthetic biology applications.

A prerequisite for widespread adoption of *V. natriegens* as a chassis for biotechnology and synthetic biology is the establishment of well-characterized genetic parts, plasmids and methods tailored to this organism. Recently, several strategies for genome engineering have been published (5,15,16) and different transformation protocols for plasmids have been established (15,17,18). In addition, several genetic parts from *E. coli* (15,17,18) and from synthetic libraries (19) have been functionally characterized in *V. natriegens*. However, to date, there exists no simple, one-stop solution for quickly building multi-gene constructs from a library of *V. natriegens*-tested DNA parts.

Over the recent years, many synthetic biology toolboxes were developed based on the Golden Gate cloning method (20), using one-pot restriction/ligation reactions based on type IIs restriction enzymes to quickly assemble DNA parts. The modular cloning (MoClo) system (21), for instance, allows the rapid assembly of multi-gene constructs in a hierarchical manner, and several toolboxes with basic genetic parts (promoters, ribosome binding sites, terminators, etc.) were published for important model organisms such as *E. coli* (22,23), *S. cerevisiae* (24) and *B. subtilis* (25).

Here we describe the design, construction and evaluation of a novel Golden-Gate-based toolbox, termed the Marburg Collection, optimized for working in *V. natriegens*. Through a detailed, quantitative characterization of the genetic parts in this new chassis organism, this toolbox will provide the growing *V. natriegens* community with the resources to work efficiently with this unique bacterium. We hope that this will contribute to fully harnessing the potential of *V. natriegens* as a novel, ultra-fast-growing chassis organism for synthetic biology and biotechnology.

## Results and Discussion

### Rationale of the Marburg Collection

The general cloning strategy of the Marburg Collection is based on hierarchical Golden-Gate-based DNA assembly (20,21). The cloning scheme starts with a library of level 0 plasmids storing basic genetic parts (e.g., promoters, coding sequences (CDSs), terminators), which are assembled into a single transcription unit (TU) harbored on a level 1 plasmid. Level 1 plasmids can then be combined to create higher level plasmids (level 2, level 3, etc.) carrying multiple transcription units. This hierarchical cloning workflow is enabled by alternating type IIs restriction enzymes, which cleave DNA outside of their recognition sequence. Thereby, a single enzyme can result in a large range of fusion sites, which define the order of the assembled DNA fragments (20).

A detailed overview on the assembly scheme of the Marburg Collection is provided in Figure 1A. The Marburg Collection comes with a library of level 0 plasmids, each carrying one basic genetic part.

**Figure 1.**
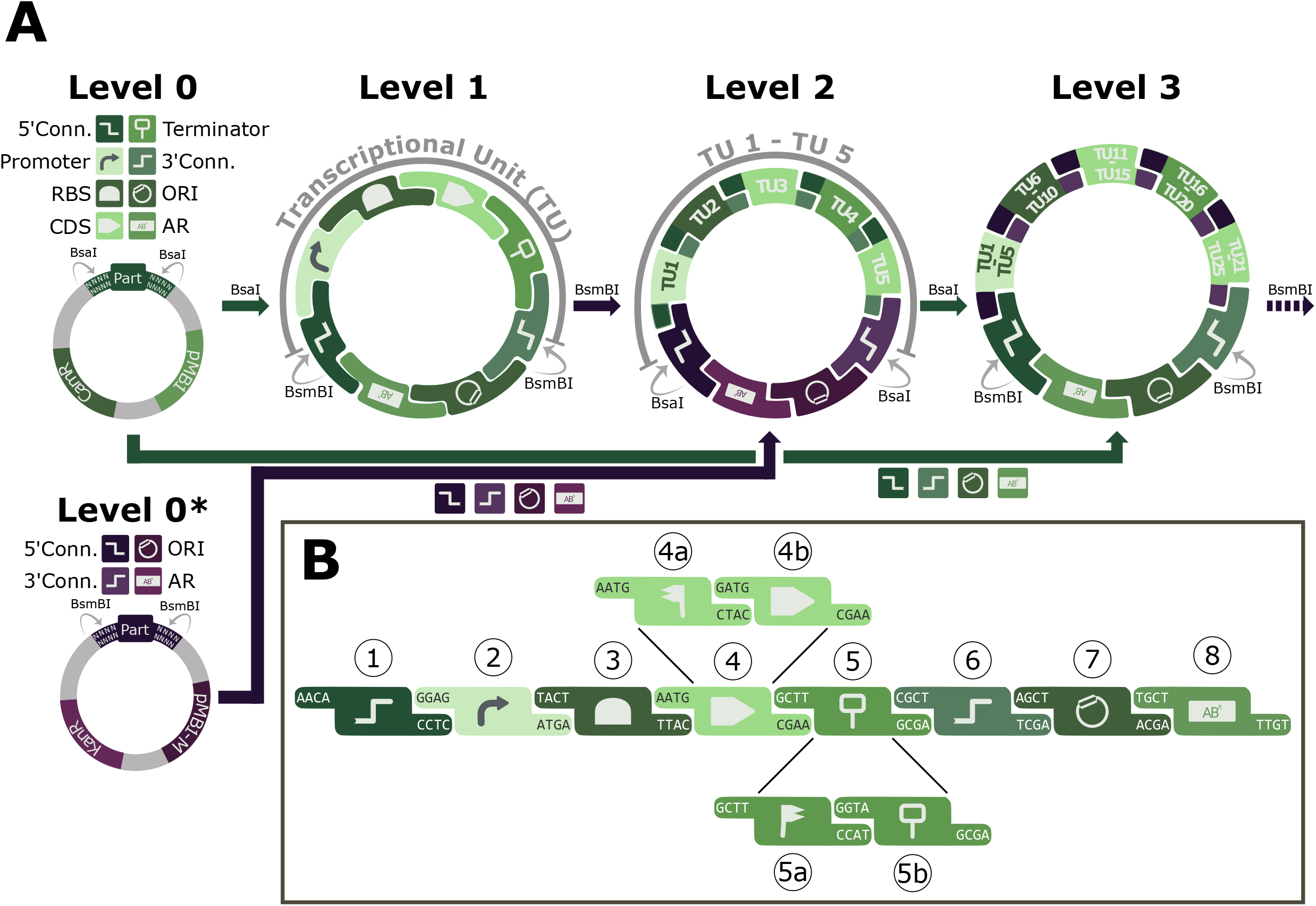
General overview and fusion sites. (**A**) Basic genetic parts (level 0) are stored on plasmids with a chloramphenicol resistance marker and can be used to assemble a plasmid harboring a single transcriptional unit (TU) (level 1) by using the type IIs restriction enzyme BsaI. All sequences of the level 1 plasmid are provided as level 0 parts (shown in green) and no entry vector is required. Up to five level 1 plasmids can be used to create a level 2 plasmid by using BsmBI, also a type IIs restriction enzyme. Backbone components of level 2 plasmids are provided as level 0* parts (shown in purple), which are stored on plasmids with a kanamycin resistance marker. Further cloning (e.g., level 3, level 4, and so on) is possible by alternating between BsaI and level 0 parts (for odd number level plasmids) and BsmBI and level 0* parts (for even number level plasmids). (**B**) Defined 4 bp overhangs ensure assembly of level 0 parts in correct order. Fusion sites are largely based on the Phytobrick standard. Part categories can be split into subcategories to introduce additional functions, e.g., the N- and/or C-terminal tagging of proteins.

Level 0 parts are flanked on both sides by inward-facing BsaI restriction sites, which generate the overhangs for the directional assembly of these basic parts into a level 1 plasmid (Figure 1B). A unique feature of the Marburg Collection is the possibility to assemble plasmids completely *de novo* from individual level 0 parts, by providing separate parts for the plasmid replication origin (ORI) and the antibiotic resistance marker (AR). This offers users the ability to freely choose any combination of these parts, which are traditionally combined on a plasmid backbone. Possible applications of this flexibility include, for instance, the construction of two compatible plasmids for co-transformation or the adjustment of gene expression levels by selecting an ORI with a desired plasmid copy number.

In the Marburg Collection, the TU parts (promoter, RBS, CDS, terminator) are linked to the ORI and AR parts by novel 5’ and 3’ connector parts (Figure 1A). These connector parts carry BsmBI restriction sites, which, upon restriction digestion, generate specific 4 bp fusion sites that determine the position of the respective TU in the subsequent level 2 assembly (Figure 1B, Figure 2). Up to five level 1 TUs can be used for the construction of a level 2 plasmid. For a level 2 assembly, the ORI, AR, and connector parts are supplied as level 0* parts. Level 0* parts can be regarded as basic parts in the assembly of level 2 plasmids. They differ from level 0 parts in the fact that they are stored in plasmids carrying a kanamycin marker instead of a chloramphenicol marker, and that the restriction enzyme BsmBI is used for the release of parts (instead of BsaI in case of level 0 parts). With these level 0* parts, the assembly of level 2 plasmids has the same flexibility of choosing the ORI and AR parts as described for level 1 plasmids. Further cloning is again enabled through level 0* connector parts carrying BsaI recognition sites, which allow for the construction of level 3 plasmids containing up to 25 Tus (Figure 1A). The backbone components for level 3 plasmids are provided as level 0 parts. Theoretically, cloning can proceed past level 3 by combining up to five plasmids from the previous level. As a rule, odd number level plasmids are built with BsaI and level 0 parts, while even number level plasmids are created using BsmBI and level 0* parts. The creation of new project-specific level 0 parts is explained in detail in the Supplementary Text.

**Figure 2.**
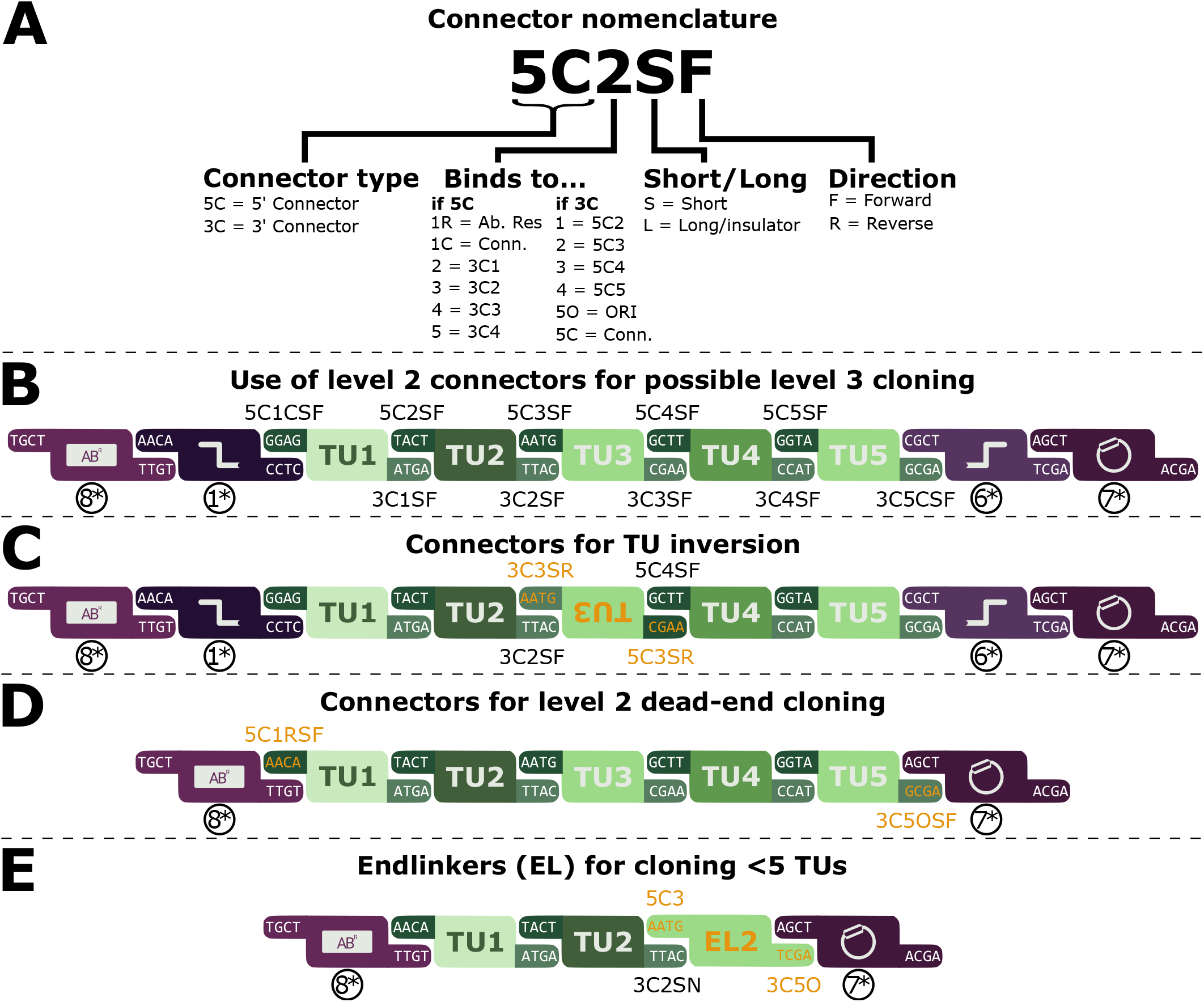
Nomenclature of connector parts and options for level 2 assembly. (**A**) 5’ and 3’ connector level 0 parts are labelled according to a defined nomenclature. The characters describe the connector type, the connectivity, the length of the connector (presence or absence of insulator sequence) and the direction of the TU in the resulting level 2 plasmid. (**B**) Standard layout of a level 2 plasmid with five TUs. Bases facilitating the assembly are indicated in the overhangs of each fragment. The nomenclature of the connectors integrated into the level 1 plasmids are displayed above and below the TU overhangs. (**C**) Assembly example for the inversion of a single TU. Deviations from the previous layout are highlighted in orange. In the inverted TU, the 3’ connector of TU3 (3C3SR) binds to the 3’ connector of TU2 (3C2SF) and 5’ connector of TU3 (5C3SR) binds to the 5’ connector of TU4 (5C4SF). (**D**) Level 0* connector parts (1* and 6*) can be omitted to reduce the number of fragments in the assembly reaction while preventing further cloning into level 3. The 5’ connector of TU1 (5C1RSF) and the 3’ connector of TU5 (3C5OSF) bind directly to the AR and ORI part, respectively. (**E**) Endlinker (EL) parts can be used to create level 2 plasmids with less than 5 TUs. EL parts connect the last TU with the ORI part.

### Adaptable assembly of level 1 plasmids

The 4 bp fusion sites flanking level 0 parts determine the order in which these parts assemble into level 1 plasmids (Figure 1B). The sequences of these fusion sites are based on the bacterial version of the Phytobrick standard, developed in the framework of the International Genetically Engineered Machine (iGEM) competition (http://2016.igem.org/Resources/Plant_Synthetic_Biology/PhytoBricks). As such, these fusion sites, and consequently the parts of this toolbox, are largely compatible with existing modular cloning toolboxes such as MoClo (21) and Loop Assembly (26), allowing flexible exchange of level 0 parts between research groups using different toolboxes. However, we note that while BsaI is used in all the mentioned toolboxes for the construction of level 1 plasmids, different enzymes are used for further assembly steps. This fact might require elimination of incompatible restriction sites if parts from different toolboxes are to be used in higher level plasmids. In addition to the standard Phytobrick fusion sites, we designed novel fusion sites for Marburg Collection-specific part categories, namely ORI, AR and 5’ and 3’ connector parts, as described previously (27). To facilitate standard cloning workflows, we defined eight level 0 part categories required for the construction of a level 1 plasmid (Figure 1B). The elements of each part category typically fulfill the same biological function; For example, category 2 represents promoter parts and category 4 represents CDS parts. All part categories and the fusion sites by which they are connected are shown in Figure 1B. In addition, some part-categories can be split into subcategories to enable the construction of N- and/or C-terminal tagging of proteins. For N-terminal tags, the CDS category 4 is split into 4a and 4b parts, where 4a represents the N-terminal tag and 4b the CDS to be tagged. In case of C-terminal tags, the terminator part category (5) is split into a C-terminal tag (5a) and a terminator for C-terminally tagged proteins (5b) (Figure 1B). Note that when using parts of a subcategory, all corresponding parts need to be added for a successful assembly, e.g., 4a and 4b instead of 4.

### Flexible orientation and positioning of transcription units in level 2 and beyond

A major difference between the Marburg Collection and existing cloning toolboxes is its superior flexibility, which offers a wide range of cloning possibilities. This is especially evident in the assembly of level 2 plasmids, in which the novel 5’ and 3’ connector parts play a central role. Our nomenclature system describes the properties of each connector part (Figure 2A). The first two characters (5C or 3C) discriminate between 5’ and 3’ connector parts, followed by a number, which describes the ‘connectivity’ of that connector. Generally, a 5’ connector with the connectivity *n* binds to a 3’ connector with the connectivity *n-1*. The next letter is either S (short), or L (long). We initially intended to include insulators into the connector parts to reduce crosstalk between neighboring TUs. Even though the implementation of such insulators could not be realized in this project, we include this letter in the nomenclature to facilitate future efforts in this direction. The final letter in the connector nomenclature is either F (forward) or R (reverse), indicating whether the respective TU will be included in forward (5’ to 3’) or in reverse orientation (3’ to 5’) in the resulting level 2 plasmid.

An overview on some possible level 2 assemblies and the corresponding use of different connector parts is shown in Figure 2B-E. In the standard scheme of a level 2 plasmid with five TUs, the TUs are joined by fusion sites in their flanking connector parts, as determined in the previous level 1 assembly (Figure 2B). In this setup, the 3’ connector of one TU binds to the 5’ connector of the following TU. As an exception, the 5’ connector of the first TU and the 3’ connector of the last TU bind to level 0* connectors to allow higher level assemblies. Additionally, the Marburg Collection allows for the inversion of individual TUs in a level 2 plasmid. This was achieved by designing the overhangs so that the 3’ connector of the inverted TU binds to the 3’ connector of the upstream TU and the 5’ connector of the inverted TU binds to the 5’ connector of the downstream TU (Figure 2C). In this way, it is possible to invert any single or even multiple TUs to achieve alternating orientations, which has previously been shown to reduce crosstalk caused by transcriptional read-through (28).

In cases when it is not desired to proceed past level 2, it is possible to omit the level 0* connector parts and thereby decrease the number of fragments in the assembly, presumably increasing the cloning efficiency. This is achieved by assembling TU1 and TU5 with specific 5’ and 3’ connectors that directly connect to the AR and ORI parts, respectively (Figure 2D). If less than five TUs are required for an application, unused positions can be skipped with end linker (EL) parts, which connect the 3’ connector of the last TU with the ORI part. For example, a level 2 plasmid can be composed of only two TUs plus an EL that skips the positions three to five (Figure 2E). The Marburg Collection provides a set of four ELs, each of which binds to the 3’ connector of the TU in the different positions. This allows for the construction of level 2 plasmids with one to five TUs. An alternative approach with the same outcome would be to create the last TU in the final plasmid with a 3’ connector that directly binds to the ORI. However, the approach to use EL parts enables re-using of level 1 plasmids in case a derivative of the level 2 plasmid with additional TUs is required later in the project.

### Pre-assembled plasmid backbones and dropout parts enable “convenience cloning”

By supplying traditional plasmid backbone elements (ORI and AR) as separate genetic parts, the Marburg Collection offers users with maximal modularity for *de novo* plasmid assembly. However, when the same combination of ORI and AR are used for the majority of experiments in a project, it can be convenient to start with a set of pre-assembled backbones, resembling position vectors of existing Golden Gate toolboxes (21–23). For this reason, we provide a set of pre-assembled backbone plasmids, which feature different 5’ and 3’ connector parts defining the TU’s position in the subsequent level 2 cloning (Figure 3A, left). These pre-assembled vectors contain a “dropout part”, carrying a fluorescent protein expression cassette for either sfGFP or mScarlet-I, that can be substituted by TU parts (promoter, RBS, CDS, terminator) during a level 1 assembly. As a result, the replacement of this dropout with the respective TU parts is indicated by the loss of the fluorescent signal, which enables easy selection of correct colonies grown on agar plates. This dropout therefore assists in the selection of positive clones, reducing the required screening effort. In addition to dropout parts spanning an entire TU, we also developed dropout parts that serve as placeholders for individual level 0 parts (Figure 3A, right). This allows for the construction of pre-assembled plasmids, carrying a dropout in one position and standard level 0 parts for the rest of this plasmid. This approach is useful if a high number of plasmids differing by only a single level 0 part are required. Applications of this strategy include, e.g., the use of a promoter dropout plasmid for the screening of promoter libraries or a CDS dropout plasmid as a protein expression plasmid.

**Figure 3.**
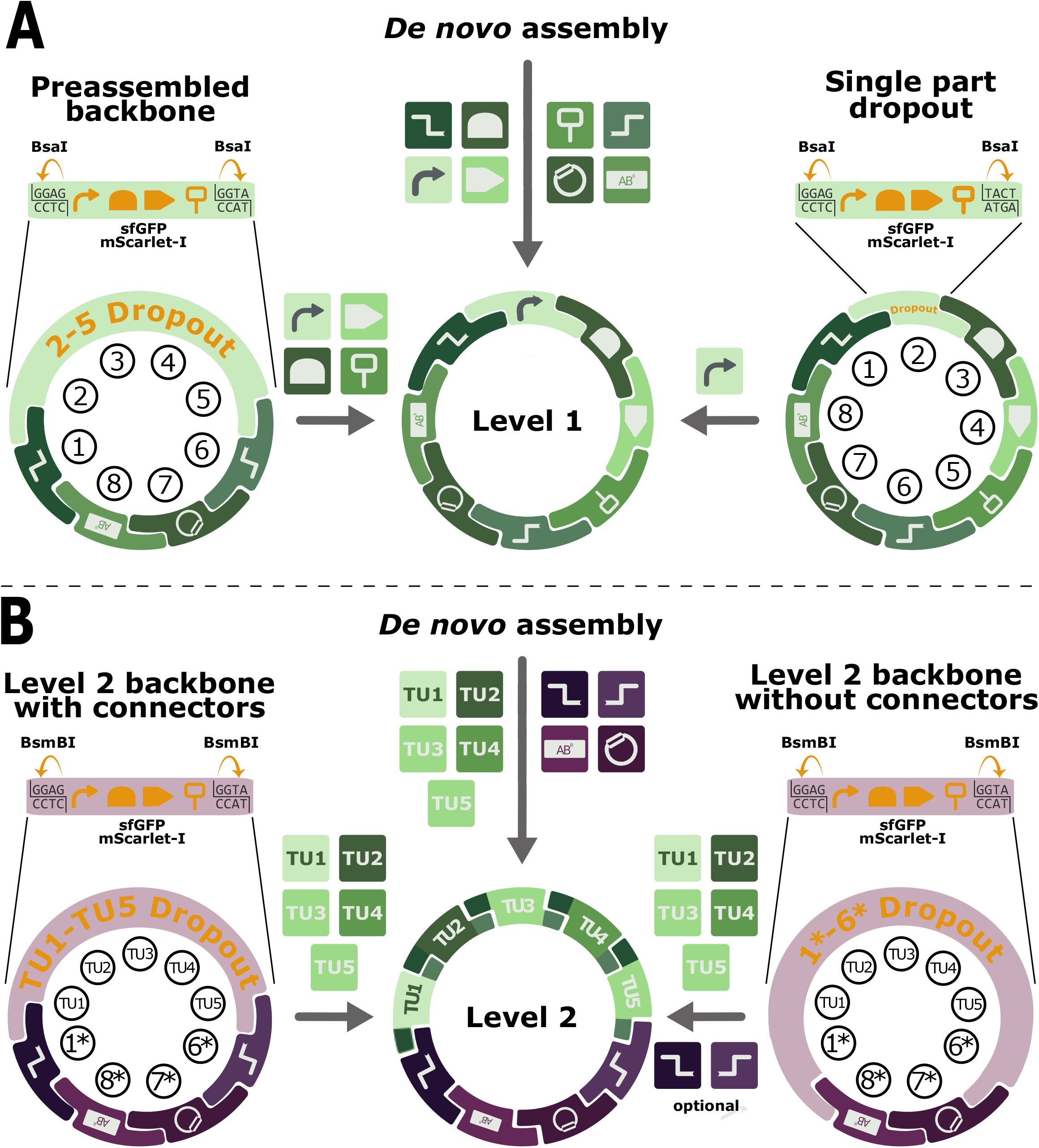
Dropout parts serve as placeholder and enable construction of pre-assembled plasmids. (**A**) Dropout parts carry a full expression cassette for the fluorescent proteins sfGFP or mScarlet-I to enable visible distinction of correct colonies and outward facing BsaI or BsmBI recognition sites. These restriction enzyme recognition sites remain on the plasmid and allow for dropout replacement. (A, left) A dropout part spanning the part categories 2-5 can be combined with the remaining part categories (1, 6–8) to create a pre-assembled backbone plasmid. Once assembled, this backbone can be used to integrate the part categories covered by the dropout part (promoter, RBS, CDS, terminator) to obtain the functional level 1 plasmid. (**A**, right) Dropout parts are available as placeholders for individual part categories. A plasmid with a promoter dropout is shown as an example, however, a set of dropout parts were created for the part categories 1-6. (**B**) Dropout parts are also available as level 0* parts (in purple) and allow for the construction of pre-assembled level 2 backbones with or without level 0* connector parts.

In fact, during this project, we typically used the approach to first create dropout plasmids and then subsequently replace the dropout with the respective parts. We found this strategy to be extremely reliable and consequently decided to provide parts for the creation of level 2 dropout plasmids. Thereby, users of the Marburg Collection can create backbones for level 2 plasmids in a similar way as described previously for level 1 plasmids. This can be achieved by preassembling either level 0* ORI and AR parts (Figure 3B, left) or by additionally including the level 0* connectors (Figure 3B, right). These backbones can then host multiple TUs with all options as previously described (Figure 2B-E).

In total, the Marburg Collection consists of 175 individual level 0 and level 0* parts (Figure 4). In addition to structural parts, such as 5’ and 3’ connector parts required for the various assembly options described above, we created a large library of thoroughly characterized regulatory parts (see below). Most of these regulatory parts are natural or synthetic sequences, which have been widely used in *E. coli*. Finally, we added eight parts encoding codon-optimized fluorescent proteins for maximal expression in *V. natriegens*, enabling the quantitative characterization of Marburg collection parts. Plasmid maps are provided in the Supplementary Information.

**Figure 4.**
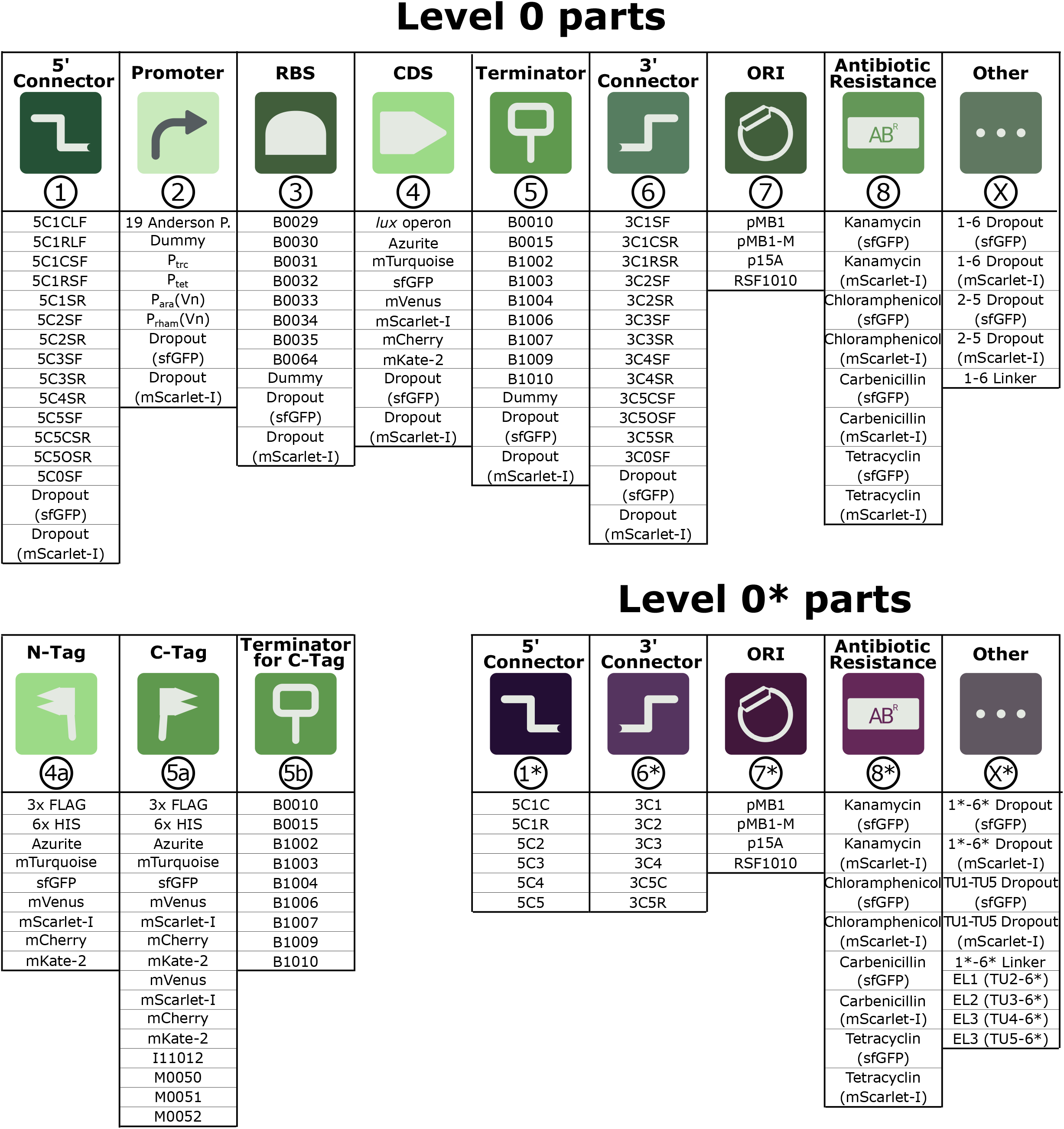
Table of all parts contained in the Marburg Collection. List of all level 0 parts (top), level 0 part subcategories (bottom left) and level 0* parts (bottom right). Numbers in circles refer to part categories and the corresponding fusion sites can be obtained from Figure 1B (level 0) and Figure 2B (level 0*). Part category X and X* translate as *miscellaneous* and contain parts which do not fall into any standard category.

### Quantitative characterization of genetic parts in *V. natriegens*

To provide users of the Marburg Collection with quantitative information about the genetic parts of the Marburg Collection, we experimentally characterized their behavior in *V. natriegens*. The modular nature of the cloning toolbox presented in this work inevitably leads to a “chicken and egg” problem for the choice of the plasmid context for the characterization of a particular part category. For example, it is necessary to choose a reporter gene for the experiments to characterize promoter parts, and conversely the validation of those reporter genes requires the availability of a suitable promoter. Several experimental iterations were necessary for the selection of a suitable plasmid context for the characterization of each part category. The data presented here represent the last round of this iteration. As mentioned before, the general experimental strategy for the construction of the test plasmids was to firstly create a pre-assembled plasmid containing a dropout part in the position of the respective part category. In a subsequent step, we replaced this dropout with the level 0 parts to create the plasmids which were then used for the characterization experiments. Due to the high efficiency of dropout replacement cloning, the assembly reaction was directly transformed into chemically competent *V. natriegens* cells. Four colonies of each cloning reaction were used in the experiments, representing biological replicates.

### Characterization of promoter parts

We characterized a range of constitutive and inducible promoter parts with mScarlet-I as the reporter gene. The short synthetic promoters obtained from the Anderson Promoter Library (http://parts.igem.org/Promoters/Catalog/Anderson) spanned at least three orders of magnitude in promoter activity, while the weakest promoters did not produce a signal significantly higher than an empty plasmid control strain (Figure 5B). The relative strengths of these promoters in *V. natriegens* were strongly correlated (*R^2^* = 0.88) with previous results from *E. coli* (http://parts.igem.org/Promoters/Catalog/Anderson) (Figure 5C). This is especially remarkable considering our experimental setup differed not only in the organism, but also in the reporter gene and measurement protocol. This strong correlation suggests that the transcription machinery is highly conserved between *E. coli* and *V. natriegens*, and therefore is likely that promoters from *E. coli* are generally functional in *V. natriegens* and *vice versa*. Indeed, recent measurements confirmed that the absolute promoter activity (RNAP/s) of one member of the Anderson Promoter Library (J23101) is highly similar between *V. natriegens* and *E. coli (Shao et al. 2021). We* also characterized four inducible promoters by first testing two well-established parts from *E. coli*, obtaining a ~170-fold induction of the P_tet_ promoter when induced with anhydrotetracycline (ATc) (Figure 5D), but with almost no visible induction for P_trc_ when induced with Isopropyl β-D-1-thiogalactopyranoside (IPTG) (Figure 5E). Then, we established endogenous sequences of *V. natriegens* as novel, tightly regulated inducible promoters. By using putative promoter regions of the arabinose and rhamnose utilization operons, respectively, we measured a high level of induction with high inducer concentrations, but only at high OD_600_ (Figures 5F and 5G), indicating a growth-phase dependent regulation of both promoters. However, this pattern was not visible for P_tet_, which did not show an obvious growth-phase dependency (Supplementary Figure S1). Note that these experiments were conducted in a *V. natriegens* strain carrying the functional arabinose and rhamnose utilization systems. In addition, while P_tet_ and P_trc_ carry the regulatory protein within the level 0 promoter part, the endogenous promoters rely solely on the chromosome-encoded copy of the regulators AraC and RhaS. It is therefore likely that the strong growth-phase dependency is due to the natural reaction of *V. natriegens* to alternative carbon sources. Further modification of the *V. natriegens* strain might abolish the growth-phase dependent regulation, as shown for the arabinose inducible P_BAD_ promoter of *E. coli* (29). For example, constitutive expression of the transporters and deletion of the genes encoding the arabinose and rhamnose catabolizing enzymes can potentially lead to novel, tightly regulated inducible promoters for *V. natriegens* in the future.

**Figure 5.**
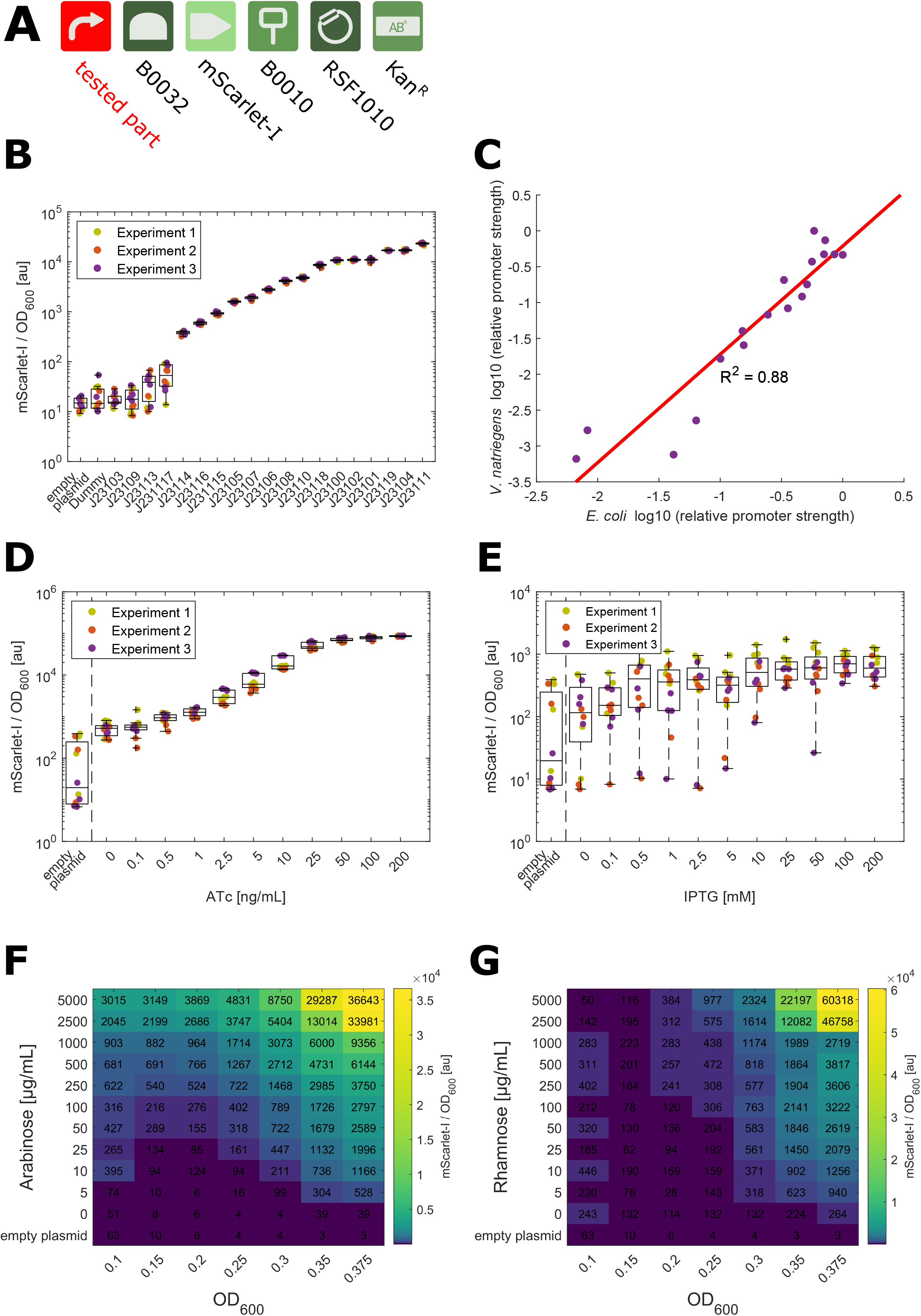
Results of promoter part characterization. (**A**) Plasmid pDS_114_1R1 kan (Table S5) was used as a dropout plasmid for all promoter experiments. (**B**) Characterization of synthetic promoters from the Anderson Promoter library. The fluorescent signal of mScarlet-I was normalized by the OD_600_ during the exponential growth phase (see Materials and Methods). Twelve data points are shown for each sample, representing four biological replicates from three independent experiments (shown in distinct colors). Data are represented with an overlaid boxplot and samples are sorted according to their median. (**C**) Correlation of promoter strength of promoter strength obtained from the experiment shown in 5A and published data (http://parts.igem.org/Promoters/Catalog/Anderson) for *V. natriegens* and *E. coli*, respectively. LoglO of the relative promoter strength is plotted and trend line, as well as R^2^ was calculated based on linear regression. Dose response curve of P_tet_ induced by ATc (**D**) and P_trc_ induced by IPTG (**E**). Heatmaps for promoters obtained from the arabinose (**F**) and rhamnose (**G**) operon show growth phase dependency. OD_600_ values displayed on X-Axis indicate the OD_600_ threshold used for the data analysis (see Materials and Methods). OD_600_ = 0.1 was used as the default value in other experiments. The values in the heatmap represent the absolute reporter signal in exponential phase (mean of seven data points around OD_600_ = OD_600_ threshold) divided by the signal of blank wells with LBv2 medium. The median of 12 data points (4 biological replicates from 3 independent experiments) was calculated and plotted in the heat map.

### Characterization of RBS parts

Using ribosome binding site (RBS) sequences from the iGEM Registry of Standard Biological Parts, we tested and characterized eight RBS in our *V. natriegens* chassis. The test constructs were created with the P_tet_ promoter to enable expression of the mScarlet-I reporter gene, while also preventing metabolic burden prior to the start of the experiment, which may have incurred due to leaky basal expression of some of the other promoters. The tested RBS spanned at least five orders of magnitude in translation strength when induced with a moderate ATc concentration (25 ng/mL) (Figure 6A). Two RBS did not result in a detectable signal at this concentration. However, at full induction of P_tet_ a slight signal was indeed visible (Supplementary Figure S2). Therefore, we conclude that these two RBS are functional, but extremely weak and might be suitable for the expression of toxic proteins.

**Figure 6.**
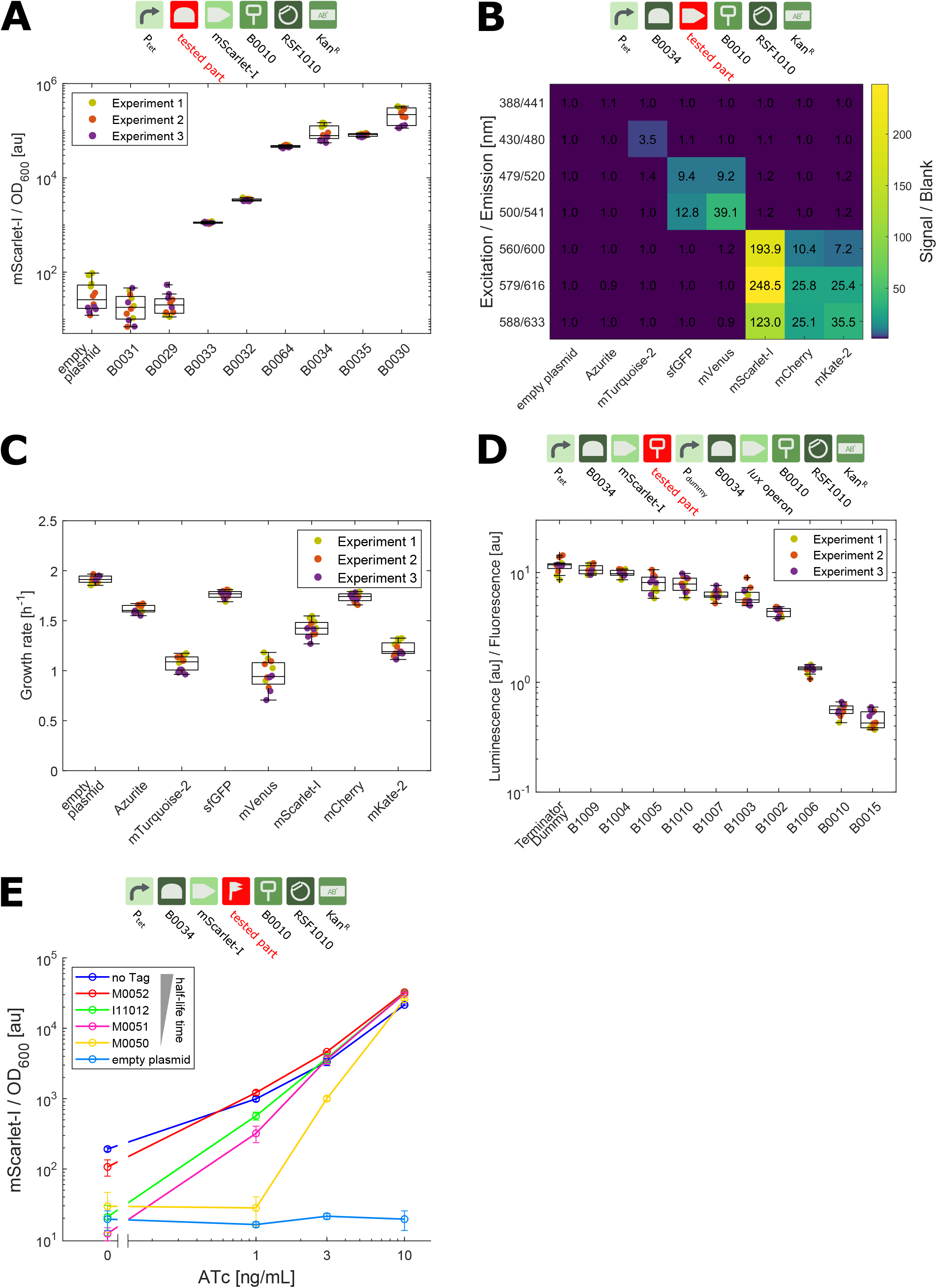
Results for the part categories RBS, CDS, terminator and degradation tags. (**A**) Results of different RBS parts, obtained from a construct with mScarlet-I and P_tet_ as the promoter part. Expression of mScarlet-I was induced by addition of 25 ng/mL ATc during the experiment. Plasmid pDS_108_1R1 kan (Table S5) was used as a dropout plasmid for the RBS characterization. (**B**) Heatmap displaying the results of the characterization of the codon-optimized fluorescent proteins in samples carry the FP driven by P_tet_ and the strong RBS B0034. Samples were induced with 20 ng/mL during the experiment. Values in the heatmap represent the absolute fluorescence signal of the cells in exponential phase divided by the absolute fluorescence signal LBv2 medium. Values represent the median of 12 data points (4 biological replicates from 3 independent experiments). (**C**) Growth rate of strains from the experiment shown in Figure 6B in exponential phase. Plasmid pDS_107_1R1 kan (Table S5) was used as a dropout plasmid for the comparison of fluorescent proteins. (**D**) Characterization of terminator parts. Constructs carry mScarlet-I, transcribed from P_tet_ upstream and a dummy promoter and the *lux* operon downstream of the terminator part. Data points indicate the ratio of luminescence/OD_600_ and mScarlet-I/OD_600_ signal during exponential phase. Samples were induced with 1 ng/mL ATc during the experiment. Plasmid pDS_119_LVL2 kan (Table S5) was used as a dropout plasmid for the terminator characterization. (**E**) Characterization of degradation tags fused to C-terminus of mScarlet-I. Constructs carry P_tet_ to allow for the variable levels of induction. Markers represent the mean of 12 data (4 biological replicates from 3 independent experiments) and were obtained from the steady mScarlet-I/OD_600_ in exponential growth phase. Plasmid pDS_109_1R1 kan (Table S5) was used as a dropout plasmid for the degradation tag characterization.

### Characterization of codon-optimized fluorescent reporter genes

One highlight of the Marburg Collection are eight codon-optimized parts encoding for fluorescent proteins (FP). The characterization of these parts provides two important pieces of information. First, sensitivity of these reporters is indicated by the signal to blank ratio, and second, which of these FPs can be used simultaneously without significant crosstalk. We measured all constructs used in this experiment with different pairs of excitation/emission wavelengths adapted to the optimal wavelengths for each FP. A heatmap shows that the highest signal/blank ratio was obtained for mScarlet-I measured with 579/616 nm (excitation/emission) (Figure 6B). Not surprisingly, we observed high crosstalk between the red FPs mScarlet-I, mCherry and mKate-2, preventing their simultaneous use. The relatively low signal/blank ratio for fluorescent proteins at shorter wavelength (Azurite, mTurquoise-2, sfGFP) is possibly due to the high autofluorescence of the LBv2 medium, which becomes apparent when comparing the spectra of the FP expressing cells with sterile LBv2 medium (Supplementary Figure S3). Interestingly, the expression of the FPs affects growth at different rates, with mTurquoise-2 and mVenus having the strongest detrimental effect on the *V. natriegens* growth rate (Figure 6C). The reason behind this growth defect is currently unknown, but it is possible that inefficient folding of some FPs leads to protein aggregation and concomitant stress responses, or that some FPs are produced at higher copy number, which may pose a higher metabolic burden on the cells. Clearly, further experiments are required to unveil the origin of this growth defect.

### Characterization of terminator parts

We characterized terminator parts by their ability to prevent transcriptional read-through. To this end, we fused P_tet_-mScarlet-I to a terminator of interest and added a promoter-less *lux* operon reporter into the downstream TU to quantify transcriptional read-through. The level 2 plasmid originally contained a terminator dropout part, which was subsequently replaced by the characterized terminator parts. This strategy demonstrates that level 0 dropout parts that are initially included in level 1 plasmids can be carried along into level 2 plasmids. The observed ratio between luminescence (output) and mScarlet-I fluorescence (input) indicates the strength of the terminator, with a low ratio representing a strong terminator. Three of the 10 tested terminators (B0015, B0010, B1006) had a luminescence/fluorescence ratio at least 10-fold lower than a construct containing a terminator dummy part, suggesting a robust transcription termination mediated by these terminators (Figure 6D). The absolute mScarlet-I signal differed only slightly (< 2-fold) between the tested terminators (Supplementary Figure S4), indicating that none of the selected terminators affect the expression of the upstream TU.

### Characterization of degradation tags

The Marburg Collection contains five ssrA-derived protein degradation tags. This tag is naturally encoded in a tmRNA, which is recruited to stalled ribosomes to add 11 amino acids to the nascent polypeptide chain, thereby directing mistranslated proteins to the ClpX/P protease machinery (30,31). Adding the same amino acids to heterologously expressed protein leads to a reduced halflife time and consequently to lower protein levels (32). We characterized these *ssrA* tags by fusing them to the C-terminal end of mScarlet-I, expressed from the inducible P_tet_ promoter. After measuring the steady state fluorescence signal of different induction levels of the ssrA-tagged mScarlet-I, we see the impact of the degradation tag is highest at low ATc concentrations, while no difference between tagged and untagged constructs is visible at high ATc levels (Figure 6E). This is possibly due to a saturation of the protease machinery at high levels of ssrA-tagged proteins, and therefore the inability of the cells to degrade the majority of tagged proteins (33,34). Unexpectedly, the growth rate of the resulting strains is affected by expression of ssrA-tagged proteins compared to expression of non-tagged proteins (Supplementary Figure S5). This might be again due to a saturation of the protease machinery – in line with the finding that a CRISPRi knock-down of protease genes also led to growth reduction, possibly caused by the accumulation of mistranslated/misfolded endogenous proteins (35).

### Characterization of ORI parts

A major advantage of the Marburg Collection is the possibility to freely choose the origin of replication (ORI), allowing for the construction of plasmids for co-transformations or adjusting the gene dosage by selecting appropriate ORI parts with desired plasmid copy numbers. Specifically, we characterized four ORIs belonging to three different ORI incompatibility groups. Two of the ORIs (ColE1, pMB1-M) are derivatives of pMB1 ORIs. In addition, we tested p15A, a medium copy ORI in *E. coli* (36), and the broad-host-range ORI RSF1010 (37). In our experiment we aimed to estimate the copy number, approximated by the expression of a fluorescent reporter gene from a constitutive promoter, to investigate the plasmid stability in cultures without antibiotic selection and, lastly, to study the cell-to-cell variability in their reporter signal, which is indicative of cell-to-cell variability in plasmid copy numbers.

For this purpose, we created a set of plasmids with a constitutively expressed mScarlet-I reporter, differing only in their ORI sequence. After inoculating these cells in fresh medium with and without chloramphenicol selection, we took samples every hour and subjected them to flow cytometry analysis. The mean fluorescent signal differed between the samples, with pMB1-M and RSF1010 being the highest, followed by ColE1 and p15A (Figure 7A). Interestingly, the signal is constant over time in case of RSF1010, consistent with the stringently regulated replication mechanism of RSF1010 via the *repABC* system (38). This system relies on the three plasmid-encoded proteins RepA, RepB and RepC, which act as helicase, primase, and DNA-binding protein, respectively. In contrast, all other ORIs showed an increase in fluorescence signal 1h after the inoculation and a steady decrease thereafter (Figures 7A and 7C), compatible with the more relaxed, RNA based replication mode of these ORIs (39–41). Interestingly, similar experiments with *E. coli* DH5α found RSF1010 to result in a lower plasmid copy number and a higher cell-to-cell variability in reporter signal compared to the other tested ORIs in our experiment (42), but our data shows the opposite trend in *V. natriegens*. This could be due to organism-specific differences in expression rate of the plasmid-encoded regulators controlling plasmid replication.

**Figure 7.**
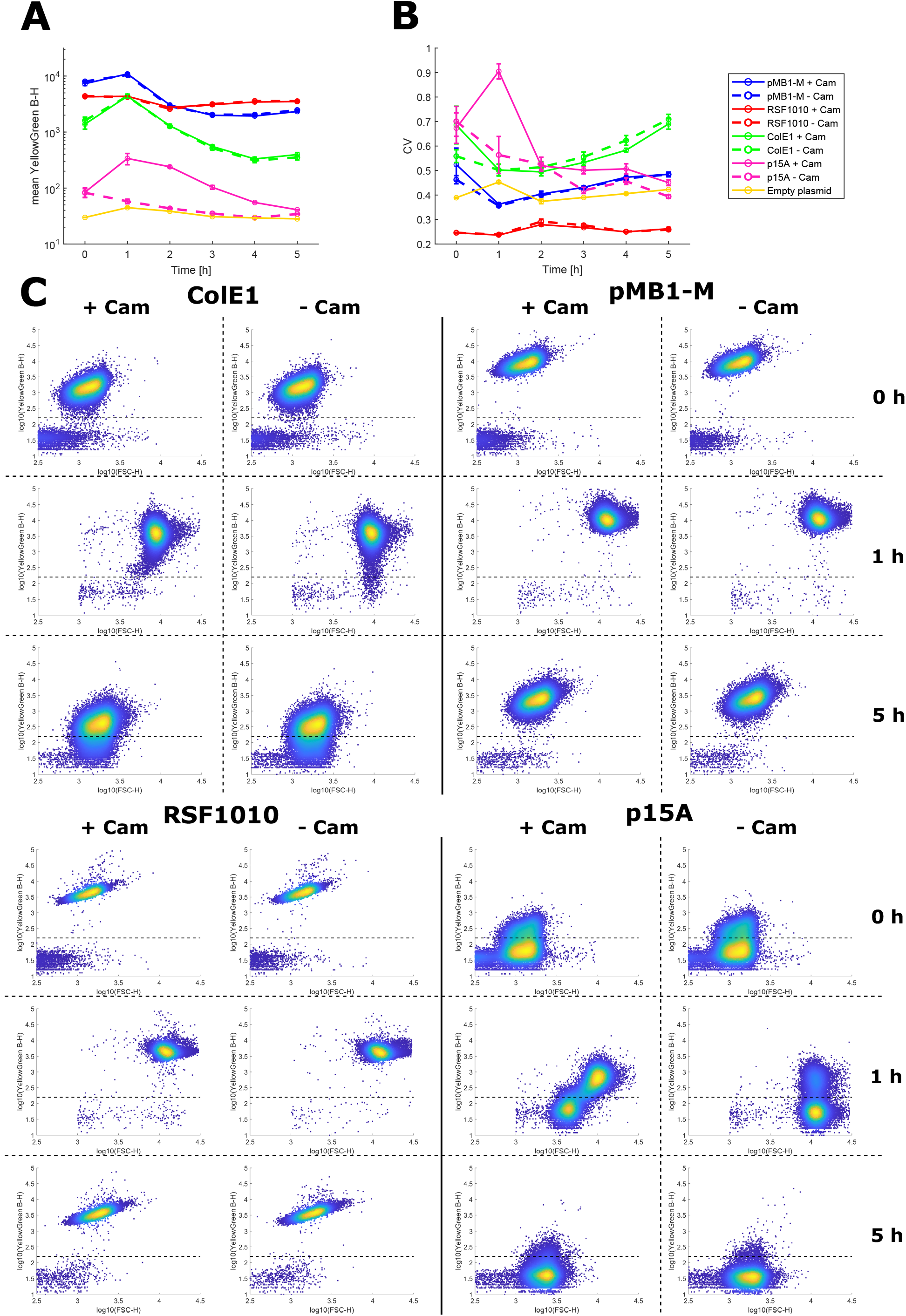
Analysis of ORI parts by flow cytometry. Samples carried plasmids with mScarlet-I driven by a constitutive promoter and differed only in the used ORI part. (**A**) Mean fluorescence signal resulting from different ORIs when samples were grown with chloramphenicol (solid lines) or without antibiotic selection (dashed lines). Each data point represents the mean of the means of all events from four biological replicates grown in two wells each. Error bars indicate the standard deviation between the means of all replicates. (**B**) Coefficient of variation (CV) of all events from one well. Data points indicate the mean CVs from four biological replicates grown in two wells each. Error bars indicate the standard deviation from mean CVs. (**C**) Density plots of representative wells for each ORI at 0 h, 1 h and 5 h grown with and without chloramphenicol selection. Dashed lines set an approximate threshold between mScarlet-I negative and positive events.

For all ORIs, except for p15A, we observed no difference in fluorescence activity for samples with and without antibiotic selection (Figures 7A), suggesting that for the duration of a standard experiment plasmids can be maintained in *V. natriegens* without selection. We note that in this experiment, the plasmids carried a moderately expressed reporter gene; for constructs resulting in strong metabolic burden and consequently high selection pressure this might lead to plasmid loss. To assess cell-to-cell variability in plasmid dopy numbers we calculated the coefficient of variation (CV) for the fluorescence reads. Cell-to-cell variability was highest for p15A (Figure 7B), with an observed bimodal distribution in the density plots (Figure 7C). ColE1 and pMB1-M were moderately variable, but this variation increased over time. RSF1010 had a very low CV, with this variability remaining constant over time (Figure 7B), and a narrow cluster in the density plots (Figure 7C). The homogeneous expression from RSF1010 carrying plasmids is also likely due to the stringent replication control of this ORI (38). From this experiment we conclude that – even in the absence of antibiotic selection – most ORIs are stably maintained in *V. natriegens*. Also, RSF1010 is the ORI that leads to the most homogenous distribution of fluorescence signal in a cell population. The bimodal distribution of p15A copy number is consistent with a recent single-cell report, in which tracking of fluorescently labeled DNA binding proteins targeted to p15A suggested a high cell-to-cell variability and rapid plasmid loss in *V. natriegens* (43). On the other hand, an earlier classical agar plate-based plasmid stability assay indicated a very high stability of this p15A in a population of *V. natriegens* cells (Tschirhart et al., 2019). Together, these observations suggest that although the p15A plasmid is stably maintained in a resistant fraction of the population, stochastic plasmid loss leads to another subpopulation of slow or non-growing cells. This model is consistent with our observation that after dilution to fresh, chloramphenicol-containing medium (t=1h), the non-fluorescent sub-population showed a markedly lower increase in the forward-scatter signal (=smaller cell size) compared to the fluorescent subpopulation (Figure 7C).

### Characterization of AR parts

Finally, we characterized the antibiotic resistance (AR) parts for chloramphenicol, kanamycin and carbenicillin. In early tests we observed that depending on the AR part used, the growth of *V. natriegens* was differently affected when proteins were overexpressed (data not shown). To investigate this effect further, we used the *luxCDABE* cassette, encoding a bacterial luciferase operon as heterologous proteins, given that we found *V. natriegens* to be susceptible toward high expression of this reporter. We induced *lux* operon expression with varying concentrations of ATc, and the resulting metabolic burden of the cells was assessed by cell growth rate. Interestingly, we observed only a slight decrease in growth for the strain carrying the chloramphenicol AR part (Cam^R^) at high ATc levels (Figure 8A). In contrast, strains with a kanamycin AR part (Kan^R^) completely ceased growth at high ATc concentrations. However, codon-optimization of the Kan^R^ CDS for expression in *V. natriegens* allowed cells to cope better with the metabolic burden incurred by *lux* operon expression (Figure 8A). The carbenicillin resistance cassette commonly used in *E. coli* appeared to be only functional in certain plasmid contexts in *V. natriegens* (data not shown). Therefore, we replaced the promoter of the resistance gene with two constitutive promoters of different strength. Carb^R^ under control of the weaker promoter (J23114) also led to growth arrest at high ATc concentrations (strong *lux* operon overexpression) (Figure 8A), like the non-codon optimized Kan^R^. In contrast, with the J23106 promoter (~ 10x stronger than J23114), high expression of the *lux* operon was better tolerated (Figure 8A). Investigation of luminescence signal shows no increase in expression at ATc concentrations above 10 ng/mL (Figure 8B), which is far below the saturating concentration of the promoter (cf. Figure 5C). Thus, it is possible that at high ATc concentrations *V. natriegens* reduces the plasmid copy number to evade toxic levels of metabolic burden on the cell. This decreased copy number and consequently lower gene dose of the AR sequence can possibly be tolerated better in case of potent (Cam^R^) or highly expressed (Kan^R^ - codon optimized, Carb^R^ - strong promoter) resistance cassettes.

**Figure 8.**
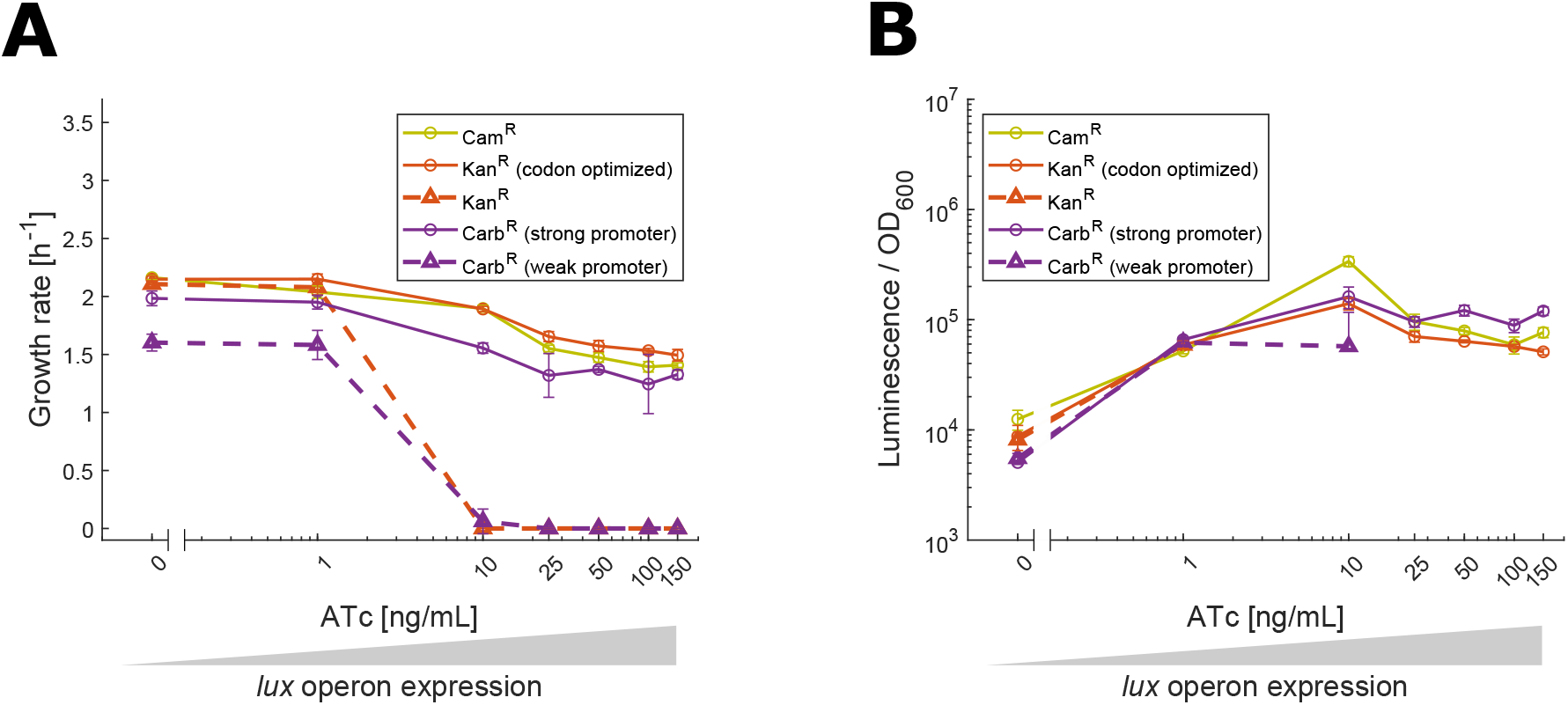
Susceptibility towards metabolic burden for plasmids with different antibiotics. (**A**) Growth rate of strains carrying plasmids with the *lux* operon, driven by P_tet_, differing in the AR part. *lux* operon expression is expected to increase with increasing ATc concentrations. Data points represent mean growth rate calculated from four biological replicates and error bars indicate the standard deviation. (**B**) Luminescence/OD_600_ acquired during the experiment. Data points represent mean luminescence/OD_600_ calculated from four biological replicates and error bars indicate the standard deviation of the mean. Discontinuous lines indicate the absence of noticeable growth of the respective wells.

### Conclusion

We have described a highly flexible cloning toolbox of essential parts for engineering the emerging model organism *V. natriegens*. The hierarchical cloning workflow permits efficient construction of multigene plasmids, taking advantage of *de novo* plasmid assembly with separate ORI and AR parts. We provide novel dropout parts as a highly efficient strategy for the construction of a large number of similar plasmids. Using this approach, we systematically characterized the regulatory parts of the toolbox and thereby provide the growing *V. natriegens* community with a comprehensive collection of genetic parts and a corresponding set of quantitative data.

In our first attempts to characterize the genetic parts of this toolbox, we noticed a high susceptibility of *V. natriegens* toward metabolic burden caused by high expression of reporter genes. This became evident by a high colony-to-colony variation after transformation of constructs with very strong expression of reporter genes and consequently the formation of suppressor mutants (data not shown). We coped with this phenomenon by using tightly regulated inducible promoters, whenever possible, to prevent metabolic burden during strain preparation, while permitting induction and high protein expression in the experiments. Additionally, we always screened multiple colonies from the transformation plate as biological replicates, allowing us to identify colony-to-colony variation in their physiological behavior, which would serve as a red flag indicative for toxic effects of the transformed plasmid. Following these precautionary measures, we were able to generate the robust quantitative data described here.

In summary, we anticipate the Marburg Collection to serve as a valuable resource for *V. natriegens* synthetic biology and the corresponding part characterization as the foundation for rational design of plasmids.

## Materials and Methods

### Strains and growth media

The *V. natriegens* strain used in this study is based on the wildtype strain ATCC14048, acquired through the German Collection of Microorganisms and Cell Cultures GmbH (DSMZ). Unless otherwise indicated, for all experiments we used a derivate of ATCC14048 with a deletion of the *dns* gene. To create the knockout, we replaced the *dns* gene with a chloramphenicol resistance cassette through natural transformation, exactly as described previously (5). Successful replacement was first selected by plating transformed cells on LBv2 agar plates containing chloramphenicol (2 μg/mL), then isolating plasmids, and verifying sequences by Sanger sequencing. After sequence verification, the plasmid required for natural transformation, pMMB-TfoX, was cured by incubating cells in antibiotic free LBv2 with 100 μM IPTG to induce TfoX expression, intentionally inflicting a metabolic burden on the cells to expel the plasmid. The chloramphenicol resistance cassette which replaced the *dns* gene was flanked by Flp/frt sites to allow removal of the resistance marker by expressing the Flp recombinase from the plasmid pBR-flp (44). Finally, loss of chloramphenicol resistance was confirmed by colony PCR and Sanger sequencing. The resulting mutant strain showed a ~3000-fold increased efficiency for plasmid transformation (based on the median CFU/μg plasmid DNA), as well as a higher reproducibility between batches of competent cells (Supplementary Figure S6).

Routine DNA assembly was performed using the commercially available NEB Turbo *E. coli* strain. Strains were grown in LB medium with appropriate antibiotics (kanamycin 50 μg/mL, chloramphenicol (25 μg/mL), carbenicillin (100 μg/mL), and tetracycline (10 μg/mL)). For all experiments involving *V. natriegens* strains, cells were cultivated in LBv2 growth medium containing v2 salts formulated as described in (Weinstock et al., 2016). Unless otherwise indicated, all downstream manipulations with *V. natriegens* ATCC14048 Δ*dns* cells were performed with antibiotic selection (kanamycin 200 μg/mL, chloramphenicol (2 μg/mL), carbenicillin (100 μg/mL)). For incubation on solid medium, 1.5 % agar and the appropriate antibiotic were added to LBv2 medium. Glycerol stocks of *V. natriegens* strains were prepared for long-term storage using the following procedure: cultures were grown for 4-8 h at 37°C and shaken at 220 rpm, then 750 μL aliquots were pelleted at 3000 x g for 5 min. After decanting the supernatant, the pellet was resuspended in 750 μL antibiotic free LBv2 and mixed with 750 μL of 50 % glycerol and stored at −80°C.

### Preparation of chemically competent *V. natriegens* cells and heat-shock transformation of plasmids

A preculture of *V. natriegens* ATCC14048 Δ*dns* was inoculated from a glycerol stock and grown overnight at 37°C and 220 rpm. At the next day 125 mL of preheated LBv2 medium (37°C) was inoculated with the overnight culture to a final OD_600_ 0.01 in a 1 L baffled shake flask. This culture was grown at 200 rpm (37°C) until an OD_600_ between 0.5 and 0.6 was reached. The culture was then transferred to pre-cooled 50 mL falcon tubes and incubated on ice for 10 min, followed by centrifugation for 10 min at 3000 x g at 4°C. The supernatant was discarded, and the pellets were resuspended in 40 mL cold TB buffer per 125 mL bacterial culture (TB buffer: 10 mM Pipes, 15 mM CaCl_2_, 250 mM KCl, pH adjusted to 6.7 with KOH, then add 55 mM MnCl_2_, filter sterilized). The cells were again incubated on ice for 10 min and further centrifuged for 10 min at 3000 x g at 4°C. The supernatant was removed, and pellets were resuspended in 5mL cold TB buffer per 125 mL starting culture and consolidated in a single falcon tube, before adding 350 μL Dimethyl sulfoxide (DMSO). After another 10 min incubation on ice, 50 μL aliquots were prepared in 1.5 mL tubes and snap frozen in liquid nitrogen. Aliquots were stored at −80°C until further use.

Chemically competent *V. natriegens* ATCC14048 Δ*dns* cells were transformed by adding DNA to an aliquot of competent cells, and incubated on ice for 30 min. After 30 min, cells were heat shocked in a water bath or heat block at 42°C for 45 s then immediately incubated on ice for 10 min before recovery. The cells were recovered in 1 mL warm LBv2 medium, shaking at 37°C for 1 h. After recovery, the cells were pelleted by centrifugation at 3000 x g for 1 min, the supernatant was decanted, and the pellet was resuspended in the remaining ~ 50 μL residual medium. The whole volume was plated on 37°C warm LBv2 plates containing the appropriate antibiotic and incubated over night at 37°C.

### Golden Gate Assembly protocol

Golden Gate Assembly reactions in a final volume of 10 μL were set up as follows: 25 fmol of plasmid DNA from each part, 1 μL T4 DNA Ligase Buffer, 0.5 μL T4 DNA Ligase and 0.5 μL of BsaI or Esp3I (BsmBI isoschizomer) restriction enzyme. The concentration of DNA was increased to 50 fmol if transformations did not produce sufficient numbers of colonies. Reactions were incubated in a thermocycler with 30 cycles of 37°C (5 min) and 16°C (10 min), followed by final digestion (60 min, 37°C) and enzyme inactivation (10 min, 80°C) steps. When a dropout part was introduced into a plasmid, the final digestion step was omitted from the cycling conditions. Highly efficient cloning reactions (e.g., creation of level 0 parts, replacing dropout placeholders), were run with a shorter protocol. In this scenario, 30 cycles were performed but incubation times were reduced to 2 min and 5 min for 37°C and 16°C, respectively, and the final digestion step was shortened to 30 min.

### Construction of level 0 and level 0* parts

All details regarding the construction of level 0 and level 0* parts are given in the Supplementary Text. The sequences of all parts are listed in Table S3 and a summary of the cloning process is given for each part in Table S4.

### Characterization of genetic parts

We constructed one dropout plasmid for the characterization of each part type (Table S5) and subsequently cloned the respective level 0 parts into these dropout plasmids. This cloning reaction was transformed directly into *V. natriegens* Δ*dns* by heat shock transformation. For all quantitative experiments, we used four biological replicates. Biological replicates refer to four colonies from the transformation plate of cloning reactions performed in *V. natriegens*. Most experiments were repeated three times, providing up to twelve data points for these experiments. Unless indicated otherwise, we characterized genetic parts with the following microplate reader workflow. First, material from glycerol stocks was resuspended in 50 μL LBv2, and then 5 μL of this suspension was used to inoculate wells of a 96 well plate in a final volume of 100 μL. This plate was incubated for 5 – 6 h at 37°C, which is the equivalent of an overnight preculture in comparable workflows for *E. coli*.

Finally, the preculture was diluted 1:2000 in fresh medium so each plate reader experiment began with a volume of 100 μL per well. Whenever inducers were used in an experiment, they were added to the medium only at the beginning of the experiment, thus were excluded from the precultures. The experiments were performed in a Biotek Synergy H1 micro plate reader. Measurements were taken in 6 min intervals with a 3 min shaking step in double orbital mode and maximum speed occurring between measurements. The OD_600_ was measured in “normal” mode with eight measurements per data point and 100 ms delay after plate movement. Fluorescence was measured with a focal height of 6.5 mm. For all experiments with mScarlet-I as the reporter gene, the excitation and emission wavelength 579 and 616 nm were used, respectively. Excitation and emission wavelengths for the characterization of the codon-optimized FPs are indicated at the Y-axis of the heat map (Figure 6B). Luminescence was measured with an integration time of 1 s and a gain of 150. Experiments were performed without lid/cover to ensure sufficient availability of oxygen and were run for at least 6 h. Experiments involving a luminescent reporter were carried out in black 96 well plates, while all other experiments were performed in transparent 96 well plates.

### Computational analysis of plate reader experiments

Sample data were first normalized by subtracting the mean of four blank wells of the background medium measurements from the sample measurements. The mid of the exponential phase was defined as the first data point with an OD_600_ > 0.1. Three measurements before and after the data point identified as the mid of exponential phase are used to calculate the expression strength. Simply, we calculated the mean of the normalized signal (signal/OD_600_) from all seven data points. Growth rates of samples were derived from the exponential part of the measured growth curve. This section of the curve is confined by the first values with OD_600_ > 0.01 and OD_600_ > 0.18, defining the beginning and end of the exponential phase in LBv2 in the plate reader measurement. A linear fit was derived with the log10 of the OD_600_ values of the respective data points and with the resulting equation we calculated the growth rate of the sample. Data analysis in this manner was conducted for each individual sample well. Whenever growth curves are shown, the data of all wells of one strain or construct were computationally synchronized by aligning all growth curves at the first data point of each well with OD_600_ > 0.01 and the mean of the OD_600_ values of the aligned growth curves is plotted. Error bars indicate the standard deviation from the mean.

### Flow cytometry experiments and data analysis

Four biological replicates for each tested ORI were inoculated in duplicates in a 96 well plate and incubated as a preculture as described above for plate reader experiments. Precultures were grown in LBv2 containing 2 μg/mL chloramphenicol. The precultures were diluted 1:2000 in LBv2 medium containing 2 μg/mL chloramphenicol or no antibiotic in a total volume of 200 μL per well in a 96 well plate. The resulting cultures were incubated in a Biotek Synergy H1 microplate reader with double orbital shaking. Samples were taken in 1 h intervals and diluted in phosphate buffered solution (PBS) to obtain suitable cell densities for the flow cytometry measurement in a BD LSRFortessa with the yellow-green laser line (561 nm). Flow rate was set to lowest possible settings (17.5 μL/s). The resulting raw data were analyzed with a custom Matlab script to gate events and after gating, 30,000 events were used for the subsequent analysis.

## Supporting information

Supplementary Plasmid Maps

## Data availability

Genetic part sequences are available in the Supplemental Materials. Plasmids are available from Addgene. Any other relevant data are available from the corresponding author upon reasonable request.

## Acknowledgements

We are grateful to Dr. Patrick Sobetzko and Prof. Anke Becker for lab space and resources to finish experiments. We thank Prof. Ankur B. Dalia for supplying plasmid pMMB-TfoX. We thank Silvia González Sierra for assistance with flow cytometry experiments. Lastly, we thank Prof. Thorsten Waldminghaus for intense discussions throughout the project.

## Author contributions

D.S., T.H., J.H., R.I. and G.F. conceived the design of the Marburg Collection. D.S., T.H., J.H., B.D. performed experiments. D.S. analyzed the data with the help of G.F. D.S., J.H., F.N., J.N.T and G.F. created figures and wrote the manuscript. G.F. supervised the study.

## Notes

The authors declare no competing financial interest.

## Supplementary Figure legends

**Figure S1. Heatmap for P_tet_ shows no growth phase dependency.** OD_600_ values displayed on X-Axis indicate the OD_600_ threshold used for the data analysis (see Materials and Methods). The values in the heatmap represent the absolute reporter signal in exponential phase (mean of seven data points around OD_600_ = OD_600_ threshold) divided by the signal of blank wells with LBv2 medium. The median of 12 data points (4 biological replicates from 3 independent experiments) was calculated and plotted in the heat map.

**Figure S2. Weak RBS with full induction.** Strains were inoculated from glycerol stock and incubated in glass test tubes in LBv2 with 200 ng/mL ATc overnight. Dilutions of 1:10 of these cultures were subjected to the micro plate reader measurement. The experiment was performed with four biological replicates and the technical measurement was repeated three times.

**Figure S3. Spectra of codon-optimized FPs in comparison to LBv2 and *V. natriegens* control strain.** FP-expressing *V. natriegens* strains were grown overnight in glass test tubes in LBv2 with 2 μg/mL chloramphenicol and 20 ng/μL ATc to induce expression from P_tet_. Of this preculture, 100 μL were transferred into a 96 well plate and then subjected to plate reader measurements. To acquire the excitation spectrum, one wavelength was set for the emission and vice versa. Spectra were measured in 1 nm steps. Wavelength used were as follows (excitation/emission): Azurite (370/470), mTurquoise (410/510), sfGFP (440/520), mVenus (460/580), mScarlet-I (520/650), mCherry (540/650), mKate-2 (540/650).

**Figure S4. Terminator characterization: luminescence and fluorescence values.** Underlying data of Figure 6D. Luminescence and fluorescence values were extracted from the exponential growth phase and normalized by OD_600_ (see Materials and Methods). Samples were induced with 1 ng/mL ATc. Twelve data points are shown for each sample, representing four biological replicates from three independent experiments (shown in different colors). Values for mScarlet-I/OD_600_ are displayed in filled circles and open triangles represent values for luminescence/OD_600_.

**Figure S5. Differences in growth rate between expression of proteins carrying degradation tags.** Growth rate of samples displayed in Figure 6E. Growth rate was calculated as previously described (see Materials and Methods). Data are reported as the mean of 12 data points (4 biological replicates from 3 independent experiments) with standard deviation.

**Figure S6. Transformation efficiency of *V. natriegens* Δ*dns* and wild type.** Plasmid pMC0_8_14_Acam (mScarlet-I) (Vn) was used for the transformation. *V. natriegens* Δ*dns* and wild type (WT) strains were transformed with 1 ng and 300 ng DNA, respectively. Three batches of competent cells were prepared independently and four aliquots each were transformed with the plasmid DNA. For WT, no colonies were obtained for four of the twelve replicates.

## Supplementary Table legends

**Table S1. Overhangs for the creation of level 0 parts.** Overhangs for creating new level 0 parts. Overhangs have to be added to the 5’ end of the primer. Reverse overhangs have to be added as the reverse complement of the indicated sequence. Underlined letters represent BsmBI recognition sites. Bolt letters indicate fusion sites for level 1 assembly. ***N*** bases in connector parts written in bold and italic indicate fusion sites for level 2 assembly that need to be chosen according to the position of the resulting TU in the subsequent cloning step (cf. Fig. 2 in the main text).

**Table S2. Overhangs for the creation of level 0* parts.** Overhangs for creating new level 0* parts. Overhangs have to be added to the 5’ end of the primer. Reverse overhangs have to be added as reverse complement of the indicated sequence. Underlined letters represent BsaI recognition sites. Bolt letters indicate fusion sites for level 1 assembly. ***N*** bases in connector parts written in bold and italic show fusion sites for level 3 assembly and depend on the position of the resulting level 2 plasmid in the subsequent cloning step.

**Table S3. Sequences of the parts in the Marburg Collection.** Each sequence contains all bases of the part, including the fusion sites after digestion with BsaI/BsmBI. The first and last four bases represent the 5’ and 3’ fusion sites, respectively. Parts marked with a hash tag are experimental parts and were not characterized or described in this publication but will be made available through Addgene.

**Table S4. Cloning of LVL0 and LVL0* parts.** Abbreviations used in methods: GGA = Golden Gate Assembly; Gib = Gibson Assembly; Digest = digestion of PCR product with BsaI. Parts marked with a hash tag are experimental parts and were not characterized or described in this publication but will be made available through Addgene.

**Table S5. Construction of plasmids used in this project.** Short nomenclature indicates the level 0 parts used in the assembly (e.g., 1_03 represents pMC0_**1_03**_5C1RLN).

