## Supplementary figures and images for "The Marburg Collection: A Golden Gate DNA Assembly Framework for Synthetic Biology Applications in *Vibrio natriegens*"

### Supplementary Figure 1

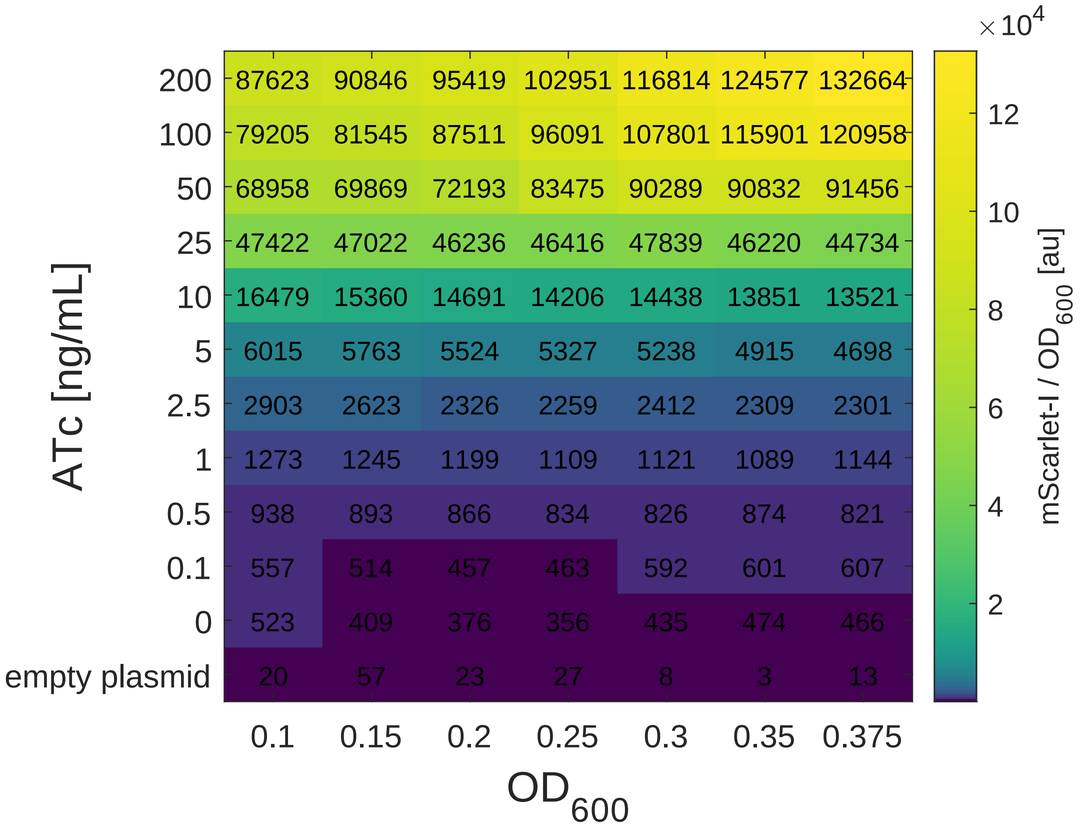

### Supplementary Figure 2

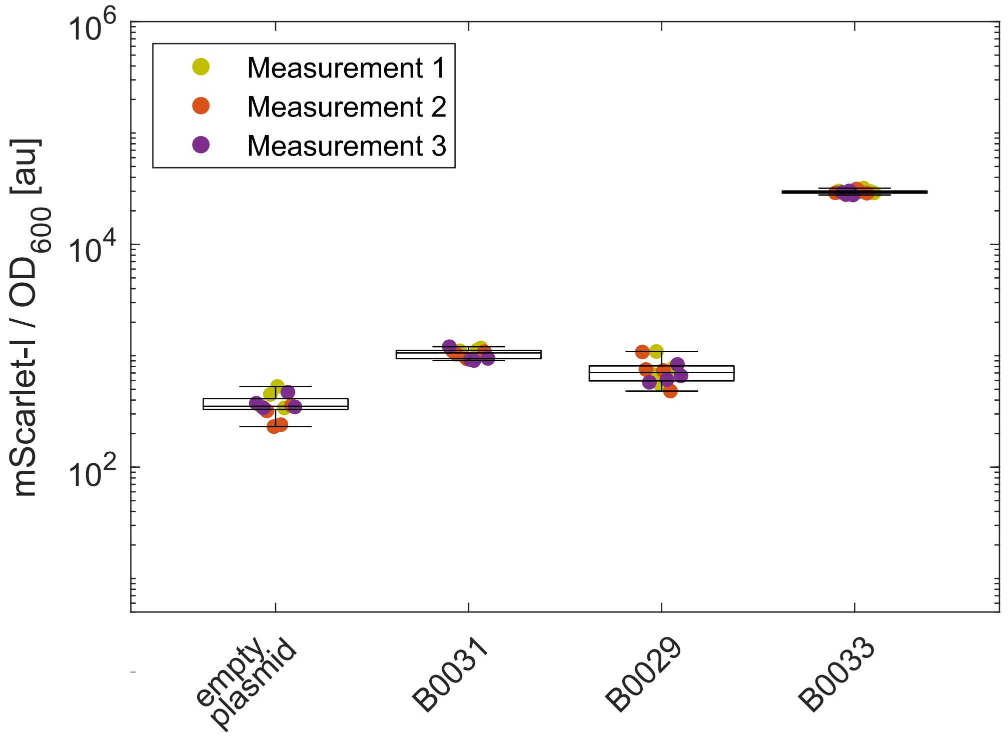

### Supplementary Figure 3

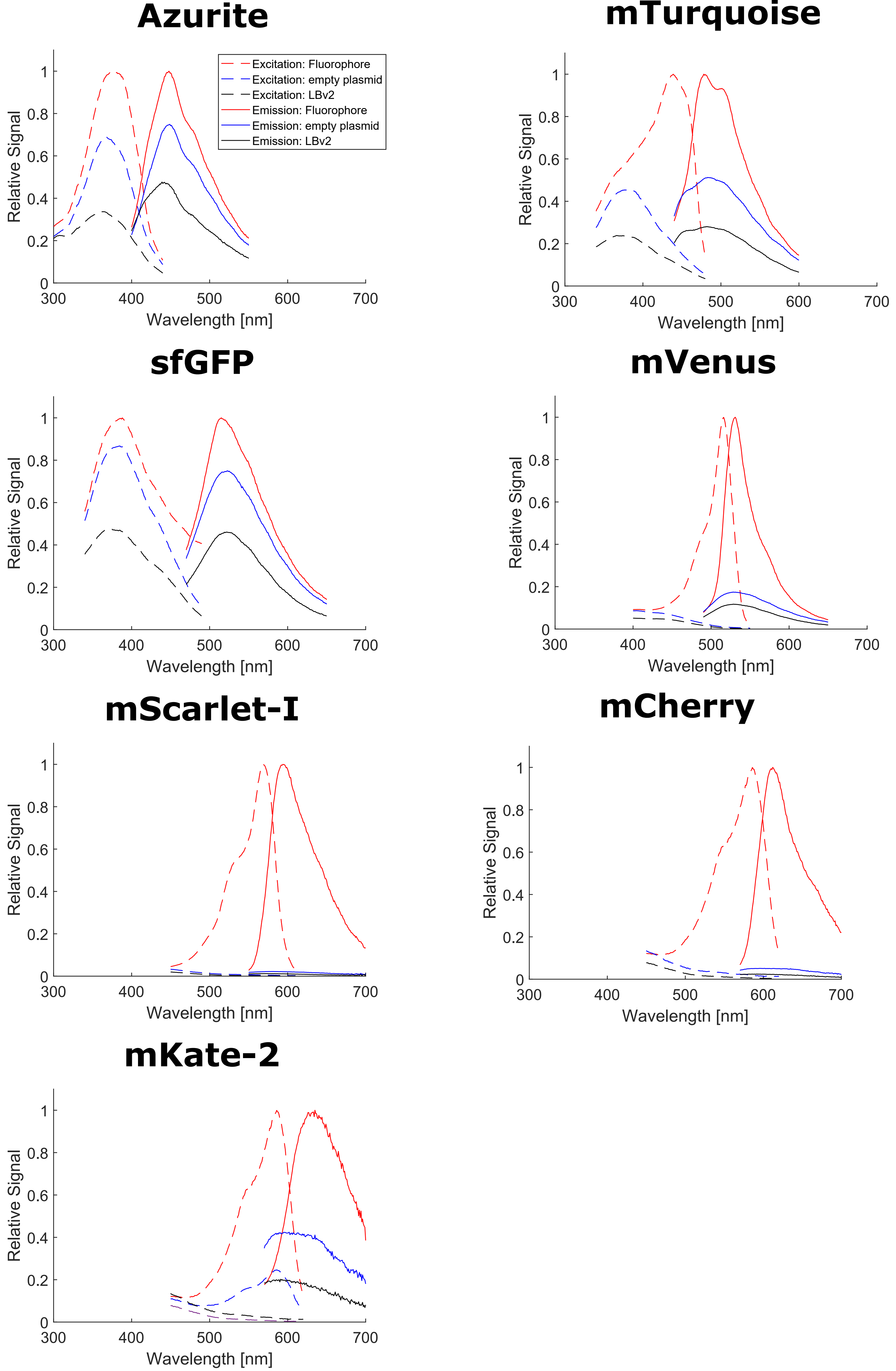

### Supplementary Figure 4

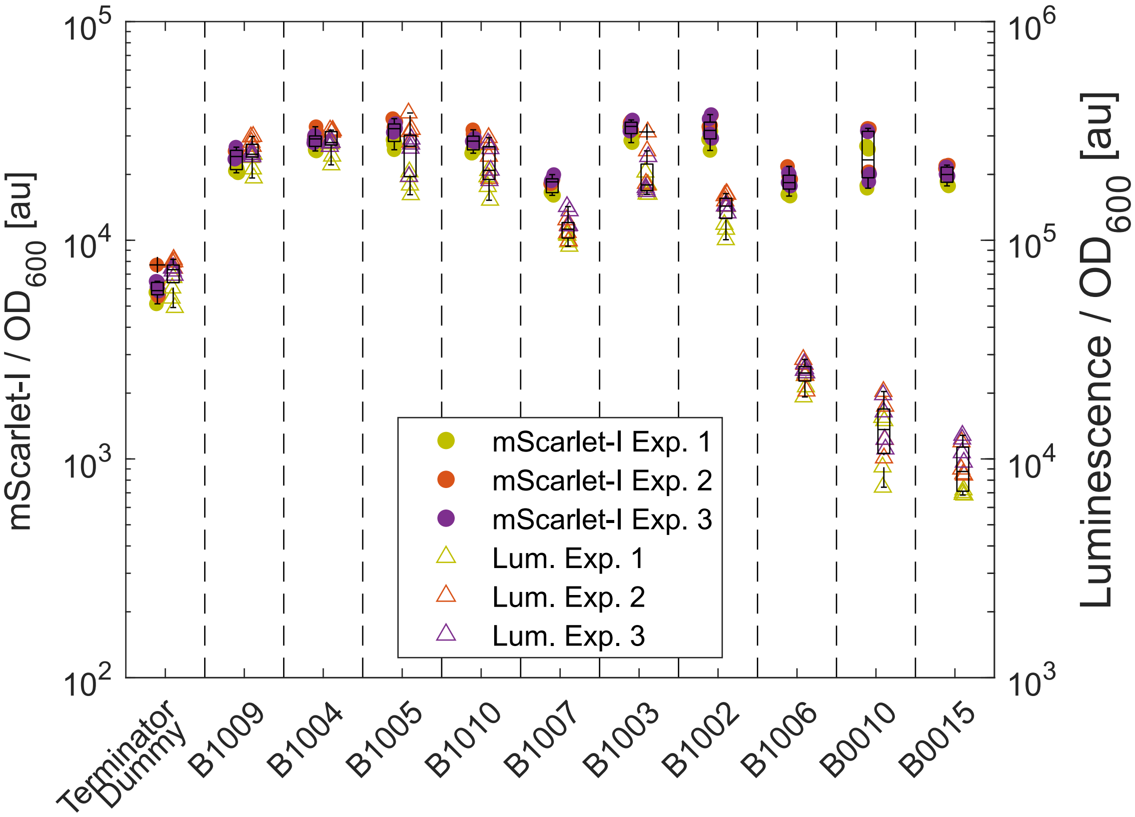

### Supplementary Figure 5

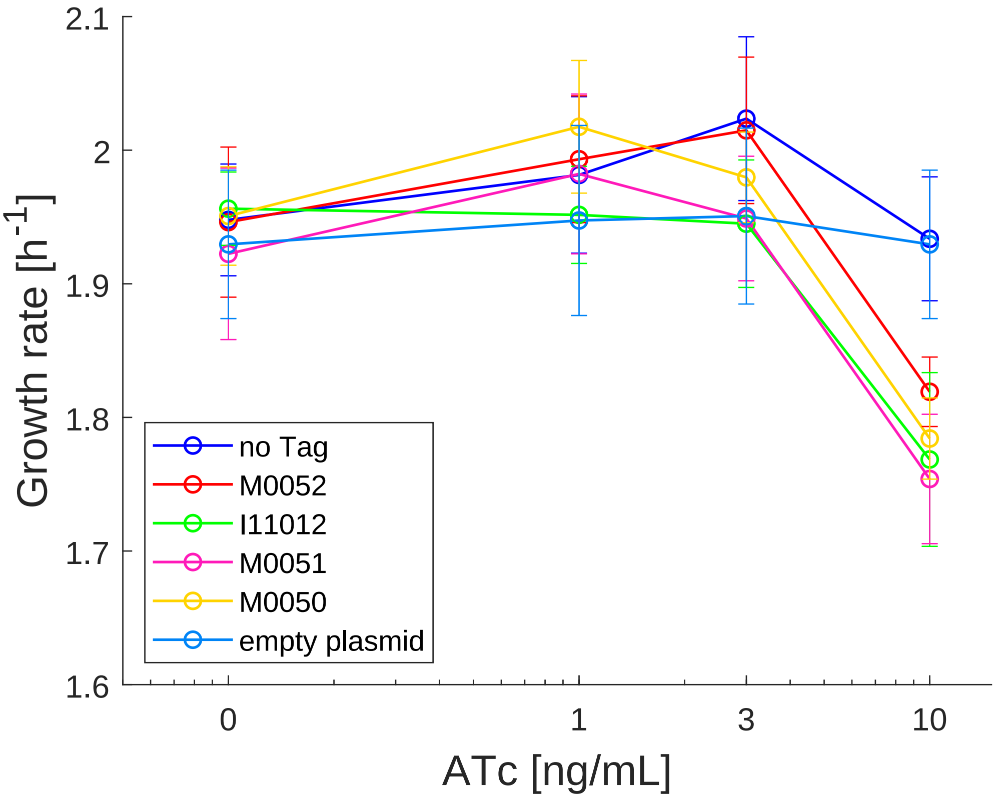

### Supplementary Figure 6

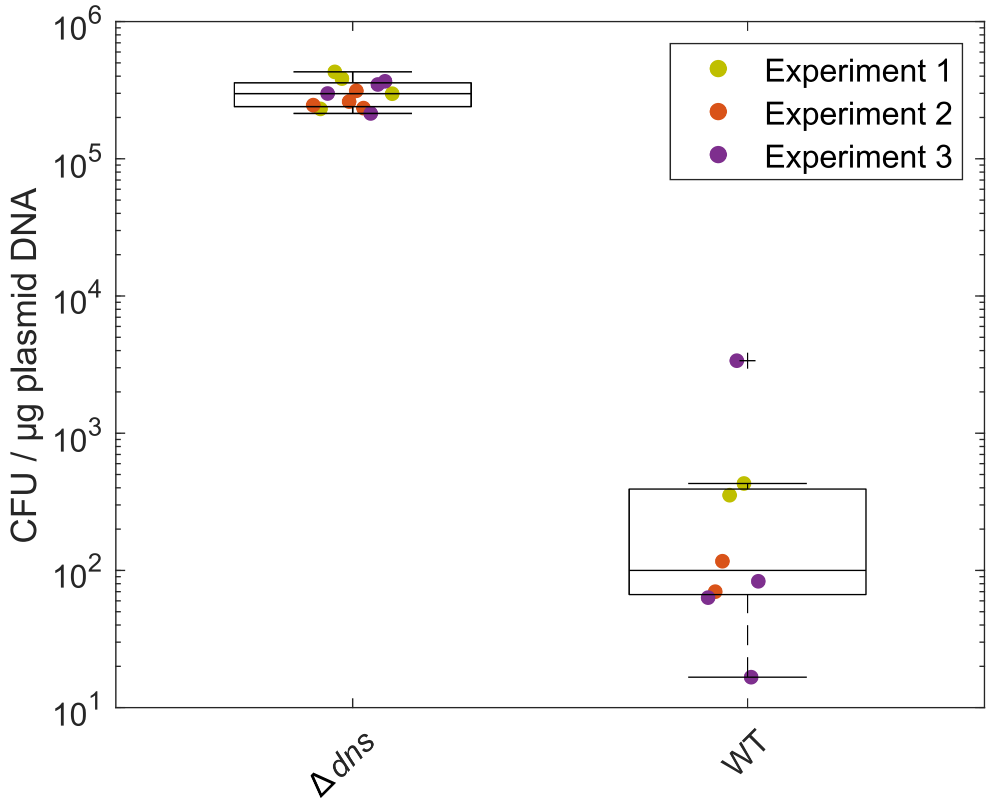
