## Supplementary Text & Tables S1-S5 for "The Marburg Collection: A Golden Gate DNA Assembly Framework for Synthetic Biology Applications in *Vibrio natriegens*"

### Contents

Supplementary text

Supplementary Figure S7 - Generation of new level 0 parts.Supplementary Table S1 - Overhangs for the creation of level 0 parts.

Supplementary Table S2 - Overhangs for the creation of level 0* parts.

Supplementary Table S3 - Sequences of the parts in the Marburg Collection.

Supplementary Table S4 - Cloning of LVL0 and LVL0* parts.

Supplementary Table S5 - Construction of plasmids used in this project.

### Supplementary text

#### Creation of level 0 and level 0* parts

Level 0 parts represent the basic genetic building blocks (e.g., promoter, RBS, CDS), which are stored in entry vectors and are used for the construction of level 1 plasmids. Based on the size of the parts, we suggest different strategies for their creation. For very short parts of < ~ 70 bp, (e.g., constitutive promoters, RBS, and connectors), two complementary oligonucleotides can be ordered which form a short double stranded DNA fragment with a four-base overhang on either side. These overhangs are compatible with the entry vector after its digestion with Type IIs enzyme BsmBI. This strategy represents the cheapest and most convenient way to generate new short parts (Figure S7A). Slightly larger fragments (~ 70 – 150 bp) can be generated by primer extension. In this method two oligonucleotides are used which overlap for ~ 20 bp. The rest of the primers will then be filled up by a DNA polymerase to generate a double stranded DNA sequence. Since this method will generate blunt-end fragments, the restriction site for the Type IIs enzyme has to be included in the primer sequence (Figure S7B). For all fragments larger than 150 bp, a double stranded DNA is required that can either be obtained by standard PCR or by DNA synthesis. Similar to the primer extension approach, restriction sites have to be included in the fragment (Figure S7C).

Parts of all categories can be created by using the same entry vector, as the fusion sites specific to a part category are included during the creation of the respective parts. The only exception are antibiotic resistance (AR) parts, which should be built by using a special AR entry vector. It is generally recommended to switch antibiotics between levels to avoid a high number of false positive colonies. For the construction of a plasmid by Golden Gate Assembly, inevitably one part has to be included, which confers resistance to the same antibiotic used for the selection of the desired plasmid. As there is always a certain fraction of re-ligated or undigested plasmids after the cloning reaction, the AR part should carry a visible marker to distinguish colonies containing the level 0 AR part instead of the correctly assembled plasmid. For this reason, resistance parts are created by using a special AR entry vector. In this case, the fluorescent reporter stays on the plasmid and instead the chloramphenicol resistance marker of the entry vector is replaced by the respective part sequence.

Level 0* parts, which are required for the creation of level 2 plasmids, are built in a similar way as described above with the difference that BsaI restriction enzyme is used instead of BsmBI, and that level 0* part entry vectors have to be used. Level 0 and level 0* parts that have the same fusion sites can also be used to generate level 0 and level 0* parts from the same DNA fragment. Therefore, the DNA fragment has to be predigested with either BsmBI or BsaI and can then be added to the Golden Gate reaction. We used this strategy to create the ORI and AR parts, which have the same sequence in level 0 and level 0*.

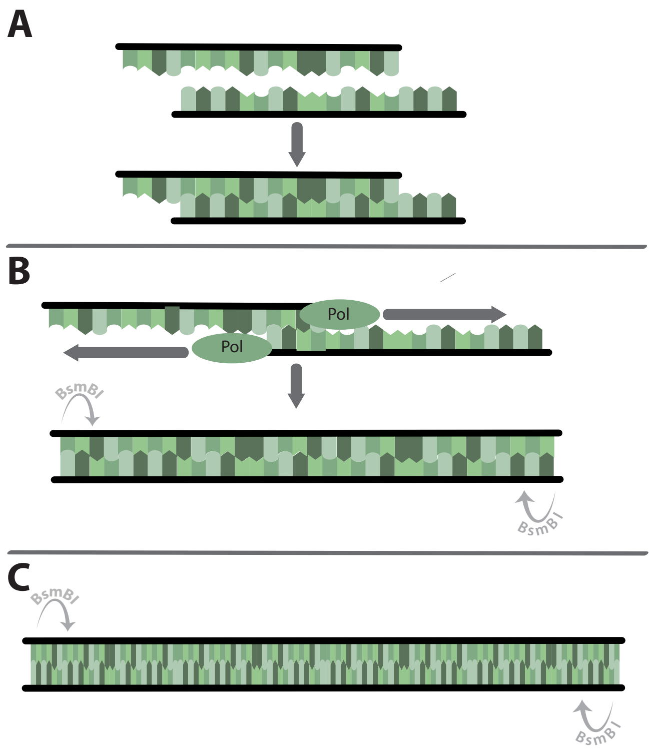


**Figure S7. Generation of new level 0 parts.** Three different approaches for the generation of new level 0 parts are depicted. (**A**) Small parts (<70 bp) were generated by annealing of oligonucleotides. (**B**) Slightly larger parts (70 – 150 bp) were created using primer extension. (**C**) Larger parts are built out of double stranded DNA fragments that were generated either by PCR or by DNA synthesis.

All parts of one category should carry the same fusion sites to ensure compatibility within the toolbox. This also has the advantage that standard overhangs for primers can be used for the creation of new level 0 parts. Tables S1 and S2 provide the standard overhangs for the fragments which are used for the creation of level 0 and level 0* plasmids, respectively. Please note that the reverse complement of the 3' overhang has to be used for the reverse primer.

**Table S1. Overhangs for the creation of level 0 parts.** Overhangs for creating new level 0 parts. Overhangs have to be added to the 5′ end of the primer. Reverse overhangs have to be added as the reverse complement of the indicated sequence. Underlined letters represent BsmBI recognition sites. Bolt letters indicate fusion sites for level 1 assembly. ***N*** bases in connector parts written in bold and italic indicate fusion sites for level 2 assembly that need to be chosen according to the position of the resulting TU in the subsequent cloning step (cf. Fig. 2 in the main text).

| Part Category | 5ʻ overhang | 3ʻ overhang |
|---|---|---|
| 1<br>5′ Connector | AACGTCTCGCTCG**AACA**CGTCTCG***NNNN*** | **GGAG**TGAGGGAGACGAA |
| 2<br>Promoter | AACGTCTCGCTCG**GGAG** | **TACT**TGAGGGAGACGAA |
| 3<br>RBS | AACGTCTCGCTCG**TACT**AGAG | TAATC**AATG**TGAGGGAGACGAA |
| 4<br>CDS | AACGTCTCGCTCG**AATG** | **GCTT**TGAGGGAGACGAA |
| 5<br>Terminator | AACGTCTCGCTCG**GCTT**AA | **CGCT**TGAGGGAGACGAA |
| 6<br>3′ Connector | AACGTCTCGCTCG**CGCT** | ***NNNN***GGAGACG**AGCT**TGAGGGAGACGAA |
| 7<br>ORI | AACGTCTCGCTCG**AGCT** | **TGCT**TGAGGGAGACGAA |
| 8<br>AR | AACGTCTCGCTCGT**TGCT** | **AACA**TGAGGGAGACGAA |
| 4a<br>N-Tag | AACGTCTCGCTCG**AATG** | GG**GATG**TGAGGGAGACGAA |
| 4b<br>N-tagged *CDS* | AACGTCTCGCTCG**GATG** | **GCTT**TGAGGGAGACGAA |
| 5a<br>C-Tag | AACGTCTCGCTCG**GCTT**TA | G**GGTA**TGAGGGAGACGAA |
| 5b<br>Terminator | AACGTCTCGCTCG**GGTA**A | **CGCT**TGAGGGAGACGAA |

**Table S2. Overhangs for the creation of level 0* parts.** Overhangs for creating new level 0* parts. Overhangs have to be added to the 5′ end of the primer. Reverse overhangs have to be added as reverse complement of the indicated sequence. Underlined letters represent BsaI recognition sites. Bolt letters indicate fusion sites for level 1 assembly. ***N*** bases in connector parts written in bold and italic show fusion sites for level 3 assembly and depend on the position of the resulting level 2 plasmid in the subsequent cloning step.

| Part Category | 5ʻ overhang | 3ʻ overhang |
|---|---|---|
| 1*<br>5′ Connector | AAGGTCTCGCTCG**AACA**GGTCTCG***NNNN*** | **GGAG**TGAGGGAGACCAA |
| 6*<br>3′ Connector | AAGGTCTCGCTCG**CGCT** | ***NNNN***GGAGACC**AGCT**TGAGGGAGACCAA |

| | | |
|---|---|---|
| 7*<br>ORI | AAGGTCTCGCTCG**AGCT** | **TGCT**TGAGGGAGACCAA |
| 8*<br>AR | AAGGTCTCGCTCGT**TGCT** | **AACA**TGAGGGAGACCAA |

Apart from the fusion sites, some additional bases were introduced for some part categories. The RBS parts carry some bases upstream and downstream to ensure the correct spacing to the promoter and the start codon of the CDS part. All parts which are part of the translated sequence carry additional bases to not disrupt the reading frame. The included bases were chosen to result in small amino acids. To allow for C-terminal tagging of proteins, the CDS part must not carry a stop codon. By default, the protein will be extended by alanine before translation will be stopped by the stop codon, which is located at the beginning of each terminator part. If the extension by one amino acid cannot be tolerated in a specific case, a stop codon can be included at the end of the CDS part. However, this will prevent applying C-terminal tags in the future. Please note that the fusion site between RBS and CDS already contains the start codon ATG, therefore the start codon of the CDS sequence should be omitted.

The additionally introduced bases should be seen as a recommendation for the future construction of novel level 0 parts rather than a strict requirement. It is only necessary to maintain the 4 bp fusion sites in order the ensure compatibility within the Marburg Collection. Another possibility is to further split part categories, by introducing additional fusion sites to achieve a higher degree of modularity. For example, one could split the promoter part in case of inducible promoters by separating the regulatory protein and the actual promoter with the respective operator sites. Likewise, parts spanning multiple categories could be designed. As an example, if promoters and RBS are to be used always in a fixed combination, the complexity of the assembly can be lowered by reducing the number of required fragments by using a "promoter-RBS" part instead of using both a promoter and an RBS part.

**Table S3. Sequences of the parts in the Marburg Collection**. Each sequence contains all bases of the part, including the fusion sites after digestion with BsaI/BsmBI. The first and last four bases represent the 5' and 3' fusion sites, respectively. Parts marked with a hash tag are experimental parts and were not characterized or described in this publication but will be made available through Addgene.

| Part | Sequence |
|---|---|
| **pMC0_1-6_01_1-6 Dropout** | AACAAGAGACCGAAAGTGAAACGTGATTTCATGCGTCATTTTGAACATTTTGTAAATCTTATTTAATAATGTGTGCGGCAATTCACATTTAATTTAT GAATGTTTTCTTAACATCGCGGCAACTCAAGAAACGGCAGGTTCGGATCTTAGCTACTAGAGAAAGAGGAGAAATACTAGATGCGTAAAGGCGA AGAACTTTTCACTGGCGTAGTACCAATTTTAGTGGAGCTGGACGGAGATGTAAATGGTCATAAGTTTTCAGTTCGAGGAGAAGGCGAAGGAGAT GCAACTAACGGTAAGCTGACACTAAAGTTTATCTGTACCACGGGTAAACTGCCGGTCCCATGGCCGACACTGGTTACTACACTTACTTATGGTGT ACAATGCTTTGCTCGTTATCCTGACCATATGAAGCAGCACGATTTCTTCAAATCGGCCATGCCTGAAGGATACGTGCAAGAACGTACAATTAGCTT TAAAGATGACGGCACCTATAAAACGCGCGCCGAAGTTAAATTCGAGGGTGACACATTGGTTAATCGTATAGAACTTAAGGGCATTGACTTCAAA GAAGATGGCAACATCCTTGGCCATAAACTGGAATATAATTTTAACTCTCACAATGTCTACATTACGGCGGATAAACAAAAGAATGGCATTAAAGC GAATTTCAAAATCCGCCACAATGTCGAAGACGGCTCGGTTCAACTGGCGGACCATTATCAGCAGAACACCCCAATCGGTGATGGCCCGGTTCTGT TACCTGATAATCATTACCTTTCTACTCAAAGCGTTTTATCTAAAGATCCTAACGAGAAGCGTGATCATATGGTTCTACTGGAATTTGTTACCGCAGC TGGTATCACGCACGGCATGGATGAGCTGTATAAATGACCAGGCATCAAATAAAACGAAAGGCTCAGTCGAAAGACTGGGCCTTTCGTTTTATCT GTTGTTTGTCGGTGAACGCTCTCTACTAGAGTCACACTGGCTCACCTTCGGGTGGGCCTTTCTGCGTTTATAGGTCTCTAGCT |

| | |
|---|---|
| **pMC0_1-6_02_1-6 Connector** | AACATATCCGTGAGAGCT |
| **pMC0_1-6_03_1-6 Dropout mScarlet-I (Vn)** | AACAAGAGACCGAAAGTGAAACGTGATTTCATGCGTCATTTTGAACATTTTGTAAATCTTATTTAATAATGTGTGCGGCAATTCACATTTAATTTAT<br>GAATGTTTTCTTAACATCGCGGCAACTCAAGAAACGGCAGGTTCGGATCTTAGCTACTAGAGAAAGAGGAGAAATACTAGATGGTTTCTAAAGG<br>TGAAGCAGTGATCAAAGAATTTATGCGCTTCAAAGTTCACATGGAAGGTTCTATGAACGGCCACGAATTTGAAATTGAAGGTGAAGGCGAAGGT<br>CGTCCATACGAAGGTACTCAAACCGCAAAACTGAAAGTTACCAAAGGTGGTCCACTGCCATTTTCTTGGGATATTCTGTCTCCACAATTTATGTAC<br>GGTTCTCGTGCATTTATCAAACACCCAGCAGATATTCCAGACTACTACAAACAATCTTTTCCGGAAGGTTTCAAATGGGAACGTGTTATGAATTTT<br>GAAGATGGTGGTGCAGTTACGGTTACCCAAGATACCTCTCTGGAAGATGGTACTCTGATCTACAAAGTTAAACTGCGTGGTACTAACTTTCCACC<br>AGATGGTCCAGTTATGCAGAAAAAAACCATGGGTTGGGAAGCATCTACCGAACGTCTGTACCCAGAAGATGGCGTTCTGAAAGGTGATATCAAA<br>ATGGCACTGCGTCTGAAAGATGGCGGTCGTTACCTGGCAGATTTCAAAACCACCTACAAAGCGAAAAAACCAGTTCAAATGCCAGGTGCATACA<br>ACGTTGATCGTAAACTGGATATTACCAGCCACAACGAAGATTACACCGTTGTTGAACAATACGAACGTTCTGAAGGCCGTCACTCTACCGGTGGT<br>ATGGATGAACTGTACAAATAACCAGGCATCAAATAAAACGAAAGGCTCAGTCGAAAGACTGGGCCTTTCGTTTTATCTGTTGTTTGTCGGTGAAC<br>GCTCTCTACTAGAGTCACACTGGCTCACCTTCGGGTGGGCCTTTCTGCGTTTATAGGTCTCTAGCT |
| **pMC0_1_01_1 Dropout** | AACAAGAGACCGAAAGTGAAACGTGATTTCATGCGTCATTTTGAACATTTTGTAAATCTTATTTAATAATGTGTGCGGCAATTCACATTTAATTTAT<br>GAATGTTTTCTTAACATCGCGGCAACTCAAGAAACGGCAGGTTCGGATCTTAGCTACTAGAGAAAGAGGAGAAATACTAGATGCGTAAAGGCGA<br>AGAACTTTTCACTGGCGTAGTACCAATTTTAGTGGAGCTGGACGGAGATGTAAATGGTCATAAGTTTTCAGTTCGAGGAGAAGGCGAAGGAGAT<br>GCAACTAACGGTAAGCTGACACTAAAGTTTATCTGTACCACGGGTAAACTGCCGGTCCCATGGCCGACACTGGTTACTACACTTACTTATGGTGT<br>ACAATGCTTTGCTCGTTATCCTGACCATATGAAGCAGCACGATTTCTTCAAATCGGCCATGCCTGAAGGATACGTGCAAGAACGTACAATTAGCTT<br>TAAAGATGACGGCACCTATAAAACGCGCGCCGAAGTTAAATTCGAGGGTGACACATTGGTTAATCGTATAGAACTTAAGGGCATTGACTTCAAA<br>GAAGATGGCAACATCCTTGGCCATAAACTGGAATATAATTTTAACTCTCACAATGTCTACATTACGGCGGATAAACAAAAGAATGGCATTAAAGC<br>GAATTTCAAAATCCGCCACAATGTCGAAGACGGCTCGGTTCAACTGGCGGACCATTATCAGCAGAACACCCCAATCGGTGATGGCCCGGTTCTGT<br>TACCTGATAATCATTACCTTTCTACTCAAAGCGTTTTATCTAAAGATCCTAACGAGAAGCGTGATCATATGGTTCTACTGGAATTTGTTACCGCAGC<br>TGGTATCACGCACGGCATGGATGAGCTGTATAAATGACCAGGCATCAAATAAAACGAAAGGCTCAGTCGAAAGACTGGGCCTTTCGTTTTATCT<br>GTTGTTTGTCGGTGAACGCTCTCTACTAGAGTCACACTGGCTCACCTTCGGGTGGGCCTTTCTGCGTTTATAGGTCTCTGGAG |
| **pMC0_1_02_5C1CLF** | AACACGTCTCGGGAGGGACCAAAACGAAAAAAGACGCTTTTCAGCGTCTTATTGTTCGTCTTTGGTACCGAGCTGTGTCTGTGGCTGAGAGAGC<br>AGGTACAAGATAAACCCGTCTATATTCTCAAATTCTTTCACTATCGCACTTGTTTTATGTGAAGTCCAGATATGGACTCTGGAACCCAACTCGGAG<br>CTCTGCGGGTAGAGCGTCAACCATTCGAACGTTGCGCGGTCTTCGGAGAACCGTAAGAGGTAATATACTCCGATATTCATTTCAGCGGTAGCTGA<br>GTCCGTCAGTTGATACGGTGCGTACCCGGAGCAAATAGGAGAGGCGGTTGTAGTGGGGGGAATGGATAGCAAGCTGCGGGTAGACCTCCAATT<br>ATTGAAGGCCTCCCAAATCGGGGGGCCTTTTTTATTGATAACAAAAGGAG |
| **pMC0_1_03_5C1RLF** | AACACGTCTCGAACAGGACCAAAACGAAAAAAGACGCTTTTCAGCGTCTTATTGTTCGTCTTTGGTACCGAGCTGTGTCTGTGGCTGAGAGAGCA<br>GGTACAAGATAAACCCGTCTATATTCTCAAATTCTTTCACTATCGCACTTGTTTTATGTGAAGTCCAGATATGGACTCTGGAACCCAACTCGGAGC<br>TCTGCGGGTAGAGCGTCAACCATTCGAACGTTGCGCGGTCTTCGGAGAACCGTAAGAGGTAATATACTCCGATATTCATTTCAGCGGTAGCTGA<br>GTCCGTCAGTTGATACGGTGCGTACCCGGAGCAAATAGGAGAGGCGGTTGTAGTGGGGGGAATGGATAGCAAGCTGCGGGTAGACCTCCAATT<br>ATTGAAGGCCTCCCAAATCGGGGGGCCTTTTTTATTGATAACAAAAGGAG |
| **pMC0_1_04_5C1CSF** | AACACGTCTCGGGAGGGAG |
| **pMC0_1_05_5C1RSF** | AACACGTCTCGAACAGGAG |
| **pMC0_1_06_5C1SR** | AACACGTCTCGAGTAGGAG |
| **pMC0_1_07_5C2LN[#]** | AACACGTCTCGTACTGGACCAAAACGAAAAAAGGCCGCTTTCGCGGCCTCTTTTCTGGAATTTGGTACCGAGTCGACACCCTACCGTATCGCTGT<br>GGCAGGCACAGGATCTAACCGGTCCTGGACCGGCGTAAGTTGCTGTACAGCCAAATCTTCCGTACTAATTCTACAGGATTTGTCAACTGCAAACG<br>AATTTCTAGTCGGAACGGATCACGTGCACCGGGGTATGCCGTGGTTGAATATCGGCACATGAGGGGCTCGTGGGACACTTCCAGAACTGTCTGG<br>CGTGCGAAGTCTCTGCTAAATGCAAAACAGTCAGGACTCGTTACAGCTCGTGTATGGGAGCTCCGGGTAGGCCGATCGTCAATTGCGGCTCGGT<br>ACCAAATTTTCGAAAAAAGACGCTGAAAAGCGTCTTTTTTCGTTTTGGTCCGGAG |
| **pMC0_1_08_5C2SF** | AACACGTCTCGTACTGGAG |
| **pMC0_1_09_5C2SR** | AACACGTCTCGCATTGGAG |
| **pMC0_1_10_5C3LF[#]** | AACACGTCTCGAATGGGACCAAAACGAAAAAAGACGCTTTTCAGCGTCTTCAATAATTGGTTTGGTACCGAGATTCGTTGTTGACTTCATCGTCAC<br>GATAAAGCACGACAAGTCCCAGCCTGTGAGTGCGACCCGAAATTGTAGGTAGACACTCACGCTCCCCCCGATATCAGACAGCGTGCTAATGAAA<br>GCCTTCCGATCTTCGGATAAGGCCCAAGCTCAGCACATGAACGGATCAGTGGTAGGTGTCTCACAACCCAATCGTGGTGACAATACGAATGTCCA<br>AATATTGCCTCTTGGCGGGTAGATTCCGTTACACTTTACAGCCATCATAACAACCAGATGAGAATCGATGGTATTCTTGGACCCGCGCCAATTATT<br>GAAGGCCTCCCAAATCGGGGGGCCTTTTTTATTGATAACAAAAGGAG |
| **pMC0_1_11_5C3SF** | AACACGTCTCGAATGGGAG |
| **pMC0_1_12_5C3SR** | AACACGTCTCGAAGCGGAG |
| **pMC0_1_13_5C4LF[#]** | AACACGTCTCGGCTTGGACCAAAACGAAAAAAGACGCTTTTCAGCGTCTTTTTAGTTAGATTTGGTACCGAGTTTATGGGCACGCTTCCGCGTTAA<br>TCATACCTCTATTCACACGGTCCCCCACCAGTGTGTCTAAAGACGTACCCATTGAACGCGGGATTGGTTGTGATAGCTGGTTGTGCCACTAAGTCC<br>TAGTCAAACGAGGCTGGCATCCCATAGCAGCAAAATCGGGTAGTAACCCTCAAAGACGTGCCCGACAGTTCATCTTCCTGTAAGAACTATTGGCT<br>TACGCACGAGTCTGTTCGTGTATCGTCTAGCGGTTAGATTCACCTGAATTGCTATGCTTCTTTCAATGGATCCTGCCCACTTCACTTTTCGAAAAAA<br>GGCCTCCCAAATCGGGGGGCCTTTTTTATTGATAACAAAAGGAG |
| **pMC0_1_14_5C4SF** | AACACGTCTCGGCTTGGAG |
| **pMC0_1_15_5C4SR** | AACACGTCTCGTACCGGAG |
| **pMC0_1_16_5C5LF[#]** | AACACGTCTCGGGTATTTTGTTATCAATAAAAAAGGCCCCCCGATTTGGGAGGCCTTTTTTTTTTTTGGTGGTCGCATGAAGCAGGCCTAGTATTC<br>CAACGTGGTGGCCAATTAACAATATTCCAACCCCAATCTGTAAAAAACTGTGCCCTACGCAGGACTCAGAGAACGGCGGAAGGTAGGTACTCGT<br>CTTCTAACCCGACGCATTAAGGCTACAGGGTGCGAAGGCCATTATTGCAAACTTTTTGTTCTCAACAACGTAAATCTACCGCACTTCAGGTGTCGT<br>TCTTGATGTACGAAGACGGTAGGCCTATGGATTGCCTTCTGTATCAGCAATTTTTCATCTCCTTCCAAGCGGCATCCATAATCTAACTAAAAAGGC<br>CTCCCAAATCGGGGGGCCTTTTTTATTGATAACAAAAGGAG |
| **pMC0_1_17_5C5SF** | AACACGTCTCGGGTAGGAG |
| **pMC0_1_18_5C5CSR** | AACACGTCTCGAGCGGGAG |
| **pMC0_1_19_5C5OS R** | AACACGTCTCGAGCTGGAG |
| **pMC0_1_20_5C0SF[#]** | AACAGAATTCGCGGCCGCTTCTAGAGGGAG |
| **pMC0_1_21_1 Dropout mScarlet-I (Vn)** | AACAAGAGACCGAAAGTGAAACGTGATTTCATGCGTCATTTTGAACATTTTGTAAATCTTATTTAATAATGTGTGCGGCAATTCACATTTAATTTAT<br>GAATGTTTTCTTAACATCGCGGCAACTCAAGAAACGGCAGGTTCGGATCTTAGCTACTAGAGAAAGAGGAGAAATACTAGATGGTTTCTAAAGG<br>TGAAGCAGTGATCAAAGAATTTATGCGCTTCAAAGTTCACATGGAAGGTTCTATGAACGGCCACGAATTTGAAATTGAAGGTGAAGGCGAAGGT<br>CGTCCATACGAAGGTACTCAAACCGCAAAACTGAAAGTTACCAAAGGTGGTCCACTGCCATTTTCTTGGGATATTCTGTCTCCACAATTTATGTAC<br>GGTTCTCGTGCATTTATCAAACACCCAGCAGATATTCCAGACTACTACAAACAATCTTTTCCGGAAGGTTTCAAATGGGAACGTGTTATGAATTTT<br>GAAGATGGTGGTGCAGTTACGGTTACCCAAGATACCTCTCTGGAAGATGGTACTCTGATCTACAAAGTTAAACTGCGTGGTACTAACTTTCCACC<br>AGATGGTCCAGTTATGCAGAAAAAAACCATGGGTTGGGAAGCATCTACCGAACGTCTGTACCCAGAAGATGGCGTTCTGAAAGGTGATATCAAA<br>ATGGCACTGCGTCTGAAAGATGGCGGTCGTTACCTGGCAGATTTCAAAACCACCTACAAAGCGAAAAAACCAGTTCAAATGCCAGGTGCATACA<br>ACGTTGATCGTAAACTGGATATTACCAGCCACAACGAAGATTACACCGTTGTTGAACAATACGAACGTTCTGAAGGCCGTCACTCTACCGGTGGT<br>ATGGATGAACTGTACAAATAACCAGGCATCAAATAAAACGAAAGGCTCAGTCGAAAGACTGGGCCTTTCGTTTTATCTGTTGTTTGTCGGTGAAC<br>GCTCTCTACTAGAGTCACACTGGCTCACCTTCGGGTGGGCCTTTCTGCGTTTATAGGTCTCTGGAG |
| **pMC0_2-5_01_2-5 Dropout** | GGAGAGAGACCGAAAGTGAAACGTGATTTCATGCGTCATTTTGAACATTTTGTAAATCTTATTTAATAATGTGTGCGGCAATTCACATTTAATTTA<br>TGAATGTTTTCTTAACATCGCGGCAACTCAAGAAACGGCAGGTTCGGATCTTAGCTACTAGAGAAAGAGGAGAAATACTAGATGCGTAAAGGCG<br>AAGAACTTTTCACTGGCGTAGTACCAATTTTAGTGGAGCTGGACGGAGATGTAAATGGTCATAAGTTTTCAGTTCGAGGAGAAGGCGAAGGAGA |

| | |
|---|---|
| | TGCAACTAACGGTAAGCTGACACTAAAGTTTATCTGTACCACGGGTAAACTGCCGGTCCCATGGCCGACACTGGTTACTACACTTACTTATGGTGTACAATGCTTTGCTCGTTATCCTGACCATATGAAGCAGCACGATTTCTTCAAATCGGCCATGCCTGAAGGATACGTGCAAGAACGTACAATTAGCTTTAAAGATGACGGCACCTATAAAACGCGCGCCGAAGTTAAATTCGAGGGTGACACATTGGTTAATCGTATAGAACTTAAGGGCATTGACTTCAAAGAAGATGGCAACATCCTTGGCCATAAACTGGAATATAATTTTAACTCTCACAATGTCTACATTACGGCGGATAAACAAAAGAATGGCATTAAAGCGAATTTCAAAATCCGCCACAATGTCGAAGACGGCTCGGTTCAACTGGCGGACCATTATCAGCAGAACACCCCAATCGGTGATGGCCCGGTTCTGTTACCTGATAATCATTACCTTTCTACTCAAAGCGTTTTATCTAAAGATCCTAACGAGAAGCGTGATCATATGGTTCTACTGGAATTTGTTACCGCAGCTGGTATCACGCACGGCATGGATGAGCTGTATAAATGACCAGGCATCAAATAAAACGAAAGGCTCAGTCGAAAGACTGGGCCTTTCGTTTTATCTGTTGTTTGTCGGTGAACGCTCTCTACTAGAGTCACACTGGCTCACCTTCGGGTGGGCCTTTCTGCGTTTATAGGTCTCTCGCT |
| **pMC0_2-5_02_2-5 Dropout mScarlet-I (Vn)** | GGAGAGAGACCGAAAGTGAAACGTGATTTCATGCGTCATTTTGAACATTTTGTAAATCTTATTTAATAATGTGTGCGGCAATTCACATTTAATTTATGAATGTTTTCTTAACATCGCGGCAACTCAAGAAACGGCAGGTTCGGATCTTAGCTACTAGAGAAAGAGGAGAAATACTAGATGGTTTCTAAAGGTGAAGCAGTGATCAAAGAATTTATGCGCTTCAAAGTTCACATGGAAGGTTCTATGAACGGCCACGAATTTGAAATTGAAGGTGAAGGCGAAGGTCGTCCATACGAAGGTACTCAAACCGCAAAACTGAAAGTTACCAAAGGTGGTCCACTGCCATTTTCTTGGGATATTCTGTCTCCACAATTTATGTACGGTTCTCGTGCATTTATCAAACACCCAGCAGATATTCCAGACTACTACAAACAATCTTTTCCGGAAGGTTTCAAATGGGAACGTGTTATGAATTTTGAAGATGGTGGTGCAGTTACGGTTACCCAAGATACCTCTCTGGAAGATGGTACTCTGATCTACAAAGTTAAACTGCGTGGTACTAACTTTCCACCAGATGGTCCAGTTATGCAGAAAAAAACCATGGGTTGGGAAGCATCTACCGAACGTCTGTACCCAGAAGATGGCGTTCTGAAAGGTGATATCAAAATGGCACTGCGTCTGAAAGATGGCGGTCGTTACCTGGCAGATTTCAAAACCACCTACAAAGCGAAAAAACCAGTTCAAATGCCAGGTGCATACAACGTTGATCGTAAACTGGATATTACCAGCCACAACGAAGATTACACCGTTGTTGAACAATACGAACGTTCTGAAGGCCGTCACTCTACCGGTGGTATGGATGAACTGTACAAATAACCAGGCATCAAATAAAACGAAAGGCTCAGTCGAAAGACTGGGCCTTTCGTTTTATCTGTTGTTTGTCGGTGAACGCTCTCTACTAGAGTCACACTGGCTCACCTTCGGGTGGGCCTTTCTGCGTTTATAGGTCTCTCGCT |
| **pMC0_2_01_2 Dropout** | GGAGAGAGACCGAAAGTGAAACGTGATTTCATGCGTCATTTTGAACATTTTGTAAATCTTATTTAATAATGTGTGCGGCAATTCACATTTAATTTATGAATGTTTTCTTAACATCGCGGCAACTCAAGAAACGGCAGGTTCGGATCTTAGCTACTAGAGAAAGAGGAGAAATACTAGATGCGTAAAGGCGAAGAACTTTTCACTGGCGTAGTACCAATTTTAGTGGAGCTGGACGGAGATGTAAATGGTCATAAGTTTTCAGTTCGAGGAGAAGGCGAAGGAGATGCAACTAACGGTAAGCTGACACTAAAGTTTATCTGTACCACGGGTAAACTGCCGGTCCCATGGCCGACACTGGTTACTACACTTACTTATGGTGTACAATGCTTTGCTCGTTATCCTGACCATATGAAGCAGCACGATTTCTTCAAATCGGCCATGCCTGAAGGATACGTGCAAGAACGTACAATTAGCTTTAAAGATGACGGCACCTATAAAACGCGCGCCGAAGTTAAATTCGAGGGTGACACATTGGTTAATCGTATAGAACTTAAGGGCATTGACTTCAAAGAAGATGGCAACATCCTTGGCCATAAACTGGAATATAATTTTAACTCTCACAATGTCTACATTACGGCGGATAAACAAAAGAATGGCATTAAAGCGAATTTCAAAATCCGCCACAATGTCGAAGACGGCTCGGTTCAACTGGCGGACCATTATCAGCAGAACACCCCAATCGGTGATGGCCCGGTTCTGTTACCTGATAATCATTACCTTTCTACTCAAAGCGTTTTATCTAAAGATCCTAACGAGAAGCGTGATCATATGGTTCTACTGGAATTTGTTACCGCAGCTGGTATCACGCACGGCATGGATGAGCTGTATAAATGACCAGGCATCAAATAAAACGAAAGGCTCAGTCGAAAGACTGGGCCTTTCGTTTTATCTGTTGTTTGTCGGTGAACGCTCTCTACTAGAGTCACACTGGCTCACCTTCGGGTGGGCCTTTCTGCGTTTATAGGTCTCTTACT |
| **pMC0_2_02_PJ23100** | GGAGTTGACGGCTAGCTCAGTCCTAGGTACAGTGCTAGCTACT |
| **pMC0_2_03_PJ23101** | GGAGTTTACAGCTAGCTCAGTCCTAGGTATTATGCTAGCTACT |
| **pMC0_2_04_PJ23102** | GGAGTTGACAGCTAGCTCAGTCCTAGGTACTGTGCTAGCTACT |
| **pMC0_2_05_PJ23103** | GGAGCTGATAGCTAGCTCAGTCCTAGGGATTATGCTAGCTACT |
| **pMC0_2_06_PJ23104** | GGAGTTGACAGCTAGCTCAGTCCTAGGTATTGTGCTAGCTACT |
| **pMC0_2_07_PJ23105** | GGAGTTTACGGCTAGCTCAGTCCTAGGTACTATGCTAGCTACT |
| **pMC0_2_08_PJ23106** | GGAGTTTACGGCTAGCTCAGTCCTAGGTATAGTGCTAGCTACT |
| **pMC0_2_09_PJ23107** | GGAGTTTACGGCTAGCTCAGCCCTAGGTATTATGCTAGCTACT |
| **pMC0_2_10_PJ23108** | GGAGCTGACAGCTAGCTCAGTCCTAGGTATAATGCTAGCTACT |
| **pMC0_2_11_PJ23109** | GGAGTTTACAGCTAGCTCAGTCCTAGGGACTGTGCTAGCTACT |
| **pMC0_2_12_PJ23110** | GGAGTTTACGGCTAGCTCAGTCCTAGGTACAATGCTAGCTACT |
| **pMC0_2_13_PJ23111** | GGAGTTGACGGCTAGCTCAGTCCTAGGTATAGTGCTAGCTACT |
| **pMC0_2_14_PJ23113** | GGAGCTGATGGCTAGCTCAGTCCTAGGGATTATGCTAGCTACT |
| **pMC0_2_15_PJ23114** | GGAGTTTATGGCTAGCTCAGTCCTAGGTACAATGCTAGCTACT |
| **pMC0_2_16_PJ23115** | GGAGTTTATAGCTAGCTCAGCCCTTGGTACAATGCTAGCTACT |
| **pMC0_2_17_PJ23116** | GGAGTTGACAGCTAGCTCAGTCCTAGGGACTATGCTAGCTACT |
| **pMC0_2_18_PJ23117** | GGAGTTGACAGCTAGCTCAGTCCTAGGGATTGTGCTAGCTACT |
| **pMC0_2_19_PJ23118** | GGAGTTGACGGCTAGCTCAGTCCTAGGTATTGTGCTAGCTACT |
| **pMC0_2_20_PJ23119** | GGAGTTGACAGCTAGCTCAGTCCTAGGTATAATGCTAGCTACT |
| **pMC0_2_21_PDummy** | GGAGCCCCTGGCGCCCCTTTACT |
| **pMC0_2_22_PTrc** | GGAGGTCTAGGGCGGCGGATTTGTCCTACTCAGGAGAGCGTTCACCGACAAACAACAGATAAAACGAAAGGCCCAGTCTTTCGACTGAGCCTTTCGTTTTATTTGATGCAGCGGGTCGACTCCCCTAGGTCACTGCCCGCTTTCCAGTCGGGAAACCTGTCGTGCCAGCTGCATTAATGAATCGGCCAACGCGCGGGGAGAGGCGGTTTGCGTATTGGGCGCCAGGGTGGTTTTTCTTTTCACCAGTGACACGGGCAACAGCTGATTGCCCTTCACCGCCTGGCCCTGAGAGAGTTGCAGCAAGCGGTCCACGCTGGTTTGCCCCAGCAGGCGAAAATCCTGTTTGATGGTGGTTAACGGCGGGATATAACATGAGCTATCTTCGGTATCGTCGTATCCCACTACCGAGATATCCGCACCAACGCGCAGCCCGGACTCGGTAATGGCGCGCATTGCGCCCAGCGCCATCTGATCGTTGGCAACCAGCATCGCAGTGGGAACGATGCCCTCATTCAGCATTTGCATGGTTTGTTGAAAACCGGACATGGCACTCCAGTCGCCTTCCCGTTCCGCTATCGGCTGAATTTGATTGCGAGTGAGATATTTATGCCAGCCAGCCAGACGCAGACGCGCCGAGACAGAACTTAATGGGCCCGCTAACAGCGCGATTTGCTGGTGACCCAATGCGACCAGATGCTCCACGCCCAGTCGCGTACCATCTTCATGGGAGAAAATAATACTGTTGATGGGTGTCTGGTCAGAGACATCAAGAAATAACGCCGGAACATTAGTGCAGGCAGCTTCCACAGCAATGGCATCCTGGTCATCCAGCGGATAGTTAATGATCAGCCCACTGACGCGTTGCGCGAGAAGATTGTGCACCGCCGCTTTACAGGCTTCGACGCCGCTTCGTTCTACCATCGACACCACCACGCTGGCACCCAGTTGATCGGCGCGAGATTTAATCGCCGCGACAATTTGCGACGGCGCGTGCAGGGCCAGACTGGAGGTGGCAACGCCAATCAGCAACGACTGTTTGCCCGCCAGTTGTTGTGCCACGCGGTTGGGAATGTAATTCAGCTCCGCCATCGCCGCTTCCACTTTTTCCCGCGTTTTCGCAGAAACGTGGCTGGC |

| | |
|---|---|
| | CTGGTTCACCACGCGGGAAACGGTCTGATAAGAGACACCGGCATACTCTGCGACATCGTATAACGTTACTGGTTTCACATTCACCACCCTGAATTGACTCTCTTCCGGGCGCTATCATGCCATACCGCGAAAGGTTTTGCGCCAAAGCTTTCCCTCGACAATTCGATAAATGTGAGCGGATAACATTGACATTGGTGAGCGGATAACAAGATACTGAGCACATCAGCAGGACGCACTGACCTACT |
| **pMC0_2_23_PTet** | GGAGTTTTGTTATCAATAAAAAAGGCCCCCCGTTAGGGAGGCTTATTGTTCGTCCCATCACGGAAAAAGGTTATGCTGCTTTTAAGACCCACTTTCACATTTAAGTTGTTTTTCTAATCCGCATATGATCAATTCAAGGCCGAATAAGAAGGCTGGCTCTGCACCTTGGTGATCAAATAATTCGATAGCTTGTCGTAATAATGGCGGCATACTATCAGTAGTAGGTGTTTCCCTTTCTTCTTTAGCGACTTGATGCTCTTGATCTTCCAATACGCAACCTAAAGTAAAATGCCCCACAGCGCTGAGTGCATATAATGCATTCTCTAGTGAAAAACCTTGTTGGCATAAAAAGGCTAATTGATTTTCGAGAGTTTCATACTGTTTTTCTGTAGGCCGTGTACCTAAATGTACTTTTGCTCCATCGCGATGACTTAGTAAAGCACATCTAAAACTTTTAGCGTTATTACGTAAAAAATCTTGCCAGCTTTCCCCTTCTAAAGGGCAAAAGTGAGTATGGTGCCTATCTAACATCTCAATGGCTAAGGCGTCGAGCAAAGCCCGCTTATTTTTTACATGCCAATACAATGTAGGCTGCTCTACACCTAGCCTCTGGGCGAGTTTACGGGTTGTTAAACCTTCGATTCCGACCTCATTAAGCAGCTCTAATGCGCTGCTAATCACTCTACTTTTATCTAATCGAGACATCATTAATTCCTAATTTTTGTTGACACTCTATCATTGATAGAGTTATTTTACCACTCCCTATCAGTGATAGAGAAAAGTGAATACT |
| **pMC0_2_29_PBDA (Vn)** | GGAGTGTTATCCATCCACTGGTAGAGGTGAGTGTTCGCTATACATATTTGTGGATTTAGGGACATTTGTTAGTGACAAAAATCACAGCGGAAAAATGTAGCGAATTTGTCCATTCATTTAGCCAGTGTGGCTATGACACAGATCTCAATTATGCGACCAATGATCCAAATTCTCAGTAAGCAACCCAATACCAGCCTAATGCAAACGTTGAATTCGCTGGTTTCTTGCTTTTCCGACCTGACAGAAGAAGGTGTTAAAAAGAACACAAAAAATCGTCCATGACGTTTTTGTCCATAAAGTTAGCAGACCTCTTTATGGGATAACATCCCTCCTCTAGCTATAACAACAAGTAGATTTAGTTTGCTGACCAAAATACT |
| **pMC0_2_33_PRham (Vn)** | GGAGGACACACTCTAATAACCAAGCCCCGCAATTCGCGGGGCTTATTATTTTTAGCCAGCCAAATGTTACGCCCTCCCCGTTATTTCAAACAGTAAATAGCTTGAATAAATAAGAAAAACCACACCTTTTACACGCCTACTCCACTTCACTTAAACCCAGGTTTTATCTGGCCTCACGCACGAGTTGTCAAAAGTTTGAATTACCGCAAGAGCTTCTTGAGAAAAACGCATCGAATACGTTTTTTCAGGGGGATTTTTGAAGTTATTTAGTGCGGAAAAACGGTGTAATACCTCTAAAGAACAAGAGGTGTTAATCTACT |
| **pMC0_2_37_2 Dropout mScarlet-I (Vn)** | GGAGAGAGACCGAAAGTGAAACGTGATTTCATGCGTCATTTTGAACATTTTGTAAATCTTATTTAATAATGTGTGCGGCAATTCACATTTAATTTATGAATGTTTTCTTAACATCGCGGCAACTCAAGAAACGGCAGGTTCGGATCTTAGCTACTAGAGAAAGAGGAGAAATACTAGATGGTTTCTAAAGGTGAAGCAGTGATCAAAGAATTTATGCGCTTCAAAGTTCACATGGAAGGTTCTATGAACGGCCACGAATTTGAAATTGAAGGTGAAGGCGAAGGTCGTCCATACGAAGGTACTCAAACCGCAAAACTGAAAGTTACCAAAGGTGGTCCACTGCCATTTTCTTGGGATATTCTGTCTCCACAATTTATGTACGGTTCTCGTGCATTTATCAAACACCCAGCAGATATTCCAGACTACTACAAACAATCTTTTCCGGAAGGTTTCAAATGGGAACGTGTTATGAATTTTGAAGATGGTGGTGCAGTTACGGTTACCCAAGATACCTCTCTGGAAGATGGTACTCTGATCTACAAAGTTAAACTGCGTGGTACTAACTTTCCACCAGATGGTCCAGTTATGCAGAAAAAAACCATGGGTTGGGAAGCATCTACCGAACGTCTGTACCCAGAAGATGGCGTTCTGAAAGGTGATATCAAAATGGCACTGCGTCTGAAAGATGGCGGTCGTTACCTGGCAGATTTCAAAACCACCTACAAAGCGAAAAAACCAGTTCAAATGCCAGGTGCATACAACGTTGATCGTAAACTGGATATTACCAGCCACAACGAAGATTACACCGTTGTTGAACAATACGAACGTTCTGAAGGCCGTCACTCTACCGGTGGTATGGATGAACTGTACAAATAACCAGGCATCAAATAAAACGAAAGGCTCAGTCGAAAGACTGGGCCTTTCGTTTTATCTGTTGTTTGTCGGTGAACGCTCTCTACTAGAGTCACACTGGCTCACCTTCGGGTGGGCCTTTCTGCGTTTATAGGTCTCTTACT |
| **pMC0_3_01_3 Dropout** | TACTAGAGACCGAAAGTGAAACGTGATTTCATGCGTCATTTTGAACATTTTGTAAATCTTATTTAATAATGTGTGCGGCAATTCACATTTAATTTATGAATGTTTTCTTAACATCGCGGCAACTCAAGAAACGGCAGGTTCGGATCTTAGCTACTAGAGAAAGAGGAGAAATACTAGATGCGTAAAGGCGAAGAACTTTTCACTGGCGTAGTACCAATTTTAGTGGAGCTGGACGGAGATGTAAATGGTCATAAGTTTTCAGTTCGAGGAGAAGGCGAAGGAGATGCAACTAACGGTAAGCTGACACTAAAGTTTATCTGTACCACGGGTAAACTGCCGGTCCCATGGCCGACACTGGTTACTACACTTACTTATGGTGTACAATGCTTTGCTCGTTATCCTGACCATATGAAGCAGCACGATTTCTTCAAATCGGCCATGCCTGAAGGATACGTGCAAGAACGTACAATTAGCTTTAAAGATGACGGCACCTATAAAACGCGCGCCGAAGTTAAATTCGAGGGTGACACATTGGTTAATCGTATAGAACTTAAGGGCATTGACTTCAAAGAAGATGGCAACATCCTTGGCCATAAACTGGAATATAATTTTAACTCTCACAATGTCTACATTACGGCGGATAAACAAAAGAATGGCATTAAAGCGAATTTCAAAATCCGCCACAATGTCGAAGACGGCTCGGTTCAACTGGCGGACCATTATCAGCAGAACACCCCAATCGGTGATGGCCCGGTTCTGTTACCTGATAATCATTACCTTTCTACTCAAAGCGTTTTATCTAAAGATCCTAACGAGAAGCGTGATCATATGGTTCTACTGGAATTTGTTACCGCAGCTGGTATCACGCACGGCATGGATGAGCTGTATAAATGACCAGGCATCAAATAAAACGAAAGGCTCAGTCGAAAGACTGGGCCTTTCGTTTTATCTGTTGTTTGTCGGTGAACGCTCTCTACTAGAGTCACACTGGCTCACCTTCGGGTGGGCCTTTCTGCGTTTATAGGTCTCTAATG |
| **pMC0_3_02_RB0029** | TACTAGAGTTCACACAGGAAACCTAATCAATG |
| **pMC0_3_03_RB0030** | TACTAGAGATTAAAGAGGAGAAATAATCAATG |
| **pMC0_3_04_RB0031** | TACTAGAGTCACACAGGAAACCTAATCAATG |
| **pMC0_3_05_RB0032** | TACTAGAGTCACACAGGAAAGTAATCAATG |
| **pMC0_3_06_RB0033** | TACTAGAGTCACACAGGACTAATCAATG |
| **pMC0_3_07_RB0034** | TACTAGAGAAAGAGGAGAAATAATCAATG |
| **pMC0_3_08_RB0035** | TACTAGAGATTAAAGAGGAGAATAATCAATG |
| **pMC0_3_09_RB0064** | TACTAGAGAAAGAGGGGAAATAATCAATG |
| **pMC0_3_10_RDummy** | TACTAGAGTGTCAGGATACCCGATAATCAATG |
| **pMC0_3_11_3 Dropout mScarlet-I (Vn)** | TACTAGAGACCGAAAGTGAAACGTGATTTCATGCGTCATTTTGAACATTTTGTAAATCTTATTTAATAATGTGTGCGGCAATTCACATTTAATTTATGAATGTTTTCTTAACATCGCGGCAACTCAAGAAACGGCAGGTTCGGATCTTAGCTACTAGAGAAAGAGGAGAAATACTAGATGGTTTCTAAAGGTGAAGCAGTGATCAAAGAATTTATGCGCTTCAAAGTTCACATGGAAGGTTCTATGAACGGCCACGAATTTGAAATTGAAGGTGAAGGCGAAGGTCGTCCATACGAAGGTACTCAAACCGCAAAACTGAAAGTTACCAAAGGTGGTCCACTGCCATTTTCTTGGGATATTCTGTCTCCACAATTTATGTACGGTTCTCGTGCATTTATCAAACACCCAGCAGATATTCCAGACTACTACAAACAATCTTTTCCGGAAGGTTTCAAATGGGAACGTGTTATGAATTTTGAAGATGGTGGTGCAGTTACGGTTACCCAAGATACCTCTCTGGAAGATGGTACTCTGATCTACAAAGTTAAACTGCGTGGTACTAACTTTCCACCAGATGGTCCAGTTATGCAGAAAAAAACCATGGGTTGGGAAGCATCTACCGAACGTCTGTACCCAGAAGATGGCGTTCTGAAAGGTGATATCAAAATGGCACTGCGTCTGAAAGATGGCGGTCGTTACCTGGCAGATTTCAAAACCACCTACAAAGCGAAAAAACCAGTTCAAATGCCAGGTGCATACAACGTTGATCGTAAACTGGATATTACCAGCCACAACGAAGATTACACCGTTGTTGAACAATACGAACGTTCTGAAGGCCGTCACTCTACCGGTGGTATGGATGAACTGTACAAATAACCAGGCATCAAATAAAACGAAAGGCTCAGTCGAAAGACTGGGCCTTTCGTTTTATCTGTTGTTTGTCGGTGAACGCTCTCTACTAGAGTCACACTGGCTCACCTTCGGGTGGGCCTTTCTGCGTTTATAGGTCTCTAATG |
| **pMC0_4_01_4 Dropout** | AATGAGAGACCGAAAGTGAAACGTGATTTCATGCGTCATTTTGAACATTTTGTAAATCTTATTTAATAATGTGTGCGGCAATTCACATTTAATTTATGAATGTTTTCTTAACATCGCGGCAACTCAAGAAACGGCAGGTTCGGATCTTAGCTACTAGAGAAAGAGGAGAAATACTAGATGCGTAAAGGCGAAGAACTTTTCACTGGCGTAGTACCAATTTTAGTGGAGCTGGACGGAGATGTAAATGGTCATAAGTTTTCAGTTCGAGGAGAAGGCGAAGGAGATGCAACTAACGGTAAGCTGACACTAAAGTTTATCTGTACCACGGGTAAACTGCCGGTCCCATGGCCGACACTGGTTACTACACTTACTTATGGTGTACAATGCTTTGCTCGTTATCCTGACCATATGAAGCAGCACGATTTCTTCAAATCGGCCATGCCTGAAGGATACGTGCAAGAACGTACAATTAGCTTTAAAGATGACGGCACCTATAAAACGCGCGCCGAAGTTAAATTCGAGGGTGACACATTGGTTAATCGTATAGAACTTAAGGGCATTGACTTCAAAGAAGATGGCAACATCCTTGGCCATAAACTGGAATATAATTTTAACTCTCACAATGTCTACATTACGGCGGATAAACAAAAGAATGGCATTAAAGCGAATTTCAAAATCCGCCACAATGTCGAAGACGGCTCGGTTCAACTGGCGGACCATTATCAGCAGAACACCCCAATCGGTGATGGCCCGGTTCTGTTACCTGATAATCATTACCTTTCTACTCAAAGCGTTTTATCTAAAGATCCTAACGAGAAGCGTGATCATATGGTTCTACTGGAATTTGTTACCGCAGCTGGTATCACGCACGGCATGGATGAGCTGTATAAATGACCAGGCATCAAATAAAACGAAAGGCTCAGTCGAAAGACTGGGCCTTTCGTTTTATCTGTTGTTTGTCGGTGAACGCTCTCTACTAGAGTCACACTGGCTCACCTTCGGGTGGGCCTTTCTGCGTTTATAGGTCTCTGCTT |

| | |
|---|---|
| **pMC0_4_02_CDSlux Operon** | AATGACTAAAAAAATTTCATTCATTATTAACGGCCAGGTTGAAATCTTTCCCGAAAGTGATGATTTAGTGCAATCCATTAATTTTGGTGATAATAG<br>TGTTTACCTGCCAATATTGAATGACTCTCATGTAAAAAACATTATTGATTGTAATGGAAATAACGAATTACGGTTGCATAACATTGTCAATTTTCTC<br>TATACGGTAGGGCAAAGATGGAAAAATGAAGAATACTCAAGACGCAGGACATACATTCGTGACTTAAAAAAATATATGGGATATTCAGAAGAA<br>ATGGCTAAGCTAGAGGCCAATTGGATATCTATGATTTTATGTTCTAAAGGCGGCCTTTATGATGTTGTAGAAAATGAACTTGGTTCTCGCCATATC<br>ATGGATGAATGGCTACCTCAGGATGAAAGTTATGTTCGGGCTTTTCCGAAAGGTAAATCTGTACATCTGTTGGCAGGTAATGTTCCATTATCTGG<br>GATCATGTCTATATTACGCGCAATTTTAACTAAGAATCAGTGTATTATAAAAACATCGTCAACCGATCCTTTTACCGCTAATGCATTAGCGTTAAGT<br>TTTATTGATGTAGACCCTAATCATCCGATAACGCGCTCTTTATCTGTTATATATTGGCCCCACCAAGGTGATACATCACTCGCAAAAGAAATTATGC<br>GACATGCGGATGTTATTGTCGCTTGGGGAGGGCCAGATGCGATTAATTGGGCGGTAGAGCACGCGCCATCTTATGCTGATGTGATTAAATTTGG<br>TTCTAAAAAGAGTCTTTGCATTATCGATAATCCTGTTGATTTGACGTCCGCAGCGACAGGTGCGGCTCATGATGTTTGTTTTTACGATCAGCGAGC<br>TTGTTTTTCTGCCCAAAACATATATTACATGGGAAATCATTATGAGGAATTTAAGTTAGCGTTGATAGAAAAACTTAATCTATATGCGCATATATTA<br>CCGAATGCCAAAAAAGATTTTGATGAAAAGGCGGCCTATTCTTTAGTTCAAAAAGAAAGCCTGTTTGCTGGATTAAAAGTAGAGGTGGATATTCA<br>TCAACGTTGGATGATTATTGAGTCAAATGCAGGTGTGGAATTTAATCAACCACTTGGCAGATGTGTGTACCTTCATCACGTCGATAATATTGAGC<br>AAATATTGCCTTATGTTCAAAAAAATAAGACGCAAACCATATCTATTTTTCCTTGGGAGTCATCATTTAAATATCGAGATGCGTTAGCATTAAAAG<br>GTGCGGAAAGGATTGTAGAAGCAGGAATGAATAACATATTTCGAGTTGGTGGATCTCATGACGGAATGAGGCCGTTGCAACGATTAGTGACAT<br>ATATTTCTCATGAAAGGCCATCTAACTATACGGCTAAGGATGTTGCGGTTGAAATAGAACAGACTCGATTCCTGGAAGAAGATAAGTTCCTTGTA<br>TTTGTCCCATAATAGGTAAAAAGTATGGAAAATGAATCAAAATATAAAACCATCGACCACGTTATTTGTGTTGAAGGAAATAAAAAAATTCATGT<br>TTGGGAAACGCTGCCAGAAGAAAACAGCCCAAAGAGAAAGAATGCCATTATTATTGCGTCTGGTTTTGCCCGCAGGATGGATCATTTTGCTGGT<br>CTGGCGGAATATTTATCGCGGAATGGATTTCATGTGATCCGCTATGATTCGCTTCACCATGTTGGCTTGAGTTCAGGGACAATTGATGAATTTACA<br>ATGTCTATAGGAAAGCAGAGCTTGTTAGCAGTGGTTGATTGGTTAACTACACGAAAAATAAATAACTTCGGTATGTTGGCTTCCAGCTTATCTGC<br>GCGGATAGCTTATGCAAGCCTATCTGAAATCAATGCTTCGTTTTTAATCACCGCAGTCGGTGTTGTTAACTTAAGATATTCTCTTGAAAGAGCTTTA<br>GGGTTTGATTATCTCAGTCTACCCATTAATGAATTGCCGGATAATCTGGATTTTGAAGGCCATAAATTGGGTGCTGAAGTCTTTGCGAGAGATTG<br>TCTTGATTTTGGTTGGGAAGATTTAGCTTCTACAATTAATAACATGATGTATCTTGATATACCGTTTATTGCTTTTACTGCAAATAACGATAATTGG<br>GTCAAGCAAGATGAAGTTATCACATTGTTATCAAATATTCGTAGTAATCGATGCAAGATATATTCTTTGTTAGGAAGTTCGCATGACTTGAGTGAA<br>AATTTAGTGGTCCTGCGCAATTTTTATCAATCGGTTACGAAAGCCGCTATCGCGATGGATAATGATCATCTGGATATTGATGTTGATATTACTGAA<br>CCGTCATTTGAACATTTAACTATTGCGACAGTCAATGAACGCCGAATGAGAATTGAGATTGAAAATCAAGCAATTTCTCTGTCTTAAAATCTATTG<br>AGATATTCTATCACTCAAATAGCAATATAAGGACTCTCTATGAAATTTGGAAACTTTTTGCTTACATACCAACCTCCCCAATTTTCTCAAACAGAGG<br>TAATGAAACGTTTGGTTAAATTAGGTCGCATCTCTGAGGAGTGTGGTTTTGATACCGTATGGTTACTGGAGCATCATTTCACGGAGTTTGGTTTGC<br>TTGGTAACCCTTATGTCGCTGCTGCATATTTACTTGGCGCGACTAAAAAATTGAATGTAGGAACTGCCGCTATTGTTCTTCCCACAGCCCATCCAG<br>TACGCCAACTTGAAGATGTGAATTTATTGGATCAAATGTCAAAAGGACGATTTCGGTTTGGTATTTGCCGAGGGCTTTACAACAAGGACTTTCGC<br>GTATTCGGCACAGATATGAATAACAGTCGCGCCTTAGCGGAATGCTGGTACGGGCTGATAAAGAATGGCATGACAGAGGGATATATGGAAGCT<br>GATAATGAACATATCAAGTTCCATAAGGTAAAAGTAAACCCCGCGGCGTATAGCAGAGGTGGCGCACCGGTTTATGTGGTGGCTGAATCAGCTT<br>CGACGACTGAGTGGGCTGCTCAATTTGGCCTACCGATGATATTAAGTTGGATTATAAATACTAACGAAAAGAAAGCACAACTTGAGCTTTATAAT<br>GAAGTGGCTCAAGAATATGGGCACGATATTCATAATATCGACCATTGCTTATCATATATAACATCTGTAGATCATGACTCAATTAAAGCGAAAGA<br>GATTTGCCGGAAATTTCTGGGGCATTGGTATGATTCTTATGTGAATGCTACGACTATTTTTGATGATTCAGACCAAACAAGAGGTTATGATTTCAA<br>TAAAGGGCAGTGGCGTGACTTTGTATTAAAAGGACATAAAGATACTAATCGCCGTATTGATTACAGTTACGAAATCAATCCCGTGGGAACGCCG<br>CAGGAATGTATTGACATAATTCAAAAAGACATTGATGCTACAGGAATATCAAATATTTGTTGTGGATTTGAAGCTAATGGAACAGTAGACGAAAT<br>TATTGCTTCCATGAAGCTCTTCCAGTCTGATGTCATGCCATTTCTTAAAGAAAAACAACGTTCGCTATTATATTAGCTAAGGAGAAAGAAATGAAA<br>TTTGGATTGTTCTTCCTTAACTTCATCAATTCAACAACTGTTCAAGAACAAAGTATAGTTCGTATGCAGGAAATAACGGAGTATGTTGATAAGTTG<br>AATTTTGAACAGATTTTAGTGTATGAAAATCATTTTTCAGATAATGGTGTTGTCGGCGCTCCTCTGACTGTTTCTGGTTTTCTGCTCGGTTTAACAG<br>AGAAAATTAAAATTGGTTCATTAAATCACATCATTACAACTCATCATCCTGTCCGCATAGCGGAGGAAGCGTGCTTATTGGATCAGTTAAGTGAA<br>GGGAGATTTATTTTAGGGTTTAGTGATTGCGAAAAAAAAGATGAAATGCATTTTTTTAATCGCCCGGTTGAATATCAACAGCAACTATTTGAAGA<br>GTGTTATGAAATCATTAACGATGCTTTAACAACAGGCTATTGTAATCCAGATAACGATTTTTATAGCTTCCCTAAAATATCTGTAAATCCCCATGCT<br>TATACGCCAGGCGGACCTCGGAAATATGTAACAGCAACCAGTCATCATATTGTTGAGTGGGCGGCCAAAAAAGGTATTCCTCTCATCTTTAAGTG<br>GGATGATTCTAATGATGTTAGATATGAATATGCTGAAAGATATAAAGCCGTTGCGGATAAATATGACGTTGACCTATCAGAGATAGACCATCAGT<br>TAATGATATTAGTTAACTATAACGAAGATAGTAATAAAGCTAAACAAGAGACTCGTGCATTTATTAGTGATTATGTTCTTGAAATGCACCCTAATG<br>AAAATTTCGAAAATAAACTTGAAGAAATAATTGCAGAAAACGCTGTCGGAAATTATACGGAGTGTATAACTGCGGCTAAGTTGGCAATTGAAAA<br>GTGTGGTGCGAAAAGTGTATTGCTGTCCTTTGAACCAATGAATGATTTGATGAGCCAAAAAAATGTAATCAATATTGTTGATGATAATATTAAGA<br>AGTACCACATGGAATATACCTAATAGATTTCGAGTTGCAGCGAGGCGGCAAGTGAACGAATCCCCAGGAGCATAGATAACTATGTGACTGGGGT<br>GAGTGAAAGCAGCCAACAAAGCAGCAGCTTGAAAGATGAAGGGTATAAAAGAGTATGACAGCAGTGCTGCCATACTTTCTAATATTATCTTGAG<br>GAGTAAAACAGGTATGACTTCATATGTTGATAAACAAGAAATTACAGCAAGCTCAGAAATTGATGATTTGATTTTTTCGAGCGATCCATTAGTGT<br>GGTCTTACGACGAGCAGGAAAAAATCAGAAAGAAACTTGTGCTTGATGCATTTCGTAATCATTATAAACATTGTCGAGAATATCGTCACTACTGT<br>CAGGCACACAAAGTAGATGACAATATTACGGAAATTGATGACATACCTGTATTCCCAACATCGGTTTTTAAGTTTACTCGCTTATTAACTTCTCAG<br>GAAAACGAGATTGAAAGTTGGTTTACCAGTAGCGGCACGAATGGTTTAAAAAGTCAGGTGGCGCGTGACAGATTAAGTATTGAGAGACTCTTA<br>GGCTCTGTGAGTTATGGCATGAAATATGTTGGTAGTTGGTTTGATCATCAAATAGAATTAGTCAATTTGGGACCAGATAGATTTAATGCTCATAAT<br>ATTTGGTTTAAATATGTTATGAGTTTGGTGGAATTGTTATATCCTACGACATTTACCGTAACAGAAGAACGAATAGATTTTGTTAAAACATTGAAT<br>AGTCTTGAACGAATAAAAAATCAAGGGAAAGATCTTTGTCTTATTGGTTCGCCATACTTTATTTATTTACTCTGCCATTATATGAAAGATAAAAAA<br>ATCTCATTTTCTGGAGATAAAAGCCTTTATATCATAACCGGAGGCGGCTGGAAAAGTTACGAAAAAGAATCTCTGAAACGTGATGATTTCAATCA<br>TCTTTTATTTGATACTTTCAATCTCAGTGATATTAGTCAGATCCGAGATATATTTAATCAAGTTGAACTCAACACTTGTTTCTTTGAGGATGAAATG<br>CAGCGTAAACATGTTCCGCCGTGGGTATATGCGCGAGCGCTTGATCCTGAAACGTTGAAACCTGTACCTGATGGAACGCCGGGGTTGATGAGTT<br>ATATGGATGCGTCAGCAACCAGTTATCCAGCATTTATTGTTACCGATGATGTCGGGATAATTAGCAGAGAATATGGTAAGTATCCCGGCGTGCTC<br>GTTGAAATTTTACGTCGCGTCAATACGAGGACGCAGAAAGGGTGTGCTTTGAGCTTAACCGAAGCGTTTGATAGTGCTT |
| **pMC0_4_09_CDSAzurite (Vn)** | AATGTCTAAAGGTGAAGAACTGTTTACCGGTGTTGTTCCAATTCTGGTTGAACTGGATGGTGATGTTAACGGTCACAAATTTTCTGTTTCTGGTGA<br>AGGCGAAGGTGATGCAACCTACGGTAAACTGACCCTGAAATTTATCTGTACCACTGGTAAACTGCCAGTTCCATGGCCAACTCTGGTTACCACTC<br>TGTCTCACGGTGTTCAATGTTTTTCTCGTTACCCAGATCACATGAAACAGCACGATTTTTTCAAAAGCGCAATGCCAGAAGGTTACGTTCAAGAAC<br>GTACCATCTTCTTCAAAGATGACGGTAACTACAAAACCCGTGCGGAAGTGAAATTTGAAGGTGATACCCTGGTTAACCGTATCGAACTGAAAGGT<br>ATCGACTTCAAAGAGGATGGCAACATTCTGGGTCACAAACTGGAATACAACTTTAACTCTCACAACATCTACATCATGGCGGACAAACAAAAAAA<br>CGGCATCAAAGTGAACTTCAAGATTCGCCACAACATCGAAGATGGTTCTGTTCAACTGGCAGATCACTACCAACAAAACACCCCAATTGGTGATG<br>GTCCAGTTCTGCTGCCAGATAACCACTACCTGTCTACCCAATCTGCACTGTCTAAAGATCCAAACGAAAAACGTGATCACATGGTGCTGCTGGAAT<br>TTCGTACCGCAGCAGGTATTACCCACGGTATGGATGAACTGTACAAAGCTT |
| **pMC0_4_10_CDSmTurquoise (Vn)** | AATGGTTTCTAAAGGTGAAGAACTGTTTACCGGTGTTGTTCCAATTCTGGTTGAACTGGATGGTGATGTTAACGGTCACAAATTTTCTGTTTCTGG<br>TGAAGGCGAAGGTGATGCAACCTACGGTAAACTGACCCTGAAATTTATCTGTACCACTGGTAAACTGCCAGTTCCATGGCCAACTCTGGTTACCA<br>CTCTGTCTTGGGGTGTTCAATGTTTTGCACGTTACCCAGATCACATGAAACAACACGATTTTTTCAAAAGCGCAATGCCAGAAGGTTACGTTCAAG<br>AACGTACCATCTTCTTCAAAGATGACGGTAACTACAAAACCCGTGCGGAAGTGAAATTTGAAGGTGATACCCTGGTTAACCGTATCGAACTGAAA<br>GGTATCGACTTCAAAGAGGATGGCAACATTCTGGGTCACAAACTGGAATACAACTACTTTAGCGATAACGTGTACATCACCGCAGATAAACAAA<br>AAAACGGTATCAAGGCGAACTTCAAAATCCGTCACAACATTGAAGATGGTGGTGTTCAACTGGCAGATCACTACCAACAAAACACTCCAATTGGT<br>GATGGTCCAGTTCTGCTGCCAGATAACCACTACCTGTCTACCCAATCTAAACTGTCTAAAGACCCAAACGAAAAACGTGATCACATGGTTCTGCTG<br>GAATTTGTTACCGCAGCAGGTATTACCTTAGGTATGGATGAACTGTACAAAGCTT |
| **pMC0_4_11_CDSmVenus (Vn)** | AATGGTTTCTAAAGGTGAAGAACTGTTTACCGGTGTTGTTCCAATTCTGGTTGAACTGGATGGTGATGTTAACGGTCACAAATTTTCTGTTTCTGG<br>TGAAGGCGAAGGTGATGCAACCTACGGTAAACTGACCCTGAAACTGATTTGTACCACTGGTAAACTGCCAGTTCCATGGCCAACTCTGGTTACCA<br>CCTTAGGTTACGGTCTGCAATGTTTTGCACGTTACCCAGATCACATGAAACAACACGATTTTTTCAAAAGCGCAATGCCAGAAGGTTACGTTCAAG<br>AACGTACCATCTTCTTCAAAGATGACGGTAACTACAAAACCCGTGCGGAAGTGAAATTTGAAGGTGATACCCTGGTTAACCGTATCGAACTGAAA<br>GGTATCGACTTCAAAGAGGATGGCAACATTCTGGGTCACAAACTGGAATACAACTACAACTCTCACAACGTTTACATCACCGCAGATAAACAAAA<br>AAACGGCATCAAAGCGAACTTCAAAATCCGTCACAACATTGAAGATGGTGGTGTTCAACTGGCAGATCACTACCAACAAAACACCCCAATTGGTG |

| | |
|---|---|
| | ATGGTCCAGTTCTGCTGCCAGATAACCACTACCTGTCTTACCAATCTAAACTGAGCAAAGACCCAAACGAAAAACGTGATCACATGGTTCTGCTG<br>GAATTTGTTACCGCAGCAGGTATTACCTTAGGTATGGATGAACTGTACAAAGCTT |
| **pMC0_4_12_CDSmScarlet-I (Vn)** | AATGGTTTCTAAAGGTGAAGCAGTGATCAAAGAATTTATGCGCTTCAAAGTTCACATGGAAGGTTCTATGAACGGCCACGAATTTGAAATTGAA<br>GGTGAAGGCGAAGGTCGTCCATACGAAGGTACTCAAACCGCAAAACTGAAAGTTACCAAAGGTGGTCCACTGCCATTTTCTTGGGATATTCTGTC<br>TCCACAATTTATGTACGGTTCTCGTGCATTTATCAAACACCCAGCAGATATTCCAGACTACTACAAACAATCTTTTCCGGAAGGTTTCAAATGGGA<br>ACGTGTTATGAATTTTGAAGATGGTGGTGCAGTTACGGTTACCCAAGATACCTCTCTGGAAGATGGTACTCTGATCTACAAAGTTAAACTGCGTG<br>GTACTAACTTTCCACCAGATGGTCCAGTTATGCAGAAAAAAACCATGGGTTGGGAAGCATCTACCGAACGTCTGTACCCAGAAGATGGCGTTCTG<br>AAAGGTGATATCAAAATGGCACTGCGTCTGAAAGATGGCGGTCGTTACCTGGCAGATTTCAAAACCACCTACAAAGCGAAAAAACCAGTTCAAA<br>TGCCAGGTGCATACAACGTTGATCGTAAACTGGATATTACCAGCCACAACGAAGATTACACCGTTGTTGAACAATACGAACGTTCTGAAGGCCGT<br>CACTCTACCGGTGGTATGGATGAACTGTACAAAGCTT |
| **pMC0_4_13_CDSmCherry (Vn)** | AATGGTTTCTAAAGGTGAAGAGGATAACATGGCGATCATCAAAGAATTTATGCGCTTCAAAGTTCACATGGAAGGTTCTGTTAACGGCCACGAA<br>TTTGAAATTGAAGGTGAAGGCGAAGGTCGTCCATACGAAGGTACTCAAACCGCAAAACTGAAAGTTACCAAAGGTGGTCCACTGCCATTTGCAT<br>GGGATATTCTGTCTCCACAGTTTATGTACGGTAGCAAAGCATACGTTAAACACCCAGCAGATATTCCAGATTACCTGAAACTGTCTTTTCCGGAAG<br>GTTTCAAATGGGAACGTGTTATGAATTTTGAAGATGGTGGTGTTGTTACGGTTACCCAAGATTCTTCTCTGCAAGATGGTGAGTTTATCTACAAA<br>GTTAAACTGCGTGGCACCAACTTTCCATCTGATGGTCCAGTTATGCAGAAAAAAACCATGGGTTGGGAAGCATCTTCTGAACGTATGTACCCAGA<br>AGATGGCGCACTGAAAGGTGAAATTAAACAACGTCTGAAACTTAAAGATGGCGGTCACTACGATGCAGAAGTTAAAACCACCTACAAAGCGAA<br>AAAACCAGTTCAACTGCCAGGTGCATACAACGTTAACATTAAACTGGATATCACCAGCCACAACGAAGATTACACCATTGTTGAACAATACGAAC<br>GTGCAGAAGGCCGTCACTCTACCGGTGGTATGGATGAACTGTACAAAGCTT |
| **pMC0_4_14_CDSmKate-2 (Vn)** | AATGGTTTCTGAACTGATTAAAGAAAACATGCACATGAAACTGTACATGGAAGGTACTGTTAACAACCACCACTTCAAATGTACCTCTGAAGGTG<br>AAGGTAAACCATACGAAGGTACTCAAACCATGCGTATTAAAGCAGTTGAAGGTGGTCCACTGCCATTTGCATTTGATATTCTGGCAACCTCTTTTA<br>TGTACGGCAGCAAAACCTTTATCAACCACACTCAAGGTATCCCGGATTTTTTCAAACAAAGCTTTCCAGAAGGTTTCACCTGGGAACGTGTTACCA<br>CCTACGAAGATGGTGGTGTTCTGACCGCAACTCAAGATACCTCTCTGCAAGATGGTTGTCTGATCTACAACGTTAAAATCCGTGGTGTTAACTTTC<br>CATCTAACGGTCCAGTTATGCAGAAAAAAACCTTAGGTTGGGAAGCATCTACCGAAACTCTGTACCCAGCGGATGGTGGTCTGGAAGGTCGTGC<br>AGATATGGCACTGAAACTGGTTGGTGGTGGTCACCTGATTTGTAACCTGAAAACCACCTACCGTTCTAAAAAACCAGCGAAAAATCTGAAAATGC<br>CAGGTGTTTACTACGTTGATCGTCGTCTGGAACGTATCAAAGAAGCAGATAAAGAAACCTACGTGGAACAACACGAAGTTGCAGTTGCACGTTA<br>CTGTGATCTGCCATCTAAACTGGGTCACCGTGCTT |
| **pMC0_4_16_CDSVioA[#]** | AATGAGCACGTATTCTGACATTTGCATCGTTGGCGCCGGCATAGGAGGCTTGACTTGCGCCAACAACCTGATCGACGCCGCCGCCGGCAGGAAC<br>CTGCGCATCCGCGTATTCGACCTGAATGCCACCGTAGGCGGCCGCATCCAGTCGCGGAAAATAGATGGCGAGGAAATCGCCGAACTCGGCGCC<br>GCCCGCTACTCGCCGCAGCTGCATCCGCATTTCCAGCAACTCATGCAGGGCAGCGGCTTGCCGCATGCGGTCTACCCGTTCACCGAGGTCATCTC<br>CCACGATAGCGTGCTGGAGGAGCTGAAGGCAACGCTGGATGAGCTGAGCCCGATGCTGAAAATGCATCCGAACGACTCCTTCCTCGAGTTCGTC<br>AGCCATTACCTGGGCGCCGCCAAGGCCAGCCACATCATCAAGGCGACCGGCTATGACGCCCTGCTGCTGCCGATGGTGTCGGCGGCCATGGCCT<br>ACGACATCATCAAGAAGCACCCGGAAACGCAGCACTTTACGGAAAACGCCGCCAACCAGTGGCACTACGCCACCGACGGCTACCACGAATTGCT<br>GTGCCAGTTGCAGCACCAGGCCCAGGTCGCCGGGGTGGAATTCAGGCTCGAACACCGCTTGCTGTCCGTTGAAAAATCGGGCGCCGACCATGTG<br>CTCGCCTTCAGCCACCATGGCGACACGCAGATGCACCGCACGCGCCATCTGGTGATGGCCATCCCGCCGTCCGCCATGCCGCGCCTGAACCTGGA<br>TTTCCCGAACGCCTGGAGTCCGTTCCAATACGACTCGCTGCCCCTGTTCAAGGGATTCTTCACGTTCGACACAGCCTGGTGGGATGCGCTGGGGC<br>TGACCGACAAGGTGCTGATGGCGGCAAATCCCCTGCGCAAGATCTACTTCAAGAGCGACAAATACGTGCTGTTCTACACCGACAGCAAAAGCGC<br>CACCTACTGGCGGGACAGCCTGGAGCTTGGTGAGGACGTATACCTGGAGCGTGTCCGCAGCCACCTGGAAGAAGTCCTGCCGCTCGATGGCCA<br>GCCTCTGCCGCAGATCAAGGCGCACTTCCACAAGTTCTGGCCGCATGGCGTCGAGTTTTGCGTGGAGCCGGAAGCCGACCACCCGGCCATCCTG<br>CTGCACCGGGACGGCATCATCTCCTGCTCGGATGCCTATACCGCGCATTGCGGCTGGATGGAAGGCAGCCTGATCAGCGCCCAGCACGCCAGCG<br>GCCTGTTGCTGCAGCGCCTCGATCAACGGACGGAAGAAGAAGCTGCCAACGATACCTTCATCACTTCCTCGACCGAGCGCGCAGCTT |
| **pMC0_4_17_CDSVioB[#]** | AATGAGCCTACTTGACTTCCCCCGCCTGCATTTTCGGGGTTTTGCCCGCGCCAATGTGCCGACGGGGAATCGCAATACGCACGGCAACATCGATA<br>TCGCCACGAATGCGGTATCGATGGCGGGCGAGGCTGTCGACCTGAGCCGGCCGCCAGCCGAATTCCATGCGCACCTGAAACAGCTCGCCCCCCG<br>CTTCAACGCACAGGGCAAGCCCGATCCGGACGGCATCTTCAGCCAGGCGACAGGCTATAATTTTTGCGGGAACAACCATTTCTCGTGGGAAAAC<br>GCGCGGATCACGGGCGTCCAGTTGCGTGATGGCGAGGTCGATACCCAGGACGCGCTGGTGGGCGCCAAGCTGGGCCTGTGGGGCCACTACAA<br>CGAGTACCTGCGCACGACGTTCAACCGCGCAAGGTGGATCGACAACAACCCGGCGCAGCCCGACACCACGCTGATCTACGCGGGCCAGTTCACC<br>TTGAGCGACAAGCTGGCCACGCCCAATACGCCCACGCTGTTCACGGCCGACATCGCGCAGGCGCACTCGGTACGCTGGCTCGGCAGCGGCCACA<br>TCACGGAACGCAGCGGGCATTTCCTGGACGAGGAATTCGGCCGCTCCAGGCTGTTCCAGTTTTCCGTGGCCAAGCAGGACCCGCATTTCCTGTTC<br>AATCCGGACCTGCCGCTGCCGGCCAGTATGCATGCCTTGCAGCAAGCCCTGGCCGACGACGAGGTGCTGGGCCTGACGGTGCAATACTGCCTGT<br>TCAATATGTCGACGCCGCAAAAACCCGATTCGCCCGTGTTCTACGACCTGGCCGGCAGCATCGGCCTGTGGCGGCGCGGCGAGCTGGCCACCTA<br>TCCGGCCGGCCGCCTGCTGCAGCCGCGCCAGGGCAGCCTGGGGCCGGTGCTGGTGAAAGTGCATGCGGATCGCGTGAGTTTCAACATGCCTAC<br>CGCAATCCCGTTTACTACCCGTGATGCCGGGGCTGTATCCGAGCAGCATCCGACACATGCGCTGGGTGGTAAACAAGCATTGGGCGATCTGCTG<br>CTGCACGATGGGGCAGGGACCGTACTCGCCCGTATTCCGGAACAGTTGTATCGCGACTATTGGCGTCATCATGGCGTGTTTGATGTTCCATTACA<br>GCATGCGGGAGCTGCACCAGGTTCTCTTTCGCTCGGTTCAGCCCAGGCACAGTGGGATGAAGCCGATTGGGTCCTCCAAAGCGATTCTAACCAG<br>CTGTACCTGGAGGCACCGAATCGCAACAAACACGAGCAGTTTCCGCAGACCATCACGGTTCAATCGCGTTTCCGTGGTGAACTGGCTGCACCTCC<br>GTCACTGGCCGAAGCGGAAGATGGCGTCCTCTTAGCCGTCGAGCAGCAACCGAGTCCGTTAGGCCACGGCTATACCACGCTGACCCTCACAGGT<br>CGTAAACCGGGAGCAACGCGCATTGTCCTTGGAACTGGCAAAGCGAAACAGTATCTTGGCGTGCGGGTTCTGCCGGATGACTGGGACTTAGAC<br>GACGTTCCAGCAGAACAAGTGGACTATGCCTTCTTGTACCGCCATGTGATGAGTTACTACGAATTGGTTTACCCCTTTATGTCCGATAAGGTGTTC<br>AGCCTGGCTGATCAGTGCAAATGCGAAACGTATTCACGCCTGATGTGGCAGATGTGTGACCCGCAAAATCGGGACAAAAGCTACTACATGCCCA<br>GCACTCGCGAACTGTCGCTGCCTAAATCCCGCTTGTTTCTGAAATATCTGACCCAAGTAGAAGCCGCCGCGGCCGCAAAAGCGGCGGCTCCGGA<br>ACCAGCGGCGCCTCATGCGATTGGCGGCAAAGCTGAACTGATAGATGAGCTGAAGAAGGCGATTGACCTGGAATTAAGCCTGATGCTGCAGTA<br>TCTGTACGCTGCGTATAGCATCCCGAATTATGCGCAAGGTGCCGCGTTGGTGCAGTCTGGTCGCTGGCTTCCCGCGGAACTGGAGTTAGCGTGT<br>GGCGCCGAGGATCGGCGCCGCAACAGCGGCACGCGCGGCGCGCTGCTGGAAATCGCCCATGAAGAAATGATTCACTACTTATTGGTGAACAAT<br>GTATTGATGGCGCTTGGCGAACCGTTTTACAGCGGTACCCCGCTGCTGGGCCAGCAGGCGCGCCAGCGTTTCGGCCTGGACACGGAATTTGCGT<br>TCGAACCATTTTCCGAACACGTGCTGGCCCGCTTCGTGCGTTTTGAATGGCCCGACTACATTCCCACGCCGGGCAAATCCATCGCCACCTTCTATA<br>TCGCGATCCGCCAGGCCCTGGCCGAGCTGCCCGGCCTGTTCGAAAGCGGCGGCGGCAAGCGCGGCGGCGAGCACCACCTGTTCCTGAAAGAAC<br>TGACCAACCGCGCCTATCCCGGCTACCAGCTGGAAGTATCCGACCGCGACAGCGCGTTGTTCGCCATCGATTTCGTCACGGAACAGGGCGAAGG<br>CGTGGCCGTCGATTCGCCGCATTTCGCCTCCTCGCACTTCCAGCGGCTGCGCGCCATCGCCGGCAGGTTTTCGGCCTGCGACAAGCCGTTCGAAC<br>CGGCGCTGCCGGCGCTGAAGAATCCCGTGCTGGAAGCGCGCGCGGACTGCAGCGTGGTGACCGATCAGAAGGCGCGCGCGCTGATGCAGCTG<br>TATCAGGGCTGCTATGAACTGACCTTCCTGATGATGGCGCACCATTTTGCGCAGCAGCCGCTGGGCAGCCTGCGCCGCTCGCGCCTGATGAACGC<br>GTCCATCGACATCATGACAGGCCTGTTGCGCCCCCTGTCGGCGGCCCTGATGAACATGCCGTCCGGCCTGCCTGGCCGCCATGCTGGACCGCCCG<br>TGCCCGAGCCGGTCAGCAGCCGGGTCAGCAGCGACTACAGCCTGGGCTGCGACATGCTGGCGCAGAGATGCCTGGCGCTGGCGCAGTACGCGC<br>GCAGCCTGGAGAGCGATGCTATCGGCATGGCGCCGATAGAAATGTTGGAGTTTTTAATCAGCAACTTACCGATTTATCTCGGGGAAAGATGTC<br>AAGAGAGGCTGCTT |
| **pMC0_4_18_CDSVioC[#]** | AATGCATAAAATCATTATCGTCGGCGGAGGCCTGGCAGGCAGCCTCAGCGCTATTTATCTGGCGCAACGGGGGCACGATGTCCACGTTGTCGAA<br>AAGCGCGGCGATCCGCTGCTGGAGAATGCCGCAAACGCCGACCCCGTCAACTCGCGCGCCATCGGCGTGAGCATGACGGTACGCGGCATCAAG<br>GCCGTCCTGGCCGCCGGCATCAGCAAGCAGGAGCTCGACCAGTGCGGCGAACCCATCGTCGGCATGGCATTCAGCGTGGGCGGCCGGCACCGG<br>ATACGCGAGCTGACCCCGCTCGAAGGCCTGTTCCCCCTGTCGCTGGACCGCACCGCCTTCCAGCGCCTGCTGAACCGGCATGCCGCCCTGCATGA<br>GGTGAAGTATTACTTTGAGCATAAATGCCTGGATGTCGACCTGGAAAGAAAGATCGTGCTGATCCAGGGCCCGGACGGCGCCTTGCAGAAGCT<br>GCATGGCGACCTGGTCATTGGCGCCGACGGCGCCCACTCTGCCGTGCGGCGCGCCATGCAAAGCGGCATGCGCCGTTTCGAGTTCAGGCAAAGT<br>TACTTCCGCCACGGCTACAAGACGCTGGTGTTGCCGAACGCGGCGGATCTGGGTTTCAGGAAGGATTTGCTGTACTTCTTCGGCATGGATTCCAA<br>GGGCCTGTTTGCCGGCCGCGCGGCCACCATCCCGGACGGCAGCATCAGCTTTGCCCTGTGCCTGCCCTACACCGGCACGCCCAGCCTGGGCACG<br>CTCAACCGCGAAGCCATGGCCGATTTCTTCAGCCGCTACTTCGGCACCCTGCCGCCGGACCGTCGCAAGGAAATGCTGGACCAGTTCATGGCGCT<br>GCCCAGCAACGACCTCATCAATGTCCGTTCCAGCACCTTCCACTACAAGGCCAATATCCTGCTGATCGGCGATGCGGCGCATGCCACCGCCCCGT |

| | |
|---|---|
| | TCCTCGGGCAAGGCATGAACATGGCGCTGGAGGACGTCCACGTATTCGTGTCCCTGCTGGAAAAGCACGGCAATGCCCTGGGCCCTGCCCTGTC<br>CGAATTCACGCAGCAGCGCAAGGTGCAGGCGGACGCCATGCAGGACATGGCGATCGCCAACTATGAGGCGCTGAGCAATCCGAACCTGATTTT<br>CTTCCTGCAGACGCGCTACACGCGCTACATGCACAAGAAATTCCCCCGTGTTTATCCGCCAGACATGGCGGAGAAACTGTACTTCACATCGGTTC<br>CTTACGATGAATTGCAGCAAATCCAGAAGAAACAAAACGTTTGGTACAAACTTGGAAGGGTAAATGCTT |
| **pMC0_4_20_CDSVioE [#]** | AATGCCGACACACGTCAGCCCGCCGCTGCTGCCGATGCAATGGAGCAGCGCCTATGTTTCCTACTGGACGCCGATGCAGGCGGATGACCAGGTC<br>ACCTCCGGCTATTGCTGGTTCGACTATGCGCGCAATATCTGCCGCATCGACGGCCTGTTCAACCCCTGGTCGGAAAAGGAACATGGACACCTGCT<br>GTGGATGTCGGAAATCGGCGACGCCAGGCGCGAACAAAGCCGCAAGCAGAAAGTGGCCTACGCAAGGCAAGCGGAGGCGGCTGGCGAGCAG<br>CTGCAGGGCACGGCGCTGGCCGATGAGGTGACCCCGTTCCATGAGCTGTTCCTGCCGCAGGCGGTGCTGCTGGACGGCGGTGCCCGTCACGAC<br>GGCCGCCATACCGTGCTGGGCCGGGAGGCGGACGCCTGGGTAGTCGAGCGGGCGGGCAAGCCGCCATCGGTCTTTTACCTGGAGGCCGGTGG<br>CAACCGCCTGCTGCGCATGGTCACCGGCAATGACCCGCAGCACCTGTCGGTACGCGACTTTCCCAACCTGTTTGTCAGCGACATTCCGGACAGCG<br>TCTTTACGTCTTGCAACACCGCTT |
| **pMC0_4_21_4 Dropout mScarlet-I (Vn)** | AATGAGAGACCGAAAGTGAAACGTGATTTCATGCGTCATTTTGAACATTTTGTAAATCTTATTTAATAATGTGTGCGGCAATTCACATTTAATTTAT<br>GAATGTTTTCTTAACATCGCGGCAACTCAAGAAACGGCAGGTTCGGATCTTAGCTACTAGAGAAAGAGGAGAAATACTAGATGGTTTCTAAAGG<br>TGAAGCAGTGATCAAAGAATTTATGCGCTTCAAAGTTCACATGGAAGGTTCTATGAACGGCCACGAATTTGAAATTGAAGGTGAAGGCGAAGGT<br>CGTCCATACGAAGGTACTCAAACCGCAAAACTGAAAGTTACCAAAGGTGGTCCACTGCCATTTTCTTGGGATATTCTGTCTCCACAATTTATGTAC<br>GGTTCTCGTGCATTTATCAAACACCCAGCAGATATTCCAGACTACTACAAACAATCTTTTCCGGAAGGTTTCAAATGGGAACGTGTTATGAATTTT<br>GAAGATGGTGGTGCAGTTACGGTTACCCAAGATACCTCTCTGGAAGATGGTACTCTGATCTACAAAGTTAAACTGCGTGGTACTAACTTTCCACC<br>AGATGGTCCAGTTATGCAGAAAAAAACCATGGGTTGGGAAGCATCTACCGAACGTCTGTACCCAGAAGATGGCGTTCTGAAAGGTGATATCAAA<br>ATGGCACTGCGTCTGAAAGATGGCGGTCGTTACCTGGCAGATTTCAAAACCACCTACAAAGCGAAAAAACCAGTTCAAATGCCAGGTGCATACA<br>ACGTTGATCGTAAACTGGATATTACCAGCCACAACGAAGATTACACCGTTGTTGAACAATACGAACGTTCTGAAGGCCGTCACTCTACCGGTGGT<br>ATGGATGAACTGTACAAATAACCAGGCATCAAATAAAACGAAAGGCTCAGTCGAAAGACTGGGCCTTTCGTTTTATCTGTTGTTTGTCGGTGAAC<br>GCTCTCTACTAGAGTCACACTGGCTCACCTTCGGGTGGGCCTTTCTGCGTTTATAGGTCTCTGCTT |
| **pMC0_4_22_ftsZ [#]** | AATGTTTGAACCGATGATGGAAATGTCTGACGATGCAGTAATCAAGGTCGTTGGAGTTGGTGGCGGCGGTGGTAACGCTGTTGAACACATGGT<br>GCGTGAATCCATCGAAGGTGTGGAATTCATCAGCGTTAACACTGATGCGCAAGCACTTCGTAAGACAAGCGTTGGCAATGTTATTCAGATTGGT<br>GGTGATATCACCAAAGGTCTGGGTGCAGGTGCGAATCCACAGGTCGGCCGTGAGGCAGCTCTCGAAGATAGAGACAGAATTAAAGATTCCATA<br>ACTGGTGCCGATATGGTTTTTATCGCAGCAGGTATGGGTGGTGGTACAGGTACTGGTGCAGCTCCTGTTATCGCTGAAGTAGCCAAAGAGCTAG<br>GCATTCTTACTGTTGCAGTGGTAACTAAACCATTCAGCTTTGAAGGTAAGAAGCGTTTGGCGTTTGCTGAACAAGGTATCGATGAGCTTTCTAAA<br>CACGTTGACTCATTGATTACGATTCCAAACGAAAAGCTACTTAAAGTTCTTGGTCGTGGCGTGACGTTGCTAGAAGCGTTTGCAAGTGCGAATGA<br>TGTTCTTAAAAATGCAGTACAAGGTATTGCGGAACTAATTACTCGTCCGGGTATGATCAACGTCGACTTTGCTGACGTACGTACTGTTATGTCTGA<br>AATGGGCCACGCAATGATGGGTAGCGGTATCGCGAAAGGCGAAGACCGCGCTGAAGAAGCGGCTGAAATGGCAATTTCTAGTCCACTTCTGGA<br>AGACATCGATCTAGCTGGTGCACGTGGTGTTCTTGTTAACATTACTGCTGGCTTGGATATGCGTCTTGACGAGTTTGAGACAGTGGGTAATACAG<br>TTAAGGCATTTGCTTCTGATAATGCAACTGTAGTTATCGGTACTTCTCTAGACCCAGATATGACGGATGAAATCCGCGTAACTGTTGTAGCGACAG<br>GTATTGGTAACGAGAGAAAACCAGATATCACCCTTGTTGCTGGTGGTAAAGCGAAAGTAGCACCAACTACACAAGCTCAGCCACAACAACAAGC<br>AATGGCAACTCAAGCTGAAGAGAAGCCGGCACAAACTCTGCAACCTCAAGTGCAAGAGAAGCCACAGGTAACTCCTCAGCCGACAAATACAGC<br>GTCTTCTGCTCCTGCTTCGTCGAGCCAAAGCTCCGCGGCACCAAAGCAAGAGAAAGAGAGCGGTTATTTAGATATTCCGGCATTCTTGCGTCGCC<br>AGGCTGATGCTT |
| **pMC0_4_23_CDSsfgfp (Vn)** | AATGCGTAAAGGTGAAGAACTGTTTACCGGTGTTGTTCCAATTCTGGTTGAACTGGATGGTGATGTTAACGGTCACAAATTTTCTGTTCGTGGTG<br>AAGGCGAAGGTGATGCAACCAACGGTAAACTGACCCTGAAATTTATCTGTACCACTGGTAAACTGCCAGTTCCATGGCCAACTCTGGTTACCACT<br>CTGACCTACGGTGTTCAATGTTTTGCACGTTACCCAGATCACATGAAACAACACGATTTTTTCAAAAGCGCAATGCCAGAAGGTTACGTTCAAGA<br>ACGTACCATCTCTTTTAAAGATGACGGCACCTACAAAACCCGTGCGGAAGTTAAATTTGAAGGTGATACCCTGGTTAACCGCATTGAACTGAAAG<br>GCATCGATTTTAAAGAAGATGGTAACATCCTGGGCCACAAACTGGAATACAACTTTAACTCTCACAACGTGTACATCACCGCAGACAAACAAAAA<br>AACGGTATCAAAGCGAACTTCAAGATCCGTCACAACGTTGAAGATGGTTCTGTTCAACTGGCAGATCACTACCAACAAAACACCCCAATTGGTGA<br>TGGTCCAGTTCTGCTGCCAGATAACCACTACCTGTCTACCCAAAGCGTTCTGTCTAAAGATCCAAACGAAAAACGTGATCACATGGTGCTGCTGG<br>AATTTGTTACCGCAGCAGGTATTACCCACGGTATGGATGAACTGTACAAAGCAGCTT |
| **pMC0_4a_04_N3xFLAG** | AATGGATTATAAGGATCATGATGGTGATTATAAGGATCATGATATCGACTACAAAGACGATGACGACAAGGGGATG |
| **pMC0_4a_05_N6xHis** | AATGCACCATCACCACCATCATGGGATG |
| **pMC0_4a_06_NAzurite (Vn)** | AATGTCTAAAGGTGAAGAACTGTTTACCGGTGTTGTTCCAATTCTGGTTGAACTGGATGGTGATGTTAACGGTCACAAATTTTCTGTTTCTGGTGA<br>AGGCGAAGGTGATGCAACCTACGGTAAACTGACCCTGAAATTTATCTGTACCACTGGTAAACTGCCAGTTCCATGGCCAACTCTGGTTACCACTC<br>TGTCTCACGGTGTTCAATGTTTTTCTCGTTACCCAGATCACATGAAACAGCACGATTTTTTCAAAAGCGCAATGCCAGAAGGTTACGTTCAAGAAC<br>GTACCATCTTCTTCAAAGATGACGGTAACTACAAAACCCGTGCGGAAGTGAAATTTGAAGGTGATACCCTGGTTAACCGTATCGAACTGAAAGGT<br>ATCGACTTCAAAGAGGATGGCAACATTCTGGGTCACAAACTGGAATACAACTTTAACTCTCACAACATCTACATCATGGCGGACAAACAAAAAAA<br>CGGCATCAAAGTGAACTTCAAGATTCGCCACAACATCGAAGATGGTTCTGTTCAACTGGCAGATCACTACCAACAAAACACCCCAATTGGTGATG<br>GTCCAGTTCTGCTGCCAGATAACCACTACCTGTCTACCCAATCTGCACTGTCTAAAGATCCAAACGAAAAACGTGATCACATGGTGCTGCTGGAAT<br>TTCGTACCGCAGCAGGTATTACCCACGGTATGGATGAACTGTACAAAGGGATG |
| **pMC0_4a_07_NmTurquoise 2 (Vn)** | AATGGTTTCTAAAGGTGAAGAACTGTTTACCGGTGTTGTTCCAATTCTGGTTGAACTGGATGGTGATGTTAACGGTCACAAATTTTCTGTTTCTGG<br>TGAAGGCGAAGGTGATGCAACCTACGGTAAACTGACCCTGAAATTTATCTGTACCACTGGTAAACTGCCAGTTCCATGGCCAACTCTGGTTACCA<br>CTCTGTCTTGGGGTGTTCAATGTTTTGCACGTTACCCAGATCACATGAAACAACACGATTTTTTCAAAAGCGCAATGCCAGAAGGTTACGTTCAAG<br>AACGTACCATCTTCTTCAAAGATGACGGTAACTACAAAACCCGTGCGGAAGTGAAATTTGAAGGTGATACCCTGGTTAACCGTATCGAACTGAAA<br>GGTATCGACTTCAAAGAGGATGGCAACATTCTGGGTCACAAACTGGAATACAACTACTTTAGCGATAACGTGTACATCACCGCAGATAAACAAA<br>AAAACGGTATCAAGGCGAACTTCAAAATCCGTCACAACATTGAAGATGGTGGTGTTCAACTGGCAGATCACTACCAACAAAACACTCCAATTGGT<br>GATGGTCCAGTTCTGCTGCCAGATAACCACTACCTGTCTACCCAATCTAAACTGTCTAAAGACCCAAACGAAAAACGTGATCACATGGTTCTGCTG<br>GAATTTGTTACCGCAGCAGGTATTACCTTAGGTATGGATGAACTGTACAAAGGGATG |
| **pMC0_4a_09_NmVenus (Vn)** | AATGGTTTCTAAAGGTGAAGAACTGTTTACCGGTGTTGTTCCAATTCTGGTTGAACTGGATGGTGATGTTAACGGTCACAAATTTTCTGTTTCTGG<br>TGAAGGCGAAGGTGATGCAACCTACGGTAAACTGACCCTGAAACTGATTTGTACCACTGGTAAACTGCCAGTTCCATGGCCAACTCTGGTTACCA<br>CCTTAGGTTACGGTCTGCAATGTTTTGCACGTTACCCAGATCACATGAAACAACACGATTTTTTCAAAAGCGCAATGCCAGAAGGTTACGTTCAAG<br>AACGTACCATCTTCTTCAAAGATGACGGTAACTACAAAACCCGTGCGGAAGTGAAATTTGAAGGTGATACCCTGGTTAACCGTATCGAACTGAAA<br>GGTATCGACTTCAAAGAGGATGGCAACATTCTGGGTCACAAACTGGAATACAACTACAACTCTCACAACGTTTACATCACCGCAGATAAACAAAA<br>AAACGGCATCAAAGCGAACTTCAAAATCCGTCACAACATTGAAGATGGTGGTGTTCAACTGGCAGATCACTACCAACAAAACACCCCAATTGGTG<br>ATGGTCCAGTTCTGCTGCCAGATAACCACTACCTGTCTTACCAATCTAAACTGAGCAAAGACCCAAACGAAAAACGTGATCACATGGTTCTGCTG<br>GAATTTGTTACCGCAGCAGGTATTACCTTAGGTATGGATGAACTGTACAAAGGGATG |
| **pMC0_4a_10_NmScarlet-I (Vn)** | AATGGTTTCTAAAGGTGAAGCAGTGATCAAAGAATTTATGCGCTTCAAAGTTCACATGGAAGGTTCTATGAACGGCCACGAATTTGAAATTGAA<br>GGTGAAGGCGAAGGTCGTCCATACGAAGGTACTCAAACCGCAAAACTGAAAGTTACCAAAGGTGGTCCACTGCCATTTTCTTGGGATATTCTGTC<br>TCCACAATTTATGTACGGTTCTCGTGCATTTATCAAACACCCAGCAGATATTCCAGACTACTACAAACAATCTTTTCCGGAAGGTTTCAAATGGGA<br>ACGTGTTATGAATTTTGAAGATGGTGGTGCAGTTACGGTTACCCAAGATACCTCTCTGGAAGATGGTACTCTGATCTACAAAGTTAAACTGCGTG<br>GTACTAACTTTCCACCAGATGGTCCAGTTATGCAGAAAAAAACCATGGGTTGGGAAGCATCTACCGAACGTCTGTACCCAGAAGATGGCGTTCTG<br>AAAGGTGATATCAAAATGGCACTGCGTCTGAAAGATGGCGGTCGTTACCTGGCAGATTTCAAAACCACCTACAAAGCGAAAAAACCAGTTCAAA<br>TGCCAGGTGCATACAACGTTGATCGTAAACTGGATATTACCAGCCACAACGAAGATTACACCGTTGTTGAACAATACGAACGTTCTGAAGGCCGT<br>CACTCTACCGGTGGTATGGATGAACTGTACAAAGGGATG |
| **pMC0_4a_11_NmCherry (Vn)** | AATGGTTTCTAAAGGTGAAGAGGATAACATGGCGATCATCAAAGAATTTATGCGCTTCAAAGTTCACATGGAAGGTTCTGTTAACGGCCACGAA<br>TTTGAAATTGAAGGTGAAGGCGAAGGTCGTCCATACGAAGGTACTCAAACCGCAAAACTGAAAGTTACCAAAGGTGGTCCACTGCCATTTGCAT |

| | |
|---|---|
| | GGGATATTCTGTCTCCACAGTTTATGTACGGTAGCAAAGCATACGTTAAACACCCAGCAGATATTCCAGATTACCTGAAACTGTCTTTTCCGGAAG<br>GTTTCAAATGGGAACGTGTTATGAATTTTGAAGATGGTGGTGTTGTTACGGTTACCCAAGATTCTTCTCTGCAAGATGGTGAGTTTATCTACAAA<br>GTTAAACTGCGTGGCACCAACTTTCCATCTGATGGTCCAGTTATGCAGAAAAAAACCATGGGTTGGGAAGCATCTTCTGAACGTATGTACCCAGA<br>AGATGGCGCACTGAAAGGTGAAATTAAACAACGTCTGAAACTTAAAGATGGCGGTCACTACGATGCAGAAGTTAAAACCACCTACAAAGCGAA<br>AAAACCAGTTCAACTGCCAGGTGCATACAACGTTAACATTAAACTGGATATCACCAGCCACAACGAAGATTACACCATTGTTGAACAATACGAAC<br>GTGCAGAAGGCCGTCACTCTACCGGTGGTATGGATGAACTGTACAAAGGGATG |
| **pMC0_4a_12_NmKate (Vn)** | AATGGTTTCTGAACTGATTAAAGAAAACATGCACATGAAACTGTACATGGAAGGTACTGTTAACAACCACCACTTCAAATGTACCTCTGAAGGTG<br>AAGGTAAACCATACGAAGGTACTCAAACCATGCGTATTAAAGCAGTTGAAGGTGGTCCACTGCCATTTGCATTTGATATTCTGGCAACCTCTTTTA<br>TGTACGGCAGCAAAACCTTTATCAACCACACTCAAGGTATCCCGGATTTTTTCAAACAAAGCTTTCCAGAAGGTTTCACCTGGGAACGTGTTACCA<br>CCTACGAAGATGGTGGTGTTCTGACCGCAACTCAAGATACCTCTCTGCAAGATGGTTGTCTGATCTACAACGTTAAAATCCGTGGTGTTAACTTTC<br>CATCTAACGGTCCAGTTATGCAGAAAAAAACCTTAGGTTGGGAAGCATCTACCGAAACTCTGTACCCAGCGGATGGTGGTCTGGAAGGTCGTGC<br>AGATATGGCACTGAAACTGGTTGGTGGTGGTCACCTGATTTGTAACCTGAAAACCACCTACCGTTCTAAAAAACCAGCGAAAAATCTGAAAATGC<br>CAGGTGTTTACTACGTTGATCGTCGTCTGGAACGTATCAAAGAAGCAGATAAAGAAACCTACGTGGAACAACACGAAGTTGCAGTTGCACGTTA<br>CTGTGATCTGCCATCTAAACTGGGTCACCGTGGGATG |
| **pMC0_4a_13_Nsfgfp(Vn)** | AATGCGTAAAGGTGAAGAACTGTTTACCGGTGTTGTTCCAATTCTGGTTGAACTGGATGGTGATGTTAACGGTCACAAATTTTCTGTTCGTGGTG<br>AAGGCGAAGGTGATGCAACCAACGGTAAACTGACCCTGAAATTTATCTGTACCACTGGTAAACTGCCAGTTCCATGGCCAACTCTGGTTACCACT<br>CTGACCTACGGTGTTCAATGTTTTGCACGTTACCCAGATCACATGAAACAACACGATTTTTTCAAAAGCGCAATGCCAGAAGGTTACGTTCAAGA<br>ACGTACCATCTCTTTTAAAGATGACGGCACCTACAAAACCCGTGCGGAAGTTAAATTTGAAGGTGATACCCTGGTTAACCGCATTGAACTGAAAG<br>GCATCGATTTTAAAGAAGATGGTAACATCCTGGGCCACAAACTGGAATACAACTTTAACTCTCACAACGTGTACATCACCGCAGACAAACAAAAA<br>AACGGTATCAAAGCGAACTTCAAGATCCGTCACAACGTTGAAGATGGTTCTGTTCAACTGGCAGATCACTACCAACAAAACACCCCAATTGGTGA<br>TGGTCCAGTTCTGCTGCCAGATAACCACTACCTGTCTACCCAAAGCGTTCTGTCTAAAGATCCAAACGAAAAACGTGATCACATGGTGCTGCTGG<br>AATTTGTTACCGCAGCAGGTATTACCCACGGTATGGATGAACTGTACAAAGCAGGGATG |
| **pMC0_4b_01_ftsA [#]** | GATGACTAAGGCCGCAGACGACAACATTATTGTTGGTCTTGATATAGGCACTGCAACCGTATCAGCTCTAGTGGGCGAAATTTTACCAGATGGCC<br>AAATTAATATAATTGGTGCGGGTAGTAGCCCTTCGCGCGGCATGGACAAAGGTGGCGTGAATGACTTGGAGTCGGTTGTAAAATCAGTACAACG<br>CGCAATTGATCAAGCTGAGTTAATGGCCGAGTGCCAAATCAGCCGAGTATTCATTTCGTTGTCTGGTAAGCATATCGCAAGCCGAATCGAAAAAG<br>GTATGGGTACGATATCTGATGAGGAAGTTTCTCAAGAGGATATGGACAGAGCTATCCATACCGCAAAATCAATTAAAATCGGTGATGAACAGCG<br>AATCCTTCATGTGATTCCACAAGAGTTTACCATCGACTACCAGGAAGGGATTAAAAACCCGCTTGGTCTATCTGGGGTAAGAATGGAAGTAAGTG<br>TCCATCTTATCACCTGCCATAATGACATGGCGCGCAATATCATTAAGGCGGTAGAACGTTGCGGCCTTAAGGTAGAGCAACTTGTATTTTCTGGG<br>CTTGCTGCAAGTAATGCAGTAATTACCGAGGATGAAAGAGAGCTTGGAGTTTGTGTGGTCGATATTGGCGCAGGCACGATGGATGTTGCAATCT<br>GGACGGGTGGTGCGCTACGTCACACAGAAGTATTTTCGTACGCGGGAAATGCGGTAACAAGCGATATTGCTTTTGCTTTTGGTACGCCAGTAAG<br>CGATGCTGAAGAAATAAAAGTAAAATATGGCTGTGCTTTAAGTGAACTGGTCAGTAAGGATGACACGGTTAACGTTCCAAGCGTAGGTGGGCG<br>ACCTTCACGTAGTCTGCAACGTCAAACATTGTCAGAAGTGATAGAGCCACGCTACTCTGAGCTTATGGGACTGGTTAACCAAACGATCGATACCG<br>TTCAGGCAAAATTACGCGCAGAGGGCATTAAGCACCATCTGGCAGCCGGTGTAGTACTGACTGGCGGCGCTGCGCAAATGGACGGGGTCGTCG<br>AATGTGCGGAACGCGTATTCCGCAACCAAGTAAGGGTCGGCAAACCGCTCGAAGTCAGCGGCTTAACTGACTATGTAAAAGAGCCGTACCATTC<br>TACGGCAGTTGGATTACTTCATTACGCAAGAGACATGCAGTCCAGCGACGATAGCGATTACAATGAACCTAAACGCTCATCTGTTACTGGCTTTTT<br>CGGTAAATTGCGTAACTGGATACAAAAAGAGTTTGCTT |
| **pMC0_5_01_5 Dropout** | GCTTAGAGACCGAAAGTGAAACGTGATTTCATGCGTCATTTTGAACATTTTGTAAATCTTATTTAATAATGTGTGCGGCAATTCACATTTAATTTAT<br>GAATGTTTTCTTAACATCGCGGCAACTCAAGAAACGGCAGGTTCGGATCTTAGCTACTAGAGAAAGAGGAGAAATACTAGATGCGTAAAGGCGA<br>AGAACTTTTCACTGGCGTAGTACCAATTTTAGTGGAGCTGGACGGAGATGTAAATGGTCATAAGTTTTCAGTTCGAGGAGAAGGCGAAGGAGAT<br>GCAACTAACGGTAAGCTGACACTAAAGTTTATCTGTACCACGGGTAAACTGCCGGTCCCATGGCCGACACTGGTTACTACACTTACTTATGGTGT<br>ACAATGCTTTGCTCGTTATCCTGACCATATGAAGCAGCACGATTTCTTCAAATCGGCCATGCCTGAAGGATACGTGCAAGAACGTACAATTAGCTT<br>TAAAGATGACGGCACCTATAAAACGCGCGCCGAAGTTAAATTCGAGGGTGACACATTGGTTAATCGTATAGAACTTAAGGGCATTGACTTCAAA<br>GAAGATGGCAACATCCTTGGCCATAAACTGGAATATAATTTTAACTCTCACAATGTCTACATTACGGCGGATAAACAAAAGAATGGCATTAAAGC<br>GAATTTCAAAATCCGCCACAATGTCGAAGACGGCTCGGTTCAACTGGCGGACCATTATCAGCAGAACACCCCAATCGGTGATGGCCCGGTTCTGT<br>TACCTGATAATCATTACCTTTCTACTCAAAGCGTTTTATCTAAAGATCCTAACGAGAAGCGTGATCATATGGTTCTACTGGAATTTGTTACCGCAGC<br>TGGTATCACGCACGGCATGGATGAGCTGTATAAATGACCAGGCATCAAATAAAACGAAAGGCTCAGTCGAAAGACTGGGCCTTTCGTTTTATCT<br>GTTGTTTGTCGGTGAACGCTCTCTACTAGAGTCACACTGGCTCACCTTCGGGTGGGCCTTTCTGCGTTTATAGGTCTCTCGCT |
| **pMC0_5_02_TB0010** | GCTTAACCAGGCATCAAATAAAACGAAAGGCTCAGTCGAAAGACTGGGCCTTTCGTTTTATCTGTTGTTTGTCGGTGAACGCTCTCCGCT |
| **pMC0_5_03_TB0015** | GCTTAACCAGGCATCAAATAAAACGAAAGGCTCAGTCGAAAGACTGGGCCTTTCGTTTTATCTGTTGTTTGTCGGTGAACGCTCTCTACTAGAGTC<br>ACACTGGCTCACCTTCGGGTGGGCCTTTCTGCGTTTATACGCT |
| **pMC0_5_04_TB1002** | GCTTAACGCAAAAAACCCCGCTTCGGCGGGGTTTTTTCGCCGCT |
| **pMC0_5_05_TB1003** | GCTTAACGCCAAAAACCCCGCTTCGGCGGGGTTTTTCCGCCGCT |
| **pMC0_5_06_TB1004** | GCTTAACGCCGAAAACCCCGCTTCGGCGGGGTTTTGCCGCCGCT |
| **pMC0_5_07_TB1005** | GCTTAACGCCGCAAACCCCGCTTCGGCGGGGTTTCGCCGCCGCT |
| **pMC0_5_08_TB1006** | GCTTAAAAAAAAAAAACCCCGCCCCTGACAGGGCGGGGTTTTTTTTCGCT |
| **pMC0_5_09_TB1007** | GCTTAACGCAAAAAACCCCGCCCCTGACAGGGCGGGGTTTTTTCGCCGCT |
| **pMC0_5_11_TB1009** | GCTTAACGCCGAAAACCCCGCCCCTGACAGGGCGGGGTTTTGCCGCCGCT |
| **pMC0_5_12_TB1010** | GCTTAACGCCGCAAACCCCGCCCCTGACAGGGCGGGGTTTCGCCGCCGCT |
| **pMC0_5_13_TDummy** | GCTTAAACTCAGTTGTAGTAACGAGCGGATAGATTCCCAGACCCACCTTCACGGGCGGTAGCAGGACCTCAATAATAGGATTTTCGCGCT |
| **pMC0_5_14_5 Dropout mScarlet-I (Vn)** | GCTTAGAGACCGAAAGTGAAACGTGATTTCATGCGTCATTTTGAACATTTTGTAAATCTTATTTAATAATGTGTGCGGCAATTCACATTTAATTTAT<br>GAATGTTTTCTTAACATCGCGGCAACTCAAGAAACGGCAGGTTCGGATCTTAGCTACTAGAGAAAGAGGAGAAATACTAGATGGTTTCTAAAGG<br>TGAAGCAGTGATCAAAGAATTTATGCGCTTCAAAGTTCACATGGAAGGTTCTATGAACGGCCACGAATTTGAAATTGAAGGTGAAGGCGAAGGT<br>CGTCCATACGAAGGTACTCAAACCGCAAAACTGAAAGTTACCAAAGGTGGTCCACTGCCATTTTCTTGGGATATTCTGTCTCCACAATTTATGTAC<br>GGTTCTCGTGCATTTATCAAACACCCAGCAGATATTCCAGACTACTACAAACAATCTTTTCCGGAAGGTTTCAAATGGGAACGTGTTATGAATTTT<br>GAAGATGGTGGTGCAGTTACGGTTACCCAAGATACCTCTCTGGAAGATGGTACTCTGATCTACAAAGTTAAACTGCGTGGTACTAACTTTCCACC<br>AGATGGTCCAGTTATGCAGAAAAAAACCATGGGTTGGGAAGCATCTACCGAACGTCTGTACCCAGAAGATGGCGTTCTGAAAGGTGATATCAAA<br>ATGGCACTGCGTCTGAAAGATGGCGGTCGTTACCTGGCAGATTTCAAAACCACCTACAAAGCGAAAAAACCAGTTCAAATGCCAGGTGCATACA<br>ACGTTGATCGTAAACTGGATATTACCAGCCACAACGAAGATTACACCGTTGTTGAACAATACGAACGTTCTGAAGGCCGTCACTCTACCGGTGGT<br>ATGGATGAACTGTACAAATAACCAGGCATCAAATAAAACGAAAGGCTCAGTCGAAAGACTGGGCCTTTCGTTTTATCTGTTGTTTGTCGGTGAAC<br>GCTCTCTACTAGAGTCACACTGGCTCACCTTCGGGTGGGCCTTTCTGCGTTTATAGGTCTCTCGCT |
| **pMC0_5a_04_C3xFLAG** | GCTTTAGATTATAAGGATCATGATGGTGATTATAAGGATCATGATATCGACTACAAAGACGATGACGACAAGGGGTA |
| **pMC0_5a_05_C6xHis** | GCTTTACACCATCACCACCATCATGGGTA |
| **pMC0_5a_06_CI11012** | GCTTTAGCAGCAAACGACGAAAACTACGCTGCTGCTGTTTAGGGGTA |

| | |
|---|---|
| **pMC0_5a_07_CM0050** | GCTTTAGCTGCTAACGACGAAAACTACGCTCTGGCTGCTTAGGGGTA |
| **pMC0_5a_08_CM0051** | GCTTTAGCTGCTAACGACGAAAACTACAACTACGCTGACGCTTCTTAGGGGTA |
| **pMC0_5a_09_CM0052** | GCTTTAGCTGCTAACGACGAAAACTACGCTGACGCTTCTTAGGGGTA |
| **pMC0_5a_10_CAzurite (Vn)** | GCTTTATCTAAAGGTGAAGAACTGTTTACCGGTGTTGTTCCAATTCTGGTTGAACTGGATGGTGATGTTAACGGTCACAAATTTTCTGTTTCTGGTGAAGGCGAAGGTGATGCAACCTACGGTAAACTGACCCTGAAATTTATCTGTACCACTGGTAAACTGCCAGTTCCATGGCCAACTCTGGTTACCACTCTGTCTCACGGTGTTCAATGTTTTTCTCGTTACCCAGATCACATGAAACAGCACGATTTTTTCAAAAGCGCAATGCCAGAAGGTTACGTTCAAGAACGTACCATCTTCTTCAAAGATGACGGTAACTACAAAACCCGTGCGGAAGTGAAATTTGAAGGTGATACCCTGGTTAACCGTATCGAACTGAAAGGTATCGACTTCAAAGAGGATGGCAACATTCTGGGTCACAAACTGGAATACAACTTTAACTCTCACAACATCTACATCATGGCGGACAAACAAAAAAACGGCATCAAAGTGAACTTCAAGATTCGCCACAACATCGAAGATGGTTCTGTTCAACTGGCAGATCACTACCAACAAAACACCCCAATTGGTGATGGTCCAGTTCTGCTGCCAGATAACCACTACCTGTCTACCCAATCTGCACTGTCTAAAGATCCAAACGAAAAACGTGATCACATGGTGCTGCTGGAATTTCGTACCGCAGCAGGTATTACCCACGGTATGGATGAACTGTACAAAGGGTA |
| **pMC0_5a_11_CmTurquoise (Vn)** | GCTTTAGTTTCTAAAGGTGAAGAACTGTTTACCGGTGTTGTTCCAATTCTGGTTGAACTGGATGGTGATGTTAACGGTCACAAATTTTCTGTTTCTGGTGAAGGCGAAGGTGATGCAACCTACGGTAAACTGACCCTGAAATTTATCTGTACCACTGGTAAACTGCCAGTTCCATGGCCAACTCTGGTTACCACTCTGTCTTGGGGTGTTCAATGTTTTGCACGTTACCCAGATCACATGAAACAACACGATTTTTTCAAAAGCGCAATGCCAGAAGGTTACGTTCAAGAACGTACCATCTTCTTCAAAGATGACGGTAACTACAAAACCCGTGCGGAAGTGAAATTTGAAGGTGATACCCTGGTTAACCGTATCGAACTGAAAGGTATCGACTTCAAAGAGGATGGCAACATTCTGGGTCACAAACTGGAATACAACTACTTTAGCGATAACGTGTACATCACCGCAGATAAACAAAAAAACGGTATCAAGGCGAACTTCAAAATCCGTCACAACATTGAAGATGGTGGTGTTCAACTGGCAGATCACTACCAACAAAACACTCCAATTGGTGATGGTCCAGTTCTGCTGCCAGATAACCACTACCTGTCTACCCAATCTAAACTGTCTAAAGACCCAAACGAAAAACGTGATCACATGGTTCTGCTGGAATTTGTTACCGCAGCAGGTATTACCTTAGGTATGGATGAACTGTACAAAGGGTA |
| **pMC0_5a_13_CmVenus (Vn)** | GCTTTAGTTTCTAAAGGTGAAGAACTGTTTACCGGTGTTGTTCCAATTCTGGTTGAACTGGATGGTGATGTTAACGGTCACAAATTTTCTGTTTCTGGTGAAGGCGAAGGTGATGCAACCTACGGTAAACTGACCCTGAAACTGATTTGTACCACTGGTAAACTGCCAGTTCCATGGCCAACTCTGGTTACCACCTTAGGTTACGGTCTGCAATGTTTTGCACGTTACCCAGATCACATGAAACAACACGATTTTTTCAAAAGCGCAATGCCAGAAGGTTACGTTCAAGAACGTACCATCTTCTTCAAAGATGACGGTAACTACAAAACCCGTGCGGAAGTGAAATTTGAAGGTGATACCCTGGTTAACCGTATCGAACTGAAAGGTATCGACTTCAAAGAGGATGGCAACATTCTGGGTCACAAACTGGAATACAACTACAACTCTCACAACGTTTACATCACCGCAGATAAACAAAAAAACGGCATCAAAGCGAACTTCAAAATCCGTCACAACATTGAAGATGGTGGTGTTCAACTGGCAGATCACTACCAACAAAACACCCCAATTGGTGATGGTCCAGTTCTGCTGCCAGATAACCACTACCTGTCTTACCAATCTAAACTGAGCAAAGACCCAAACGAAAAACGTGATCACATGGTTCTGCTGGAATTTGTTACCGCAGCAGGTATTACCTTAGGTATGGATGAACTGTACAAAGGGTA |
| **pMC0_5a_14_CmScarlet-I (Vn)** | GCTTTAGTTTCTAAAGGTGAAGCAGTGATCAAAGAATTTATGCGCTTCAAAGTTCACATGGAAGGTTCTATGAACGGCCACGAATTTGAAATTGAAGGTGAAGGCGAAGGTCGTCCATACGAAGGTACTCAAACCGCAAAACTGAAAGTTACCAAAGGTGGTCCACTGCCATTTTCTTGGGATATTCTGTCTCCACAATTTATGTACGGTTCTCGTGCATTTATCAAACACCCAGCAGATATTCCAGACTACTACAAACAATCTTTTCCGGAAGGTTTCAAATGGGAACGTGTTATGAATTTTGAAGATGGTGGTGCAGTTACGGTTACCCAAGATACCTCTCTGGAAGATGGTACTCTGATCTACAAAGTTAAACTGCGTGGTACTAACTTTCCACCAGATGGTCCAGTTATGCAGAAAAAAACCATGGGTTGGGAAGCATCTACCGAACGTCTGTACCCAGAAGATGGCGTTCTGAAAGGTGATATCAAAATGGCACTGCGTCTGAAAGATGGCGGTCGTTACCTGGCAGATTTCAAAACCACCTACAAAGCGAAAAAACCAGTTCAAATGCCAGGTGCATACAACGTTGATCGTAAACTGGATATTACCAGCCACAACGAAGATTACACCGTTGTTGAACAATACGAACGTTCTGAAGGCCGTCACTCTACCGGTGGTATGGATGAACTGTACAAAGGGTA |
| **pMC0_5a_15_CmCherry (Vn)** | GCTTTAGTTTCTAAAGGTGAAGAGGATAACATGGCGATCATCAAAGAATTTATGCGCTTCAAAGTTCACATGGAAGGTTCTGTTAACGGCCACGAATTTGAAATTGAAGGTGAAGGCGAAGGTCGTCCATACGAAGGTACTCAAACCGCAAAACTGAAAGTTACCAAAGGTGGTCCACTGCCATTTGCATGGGATATTCTGTCTCCACAGTTTATGTACGGTAGCAAAGCATACGTTAAACACCCAGCAGATATTCCAGATTACCTGAAACTGTCTTTTCCGGAAGGTTTCAAATGGGAACGTGTTATGAATTTTGAAGATGGTGGTGTTGTTACGGTTACCCAAGATTCTTCTCTGCAAGATGGTGAGTTTATCTACAAAGTTAAACTGCGTGGCACCAACTTTCCATCTGATGGTCCAGTTATGCAGAAAAAAACCATGGGTTGGGAAGCATCTTCTGAACGTATGTACCCAGAAGATGGCGCACTGAAAGGTGAAATTAAACAACGTCTGAAACTTAAAGATGGCGGTCACTACGATGCAGAAGTTAAAACCACCTACAAAGCGAAAAAACCAGTTCAACTGCCAGGTGCATACAACGTTAACATTAAACTGGATATCACCAGCCACAACGAAGATTACACCATTGTTGAACAATACGAACGTGCAGAAGGCCGTCACTCTACCGGTGGTATGGATGAACTGTACAAAGGGTA |
| **pMC0_5a_16_CmKate-2 (Vn)** | GCTTTAGTTTCTGAACTGATTAAAGAAAACATGCACATGAAACTGTACATGGAAGGTACTGTTAACAACCACCACTTCAAATGTACCTCTGAAGGTGAAGGTAAACCATACGAAGGTACTCAAACCATGCGTATTAAAGCAGTTGAAGGTGGTCCACTGCCATTTGCATTTGATATTCTGGCAACCTCTTTTATGTACGGCAGCAAAACCTTTATCAACCACACTCAAGGTATCCCGGATTTTTTCAAACAAAGCTTTCCAGAAGGTTTCACCTGGGAACGTGTTACCACCTACGAAGATGGTGGTGTTCTGACCGCAACTCAAGATACCTCTCTGCAAGATGGTTGTCTGATCTACAACGTTAAAATCCGTGGTGTTAACTTTCCATCTAACGGTCCAGTTATGCAGAAAAAAACCTTAGGTTGGGAAGCATCTACCGAAACTCTGTACCCAGCGGATGGTGGTCTGGAAGGTCGTGCAGATATGGCACTGAAACTGGTTGGTGGTGGTCACCTGATTTGTAACCTGAAAACCACCTACCGTTCTAAAAAACCAGCGAAAAATCTGAAAATGCCAGGTGTTTACTACGTTGATCGTCGTCTGGAACGTATCAAAGAAGCAGATAAAGAAACCTACGTGGAACAACACGAAGTTGCAGTTGCACGTTACTGTGATCTGCCATCTAAACTGGGTCACCGTGGGTA |
| **pMC0_5a_17_CsfGFP(Vn)** | GCTTTACGTAAAGGTGAAGAACTGTTTACCGGTGTTGTTCCAATTCTGGTTGAACTGGATGGTGATGTTAACGGTCACAAATTTTCTGTTCGTGGTGAAGGCGAAGGTGATGCAACCAACGGTAAACTGACCCTGAAATTTATCTGTACCACTGGTAAACTGCCAGTTCCATGGCCAACTCTGGTTACCACTCTGACCTACGGTGTTCAATGTTTTGCACGTTACCCAGATCACATGAAACAACACGATTTTTTCAAAAGCGCAATGCCAGAAGGTTACGTTCAAGAACGTACCATCTCTTTTAAAGATGACGGCACCTACAAAACCCGTGCGGAAGTTAAATTTGAAGGTGATACCCTGGTTAACCGCATTGAACTGAAAGGCATCGATTTTAAAGAAGATGGTAACATCCTGGGCCACAAACTGGAATACAACTTTAACTCTCACAACGTGTACATCACCGCAGACAAACAAAAAAACGGTATCAAAGCGAACTTCAAGATCCGTCACAACGTTGAAGATGGTTCTGTTCAACTGGCAGATCACTACCAACAAAACACCCCAATTGGTGATGGTCCAGTTCTGCTGCCAGATAACCACTACCTGTCTACCCAAAGCGTTCTGTCTAAAGATCCAAACGAAAAACGTGATCACATGGTGCTGCTGGAATTTGTTACCGCAGCAGGTATTACCCACGGTATGGATGAACTGTACAAAGCAGGGTA |
| **pMC0_5b_01_TB0010** | GGTAACCAGGCATCAAATAAAACGAAAGGCTCAGTCGAAAGACTGGGCCTTTCGTTTTATCTGTTGTTTGTCGGTGAACGCTCTCCGCT |
| **pMC0_5b_02_TB0015** | GGTAACCAGGCATCAAATAAAACGAAAGGCTCAGTCGAAAGACTGGGCCTTTCGTTTTATCTGTTGTTTGTCGGTGAACGCTCTCTACTAGAGTCACACTGGCTCACCTTCGGGTGGGCCTTTCTGCGTTTATACGCT |
| **pMC0_5b_03_TB1002** | GGTAACGCAAAAAACCCCGCTTCGGCGGGGTTTTTTCGCCGCT |
| **pMC0_5b_04_TB1003** | GGTAACGCCAAAAACCCCGCTTCGGCGGGGTTTTTCCGCCGCT |
| **pMC0_5b_05_TB1004** | GGTAACGCCGAAAACCCCGCTTCGGCGGGGTTTTGCCGCCGCT |
| **pMC0_5b_06_TB1005** | GGTAACGCCGCAAACCCCGCTTCGGCGGGGTTTCGCCGCCGCT |
| **pMC0_5b_07_TB1006** | GGTAAAAAAAAAAACCCCGCCCCTGACAGGGCGGGGTTTTTTTTCGCT |
| **pMC0_5b_08_TB1007** | GGTAACGCAAAAAACCCCGCCCCTGACAGGGCGGGGTTTTTTCGCCGCT |
| **pMC0_5b_10_TB1009** | GGTAACGCCGAAAACCCCGCCCCTGACAGGGCGGGGTTTTGCCGCCGCT |
| **pMC0_5b_11_TB1010** | GGTAACGCCGCAAACCCCGCCCCTGACAGGGCGGGGTTTCGCCGCCGCT |

| | |
|---|---|
| **pMC0_6_01_6 Dropout** | CGCTAGAGACCGAAAGTGAAACGTGATTTCATGCGTCATTTTGAACATTTTGTAAATCTTATTTAATAATGTGTGCGGCAATTCACATTTAATTTAT<br>GAATGTTTTCTTAACATCGCGGCAACTCAAGAAACGGCAGGTTCGGATCTTAGCTACTAGAGAAAGAGGAGAAATACTAGATGCGTAAAGGCGA<br>AGAACTTTTCACTGGCGTAGTACCAATTTTAGTGGAGCTGGACGGAGATGTAAATGGTCATAAGTTTTCAGTTCGAGGAGAAGGCGAAGGAGAT<br>GCAACTAACGGTAAGCTGACACTAAAGTTTATCTGTACCACGGGTAAACTGCCGGTCCCATGGCCGACACTGGTTACTACACTTACTTATGGTGT<br>ACAATGCTTTGCTCGTTATCCTGACCATATGAAGCAGCACGATTTCTTCAAATCGGCCATGCCTGAAGGATACGTGCAAGAACGTACAATTAGCTT<br>TAAAGATGACGGCACCTATAAAACGCGCGCCGAAGTTAAATTCGAGGGTGACACATTGGTTAATCGTATAGAACTTAAGGGCATTGACTTCAAA<br>GAAGATGGCAACATCCTTGGCCATAAACTGGAATATAATTTTAACTCTCACAATGTCTACATTACGGCGGATAAACAAAAGAATGGCATTAAAGC<br>GAATTTCAAAATCCGCCACAATGTCGAAGACGGCTCGGTTCAACTGGCGGACCATTATCAGCAGAACACCCCAATCGGTGATGGCCCGGTTCTGT<br>TACCTGATAATCATTACCTTTCTACTCAAAGCGTTTTATCTAAAGATCCTAACGAGAAGCGTGATCATATGGTTCTACTGGAATTTGTTACCGCAGC<br>TGGTATCACGCACGGCATGGATGAGCTGTATAAATGACCAGGCATCAAATAAAACGAAAGGCTCAGTCGAAAGACTGGGCCTTTCGTTTTATCT<br>GTTGTTTGTCGGTGAACGCTCTCTACTAGAGTCACACTGGCTCACCTTCGGGTGGGCCTTTCTGCGTTTATAGGTCTCTAGCT |
| **pMC0_6_02_3C1SF** | CGCTTACTGGAGACGAGCT |
| **pMC0_6_03_3C1CLR [#]** | CGCTGGACCAAAACGAAAAAAGACGCTTTTCAGCGTCTTATTGTTCGTCTTTGGTACCGAGCTGTGTCTGTGGCTGAGAGAGCAGGTACAAGATA<br>AACCCGTCTATATTCTCAAATTCTTTCACTATCGCACTTGTTTTATGTGAAGTCCAGATATGGACTCTGGAACCCAACTCGGAGCTCTGCGGGTAG<br>AGCGTCAACCATTCGAACGTTGCGCGGTCTTCGGAGAACCGTAAGAGGTAATATACTCCGATATTCATTTCAGCGGTAGCTGAGTCCGTCAGTTG<br>ATACGGTGCGTACCCGGAGCAAATAGGAGAGGCGGTTGTAGTGGGGGGAATGGATAGCAAGCTGCGGGTAGACCTCCAATTATTGAAGGCCTC<br>CCAAATCGGGGGGCCTTTTTTATTGATAACAAAACTCCGGAGACGAGCT |
| **pMC0_6_04_3C1RLR [#]** | CGCTGGACCAAAACGAAAAAAGACGCTTTTCAGCGTCTTATTGTTCGTCTTTGGTACCGAGCTGTGTCTGTGGCTGAGAGAGCAGGTACAAGATA<br>AACCCGTCTATATTCTCAAATTCTTTCACTATCGCACTTGTTTTATGTGAAGTCCAGATATGGACTCTGGAACCCAACTCGGAGCTCTGCGGGTAG<br>AGCGTCAACCATTCGAACGTTGCGCGGTCTTCGGAGAACCGTAAGAGGTAATATACTCCGATATTCATTTCAGCGGTAGCTGAGTCCGTCAGTTG<br>ATACGGTGCGTACCCGGAGCAAATAGGAGAGGCGGTTGTAGTGGGGGGAATGGATAGCAAGCTGCGGGTAGACCTCCAATTATTGAAGGCCTC<br>CCAAATCGGGGGGCCTTTTTTATTGATAACAAAATGTTGGAGACGAGCT |
| **pMC0_6_05_3C1CSR** | CGCTCTCCGGAGACGAGCT |
| **pMC0_6_06_3C1RSR** | CGCTTGTTGGAGACGAGCT |
| **pMC0_6_07_3C2SN** | CGCTAATGGGAGACGAGCT |
| **pMC0_6_08_3C2LR [#]** | CGCTGGACCAAAACGAAAAAAGGCCGCTTTCGCGGCCTCTTTTCTGGAATTTGGTACCGAGTCGACACCCTACCGTATCGCTGTGGCAGGCACAG<br>GATCTAACCGGTCCTGGACCGGCGTAAGTTGCTGTACAGCCAAATCTTCCGTACTAATTCTACAGGATTTGTCAACTGCAAACGAATTTCTAGTCG<br>GAACGGATCACGTGCACCGGGGTATGCCGTGGTTGAATATCGGCACATGAGGGGCTCGTGGGACACTTCCAGAACTGTCTGGCGTGCGAAGTC<br>TCTGCTAAATGCAAAACAGTCAGGACTCGTTACAGCTCGTGTATGGGAGCTCCGGGTAGGCCGATCGTCAATTGCGGCTCGGTACCAAATTTTCG<br>AAAAAAGACGCTGAAAAGCGTCTTTTTTCGTTTTGGTCCAGTAGGAGACGAGCT |
| **pMC0_6_09_3C2SR** | CGCTAGTAGGAGACGAGCT |
| **pMC0_6_10_3C3SN** | CGCTGCTTGGAGACGAGCT |
| **pMC0_6_11_3C3LR [#]** | CGCTGGACCAAAACGAAAAAAGACGCTTTTCAGCGTCTTCAATAATTGGTTTGGTACCGAGATTCGTTGTTGACTTCATCGTCACGATAAAGCAC<br>GACAAGTCCCAGCCTGTGAGTGCGACCCGAAATTGTAGGTAGACACTCACGCTCCCCCCGATATCAGACAGCGTGCTAATGAAAGCCTTCCGATC<br>TTCGGATAAGGCCCAAGCTCAGCACATGAACGGATCAGTGGTAGGTGTCTCACAACCCAATCGTGGTGACAATACGAATGTCCAAATATTGCCTC<br>TTGGCGGGTAGATTCCGTTACACTTTACAGCCATCATAACAACCAGATGAGAATCGATGGTATTCTTGGACCCGCGCCAATTATTGAAGGCCTCC<br>CAAATCGGGGGGCCTTTTTTATTGATAACAAAACATTGGAGACGAGCT |
| **pMC0_6_12_3C3SR** | CGCTCATTGGAGACGAGCT |
| **pMC0_6_13_3C4SF** | CGCTGGTAGGAGACGAGCT |
| **pMC0_6_14_3C4LR** | CGCTGGACCAAAACGAAAAAAGACGCTTTTCAGCGTCTTTTTAGTTAGATTTGGTACCGAGTTTATGGGCACGCTTCCGCGTTAATCATACCTCTA<br>TTCACACGGTCCCCCACCAGTGTGTCTAAAGACGTACCCATTGAACGCGGGATTGGTTGTGATAGCTGGTTGTGCCACTAAGTCCTAGTCAAACG<br>AGGCTGGCATCCCATAGCAGCAAAATCGGGTAGTAACCCTCAAAGACGTGCCCGACAGTTCATCTTCCTGTAAGAACTATTGGCTTACGCACGAG<br>TCTGTTCGTGTATCGTCTAGCGGTTAGATTCACCTGAATTGCTATGCTTCTTTCAATGGATCCTGCCCACTTCACTTTTCGAAAAAAGGCCTCCCAA<br>ATCGGGGGGCCTTTTTTATTGATAACAAAAAAGCGGAGACGAGCT |
| **pMC0_6_15_3C4SR** | CGCTAAGCGGAGACGAGCT |
| **pMC0_6_16_3C5CSF** | CGCTCGCTGGAGACGAGCT |
| **pMC0_6_17_3C5OSF** | CGCTAGCTGGAGACGAGCT |
| **pMC0_6_18_3C5LR [#]** | CGCTTTTTGTTATCAATAAAAAAGGCCCCCCGATTTGGGAGGCCTTTTTTTTTGGTGGTCGCATGAAGCAGGCCTAGTATTCCAACGTGGTGGCCA<br>ATTAACAATATTCCAACCCCAATCTGTAAAAAACTGTGCCCTACGCAGGACTCAGAGAACGGCGGAAGGTAGGTACTCGTCTTCTAACCCGACGC<br>ATTAAGGCTACAGGGTGCGAAGGCCATTATTGCAAACTTTTTGTTCTCAACAACGTAAATCTACCGCACTTCAGGTGTCGTTCTTGATGTACGAAG<br>ACGGTAGGCCTATGGATTGCCTTCTGTATCAGCAATTTTTCATCTCCTTCCAAGCGGCATCCATAATCTAACTAAAAAGGCCTCCCAAATCGGGGG<br>GCCTTTTTTATTGATAACAAAATACCGGAGACGAGCT |
| **pMC0_6_19_3C5SR** | CGCTTACCGGAGACGAGCT |
| **pMC0_6_20_3C0SF** | CGCTTACTAGTAGCGGCCGCTGCAGAGCT |
| **pMC0_6_21_6 Dropout mScarlet-I (Vn)** | CGCTAGAGACCGAAAGTGAAACGTGATTTCATGCGTCATTTTGAACATTTTGTAAATCTTATTTAATAATGTGTGCGGCAATTCACATTTAATTTAT<br>GAATGTTTTCTTAACATCGCGGCAACTCAAGAAACGGCAGGTTCGGATCTTAGCTACTAGAGAAAGAGGAGAAATACTAGATGGTTTCTAAAGG<br>TGAAGCAGTGATCAAAGAATTTATGCGCTTCAAAGTTCACATGGAAGGTTCTATGAACGGCCACGAATTTGAAATTGAAGGTGAAGGCGAAGGT<br>CGTCCATACGAAGGTACTCAAACCGCAAAACTGAAAGTTACCAAAGGTGGTCCACTGCCATTTTCTTGGGATATTCTGTCTCCACAATTTATGTAC<br>GGTTCTCGTGCATTTATCAAACACCCAGCAGATATTCCAGACTACTACAAACAATCTTTTCCGGAAGGTTTCAAATGGGAACGTGTTATGAATTTT<br>GAAGATGGTGGTGCAGTTACGGTTACCCAAGATACCTCTCTGGAAGATGGTACTCTGATCTACAAAGTTAAACTGCGTGGTACTAACTTTCCACC<br>AGATGGTCCAGTTATGCAGAAAAAAACCATGGGTTGGGAAGCATCTACCGAACGTCTGTACCCAGAAGATGGCGTTCTGAAAGGTGATATCAAA<br>ATGGCACTGCGTCTGAAAGATGGCGGTCGTTACCTGGCAGATTTCAAAACCACCTACAAAGCGAAAAAACCAGTTCAAATGCCAGGTGCATACA<br>ACGTTGATCGTAAACTGGATATTACCAGCCACAACGAAGATTACACCGTTGTTGAACAATACGAACGTTCTGAAGGCCGTCACTCTACCGGTGGT<br>ATGGATGAACTGTACAAATAACCAGGCATCAAATAAAACGAAAGGCTCAGTCGAAAGACTGGGCCTTTCGTTTTATCTGTTGTTTGTCGGTGAAC<br>GCTCTCTACTAGAGTCACACTGGCTCACCTTCGGGTGGGCCTTTCTGCGTTTATAGGTCTCTAGCT |
| **pMC0_7_01_OColE1** | AGCTGAACCGTAAAAAGGCCGCGTTGCTGGCGTTTTTCCACAGGCTCCGCCCCCCTGACGAGCATCACAAAAATCGACGCTCAAGTCAGAGGTG<br>GCGAAACCCGACAGGACTATAAAGATACCAGGCGTTTCCCCCTGGAAGCTCCCTCGTGCGCTCTCCTGTTCCGACCCTGCCGCTTACCGGATACCT<br>GTCCGCCTTTCTCCCTTCGGGAAGCGTGGCGCTTTCTCATAGCTCACGCTGTAGGTATCTCAGTTCGGTGTAGGTCGTTCGCTCCAAGCTGGGCTG<br>TGTGCACGAACCCCCCGTTCAGCCCGACCGCTGCGCCTTATCCGGTAACTATCGTCTTGAGTCCAACCCGGTAAGACACGACTTATCGCCACTGGC<br>AGCAGCCACTGGTAACAGGATTAGCAGAGCGAGGTATGTAGGCGGTGCTACAGAGTTCTTGAAGTGGTGGCCTAACTACGGCTACACTAGAAG |

| | |
|---|---|
| | AACAGTATTTGGTATCTGCGCTCTGCTGAAGCCAGTTACCTTCGGAAAAAGAGTTGGTAGCTCTTGATCCGGCAAACAAACCACCGCTGGTAGCG<br>GTGGTTTTTTTGTTTGCAAGCAGCAGATTACGCGCAGAAAAAAAGGATCTCAAGAAGATCCTTTGATCTTTTCTACGGGGTCTGACGCTCAGTGG<br>AACGAAAACTCACGTTAAGGGATTTTGGTCATGAGATTATCAAAAAGGATCTTCACTGCT |
| **pMC0_7_02_Op15A** | AGCTGCGCTAGCGGAGTGTATACTGGCTTACTATGTTGGCACTGATGAGGGTGTCAGTGAAGTGCTTCATGTGGCAGGAGAAAAAAGGCTGCA<br>CCGGTGCGTCAGCAGAATATGTGATACAGGATATATTCCGCTTCCTCGCTCACTGACTCGCTACGCTCGGTCGTTCGACTGCGGCGAGCGGAAAT<br>GGCTTACGAACGGGGCGGAGATTTCCTGGAAGATGCCAGGAAGATACTTAACAGGGAAGTGAGAGGGCCGCGGCAAAGCCGTTTTTCCATAGG<br>CTCCGCCCCCCTGACAAGCATCACGAAATCTGACGCTCAAATCAGTGGTGGCGAAACCCGACAGGACTATAAAGATACCAGGCGTTTCCCCTGGC<br>GGCTCCCTCGTGCGCTCTCCTGTTCCTGCCTTTCGGTTTACCGGTGTCATTCCGCTGTTATGGCCGCGTTTGTCTCATTCCACGCCTGACACTCAGT<br>TCCGGGTAGGCAGTTCGCTCCAAGCTGGACTGTATGCACGAACCCCCCGTTCAGTCCGACCGCTGCGCCTTATCCGGTAACTATCGTCTTGAGTC<br>CAACCCGGAAAGACATGCAAAAGCACCACTGGCAGCAGCCACTGGTAATTGATTTAGAGGAGTTAGTCTTGAAGTCATGCGCCGGTTAAGGCTA<br>AACTGAAAGGACAAGTTTTGGTGACTGCGCTCCTCCAAGCCAGTTACCTCGGTTCAAAGAGTTGGTAGCTCAGAGAACCTTCGAAAAACCGCCCT<br>GCAAGGCGGTTTTTTCGTTTTCAGAGCAAGAGATTACGCGCAGACCAAAACGATCTCAAGAAGATCATCTTATTAAGGGGTCTGACGCTCAGTG<br>GAACGAAAACTCACGTTAAGGGATTTTGGTCATGAGATTATCAAAAAGGATCTTCACCTAGATCCTTTTAAATTTGCT |
| **pMC0_7_04_OpMB1-M** | AGCTACGGTTATCCACAGAATCAGGGGATAACGCAGGAAAGAACATGTGAGCAAAAGGCCAGCAAAAGGCCAGGAACCGTAAAAAGGCCGCG<br>TTGCTGGCGTTTTTCCATAGGCTCCGCCCCCCTGACGAGCATCACAAAAATCGACGCTCAAGTCAGAGGTGGCGAAACCCGACAGGACTATAAA<br>GATACCAGGCGTTTCCCCCTGGAAGCTCCCTCGTGCGCTCTCCTGTTCCGACCCTGCCGCTTACCGGATACCTGTCCGCCTTTCTCCCTTCGGGAA<br>GCGTGGCGCTTTCTCATAGCTCACGCTGTAGGTATCTCAGTTCGGTGTAGGTCGTTCGCTCCAAGCTGGGCTGTGTGCACGAACCCCCCGTTCAG<br>CCCGACCGCTGCGCCTTATCCGGTAACTATCGTCTTGAGTCCAACCCGGTAAGACACGACTTATCGCCACTGGCAGCAGCCACTGGTAACAGGAT<br>TAGCAGAGCGAGGTATGTAGGCGGTGCTACAGAGTTCTTGAAGTGGTGGCCTAACTACGGCTACACTAGAAGAACAGTATTTGGTATCTGCGCT<br>CTGCTGAAGCCAGTTACCTTCGGAAAAAGAGTTGGTAGCTCTTGATCCGGCAAACAAACCACCGCTGGTAGCGGTGGTTTTTTTGTTTGCAAGCA<br>GCAGATTACGCGCAGAAAAAAAGGATCTCAAGAAGATCCTTTGATCTTTTCTACGGGGTCTGACGCTCAGTGGAACGAAAACTCACGTTAAGGG<br>ATTTTGGTCATGATGCT |
| **pMC0_7_05_ORSF1010** | AGCTTTCTGAAAGCGACCAGGTGCTCGGCGTGGCAAGACTCGCAGCGAACCCGTAGAAAGCCATGCTCCAGCCGCCCGCATTGGAGAAATTCTT<br>CAAATTCCCGTTGCACATAGCCCGGCAATTCCTTTCCCTGCTCTGCCATAAGCGCAGCGAATGCCGGGTAATACTCGTCAACGATCTGATAGAGA<br>AGGGTTTGCTCGGGTCGGTGGCTCTGGTAACGACCAGTATCCCGATCCCGGCTGGCCGTCCTGGCCGCCACATGAGGCATGTTCCGCGTCCTTGC<br>AATACTGTGTTTACATACAGTCTATCGCTTAGCGGAAAGTTCTTTTACCCTCAGCCGAAATGCCTGCCGTTGCTAGACATTGCCAGCCAGTGCCCG<br>TCACTCCCGTACTAACTGTCACGAACCCCTGCAATAACTGTCACGCCCCCCTGCAATAACTGTCACGAACCCCTGCAATAACTGTCACGCCCCCAA<br>ACCTGCAAACCCAGCAGGGGCGGGGGCTGGCGGGGTGTTGGAAAAATCCATCCATGATTATCTAAGAATAATCCACTAGGCGCGGTTATCAGC<br>GCCCTTGTGGGGCGCTGCTGCCCTTGCCCAATATGCCCGGCCAGAGGCCGGATAGCTGGTCTATTCGCTGCGCTAGGCTACACACCGCCCCACCG<br>CTGCGCGGCAGGGGGAAAGGCGGGCAAAGCCCGCTAAACCCCACACCAAACCCCGCAGAAATACGCTGGAGCGCTTTTAGCCGCTTTAGCGGC<br>CTTTCCCCCTACCCGAAGGGTGGGGGCGCGTGTGCAGCCCCGCAGGGCCTGTCTCGGTCGATCATTCAGCCCGGCTCATCCTTCTGGCGTGGCG<br>GCAGACCGAACAAGGCGCGGTCGTGGTCGCGTTCAAGGTACGCATCCATTGCCGCCATGAGCCGATCCTCCGGCCACTCGCTGCTGTTCACCTTG<br>GCCAAAATCATGGCCCCCACCAGCACCTTGCGCCTTGTTTCGTTCTTGCGCTCTTGCTGCTGTTCCCTTGCCCGCACCCGCTGAATTTCGGCATTGA<br>TTCGCGCTCGTTGTTCTTCGAGCTTGGCCAGCCGATCCGCCGCCTTGTTGCTCCCCTTAACCATCTTGACACCCCATTGTTAATGTGCTGTCTCGTA<br>GGCTATCATGGAGGCACAGCGGCGGCAATCCCGACCCTACTTTGTAGGGGAGGGCGCACTTACCGGTTTCTCTTCGAGAAACTGGCCTAACGGC<br>CACCCTTCGGGCGGTGCGCTCTCCGAGGGCCATTGCATGGAGCCGAAAAGCAAAAGCAACAGCGAGGCAGCATGGCGATTTATCACCTTACGG<br>CGAAAACCGGCAGCAGGTCGGGCGGCCAATCGGCCAGGGCCAAGGCCGACTACATCCAGCGCGAAGGCAAGTATGCCCGCGACATGGATGAA<br>GTCTTGCACGCCGAATCCGGGCACATGCCGGAGTTCGTCGAGCGGCCCGCCGACTACTGGGATGCTGCCGACCTGTATGAACGCGCCAATGGGC<br>GGCTGTTCAAGGAGGTCGAATTTGCCCTGCCGGTCGAGCTGACCCTCGACCAGCAGAAGGCGCTGGCGTCCGAGTTCGCCCAGCACCTGACCGG<br>TGCCGAGCGCCTGCCGTATACGCTGGCCATCCATGCCGGTGGCGGCGAGAACCCGCACTGCCACCTGATGATCTCCGAGCGGATCAATGACGGC<br>ATCGAGCGGCCCGCCGCTCAGTGGTTCAAGCGGTACAACGGCAAGACCCCGGAGAAGGGCGGGGCACAGAAGACCGAAGCGCTCAAGCCCAA<br>GGCATGGCTTGAGCAGACCCGCGAGGCATGGGCCGACCATGCCAACCGGGCATTAGAGCGGGCTGGCCACGACGCCCGCATTGACCACAGAAC<br>ACTTGAGGCGCAGGGCATCGAGCGCCTGCCCGGTGTTCACCTGGGGCCGAACGTGGTGGAGATGGAAGGCCGGGGCATCCGCACCGACCGGG<br>CAGACGTGGCCCTGAACATCGACACCGCCAACGCCCAGATCATCGACTTACAGGAATACCGGGAGGCAATAGACCATGAACGCAATCGACAGA<br>GTGAAGAAATCCAGAGGCATCAACGAGTTAGCGGAGCAGATCGAACCGCTGGCCCAGAGCATGGCGACACTGGCCGACGAAGCCCGGCAGGT<br>CATGAGCCAGACCAAGCAGGCCAGCGAGGCGCAGGCGGCGGAGTGGCTGAAAGCCCAGCGCCAGACAGGGGCGGCATGGGTGGAGCTGGCC<br>AAAGAGTTGCGGGAGGTAGCCGCCGAGGTGAGCAGCGCCGCGCAGAGCGCCCGGAGCGCGTCGCGGGGGTGGCACTGGAAGCTATGGCTAA<br>CCGTGATGCTGGCTTCCATGATGCCTACGGTGGTGCTGCTGATCGCATCGTTGCTCTTGCTCGACCTGACGCCACTGACAACCGAGGACGGCTCG<br>ATCTGGCTGCGCTTGGTGGCCCGATGAAGAACGACAGGACTTTGCAGGCCATAGGCCGACAGCTCAAGGCCATGGGCTGTGAGCGCTTCGATAT<br>CGGCGTCAGGGACGCCACCACCGGCCAGATGATGAACCGGGAATGGTCAGCCGCCGAAGTGCTCCAGAACACGCCATGGCTCAAGCGGATGAA<br>TGCCCAGGGCAATGACGTGTATATCAGGCCCGCCGAGCAGGAGCGGCATGGTCTGGTGCTGGTGGACGACCTCAGCGAGTTTGACCTGGATGA<br>CATGAAAGCCGAGGGCCGGGAGCCTGCCCTGGTAGTGGAAACCAGCCCGAAGAACTATCAGGCATGGGTCAAGGTGGCCGACGCCGCAGGCG<br>GTGAACTTCGGGGGCAGATTGCCCGGACGCTGGCCAGCGAGTACGACGCCGACCCGGCCAGCGCCGACAGCCGCCACTATGGCCGCTTGGCGG<br>GCTTCACCAACCGCAAGGACAAGCACACCACCCGCGCCGGTTATCAGCCGTGGGTGCTGCTGCGTGAATCCAAGGGCAAGACCGCCACCGCTGG<br>CCCGGCGCTGGTGCAGCAGGCTGGCCAGCAGATCGAGCAGGCCCAGCGGCAGCAGGAGAAGGCCCGCAGGCTGGCCAGCCTCGAACTGCCCG<br>AGCGGCAGCTTAGCCGCCACCGGCGCACGGCGCTGGACGAGTACCGCAGCGAGATGGCCGGGCTGGTCAAGCGCTTCGGTGATGACCTCAGCA<br>AGTGCGACTTTATCGCCGCGCAGAAGCTGGCCAGCCGGGGCCGCAGTGCCGAGGAAATCGGCAAGGCCATGGCCGAGGCCAGCCCAGCGCTG<br>GCAGAGCGCAAGCCCGGCCACGAAGCGGATTACATCGAGCGCACCGTCAGCAAGGTCATGGGTCTGCCCAGCGTCCAGCTTGCGCGGGCCGAG<br>CTGGCACGGGCACCGGCACCCCGCCAGCGAGGCATGGACAGGGGCGGGCCAGATTTCAGCATGTAGTGCTTGCGTTGGTACTCACGCCTGTTAT<br>ACTATGAGTACTCACGCACAGAAGGGGGTTTTATGGAATACGAAAAAAGCGCTTCAGGGTCGGTCTACCTGATCAAAAGTGACAAGGGCTATTG<br>GTTGCCCGGTGGCTTTGGTTATACGTCAAACAAGGCCGAGGCTGGCCGCTTTTCAGTCGCTGATATGGCCAGCCTTAACCTTGACGGCTGCACCT<br>TGTCCTTGTTCCGCGAAGACAAGCCTTTCGGCCCCGGCAAGTTTCTCGGTGACTGATATGAAAGACCAAAAGGACAAGCAGACCGGCGACCTGC<br>TGGCCAGCCCTGACGCTGTACGCCAAGCGCGATATGCCGAGCGCATGAAGGCCAAAGGGATGCGTCAGCGCAAGTTCTGGCTGACCGACGACG<br>AATACGAGGCGCTGCGCGAGTGCCTGGAAGAACTCAGAGCGGCGCAGGGCGGGGGTAGTGACCCCGCCAGCGCCTAACCACCAACTGCCTGC<br>AAAGGAGGCAATCAATGGCTACCCATAAGCCTATCAATATTCTGGAGGCGTTCGCAGCAGCGCCGCCACCGCTGGACTACGTTTTGCCCAACATG<br>GTGGCCGGTACGGTCGGGGCGCTGGTGTCGCCCGGTGGTGCCGGTAAATCCATGCTGGCCCTGCAACTGGCCGCACAGATTGCAGGCGGGCCG<br>GATCTGCTGGAGGTGGGCGAACTGCCCACCGGCCCGGTGATCTACCTGCCCGCCGAAGACCCGCCCACCGCCATTCATCACCGCCTGCACGCCCT<br>TGGGGCGCACCTCAGCGCCGAGGAACGGCAAGCCGTGGCTGACGGCCTGCTGATCCAGCCGCTGATCGGCAGCCTGCCCAACATCATGGCCCC<br>GGAGTGGTTCGACGGCCTCAAGCGCGCCGCCGAGGGCCGCCGCCTGATGGTGCTGGACACGCTGCGCCGGTTCCACATCGAGGAAGAAAACGC<br>CAGCGGCCCCATGGCCCAGGTCATCGGTCGCATGGAGGCCATCGCCGCCGATACCGGGTGCTCTATCGTGTTCCTGCACCATGCCAGCAAGGGC<br>GCGGCCATGATGGGCGCAGGCGACCAGCAGCAGGCCAGCCGGGGCAGCTCGGTACTGGTCGATAACATCCGCTGGCAGTCCTACCTGTCGAGC<br>ATGACCAGCGCCGAGGCCGAGGAATGGGGTGTGGACGACGACCAGCGCCGGTTCTTCGTCCGCTTCGGTGTGAGCAAGGCCAACTATGGCGCA<br>CCGTTCGCTGATCGGTGGTTCAGGCGGCATGACGGCGGGGTGCTCAAGCCCGCCGTGCTGGAGAGGCAGCGCAAGAGCAAGGGGGTGCCCCG<br>TGGTGAAGCCTAAGAACAAGCACAGCCTCAGCCACGTCCGGCACGACCCGGCGCACTGTCTGGCCCCCGGCCTGTTCCGTGCCCTCAAGCGGGG<br>CGAGCGCAAGCGCAGCAAGCTGGACGTGACGTATGACTACGGCGACGGCAAGCGGATCGAGTTCAGCGGCCCGGAGCCGCTGGGCGCTGATG<br>ATCTGCGCATCCTGCAAGGGCTGGTGGCCATGGCTGGGCCTAATGGCCTAGTGCTTGGCCCGGAACCCAAGACCGAAGGCGGACGGCAGCTCC<br>GGCTGTTCCTGGAACCCAAGTGGGAGGCCGTCACCGCTGATGCCATGGTGGTCAAAGGTAGCTATCGGGCGCTGGCAAAGGAAATCGGGGCAG<br>AGGTCGATAGTGGTGGGGCGCTCAAGCACATACAGGACTGCATCGAGCGCCTTTGGAAGGTATCCATCATCGCCCAGAATGGCCGCAAGCGGC<br>AGGGGTTTCGGCTGCTGTCGGAGTACGCCAGCGACGAGGCGGACGGGCGCCTGTACGTGGCCCTGAACCCCTTGATCGCGCAGGCCGTCATGG<br>GTGGCGGCCAGCATGTGCGCATCAGCATGGACGAGGTGCGGGCGCTGGACAGCGAAACCGCCCGCCTGCTGCACCAGCGGCTGTGTGGCTGG<br>ATCGACCCCGGCAAAACCGGCAAGGCTTCCATAGATACCTTGTGCGGCTATGTCTGGCCGTCAGAGGCCAGTGGTTCGACCATGCGCAAGCGCC<br>GCCAGCGGGTGCGCGAGGCGTTGCCGGAGCTGGTCGCGCTGGGCTGGACGGTAACCGAGTTCGCGGCGGGCAAGTACGACATCACCCGGCCC<br>AAGGCGGCAGGCTGACCCCCCCCACTCTATTGTAAACAAGACATTTTTATCTTTTATATTCAATGGCTTATTTTCCTGCTAATTGGTAATACCATGA |

| | |
|---|---|
| | AAAATACCATGCTCAGAAAAGGCTTAACAATATTTGAAAAATTGCCTACTGAGCGCTGCCGCACAGCTCCATAGGCCGCTTTCCTGGCTTTGCTT<br>CCAGATGTATGCTCTTCTGCTCCCGAACGCCAGCAAGACGTAGCCCAGCGCGTCGGCCAGCTTGCAATTCGCGCTAACTTACATTAATTGCGTTGC<br>GCTGCT |
| **pMC0_8_09_Akan (sfgfp)(Vn)** | TGCTAAACGGGCAAGGTGTCACCACCCTGCCCTTTTTCTTTAAAACCGAAAAGATTACTTCGCGTTATGCAGGCTTCCTCGCTCACTGACTCGCTG<br>CGCTCGGTCGTTCGGCTGCGGCGAGCGGTATCAGCTCACTCAAAGGCGGTAATCTCGAGTCCCGTCAAGTCAGCGTAATGCTCTGCCAGTGTTAC<br>AACCAATTAACCAATTCTGATTAGAAAAACTCATCGAGCATCAAATGAAACTGCAATTTATTCATATCAGGATTATCAATACCATATTTTTGAAAAA<br>GCCGTTTCTGTAATGAAGGAGAAAACTCACCGAGGCAGTTCCATAGGATGGCAAGATCCTGGTATCGGTCTGCGATTCCGACTCGTCCAACATCA<br>ATACAACCTATTAATTTCCCCTCGTCAAAAATAAGGTTATCAAGTGAGAAATCACCATGAGTGACGACTGAATCCGGTGAGAATGGCAAAAGCTT<br>ATGCATTTCTTTCCAGACTTGTTCAACAGGCCAGCCATTACGCTCGTCATCAAAATCACTCGCATCAACCAAACCGTTATTCATTCGTGATTGCGCC<br>TGAGCCAGACGAAATACGCGATCGCTGTTAAAAGGACAATTACAAACAGGAATCGAATGCAACCGGCGCAGGAACACTGCCAGCGCATCAACA<br>ATATTTTCACCTGAATCAGGATATTCTTCTAATACCTGGAATGCTGTTTTCCCGGGGATCGCAGTGGTGAGTAACCATGCATCATCAGGAGTACGG<br>ATAAAATGCTTGATGGTCGGAAGAGGCATAAATTCCGTCAGCCAGTTTAGTCTGACCATCTCATCTGTAACATCATTGGCAACGCTACCTTTGCCA<br>TGTTTCAGAAACAACTCTGGCGCATCGGGCTTCCCATACAATCGATAGATTGTCGCACCTGATTGCCCGACATTATCGCGAGCCCATTTATACCCA<br>TATAAATCAGCATCCATGTTGGAATTTAATCGCGGCCTCGAGCAAGACGTTTCCCGTTGAATATGGCTCATAACACCCCTTGTATTACTGTTTATGT<br>AAGCAGACAGTTTTATTGTTCATGATGATATATTTTTATCTTGTGCAATGTAACATCAGAGATTTTGAGACACAACGTGGCTTTGTTGAATAAATC<br>GAACTTTTGCTGAGTTGAAGGATCAGATCACGCATCTTAACA |
| **pMC0_8_10_Akan (mScarlet-I)(Vn)** | TGCTAAACGGGCAAGGTGTCACCACCCTGCCCTTTTTCTTTAAAACCGAAAAGATTACTTCGCGTTATGCAGGCTTCCTCGCTCACTGACTCGCTG<br>CGCTCGGTCGTTCGGCTGCGGCGAGCGGTATCAGCTCACTCAAAGGCGGTAATCTCGAGTCCCGTCAAGTCAGCGTAATGCTCTGCCAGTGTTAC<br>AACCAATTAACCAATTCTGATTAGAAAAACTCATCGAGCATCAAATGAAACTGCAATTTATTCATATCAGGATTATCAATACCATATTTTTGAAAAA<br>GCCGTTTCTGTAATGAAGGAGAAAACTCACCGAGGCAGTTCCATAGGATGGCAAGATCCTGGTATCGGTCTGCGATTCCGACTCGTCCAACATCA<br>ATACAACCTATTAATTTCCCCTCGTCAAAAATAAGGTTATCAAGTGAGAAATCACCATGAGTGACGACTGAATCCGGTGAGAATGGCAAAAGCTT<br>ATGCATTTCTTTCCAGACTTGTTCAACAGGCCAGCCATTACGCTCGTCATCAAAATCACTCGCATCAACCAAACCGTTATTCATTCGTGATTGCGCC<br>TGAGCCAGACGAAATACGCGATCGCTGTTAAAAGGACAATTACAAACAGGAATCGAATGCAACCGGCGCAGGAACACTGCCAGCGCATCAACA<br>ATATTTTCACCTGAATCAGGATATTCTTCTAATACCTGGAATGCTGTTTTCCCGGGGATCGCAGTGGTGAGTAACCATGCATCATCAGGAGTACGG<br>ATAAAATGCTTGATGGTCGGAAGAGGCATAAATTCCGTCAGCCAGTTTAGTCTGACCATCTCATCTGTAACATCATTGGCAACGCTACCTTTGCCA<br>TGTTTCAGAAACAACTCTGGCGCATCGGGCTTCCCATACAATCGATAGATTGTCGCACCTGATTGCCCGACATTATCGCGAGCCCATTTATACCCA<br>TATAAATCAGCATCCATGTTGGAATTTAATCGCGGCCTCGAGCAAGACGTTTCCCGTTGAATATGGCTCATAACACCCCTTGTATTACTGTTTATGT<br>AAGCAGACAGTTTTATTGTTCATGATGATATATTTTTATCTTGTGCAATGTAACATCAGAGATTTTGAGACACAACGTGGCTTTGTTGAATAAATC<br>GAACTTTTGCTGAGTTGAAGGATCAGATCACGCATCTTAACA |
| **pMC0_8_11_Atet (sfgfp)(Vn)** | TGCTGAGTCACTAAGGGCTAACTAACTAATTACGTAGCAATCAACTCACTGGCTCACCTTCACGGGTGGGCCTTTCTTCGGCACGGGCAAATTGC<br>TGAATATTCCTTTTCTTAGACGTCAGGTGGCAAACGCGAAGTAATCTTTTCGGTTTTAAAGAAAAAGGGCAGGGTGGTGACACCTTGCCCGTTTTT<br>TTGCCGGAGCTAGTATTATTAGGTCGAGGTGGCCCGGCTCCATGCACCGCGACGCAACGCGGGGAGGCAGACAAGGTAAAGGGCGGCACCTAC<br>AATCCATGCCAACCCGTTCCATGTGCTCGCCGAGGCGGCATAAATCGCCGTGACGATCAGCGGTCCAATGATCGAAGTGAGGCTGGTAAGAGCC<br>GCGAGCGACCCTTGAAGCTGTCCCTGATGGTCGTCATCTACTTGGCGGGACAGCATGGCCTGCAACGCGGGCATACCGATGCCGCCGGAAGCG<br>AGAAGAATCATAATGGGGAAGGCCATCCAGCCGCGCGTCGCGAACGCCAGCAAGACGTAGCCCAGCGCGTCGGCCGCCATGCCCGCGATAATG<br>GCCTGCTTCTCGCCGAAACGTTTGGTGGCGGGGCCCGTGACGAAGGCTTGAGCGAGGGCGTGCAAGATTCCGAATACCGCAAGCGACAGGCCG<br>ATCATCGTCGCGCTCCAGCGAAAGCGGTCCTCGCCGAAAATGACCCAGAGCGCTGCCGGAACCTGTCCTACGAGTTGCATGATAAAGAAAACAG<br>TCATAAGTGCGGCGACGATAGTCATGCCCCGCGCCCACCGGAAGGAGCTGACTGGATTGAAGGCACGCAAGGGCATCGGACGGCGCTCTCCCT<br>TATGCGATTCCTGCATAAGGAAGCAGCCCAGGAGGAGGTTGAGGCCGTTGAGCACCGCCGCCGCAAGGAATGGTGCATGCAAGGAGATGGCA<br>CCCAACAGTCCCCCGGCCACGGGGCCTGCCACCATACCCACGCCGAAACAAGCGCTCATGAGCCCGAAGTGGCGAGCCCGATCTTCCCCATCGG<br>TGATGTCGGCGATATAGGCACCAGCAACCGCACCTGTGGCACCCGTGATGCCCGCCACGATGCGTCCGGCGTAGAGAATCCACAGGACGGGTG<br>TGGTCGCCATGATCGCGTAGTCGATAGTGGCTCCAAGGAGCGAAGCGAGCAGGACTGGACGGCGGCCAAAGCGGTCGGACAGGGCTCCGAGA<br>ACGGGTGCGCAAAGAAATTGCATCAACGCATAGAGCGCAAGCAGCACGCCATAGTGACTGGCAATACTGTCGGAATGGACGATATCCCGCAAG<br>AGGCCCGGCAGTACCGGCATAACCAAGCCTATGCCTACAGCGTCCAGGGTGACGGTGCCGAGAATGACGATGAGCGCATTGTTAGATTTCATAC<br>ACGGTGCCTGACTGCGTTAGCAATTTAACTGTGATAAACTACCGCATTAAAGCTTATCGATGATAAGCTGTCAAACATGAGAATCTCCTCTAGTAA<br>ACACCCCTTGTATTACTGTTTATGTAAGCAGACAGTTTTATTGTTCATGATGATATATTTTTATCTTGTGCAATGTAACATCAGAGATTTTGAGACA<br>CAACGTGGCTTTGTTGAATAAATCGAACTTTTGCTGAGTTGAAGGATCAGATCACGCATCTTAACA |
| **pMC0_8_12_Atet (mScarlet-I)(Vn)** | TGCTGAGTCACTAAGGGCTAACTAACTAATTACGTAGCAATCAACTCACTGGCTCACCTTCACGGGTGGGCCTTTCTTCGGCACGGGCAAATTGC<br>TGAATATTCCTTTTCTTAGACGTCAGGTGGCAAACGCGAAGTAATCTTTTCGGTTTTAAAGAAAAAGGGCAGGGTGGTGACACCTTGCCCGTTTTT<br>TTGCCGGAGCTAGTATTATTAGGTCGAGGTGGCCCGGCTCCATGCACCGCGACGCAACGCGGGGAGGCAGACAAGGTAAAGGGCGGCACCTAC<br>AATCCATGCCAACCCGTTCCATGTGCTCGCCGAGGCGGCATAAATCGCCGTGACGATCAGCGGTCCAATGATCGAAGTGAGGCTGGTAAGAGCC<br>GCGAGCGACCCTTGAAGCTGTCCCTGATGGTCGTCATCTACTTGGCGGGACAGCATGGCCTGCAACGCGGGCATACCGATGCCGCCGGAAGCG<br>AGAAGAATCATAATGGGGAAGGCCATCCAGCCGCGCGTCGCGAACGCCAGCAAGACGTAGCCCAGCGCGTCGGCCGCCATGCCCGCGATAATG<br>GCCTGCTTCTCGCCGAAACGTTTGGTGGCGGGGCCCGTGACGAAGGCTTGAGCGAGGGCGTGCAAGATTCCGAATACCGCAAGCGACAGGCCG<br>ATCATCGTCGCGCTCCAGCGAAAGCGGTCCTCGCCGAAAATGACCCAGAGCGCTGCCGGAACCTGTCCTACGAGTTGCATGATAAAGAAAACAG<br>TCATAAGTGCGGCGACGATAGTCATGCCCCGCGCCCACCGGAAGGAGCTGACTGGATTGAAGGCACGCAAGGGCATCGGACGGCGCTCTCCCT<br>TATGCGATTCCTGCATAAGGAAGCAGCCCAGGAGGAGGTTGAGGCCGTTGAGCACCGCCGCCGCAAGGAATGGTGCATGCAAGGAGATGGCA<br>CCCAACAGTCCCCCGGCCACGGGGCCTGCCACCATACCCACGCCGAAACAAGCGCTCATGAGCCCGAAGTGGCGAGCCCGATCTTCCCCATCGG<br>TGATGTCGGCGATATAGGCACCAGCAACCGCACCTGTGGCACCCGTGATGCCCGCCACGATGCGTCCGGCGTAGAGAATCCACAGGACGGGTG<br>TGGTCGCCATGATCGCGTAGTCGATAGTGGCTCCAAGGAGCGAAGCGAGCAGGACTGGACGGCGGCCAAAGCGGTCGGACAGGGCTCCGAGA<br>ACGGGTGCGCAAAGAAATTGCATCAACGCATAGAGCGCAAGCAGCACGCCATAGTGACTGGCAATACTGTCGGAATGGACGATATCCCGCAAG<br>AGGCCCGGCAGTACCGGCATAACCAAGCCTATGCCTACAGCGTCCAGGGTGACGGTGCCGAGAATGACGATGAGCGCATTGTTAGATTTCATAC<br>ACGGTGCCTGACTGCGTTAGCAATTTAACTGTGATAAACTACCGCATTAAAGCTTATCGATGATAAGCTGTCAAACATGAGAATCTCCTCTAGTAA<br>ACACCCCTTGTATTACTGTTTATGTAAGCAGACAGTTTTATTGTTCATGATGATATATTTTTATCTTGTGCAATGTAACATCAGAGATTTTGAGACA<br>CAACGTGGCTTTGTTGAATAAATCGAACTTTTGCTGAGTTGAAGGATCAGATCACGCATCTTAACA |
| **pMC0_8_13_Acam (sfgfp)(Vn)** | TGCTACCAATAAAAAACGCCCGGCGGCAACCGAGCGTTCTGAACAAATCCAGATGGAGTTCTGAGGTCATTACTGGATCTATCAACAGGAGTCC<br>AAGCGAGCTCGATATCAAATTACGCCCCGCCCTGCCACTCATCGCAGTACTGTTGTAATTCATTAAGCATTCTGCCGACATGGAAGCCATCACAAA<br>CGGCATGATGAACCTGAATCGCCAGCGGCATCAGCACCTTGTCGCCTTGCGTATAATATTTGCCCATGGTGAAAACGGGGGCGAAGAAGTTGTC<br>CATATTGGCCACGTTTAAATCAAAACTGGTGAAACTCACCCAGGGATTGGCTGAAACGAAAAACATATTCTCAATAAACCCTTTAGGGAAATAGG<br>CCAGGTTTTCACCGTAACACGCCACATCTTGCGAATATATGTGTAGAAACTGCCGGAAATCGTCGTGGTATTCACTCCAGAGCGATGAAAACGTT<br>TCAGTTTGCTCATGGAAAACGGTGTAACAAGGGTGAACACTATCCCATATCACCAGCTCACCGTCTTTCATTGCCATACGAAATTCCGGATGAGC<br>ATTCATCAGGCGGGCAAGAATGTGAATAAAGGCCGGATAAAACTTGTGCTTATTTTTCTTTACGGTCTTTAAAAAGGCCGTAATATCCAGCTGAA<br>CGGTCTGGTTATAGGTACATTGAGCAACTGACTGAAATGCCTCAAAATGTTCTTTACGATGCCATTGGGATATATCAACGGTGGTATATCCAGTG<br>ATTTTTTTCTCCATTTTAGCTTCCTTAGCTCCTGAAAATCTCGATAACTCAAAAAATACGCCCGGTAGTGATCTTATTTCATTATGGTGAAAGTTGG<br>AACCTCTTACGTGCCCGATCAAAACA |
| **pMC0_8_14_Acam (mScarlet-I)(Vn)** | TGCTACCAATAAAAAACGCCCGGCGGCAACCGAGCGTTCTGAACAAATCCAGATGGAGTTCTGAGGTCATTACTGGATCTATCAACAGGAGTCC<br>AAGCGAGCTCGATATCAAATTACGCCCCGCCCTGCCACTCATCGCAGTACTGTTGTAATTCATTAAGCATTCTGCCGACATGGAAGCCATCACAAA<br>CGGCATGATGAACCTGAATCGCCAGCGGCATCAGCACCTTGTCGCCTTGCGTATAATATTTGCCCATGGTGAAAACGGGGGCGAAGAAGTTGTC<br>CATATTGGCCACGTTTAAATCAAAACTGGTGAAACTCACCCAGGGATTGGCTGAAACGAAAAACATATTCTCAATAAACCCTTTAGGGAAATAGG<br>CCAGGTTTTCACCGTAACACGCCACATCTTGCGAATATATGTGTAGAAACTGCCGGAAATCGTCGTGGTATTCACTCCAGAGCGATGAAAACGTT<br>TCAGTTTGCTCATGGAAAACGGTGTAACAAGGGTGAACACTATCCCATATCACCAGCTCACCGTCTTTCATTGCCATACGAAATTCCGGATGAGC<br>ATTCATCAGGCGGGCAAGAATGTGAATAAAGGCCGGATAAAACTTGTGCTTATTTTTCTTTACGGTCTTTAAAAAGGCCGTAATATCCAGCTGAA<br>CGGTCTGGTTATAGGTACATTGAGCAACTGACTGAAATGCCTCAAAATGTTCTTTACGATGCCATTGGGATATATCAACGGTGGTATATCCAGTG |

| | |
|---|---|
| | ATTTTTTTCTCCATTTTAGCTTCCTTAGCTCCTGAAAATCTCGATAACTCAAAAAATACGCCCGGTAGTGATCTTATTTCATTATGGTGAAAGTTGG<br>AACCTCTTACGTGCCCGATCAAAACA |
| **pMC0_8_18_Akan(Vn) (sfgfp)(Vn)** | TGCTAAACGGGCAAGGTGTCACCACCCTGCCCTTTTTCTTTAAAACCGAAAAGATTACTTCGCGTTATGCAGGCTTCCTCGCTCACTGACTCGCTG<br>CGCTCGGTCGTTCGGCTGCGGCGAGCGGTATCAGCTCACTCAAAGGCGGTAATCTCGAGTCCCGTCAAGTCAGCGTAATGCTCTGCCAGTGTTAC<br>AACCAATTAACCAATTCTGATTAAAAGAACTCGTCCAGCATCAGGTGAAATTGCAGTTTGTTCATGTCTGGGTTATCGATGCCGTATTTTTGAAAC<br>AGACGTTTTTGCAGAGATGGGCTAAATTCACCCAGACAGTTCCACAGAATTGCCAGATCTTGGTAACGATCTGCAATACCAACACGACCAACATC<br>AATACAACCGATCAGTTTACCCTCGTCAAAAATCAGGTTATCCAGAGAAAAATCACCGTGGGTAACAACGCTATCTGGAGAAAATGGCAGCAGTT<br>TGTGCATTTCTTTCCAAACTTGTTCAACTGGCCAACCGTTACGTTCATCATCAAAATCAGATGCATCAACCAGACCGTTGTTCATACGAGATTGTGC<br>TTGTGCCAGACGAAAAACACGATCGCTGTTAAATGGACAGTTACAAACTGGAATAGAGTGCAGACGACGCAGAAAAACTGCCAGTGCATCAACA<br>ATGTTTTCACCAGAATCTGGGTATTCTTCCAGAACTTGAAATGCGGTTTTACCTGGAATTGCGGTGGTCAGCAGCCATGCATCATCTGGGGTACG<br>AATAAAGTGTTTAATGGTTGGCAGTGGCATAAATTCGGTCAGCCAGTTTAAACGAACCATTTCATCGGTAACATCGTTTGCAACAGAACCTTTACC<br>GTGTTTCAGAAACAGTTCTGGTGCATCTGGTTTACCGTAAAGACGGTAAATGGTTGCACCAGATTGACCAACGTTATCACGTGCCCATTTGTAACC<br>GTACAGATCTGCATCCATGTTAGAGTTCAGACGTGGACGAGAACAAGAGGTTTCACGTTGAATGTGAGACATAACACCCCTTGTATTACTGTTTA<br>TGTAAGCAGACAGTTTTATTGTTCATGATGATATATTTTTATCTTGTGCAATGTAACATCAGAGATTTTGAGACACAACGTGGCTTTGTTGAATAA<br>ATCGAACTTTTGCTGAGTTGAAGGATCAGATCACGCATCTTAACA |
| **pMC0_8_19_Akan(Vn) (mScarlet)(Vn)** | TGCTAAACGGGCAAGGTGTCACCACCCTGCCCTTTTTCTTTAAAACCGAAAAGATTACTTCGCGTTATGCAGGCTTCCTCGCTCACTGACTCGCTG<br>CGCTCGGTCGTTCGGCTGCGGCGAGCGGTATCAGCTCACTCAAAGGCGGTAATCTCGAGTCCCGTCAAGTCAGCGTAATGCTCTGCCAGTGTTAC<br>AACCAATTAACCAATTCTGATTAAAAGAACTCGTCCAGCATCAGGTGAAATTGCAGTTTGTTCATGTCTGGGTTATCGATGCCGTATTTTTGAAAC<br>AGACGTTTTTGCAGAGATGGGCTAAATTCACCCAGACAGTTCCACAGAATTGCCAGATCTTGGTAACGATCTGCAATACCAACACGACCAACATC<br>AATACAACCGATCAGTTTACCCTCGTCAAAAATCAGGTTATCCAGAGAAAAATCACCGTGGGTAACAACGCTATCTGGAGAAAATGGCAGCAGTT<br>TGTGCATTTCTTTCCAAACTTGTTCAACTGGCCAACCGTTACGTTCATCATCAAAATCAGATGCATCAACCAGACCGTTGTTCATACGAGATTGTGC<br>TTGTGCCAGACGAAAAACACGATCGCTGTTAAATGGACAGTTACAAACTGGAATAGAGTGCAGACGACGCAGAAAAACTGCCAGTGCATCAACA<br>ATGTTTTCACCAGAATCTGGGTATTCTTCCAGAACTTGAAATGCGGTTTTACCTGGAATTGCGGTGGTCAGCAGCCATGCATCATCTGGGGTACG<br>AATAAAGTGTTTAATGGTTGGCAGTGGCATAAATTCGGTCAGCCAGTTTAAACGAACCATTTCATCGGTAACATCGTTTGCAACAGAACCTTTACC<br>GTGTTTCAGAAACAGTTCTGGTGCATCTGGTTTACCGTAAAGACGGTAAATGGTTGCACCAGATTGACCAACGTTATCACGTGCCCATTTGTAACC<br>GTACAGATCTGCATCCATGTTAGAGTTCAGACGTGGACGAGAACAAGAGGTTTCACGTTGAATGTGAGACATAACACCCCTTGTATTACTGTTTA<br>TGTAAGCAGACAGTTTTATTGTTCATGATGATATATTTTTATCTTGTGCAATGTAACATCAGAGATTTTGAGACACAACGTGGCTTTGTTGAATAA<br>ATCGAACTTTTGCTGAGTTGAAGGATCAGATCACGCATCTTAACA |
| **pMC0_8_21_Acarb J23106 (sfgfp)(Vn)** | TGCTTTTCTACGGGGTCTGACGCTCAGTGGAACGAAAACTCACGTTAAGGGATTTTGGTCATGAGATTATCAAAAAGGATCTTCACCTAGATCCTT<br>TTAAATTAAAAATGAAGTTTTAAATCAATCTAAAGTATATATGAGTAAACTTGGTCTGACAGTTACCAATGCTTAATCAGTGAGGCACCTATCTCA<br>GCGATCTGTCTATTTCGTTCATCCATAGTTGCCTGACTCCCCGTCGTGTAGATAACTACGATACGGGAGGGCTTACCATCTGGCCCCAGTGCTGCA<br>ATGATACCGCGTGACCCACGCTCACCGGCTCCAGATTTATCAGCAATAAACCAGCCAGCCGGAAGGGCCGAGCGCAGAAGTGGTCCTGCAACTT<br>TATCCGCCTCCATCCAGTCTATTAATTGTTGCCGGGAAGCTAGAGTAAGTAGTTCGCCAGTTAATAGTTTGCGCAACGTTGTTGCCATTGCTGTAG<br>GCATCGTGGTGTCACGCTCGTCGTTTGGTATGGCTTCATTCAGCTCCGGTTCCCAACGATCAAGGCGAGTTACATGATCCCCCATGTTGTGCAAAA<br>AAGCGGTTAGCTCCTTCGGTCCTCCGATCGTTGTCAGAAGTAAGTTGGCCGCAGTGTTATCACTCATGGTTATGGCAGCACTGCATAATTCTCTTA<br>CTGTCATGCCATCCGTAAGATGCTTTTCTGTGACTGGTGAGTACTCAACCAAGTCATTCTGAGAATAGTGTATGCGGCGACCGAGTTGCTCTTGCC<br>CGGCGTCAACACGGGATAATACCGCGCCACATAGCAGAACTTTAAAAGTGCTCATCATTGGAAAACGTTCTTCGGGGCGAAAACTCTCAAGGAT<br>CTTACCGCTGTTGAGATCCAGTTCGATGTAACCCACTCGTGCACCCAACTGATCTTCAGCATCTTTTACTTTCACCAGCGTTTCTGGGTGAGCAAAA<br>ACAGGAAGGCAAAATGCCGCAAAAAAGGGAATAAGGGCGACACGGAAATGTTGAATACTCATTGATTATTTCTCCTCTTTCTCTAGTAGCTAGCA<br>CTATACCTAGGACTGAGCTAGCCGTAAACTCCAACA |
| **pMC0_8_23_Acarb J23106 (mScarlet-I)(Vn)** | TGCTTTTCTACGGGGTCTGACGCTCAGTGGAACGAAAACTCACGTTAAGGGATTTTGGTCATGAGATTATCAAAAAGGATCTTCACCTAGATCCTT<br>TTAAATTAAAAATGAAGTTTTAAATCAATCTAAAGTATATATGAGTAAACTTGGTCTGACAGTTACCAATGCTTAATCAGTGAGGCACCTATCTCA<br>GCGATCTGTCTATTTCGTTCATCCATAGTTGCCTGACTCCCCGTCGTGTAGATAACTACGATACGGGAGGGCTTACCATCTGGCCCCAGTGCTGCA<br>ATGATACCGCGTGACCCACGCTCACCGGCTCCAGATTTATCAGCAATAAACCAGCCAGCCGGAAGGGCCGAGCGCAGAAGTGGTCCTGCAACTT<br>TATCCGCCTCCATCCAGTCTATTAATTGTTGCCGGGAAGCTAGAGTAAGTAGTTCGCCAGTTAATAGTTTGCGCAACGTTGTTGCCATTGCTGTAG<br>GCATCGTGGTGTCACGCTCGTCGTTTGGTATGGCTTCATTCAGCTCCGGTTCCCAACGATCAAGGCGAGTTACATGATCCCCCATGTTGTGCAAAA<br>AAGCGGTTAGCTCCTTCGGTCCTCCGATCGTTGTCAGAAGTAAGTTGGCCGCAGTGTTATCACTCATGGTTATGGCAGCACTGCATAATTCTCTTA<br>CTGTCATGCCATCCGTAAGATGCTTTTCTGTGACTGGTGAGTACTCAACCAAGTCATTCTGAGAATAGTGTATGCGGCGACCGAGTTGCTCTTGCC<br>CGGCGTCAACACGGGATAATACCGCGCCACATAGCAGAACTTTAAAAGTGCTCATCATTGGAAAACGTTCTTCGGGGCGAAAACTCTCAAGGAT<br>CTTACCGCTGTTGAGATCCAGTTCGATGTAACCCACTCGTGCACCCAACTGATCTTCAGCATCTTTTACTTTCACCAGCGTTTCTGGGTGAGCAAAA<br>ACAGGAAGGCAAAATGCCGCAAAAAAGGGAATAAGGGCGACACGGAAATGTTGAATACTCATTGATTATTTCTCCTCTTTCTCTAGTAGCTAGCA<br>CTATACCTAGGACTGAGCTAGCCGTAAACTCCAACA |
| **pMC0_1*_01_5C1C** | AACAGGTCTCGGGAGGGAG |
| **pMC0_1*_02_5C1R** | AACAGGTCTCGAACAGGAG |
| **pMC0_1*_03_5C2** | AACAGGTCTCGTACTGGAG |
| **pMC0_1*_04_5C3** | AACAGGTCTCGAATGGGAG |
| **pMC0_1*_05_5C4** | AACAGGTCTCGGCTTGGAG |
| **pMC0_1*_06_5C5** | AACAGGTCTCGGGTAGGAG |
| **pMC0_6*_01_3C1** | CGCTTACTGGAGACCAGCT |
| **pMC0_6*_02_3C2** | CGCTAATGGGAGACCAGCT |
| **pMC0_6*_03_3C3** | CGCTGCTTGGAGACCAGCT |
| **pMC0_6*_04_3C4** | CGCTGGTAGGAGACCAGCT |
| **pMC0_6*_05_3C5C** | CGCTCGCTGGAGACCAGCT |
| **pMC0_6*_06_3C5O** | CGCTAGCTGGAGACCAGCT |
| **pMC0_7*_01_OColE1** | AGCTGCGCTAGCGGAGTGTATACTGGCTTACTATGTTGGCACTGATGAGGGTGTCAGTGAAGTGCTTCATGTGGCAGGAGAAAAAAGGCTGCA<br>CCGGTGCGTCAGCAGAATATGTGATACAGGATATATTCCGCTTCCTCGCTCACTGACTCGCTACGCTCGGTCGTTCGACTGCGGCGAGCGGAAAT<br>GGCTTACGAACGGGGCGGAGATTTCCTGGAAGATGCCAGGAAGATACTTAACAGGGAAGTGAGAGGGCCGCGGCAAAGCCGTTTTTCCATAGG<br>CTCCGCCCCCCTGACAAGCATCACGAAATCTGACGCTCAAATCAGTGGTGGCGAAACCCGACAGGACTATAAAGATACCAGGCGTTTCCCCTGGC<br>GGCTCCCTCGTGCGCTCTCCTGTTCCTGCCTTTCGGTTTACCGGTGTCATTCCGCTGTTATGGCCGCGTTTGTCTCATTCCACGCCTGACACTCAGT<br>TCCGGGTAGGCAGTTCGCTCCAAGCTGGACTGTATGCACGAACCCCCCGTTCAGTCCGACCGCTGCGCCTTATCCGGTAACTATCGTCTTGAGTC<br>CAACCCGGAAAGACATGCAAAAGCACCACTGGCAGCAGCCACTGGTAATTGATTTAGAGGAGTTAGTCTTGAAGTCATGCGCCGGTTAAGGCTA<br>AACTGAAAGGACAAGTTTTGGTGACTGCGCTCCTCCAAGCCAGTTACCTCGGTTCAAAGAGTTGGTAGCTCAGAGAACCTTCGAAAAACCGCCCT<br>GCAAGGCGGTTTTTTCGTTTTCAGAGCAAGAGATTACGCGCAGACCAAAACGATCTCAAGAAGATCATCTTATTAAGGGGTCTGACGCTCAGTG<br>GAACGAAAACTCACGTTAAGGGATTTTGGTCATGAGATTATCAAAAAGGATCTTCACCTAGATCCTTTTAAATTTGCT |
| **pMC0_7*_02_Op15A** | AGCTCCGCGTTGCTGGCGTTTTTCCACAGGCTCCGCCCCCCTGACGAGCATCACAAAAATCGACGCTCAAGTCAGAGGTGGCGAAACCCGACAG<br>GACTATAAAGATACCAGGCGTTTCCCCCTGGAAGCTCCCTCGTGCGCTCTCCTGTTCCGACCCTGCCGCTTACCGGATACCTGTCCGCCTTTCTCCC<br>TTCGGGAAGCGTGGCGCTTTCTCATAGCTCACGCTGTAGGTATCTCAGTTCGGTGTAGGTCGTTCGCTCCAAGCTGGGCTGTGTGCACGAACCCC<br>CCGTTCAGCCCGACCGCTGCGCCTTATCCGGTAACTATCGTCTTGAGTCCAACCCGGTAAGACACGACTTATCGCCACTGGCAGCAGCCACTGGT<br>AACAGGATTAGCAGAGCGAGGTATGTAGGCGGTGCTACAGAGTTCTTGAAGTGGTGGCCTAACTACGGCTACACTAGAAGAACAGTATTTGGT<br>ATCTGCGCTCTGCTGAAGCCAGTTACCTTCGGAAAAAGAGTTGGTAGCTCTTGATCCGGCAAACAAACCACCGCTGGTAGCGGTGGTTTTTTTGT<br>TTGCAAGCAGCAGATTACGCGCAGAAAAAAAGGATCTCAAGAAGATCCTTTGATCTTTTCTGCT |

| | |
|---|---|
| **pMC0_7*_04_OpMB1-M** | AGCTACGGTTATCCACAGAATCAGGGGATAACGCAGGAAAGAACATGTGAGCAAAAGGCCAGCAAAAGGCCAGGAACCGTAAAAAGGCCGCG<br>TTGCTGGCGTTTTTCCATAGGCTCCGCCCCCCTGACGAGCATCACAAAAATCGACGCTCAAGTCAGAGGTGGCGAAACCCGACAGGACTATAAA<br>GATACCAGGCGTTTCCCCCTGGAAGCTCCCTCGTGCGCTCTCCTGTTCCGACCCTGCCGCTTACCGGATACCTGTCCGCCTTTCTCCCTTCGGGAA<br>GCGTGGCGCTTTCTCATAGCTCACGCTGTAGGTATCTCAGTTCGGTGTAGGTCGTTCGCTCCAAGCTGGGCTGTGTGCACGAACCCCCCGTTCAG<br>CCCGACCGCTGCGCCTTATCCGGTAACTATCGTCTTGAGTCCAACCCGGTAAGACACGACTTATCGCCACTGGCAGCAGCCACTGGTAACAGGAT<br>TAGCAGAGCGAGGTATGTAGGCGGTGCTACAGAGTTCTTGAAGTGGTGGCCTAACTACGGCTACACTAGAAGAACAGTATTTGGTATCTGCGCT<br>CTGCTGAAGCCAGTTACCTTCGGAAAAAGAGTTGGTAGCTCTTGATCCGGCAAACAAACCACCGCTGGTAGCGGTGGTTTTTTTGTTTGCAAGCA<br>GCAGATTACGCGCAGAAAAAAAGGATCTCAAGAAGATCCTTTGATCTTTTCTACGGGGTCTGACGCTCAGTGGAACGAAAACTCACGTTAAGGG<br>ATTTTGGTCATGATGCT |
| **pMC0_7*_05_ORSF1010** | AGCTTTCTGAAAGCGACCAGGTGCTCGGCGTGGCAAGACTCGCAGCGAACCCGTAGAAAGCCATGCTCCAGCCGCCCGCATTGGAGAAATTCTT<br>CAAATTCCCGTTGCACATAGCCCGGCAATTCCTTTCCCTGCTCTGCCATAAGCGCAGCGAATGCCGGGTAATACTCGTCAACGATCTGATAGAGA<br>AGGGTTTGCTCGGGTCGGTGGCTCTGGTAACGACCAGTATCCCGATCCCGGCTGGCCGTCCTGGCCGCCACATGAGGCATGTTCCGCGTCCTTGC<br>AATACTGTGTTTACATACAGTCTATCGCTTAGCGGAAAGTTCTTTTACCCTCAGCCGAAATGCCTGCCGTTGCTAGACATTGCCAGCCAGTGCCCG<br>TCACTCCCGTACTAACTGTCACGAACCCCTGCAATAACTGTCACGCCCCCCTGCAATAACTGTCACGAACCCCTGCAATAACTGTCACGCCCCCAA<br>ACCTGCAAACCCAGCAGGGGCGGGGGCTGGCGGGGTGTTGGAAAAATCCATCCATGATTATCTAAGAATAATCCACTAGGCGCGGTTATCAGC<br>GCCCTTGTGGGGCGCTGCTGCCCTTGCCCAATATGCCCGGCCAGAGGCCGGATAGCTGGTCTATTCGCTGCGCTAGGCTACACACCGCCCCACCG<br>CTGCGCGGCAGGGGGAAAGGCGGGCAAAGCCCGCTAAACCCCACACCAAACCCCGCAGAAATACGCTGGAGCGCTTTTAGCCGCTTTAGCGGC<br>CTTTCCCCCTACCCGAAGGGTGGGGGCGCGTGTGCAGCCCCGCAGGGCCTGTCTCGGTCGATCATTCAGCCCGGCTCATCCTTCTGGCGTGGCG<br>GCAGACCGAACAAGGCGCGGTCGTGGTCGCGTTCAAGGTACGCATCCATTGCCGCCATGAGCCGATCCTCCGGCCACTCGCTGCTGTTCACCTTG<br>GCCAAAATCATGGCCCCCACCAGCACCTTGCGCCTTGTTTCGTTCTTGCGCTCTTGCTGCTGTTCCCTTGCCCGCTCCCGCTGAATTTCGGCATTGA<br>TTCGCGCTCGTTGTTCTTCGAGCTTGGCCAGCCGATCCGCCGCCTTGTTGCTCCCCTTAACCATCTTGACACCCCATTGTTAATGTGCTGTCTCGTA<br>GGCTATCATGGAGGCACAGCGGCGGCAATCCCGACCCTACTTTGTAGGGGAGGGCGCACTTACCGGTTTCTCTTCGAGAAACTGGCCTAACGGC<br>CACCCTTCGGGCGGTGCGCTCTCCGAGGGCCATTGCATGGAGCCGAAAAGCAAAAGCAACAGCGAGGCAGCATGGCGATTTATCACCTTACGG<br>CGAAAACCGGCAGCAGGTCGGGCGGCCAATCGGCCAGGGCCAAGGCCGACTACATCCAGCGCGAAGGCAAGTATGCCCGCGACATGGATGAA<br>GTCTTGCACGCCGAATCCGGGCACATGCCGGAGTTCGTCGAGCGGCCCGCCGACTACTGGGATGCTGCCGACCTGTATGAACGCGCCAATGGGC<br>GGCTGTTCAAGGAGGTCGAATTTGCCCTGCCGGTCGAGCTGACCCTCGACCAGCAGAAGGCGCTGGCGTCCGAGTTCGCCCAGCACCTGACCGG<br>TGCCGAGCGCCTGCCGTATACGCTGGCCATCCATGCCGGTGGCGGCGAGAACCCGCACTGCCACCTGATGATCTCCGAGCGGATCAATGACGGC<br>ATCGAGCGGCCCGCCGCTCAGTGGTTCAAGCGGTACAACGGCAAGACCCCGGAGAAGGGCGGGGCACAGAAGACCGAAGCGCTCAAGCCCAA<br>GGCATGGCTTGAGCAGACCCGCGAGGCATGGGCCGACCATGCCAACCGGGCATTAGAGCGGGCTGGCCACGACGCCCGCATTGACCACAGAAC<br>ACTTGAGGCGCAGGGCATCGAGCGCCTGCCCGGTGTTCACCTGGGGCCGAACGTGGTGGAGATGGAAGGCCGGGGCATCCGCACCGACCGGG<br>CAGACGTGGCCCTGAACATCGACACCGCCAACGCCCAGATCATCGACTTACAGGAATACCGGGAGGCAATAGACCATGAACGCAATCGACAGA<br>GTGAAGAAATCCAGAGGCATCAACGAGTTAGCGGAGCAGATCGAACCGCTGGCCCAGAGCATGGCGACACTGGCCGACGAAGCCCGGCAGGT<br>CATGAGCCAGACCAAGCAGGCCAGCGAGGCGCAGGCGGCGGAGTGGCTGAAAGCCCAGCGCCAGACAGGGGCGGCATGGGTGGAGCTGGCC<br>AAAGAGTTGCGGGAGGTAGCCGCCGAGGTGAGCAGCGCCGCGCAGAGCGCCCGGAGCGCGTCGCGGGGGTGGCACTGGAAGCTATGGCTAA<br>CCGTGATGCTGGCTTCCATGATGCCTACGGTGGTGCTGCTGATCGCATCGTTGCTCTTGCTCGACCTGACGCCACTGACAACCGAGGACGGCTCG<br>ATCTGGCTGCGCTTGGTGGCCCGATGAAGAACGACAGGACTTTGCAGGCCATAGGCCGACAGCTCAAGGCCATGGGCTGTGAGCGCTTCGATAT<br>CGGCGTCAGGGACGCACCCACCGGCCAGATGATGAACCGGGAATGGTCAGCCGCCGAAGTGCTCCAGAACACGCCATGGCTCAAGCGGATGAA<br>TGCCCAGGGCAATGACGTGTATATCAGGCCCGCCGAGCAGGAGCGGCATGGTCTGGTGCTGGTGGACGACCTCAGCGAGTTTGACCTGGATGA<br>CATGAAAGCCGAGGGCCGGGAGCCTGCCCTGGTAGTGGAAACCAGCCCGAAGAACTATCAGGCATGGGTCAAGGTGGCCGACGCCGCAGGCG<br>GTGAACTTCGGGGGCAGATTGCCCGGACGCTGGCCAGCGAGTACGACGCCGACCCGGCCAGCGCCGACAGCCGCCACTATGGCCGCTTGGCGG<br>GCTTCACCAACCGCAAGGACAAGCACACCACCCGCGCCGGTTATCAGCCGTGGGTGCTGCTGCGTGAATCCAAGGGCAAGACCGCCACCGCTGG<br>CCCGGCGCTGGTGCAGCAGGCTGGCCAGCAGATCGAGCAGGCCCAGCGGCAGCAGGAGAAGGCCCGCAGGCTGGCCAGCCTCGAACTGCCCG<br>AGCGGCAGCTTAGCCGCCACCGGCGCACGGCGCTGGACGAGTACCGCAGCGAGATGGCCGGGCTGGTCAAGCGCTTCGGTCATGACCTCAGCA<br>AGTGCGACTTTATCGCCGCGCAGAAGCTGGCCAGCCGGGGCCGCAGTGCCGAGGAAATCGGCAAGGCCATGGCCGAGGCCAGCCCAGCGCTG<br>GCAGAGCGCAAGCCCGGCCACGAAGCGGATTACATCGAGCGCACCGTCAGCAAGGTCATGGGTCTGCCCAGCGTCCAGCTTGCGCGGGCCGAG<br>CTGGCACGGGCACCGGCACCCCGCCAGCGAGGCATGGACAGGGGCGGGCCAGATTTCAGCATGTAGTGCTTGCGTTGGTACTCACGCCTGTTAT<br>ACTATGAGTACTCACGCACAGAAGGGGGTTTTATGGAATACGAAAAAAGCGCTTCAGGGTCGGTCTACCTGATCAAAAGTGACAAGGGCTATTG<br>GTTGCCCGGTGGCTTTGGTTATACGTCAAACAAGGCCGAGGCTGGCCGCTTTTCAGTCGCTGATATGGCCAGCCTTAACCTTGACGGCTGCACCT<br>TGTCCTTGTTCCGCGAAGACAAGCCTTTCGGCCCCGGCAAGTTTCTCGGTGACTGATATGAAAGACCAAAAGGACAAGCAGACCGGCGACCTGC<br>TGGCCAGCCCTGACGCTGTACGCCAAGCGCGATATGCCGAGCGCATGAAGGCCAAAGGGATGCGTCAGCGCAAGTTCTGGCTGACCGACGACG<br>AATACGAGGCGCTGCGCGAGTGCCTGGAAGAACTCAGAGCGGCGCAGGGCGGGGGTAGTGACCCCGCCAGCGCCTAACCACCAACTGCCTGC<br>AAAGGAGGCAATCAATGGCTACCCATAAGCCTATCAATATTCTGGAGGCGTTCGCAGCAGCGCCGCCACCGCTGGACTACGTTTTGCCCAACATG<br>GTGGCCGGTACGGTCGGGGCGCTGGTGTCGCCCGGTGGTGCCGGTAAATCCATGCTGGCCCTGCAACTGGCCGCACAGATTGCAGGCGGGCCG<br>GATCTGCTGGAGGTGGGCGAACTGCCCACCGGCCCGGTGATCTACCTGCCCGCCGAAGACCCGCCCACCGCCATTCATCACCGCCTGCACGCCCT<br>TGGGGCGCACCTCAGCGCCGAGGAACGGCAAGCCGTGGCTGACGGCCTGCTGATCCAGCCGCTGATCGGCAGCCTGCCCAACATCATGGCCCC<br>GGAGTGGTTCGACGGCCTCAAGCGCGCCGCCGAGGGCCGCCGCCTGATGGTGCTGGACACGCTGCGCCGGTTCCACATCGAGGAAGAAAACGC<br>CAGCGGCCCCATGGCCCAGGTCATCGGTCGCATGGAGGCCATCGCCGCCGATACCGGGTGCTCTATCGTGTTCCTGCACCATGCCAGCAAGGGC<br>GCGGCCATGATGGGCGCAGGCGACCAGCAGCAGGCCAGCCGGGGCAGCTCGGTACTGGTCGATAACATCCGCTGGCAGTCCTACCTGTCGAGC<br>ATGACCAGCGCCGAGGCCGAGGAATGGGGTGTGGACGACGACCAGCGCCGGTTCTTCGTCCGCTTCGGTGTGAGCAAGGCCAACTATGGCGCA<br>CCGTTCGCTGATCGGTGGTTCAGGCGGCATGACGGCGGGGTGCTCAAGCCCGCCGTGCTGGAGAGGCAGCGCAAGAGCAAGGGGGTGCCCCG<br>TGGTGAAGCCTAAGAACAAGCACAGCCTCAGCCACGTCCGGCACGACCCGGCGCACTGTCTGGCCCCCGGCCTGTTCCGTGCCCTCAAGCGGGG<br>CGAGCGCAAGCGCAGCAAGCTGGACGTGACGTATGACTACGGCGACGGCAAGCGGATCGAGTTCAGCGGCCCGGAGCCGCTGGGCGCTGATG<br>ATCTGCGCATCCTGCAAGGGCTGGTGGCCATGGCTGGGCCTAATGGCCTAGTGCTTGGCCCGGAACCCAAGACCGAAGGCGGACGGCAGCTCC<br>GGCTGTTCCTGGAACCCAAGTGGGAGGCCGTCACCGCTGAATGCCATGTGGTCAAAGGTAGCTATCGGGCGCTGGCAAAGGAAATCGGGGCAG<br>AGGTCGATAGTGGTGGGGCGCTCAAGCACATACAGGACTGCATCGAGCGCCTTTGGAAGGTATCCATCATCGCCCAGAATGGCCGCAAGCGGC<br>AGGGGTTTCGGCTGCTGTCGGAGTACGCCAGCGACGAGGCGGACGGGCGCCTGTACGTGGCCCTGAACCCCTTGATCGCGCAGGCCGTCATGG<br>GTGGCGGCCAGCATGTGCGCATCAGCATGGACGAGGTGCGGGCGCTGGACAGCGAAACCGCCCGCCTGCTGCACCAGCGGCTGTGTGGCTGG<br>ATCGACCCCGGCAAAACCGGCAAGGCTTCCATAGATACCTTGTGCGGCTATGTCTGGCCGTCAGAGGCCAGTGGTTCGACCATGCGCAAGCGCC<br>GCAAGCGGGTGCGCGAGGCGTTGCCGGAGCTGGTCGCGCTGGGCTGGACGGTAACCGAGTTCGCGGCGGGCAAGTACGACATCACCCGGCCC<br>AAGGCGGCAGGCTGACCCCCCCCACTCTATTGTAAACAAGACATTTTTATCTTTTATATTCAATGGCTTATTTTCCTGCTAATTGGTAATACCATGA<br>AAAATACCATGCTCAGAAAAGGCTTAACAATATTTTGAAAAATTGCCTACTGAGCGCTGCCGCACAGCTCCATAGGCCGCTTTCCTGGCTTTGCTT<br>CCAGATGTATGCTCTTCTGCTCCCGAACGCCAGCAAGACGTAGCCCAGCGCGTCGGCCAGCTTGCAATTCGCGCTAACTTACATTAATTGCGTTGC<br>GCTGCT |
| **pMC0_8*_03_ATet (sfGFP)** | TGCTGAGTCACTAAGGGCTAACTAACTAATTACGTAGCAATCAACTCACTGGCTCACCTTCACGGGTGGGCCTTTCTTCGGCACGGGCAAATTGC<br>TGAATATTCCTTTTCTTAGACGTCAGGTGGCAAACGCGAAGTAATCTTTTCGGTTTTAAAGAAAAAGGGCAGGGTGGTGACACCTTGCCCGTTTTT<br>TTGCCGGAGCTAGTATTATTAGGTCGAGGTGGCCCGGCTCCATGCACCGCGACGCAACGCGGGGAGGCAGACAAGGTAAAGGGCGGCACCTAC<br>AATCCATGCCAACCCGTTCCATGTGCTCGCCGAGGCGGCATAAATCGCCGTGACGATCAGCGGTCCAATGATCGAAGTGAGGCTGGTAAGAGCC<br>GCGAGCGACCCTTGAAGCTGTCCCTGATGGTCGTCATCTACTTGGCGGGACAGCATGGCCTGCAACGCGGGCATACCGATGCCGCCGGAAGCG<br>AGAAGAATCATAATGGGGAAGGCCATCCAGCCGCGCGTCGCGAACGCCAGCAAGACGTAGCCCAGCGCGTCGGCCGCCATGCCCGCGATAATG<br>GCCTGCTTCTCGCCGAAACGTTTGGTGGCGGGGCCCGTGACGAAGGCTTGAGCGAGGGCGTGCAAGATTCCGAATACCGCAAGCGACAGGCCG<br>ATCATCGTCGCGCTCCAGCGAAAGCGGTCCTCGCCGAAAATGACCCAGAGCGCTGCCGGAACCTGTCCTACGAGTTGCATGATAAAGAAAACAG<br>TCATAAGTGCGGCGACGATAGTCATGCCCCGCGCCCACCGGAAGGAGCTGACTGGATTGAAGGCACGCAAGGGCATCGGACGGCGCTCTCCCT<br>TATGCGATTCCTGCATAAGGAAGCAGCCCAGGAGGAGGTTGAGGCCGTTGAGCACCGCCGCCGCAAGGAATGGTGCATGCAAGGAGATGGCA |

| | |
|---|---|
| | CCCAACAGTCCCCCGGCCACGGGGCCTGCCACCATACCCACGCCGAAACAAGCGCTCATGAGCCCGAAGTGGCGAGCCCGATCTTCCCCATCGGTGATGTCGGCGATATAGGCACCAGCAACCGCACCTGTGGCACCCGTGATGCCCGCCACGATGCGTCCGGCGTAGAGAATCCACAGGACGGGTGTGGTCGCCATGATCGCGTAGTCGATAGTGGCTCCAAGGAGCGAAGCGAGCAGGACTGGACGGCGGCCAAAGCGGTCGGACAGGGCTCCGAGAACGGGTGCGCAAAGAAATTGCATCAACGCATAGAGCGCAAGCAGCACGCCATAGTGACTGGCAATACTGTCGGAATGGACGATATCCCGCAAGAGGCCCGGCAGTACCGGCATAACCAAGCCTATGCCTACAGCGTCCAGGGTGACGGTGCCGAGAATGACGATGAGCGCATTGTTAGATTTCATACACGGTGCCTGACTGCGTTAGCAATTTAACTGTGATAAACTACCGCATTAAAGCTTATCGATGATAAGCTGTCAAACATGAGAATCTCCTCTAGTAAACACCCCTTGTATTACTGTTTATGTAAGCAGACAGTTTTATTGTTCATGATGATATATTTTTATCTTGTGCAATGTAACATCAGAGATTTTGAGACACAACGTGGCTTTGTTGAATAAATCGAACTTTTGCTGAGTTGAAGGATCAGATCACGCATCTTAACA |
| **pMC0_8*_04_ATet (mScarlet-I)** | TGCTACCAATAAAAAACGCCCGGCGGCAACCGAGCGTTCTGAACAAATCCAGATGGAGTTCTGAGGTCATTACTGGATCTATCAACAGGAGTCCAAGCGAGCTCGATATCAAATTACGCCCCGCCCTGCCACTCATCGCAGTACTGTTGTAATTCATTAAGCATTCTGCCGACATGGAAGCCATCACAAACGGCATGATGAACCTGAATCGCCAGCGGCATCAGCACCTTGTCGCCTTGCGTATAATATTTGCCCATGGTGAAAACGGGGGCGAAGAAGTTGTCCATATTGGCCACGTTTAAATCAAAACTGGTGAAACTCACCCAGGGATTGGCTGAAACGAAAAACATATTCTCAATAAACCCTTTAGGGAAATAGGCCAGGTTTTCACCGTAACACGCCACATCTTGCGAATATATGTGTAGAAACTGCCGGAAATCGTCGTGGTATTCACTCCAGAGCGATGAAAACGTTTCAGTTTGCTCATGGAAAACGGTGTAACAAGGGTGAACACTATCCCATATCACCAGCTCACCGTCTTTCATTGCCATACGAAATTCCGGATGAGCATTCATCAGGCGGGCAAGAATGTGAATAAAGGCCGGATAAAACTTGTGCTTATTTTTCTTTACGGTCTTTAAAAAGGCCGTAATATCCAGCTGAACGGTCTGGTTATAGGTACATTGAGCAACTGACTGAAATGCCTCAAAATGTTCTTTACGATGCCATTGGGATATATCAACGGTGGTATATCCAGTGATTTTTTTCTCCATTTTAGCTTCCTTAGCTCCTGAAAATCTCGATAACTCAAAAAATACGCCCGGTAGTGATCTTATTTCATTATGGTGAAAGTTGGAACCTCTTACGTGCCCGATCAAAACA |
| **pMC0_8*_05_ACam (sfGFP)** | TGCTACCAATAAAAAACGCCCGGCGGCAACCGAGCGTTCTGAACAAATCCAGATGGAGTTCTGAGGTCATTACTGGATCTATCAACAGGAGTCCAAGCGAGCTCGATATCAAATTACGCCCCGCCCTGCCACTCATCGCAGTACTGTTGTAATTCATTAAGCATTCTGCCGACATGGAAGCCATCACAAACGGCATGATGAACCTGAATCGCCAGCGGCATCAGCACCTTGTCGCCTTGCGTATAATATTTGCCCATGGTGAAAACGGGGGCGAAGAAGTTGTCCATATTGGCCACGTTTAAATCAAAACTGGTGAAACTCACCCAGGGATTGGCTGAAACGAAAAACATATTCTCAATAAACCCTTTAGGGAAATAGGCCAGGTTTTCACCGTAACACGCCACATCTTGCGAATATATGTGTAGAAACTGCCGGAAATCGTCGTGGTATTCACTCCAGAGCGATGAAAACGTTTCAGTTTGCTCATGGAAAACGGTGTAACAAGGGTGAACACTATCCCATATCACCAGCTCACCGTCTTTCATTGCCATACGAAATTCCGGATGAGCATTCATCAGGCGGGCAAGAATGTGAATAAAGGCCGGATAAAACTTGTGCTTATTTTTCTTTACGGTCTTTAAAAAGGCCGTAATATCCAGCTGAACGGTCTGGTTATAGGTACATTGAGCAACTGACTGAAATGCCTCAAAATGTTCTTTACGATGCCATTGGGATATATCAACGGTGGTATATCCAGTGATTTTTTTCTCCATTTTAGCTTCCTTAGCTCCTGAAAATCTCGATAACTCAAAAAATACGCCCGGTAGTGATCTTATTTCATTATGGTGAAAGTTGGAACCTCTTACGTGCCCGATCAAAACA |
| **pMC0_8*_06_ACam (mScarlet-I)** | TGCTAAACGGGCAAGGTGTCACCACCCTGCCCTTTTTCTTTAAAACCGAAAAGATTACTTCGCGTTATGCAGGCTTCCTCGCTCACTGACTCGCTGCGCTCGGTCGTTCGGCTGCGGCGAGCGGTATCAGCTCACTCAAAGGCGGTAATCTCGAGTCCCGTCAAGTCAGCGTAATGCTCTGCCAGTGTTACAACCAATTAACCAATTCTGATTAAAAGAACTCGTCCAGCATCAGGTGAAATTGCAGTTTGTTCATGTCTGGGTTATCGATGCCGTATTTTTGAAACAGACGTTTTTGCAGAGATGGGCTAAATTCACCCAGACAGTTCCACAGAATTGCCAGATCTTGGTAACGATCTGCAATACCAACACGACCAACATCAATACAACCGATCAGTTTACCCTCGTCAAAAATCAGGTTATCCAGAGAAAAATCACCGTGGGTAACAACGCTATCTGGAGAAAATGGCAGCAGTTTGTGCATTTCTTTCCAAACTTGTTCAACTGGCCAACCGTTACGTTCATCATCAAAATCAGATGCATCAACCAGACCGTTGTTCATACGAGATTGTGCTTGTGCCAGACGAAAAACACGATCGCTGTTAAATGGACAGTTACAAACTGGAATAGAGTGCAGACGACGCAGAAAAACTGCCAGTGCATCAACAATGTTTTCACCAGAATCTGGGTATTCTTCCAGAACTTGAAATGCGGTTTTACCTGGAATTGCGGTGGTCAGCAGCCATGCATCATCTGGGGTACGAATAAAGTGTTTAATGGTTGGCAGTGGCATAAATTCGGTCAGCCAGTTTAAACGAACCATTTCATCGGTAACATCGTTTGCAACAGAACCTTTACCGTGTTTCAGAAACAGTTCTGGTGCATCTGGTTTACCGTAAAGACGGTAAATGGTTGCACCAGATTGACCAACGTTATCACGTGCCCATTTGTAACCGTACAGATCTGCATCCATGTTAGAGTTCAGACGTGGACGAGAACAAGAGGTTTCACGTTGAATGTGAGACATAACACCCCTTGTATTACTGTTTATGTAAGCAGACAGTTTTATTGTTCATGATGATATATTTTTATCTTGTGCAATGTAACATCAGAGATTTTGAGACACAACGTGGCTTTGTTGAATAAATCGAACTTTTGCTGAGTTGAAGGATCAGATCACGCATCTTAACA |
| **pMC0_8*_09_AKan (Vn)(sfGFP)** | TGCTAAACGGGCAAGGTGTCACCACCCTGCCCTTTTTCTTTAAAACCGAAAAGATTACTTCGCGTTATGCAGGCTTCCTCGCTCACTGACTCGCTGCGCTCGGTCGTTCGGCTGCGGCGAGCGGTATCAGCTCACTCAAAGGCGGTAATCTCGAGTCCCGTCAAGTCAGCGTAATGCTCTGCCAGTGTTACAACCAATTAACCAATTCTGATTAAAAGAACTCGTCCAGCATCAGGTGAAATTGCAGTTTGTTCATGTCTGGGTTATCGATGCCGTATTTTTGAAACAGACGTTTTTGCAGAGATGGGCTAAATTCACCCAGACAGTTCCACAGAATTGCCAGATCTTGGTAACGATCTGCAATACCAACACGACCAACATCAATACAACCGATCAGTTTACCCTCGTCAAAAATCAGGTTATCCAGAGAAAAATCACCGTGGGTAACAACGCTATCTGGAGAAAATGGCAGCAGTTTGTGCATTTCTTTCCAAACTTGTTCAACTGGCCAACCGTTACGTTCATCATCAAAATCAGATGCATCAACCAGACCGTTGTTCATACGAGATTGTGCTTGTGCCAGACGAAAAACACGATCGCTGTTAAATGGACAGTTACAAACTGGAATAGAGTGCAGACGACGCAGAAAAACTGCCAGTGCATCAACAATGTTTTCACCAGAATCTGGGTATTCTTCCAGAACTTGAAATGCGGTTTTACCTGGAATTGCGGTGGTCAGCAGCCATGCATCATCTGGGGTACGAATAAAGTGTTTAATGGTTGGCAGTGGCATAAATTCGGTCAGCCAGTTTAAACGAACCATTTCATCGGTAACATCGTTTGCAACAGAACCTTTACCGTGTTTCAGAAACAGTTCTGGTGCATCTGGTTTACCGTAAAGACGGTAAATGGTTGCACCAGATTGACCAACGTTATCACGTGCCCATTTGTAACCGTACAGATCTGCATCCATGTTAGAGTTCAGACGTGGACGAGAACAAGAGGTTTCACGTTGAATGTGAGACATAACACCCCTTGTATTACTGTTTATGTAAGCAGACAGTTTTATTGTTCATGATGATATATTTTTATCTTGTGCAATGTAACATCAGAGATTTTGAGACACAACGTGGCTTTGTTGAATAAATCGAACTTTTGCTGAGTTGAAGGATCAGATCACGCATCTTAACA |
| **pMC0_8*_10_AKan (Vn)(mScarlet-I)** | TGCTTTTCTACGGGGTCTGACGCTCAGTGGAACGAAAACTCACGTTAAGGGATTTTGGTCATGAGATTATCAAAAAGGATCTTCACCTAGATCCTTTTAAATTAAAAATGAAGTTTTAAATCAATCTAAAGTATATATGAGTAAACTTGGTCTGACAGTTACCAATGCTTAATCAGTGAGGCACCTATCTCAGCGATCTGTCTATTTCGTTCATCCATAGTTGCCTGACTCCCCGTCGTGTAGATAACTACGATACGGGAGGGCTTACCATCTGGCCCCAGTGCTGCAATGATACCGCGTGACCCACGCTCACCGGCTCCAGATTTATCAGCAATAAACCAGCCAGCCGGAAGGGCCGAGCGCAGAAGTGGTCCTGCAACTTTATCCGCCTCCATCCAGTCTATTAATTGTTGCCGGGAAGCTAGAGTAAGTAGTTCGCCAGTTAATAGTTTGCGCAACGTTGTTGCCATTGCTGTAGGCATCGTGGTGTCACGCTCGTCGTTTGGTATGGCTTCATTCAGCTCCGGTTCCCAACGATCAAGGCGAGTTACATGATCCCCCATGTTGTGCAAAAAAGCGGTTAGCTCCTTCGGTCCTCCGATCGTTGTCAGAAGTAAGTTGGCCGCAGTGTTATCACTCATGGTTATGGCAGCACTGCATAATTCTCTTACTGTCATGCCATCCGTAAGATGCTTTTCTGTGACTGGTGAGTACTCAACCAAGTCATTCTGAGAATAGTGTATGCGGCGACCGAGTTGCTCTTGCCCGGCGTCAACACGGGATAATACCGCGCCACATAGCAGAACTTTAAAAGTGCTCATCATTGGAAAACGTTCTTCGGGGCGAAAACTCTCAAGGATCTTACCGCTGTTGAGATCCAGTTCGATGTAACCCACTCGTGCACCCAACTGATCTTCAGCATCTTTTACTTTCACCAGCGTTTCTGGGTGAGCAAAAACAGGAAGGCAAAATGCCGCAAAAAAGGGAATAAGGGCGACACGGAAATGTTGAATACTCATTGATTATTTCTCCTCTTTCTCTAGTAGCTAGCACTATACCTAGGACTGAGCTAGCCGTAAACTCCAACA |
| **pMC0_8*_12_ACarb J23106 (sfGFP)** | TGCTTTTCTACGGGGTCTGACGCTCAGTGGAACGAAAACTCACGTTAAGGGATTTTGGTCATGAGATTATCAAAAAGGATCTTCACCTAGATCCTTTTAAATTAAAAATGAAGTTTTAAATCAATCTAAAGTATATATGAGTAAACTTGGTCTGACAGTTACCAATGCTTAATCAGTGAGGCACCTATCTCAGCGATCTGTCTATTTCGTTCATCCATAGTTGCCTGACTCCCCGTCGTGTAGATAACTACGATACGGGAGGGCTTACCATCTGGCCCCAGTGCTGCAATGATACCGCGTGACCCACGCTCACCGGCTCCAGATTTATCAGCAATAAACCAGCCAGCCGGAAGGGCCGAGCGCAGAAGTGGTCCTGCAACTTTATCCGCCTCCATCCAGTCTATTAATTGTTGCCGGGAAGCTAGAGTAAGTAGTTCGCCAGTTAATAGTTTGCGCAACGTTGTTGCCATTGCTGTAGGCATCGTGGTGTCACGCTCGTCGTTTGGTATGGCTTCATTCAGCTCCGGTTCCCAACGATCAAGGCGAGTTACATGATCCCCCATGTTGTGCAAAAAAGCGGTTAGCTCCTTCGGTCCTCCGATCGTTGTCAGAAGTAAGTTGGCCGCAGTGTTATCACTCATGGTTATGGCAGCACTGCATAATTCTCTTACTGTCATGCCATCCGTAAGATGCTTTTCTGTGACTGGTGAGTACTCAACCAAGTCATTCTGAGAATAGTGTATGCGGCGACCGAGTTGCTCTTGCCCGGCGTCAACACGGGATAATACCGCGCCACATAGCAGAACTTTAAAAGTGCTCATCATTGGAAAACGTTCTTCGGGGCGAAAACTCTCAAGGATCTTACCGCTGTTGAGATCCAGTTCGATGTAACCCACTCGTGCACCCAACTGATCTTCAGCATCTTTTACTTTCACCAGCGTTTCTGGGTGAGCAAAAACAGGAAGGCAAAATGCCGCAAAAAAGGGAATAAGGGCGACACGGAAATGTTGAATACTCATTGATTATTTCTCCTCTTTCTCTAGTAGCTAGCACTATACCTAGGACTGAGCTAGCCGTAAACTCCAACA |
| **pMC0_8*_14_ACarb J23106 (mScarlet-I)** | TGCTTTTCTACGGGGTCTGACGCTCAGTGGAACGAAAACTCACGTTAAGGGATTTTGGTCATGAGATTATCAAAAAGGATCTTCACCTAGATCCTTTTAAATTAAAAATGAAGTTTTAAATCAATCTAAAGTATATATGAGTAAACTTGGTCTGACAGTTACCAATGCTTAATCAGTGAGGCACCTATCTCAGCGATCTGTCTATTTCGTTCATCCATAGTTGCCTGACTCCCCGTCGTGTAGATAACTACGATACGGGAGGGCTTACCATCTGGCCCCAGTGCTGCAATGATACCGCGTGACCCACGCTCACCGGCTCCAGATTTATCAGCAATAAACCAGCCAGCCGGAAGGGCCGAGCGCAGAAGTGGTCCTGCAACTTTATCCGCCTCCATCCAGTCTATTAATTGTTGCCGGGAAGCTAGAGTAAGTAGTTCGCCAGTTAATAGTTTGCGCAACGTTGTTGCCATTGCTGTAG |

| | |
|---|---|
| | GCATCGTGGTGTCACGCTCGTCGTTTGGTATGGCTTCATTCAGCTCCGGTTCCCAACGATCAAGGCGAGTTACATGATCCCCCATGTTGTGCAAAA<br>AAGCGGTTAGCTCCTTCGGTCCTCCGATCGTTGTCAGAAGTAAGTTGGCCGCAGTGTTATCACTCATGGTTATGGCAGCACTGCATAATTCTCTTA<br>CTGTCATGCCATCCGTAAGATGCTTTTCTGTGACTGGTGAGTACTCAACCAAGTCATTCTGAGAATAGTGTATGCGGCGACCGAGTTGCTCTTGCC<br>CGGCGTCAACACGGGATAATACCGCGCCACATAGCAGAACTTTAAAAGTGCTCATCATTGGAAAACGTTCTTCGGGGCGAAAACTCTCAAGGAT<br>CTTACCGCTGTTGAGATCCAGTTCGATGTAACCCACTCGTGCACCCAACTGATCTTCAGCATCTTTTACTTTCACCAGCGTTTCTGGGTGAGCAAAA<br>ACAGGAAGGCAAAATGCCGCAAAAAAGGGAATAAGGGCGACACGGAAATGTTGAATACTCATTGATTATTTCTCCTCTTTCTCTAGTAGCTAGCA<br>CTATACCTAGGACTGAGCTAGCCGTAAACTCCAACA |
| **pMC0_TU2-6*_EL_01** | TACTGCTGCGATTAGACAGCCAGCAGCT |
| **pMC0_TU3-6*_EL_02** | AATGGCTGCGATTAGACAGCCAGCAGCT |
| **pMC0_TU4-6*_EL_03** | GCTTGCTGCGATTAGACAGCCAGCAGCT |
| **pMC0_TU5-6*_EL_04** | GGTAGCTGCGATTAGACAGCCAGCAGCT |
| **pMC0_1*-6*_01_Dropout sfGFP** | GGAGAGAGACGGAAAGTGAAACGTGATTTCATGCGTCATTTTGAACATTTTGTAAATCTTATTTAATAATGTGTGCGGCAATTCACATTTAATTTA<br>TGAATGTTTTCTTAACATCGCGGCAACTCAAGAAACGGCAGGTTCGGATCTTAGCTACTAGAGAAAGAGGAGAAATACTAGATGCGTAAAGGCG<br>AAGAACTTTTCACTGGCGTAGTACCAATTTTAGTGGAGCTGGACGGAGATGTAAATGGTCATAAGTTTTCAGTTCGAGGAGAAGGCGAAGGAGA<br>TGCAACTAACGGTAAGCTGACACTAAAGTTTATCTGTACCACGGGTAAACTGCCGGTCCCATGGCCGACACTGGTTACTACACTTACTTATGGTGT<br>ACAATGCTTTGCTCGTTATCCTGACCATATGAAGCAGCACGATTTCTTCAAATCGGCCATGCCTGAAGGATACGTGCAAGAACGTACAATTAGCTT<br>TAAAGATGACGGCACCTATAAAACGCGCGCCGAAGTTAAATTCGAGGGTGACACATTGGTTAATCGTATAGAACTTAAGGGCATTGACTTCAAA<br>GAAGATGGCAACATCCTTGGCCATAAACTGGAATATAATTTTAACTCTCACAATGTCTACATTACGGCGGATAAACAAAAGAATGGCATTAAAGC<br>GAATTTCAAAATCCGCCACAATGTCGAAGACGGCTCGGTTCAACTGGCGGACCATTATCAGCAGAACACCCCAATCGGTGATGGCCCGGTTCTGT<br>TACCTGATAATCATTACCTTTCTACTCAAAGCGTTTTATCTAAAGATCCTAACGAGAAGCGTGATCATATGGTTCTACTGGAATTTGTTACCGCAGC<br>TGGTATCACGCACGGCATGGATGAGCTGTATAAATGACCAGGCATCAAATAAAACGAAAGGCTCAGTCGAAAGACTGGGCCTTTCGTTTTATCT<br>GTTGTTTGTCGGTGAACGCTCTCTACTAGAGTCACACTGGCTCACCTTCGGGTGGGCCTTTCTGCGTTTATACGTCTCTCGCT |
| **pMC0_1*-6*_02_Dropout mScarlet-I** | GGAGAGAGACGGAAAGTGAAACGTGATTTCATGCGTCATTTTGAACATTTTGTAAATCTTATTTAATAATGTGTGCGGCAATTCACATTTAATTTA<br>TGAATGTTTTCTTAACATCGCGGCAACTCAAGAAACGGCAGGTTCGGATCTTAGCTACTAGAGAAAGAGGAGAAATACTAGATGGTTTCTAAAG<br>GTGAAGCAGTGATCAAAGAATTTATGCGCTTCAAAGTTCACATGGAAGGTTCTATGAACGGCCACGAATTTGAAATTGAAGGTGAAGGCGAAG<br>GTCGTCCATACGAAGGTACTCAAACCGCAAAACTGAAAGTTACCAAAGGTGGTCCACTGCCATTTTCTTGGGATATTCTGTCTCCACAATTTATGT<br>ACGGTTCTCGTGCATTTATCAAACACCCAGCAGATATTCCAGACTACTACAAACAATCTTTTCCGGAAGGTTTCAAATGGGAACGTGTTATGAATT<br>TTGAAGATGGTGGTGCAGTTACGGTTACCCAAGATACCTCTCTGGAAGATGGTACTCTGATCTACAAAGTTAAACTGCGTGGTACTAACTTTCCA<br>CCAGATGGTCCAGTTATGCAGAAAAAAACCATGGGTTGGGAAGCATCTACCGAACGTCTGTACCCAGAAGATGGCGTTCTGAAAGGTGATATCA<br>AAATGGCACTGCGTCTGAAAGATGGCGGTCGTTACCTGGCAGATTTCAAAACCACCTACAAAGCGAAAAAACCAGTTCAAATGCCAGGTGCATA<br>CAACGTTGATCGTAAACTGGATATTACCAGCCACAACGAAGATTACACCGTTGTTGAACAATACGAACGTTCTGAAGGCCGTCACTCTACCGGTG<br>GTATGGATGAACTGTACAAATAACCAGGCATCAAATAAAACGAAAGGCTCAGTCGAAAGACTGGGCCTTTCGTTTTATCTGTTGTTTGTCGGTGA<br>ACGCTCTCTACTAGAGTCACACTGGCTCACCTTCGGGTGGGCCTTTCTGCGTTTATACGTCTCTCGCT |
| **pMC0_TU1-TU5_01_Dropout sfGFP** | AACAAGAGACGGAAAGTGAAACGTGATTTCATGCGTCATTTTGAACATTTTGTAAATCTTATTTAATAATGTGTGCGGCAATTCACATTTAATTTA<br>TGAATGTTTTCTTAACATCGCGGCAACTCAAGAAACGGCAGGTTCGGATCTTAGCTACTAGAGAAAGAGGAGAAATACTAGATGGTTTCTAAAG<br>GTGAAGCAGTGATCAAAGAATTTATGCGCTTCAAAGTTCACATGGAAGGTTCTATGAACGGCCACGAATTTGAAATTGAAGGTGAAGGCGAAG<br>GTCGTCCATACGAAGGTACTCAAACCGCAAAACTGAAAGTTACCAAAGGTGGTCCACTGCCATTTTCTTGGGATATTCTGTCTCCACAATTTATGT<br>ACGGTTCTCGTGCATTTATCAAACACCCAGCAGATATTCCAGACTACTACAAACAATCTTTTCCGGAAGGTTTCAAATGGGAACGTGTTATGAATT<br>TTGAAGATGGTGGTGCAGTTACGGTTACCCAAGATACCTCTCTGGAAGATGGTACTCTGATCTACAAAGTTAAACTGCGTGGTACTAACTTTCCA<br>CCAGATGGTCCAGTTATGCAGAAAAAAACCATGGGTTGGGAAGCATCTACCGAACGTCTGTACCCAGAAGATGGCGTTCTGAAAGGTGATATCA<br>AAATGGCACTGCGTCTGAAAGATGGCGGTCGTTACCTGGCAGATTTCAAAACCACCTACAAAGCGAAAAAACCAGTTCAAATGCCAGGTGCATA<br>CAACGTTGATCGTAAACTGGATATTACCAGCCACAACGAAGATTACACCGTTGTTGAACAATACGAACGTTCTGAAGGCCGTCACTCTACCGGTG<br>GTATGGATGAACTGTACAAATAACCAGGCATCAAATAAAACGAAAGGCTCAGTCGAAAGACTGGGCCTTTCGTTTTATCTGTTGTTTGTCGGTGA<br>ACGCTCTCTACTAGAGTCACACTGGCTCACCTTCGGGTGGGCCTTTCTGCGTTTATACGTCTCTAGCT |
| **pMC0_TU1-TU5_02_Dropout mScarlet-I** | AGCTGAACCGTAAAAAAGGCCGCGTTGCTGGCGTTTTTCCACAGGCTCCGCCCCCCTGACGAGCATCACAAAAATCGACGCTCAAGTCAGAGGT<br>GGCGAAACCCGACAGGACTATAAAGATACCAGGCGTTTCCCCCTGGAAGCTCCCTCGTGCGCTCTCCTGTTCCGACCCTGCCGCTTACCGGATAC<br>CTGTCCGCCTTTCTCCCTTCGGGAAGCGTGGCGCTTTCTCATAGCTCACGCTGTAGGTATCTCAGTTCGGTGTAGGTCGTTCGCTCCAAGCTGGGC<br>TGTGTGCACGAACCCCCCGTTCAGCCCGACCGCTGCGCCTTATCCGGTAACTATCGTCTTGAGTCCAACCCGGTAAGACACGACTTATCGCCACTG<br>GCAGCAGCCACTGGTAACAGGATTAGCAGAGCGAGGTATGTAGGCGGTGCTACAGAGTTCTTGAAGTGGTGGCCTAACTACGGCTACACTAGA<br>AGAACAGTATTTGGTATCTGCGCTCTGCTGAAGCCAGTTACCTTCGGAAAAAGAGTTGGTAGCTCTTGATCCGGCAAACAAACCACCGCTGGTAG<br>CGGTGGTTTTTTTGTTTGCAAGCAGCAGATTACGCGCAGAAAAAAAGGATCTCAAGAAGATCCTTTGATCTTTTCTACGGGGTCTGACGCTCAGT<br>GGAACGAAAACTCACGTTAAGGGATTTTGGTCATGAGATTATCAAAAAGGATCTTCACTGCT |
| **pMC0_1*-6* Linker** | AACATATCCGTGAGAGCT |

**Table S4. Cloning of LVL0 and LVL0* parts.** Abbreviations used in methods: GGA = Golden Gate Assembly; Gib = Gibson Assembly; Digest = digestion of PCR product with BsaI. Parts marked with a hash tag are experimental parts and were not characterized or described in this publication but will be made available through Addgene.

| Part | fwd Oligo | fwd Oligo Sequence | rev Oligo | rev Oligo Sequence | Method | Template |
|---|---|---|---|---|---|---|
| **pMC0_1-6_01_1-6 Dropout** | oiGEM1131_Dropout_1_fwd | AACGTCTCGCTCGAACAAGAGACCGAAAGTGAAACGTGATTTCATGCG | oiGEM1142_Dropout_6_rev | TTCGTCTCCCTCAAGCTAGAGACCTATAAACGCAGAAAGGCCCACC | PCR, GGA | pMC_V_06_Part Entry sfGFP (Vn) Ori Muta |
| **pMC0_1-6_02_1-6 Connector** | aiGEM1055_1-6_Connector_fwd | CTCGAACATATCCGTGAGAGCT | aiGEM1056_1-6_Connector_rev | CTCAAGCTCTCACGGATATGTT | Annealing, GGA | / |
| | oiGEM1128_GA_Entry_Vec_fwd | CCAGGCATCAAATAAAACGAAAGG | oiGEM1127_GA_Entry_Vec_rev | CTAGTATTTCTCCTCTTTCTCTAGTAG | PCR, Gib | pMC0_1-6_01_1-6 Dropout |

| | | | | | | |
|---|---|---|---|---|---|---|
| **pMC0_1-6_03_1-6 Dropout mScarlet-I (Vn)** | oiGEM1194_GA_Scarlet_fwd | ACTAGAGAAAGAGGAGAAATACTAGATGGTTTCTAAAGGTGAAGCAG | oiGEM1195_GA_Scarlet_rev | GCCTTTCGTTTTATTTGATGCCTGGTTATTTGTACAGTTCATCCATACC | | pMC0_4_12_CDSmScarlet-I (Vn) |
| **pMC0_1_01_1 Dropout** | oiGEM1131_Dropout_1_fwd | AACGTCTCGCTCGAACAAGAGACCGAAAGTGAAACGTGATTTCATGCG | oiGEM1132_Dropout_1_rev | TTCGTCTCCCTCACTCCAGAGACCTATAAACGCAGAAAGGCCCACC | PCR, GGA | pMC_V_06_Part Entry sfGFP (Vn) Ori Muta |
| **pMC0_1_02_5C1CLF** | / | / | / | / | DNA Synthesis, GGA | / |
| **pMC0_1_03_5C1RLF** | oiGEM1097_Aqua_5Con1_fwd | GAACACGTCTCGAACAGGACCAAAACGAAAAAAGACGC | oiGEM1098_Aqua_5Con1_rev | AGCGCGAGACCACTCATCGCTCTAG | PCR, Gib | pMC0_1_02_5C1CLN |
| **pMC0_1_04_5C1CSF** | aiGEM1181_5C1CSN_fwd | CTCGAACACGTCTCGGGAGGGAG | aiGEM1182_5C1CSN_rev | CTCACTCCCTCCCGAGACGTGTT | Annealing, GGA | / |
| **pMC0_1_05_5C1RSF** | aiGEM1073_5'Con_1_Short_Res_fwd | CTCGAACACGTCTCGAACAGGAG | aiGEM1074_5'Con_1_Short_Res_rev | CTCACTCCTGTTCGAGACGTGTT | Annealing, GGA | / |
| **pMC0_1_06_5C1SR** | aiGEM1107_5'Con1_inv_fwd | CTCGAACACGTCTCGAGTAGGAG | aiGEM1108_5'Con1_inv_rev | CTCACTCCTACTCGAGACGTGTT | Annealing, GGA | / |
| **pMC0_1_07_5C2LF#** | / | / | / | / | DNA Synthesis, GGA | / |
| **pMC0_1_08_5C2SF** | aiGEM1075_5'Con_2_Short_fwd | CTCGAACACGTCTCGTACTGGAG | aiGEM1076_5'Con_2_Short_rev | CTCACTCCAGTACGAGACGTGTT | Annealing, GGA | / |
| **pMC0_1_09_5C2SR** | aiGEM1109_5'Con2_inv_fwd | CTCGAACACGTCTCGCATTGGAG | aiGEM1110_5'Con2_inv_rev | CTCACTCCAATGCGAGACGTGTT | Annealing, GGA | / |
| **pMC0_1_10_5C3LF#** | / | / | / | / | DNA Synthesis, GGA | / |
| **pMC0_1_11_5C3SF** | aiGEM1077_5'Con_3_Short_fwd | CTCGAACACGTCTCGAATGGGAG | aiGEM1078_5'Con_3_Short_rev | CTCACTCCCATTCGAGACGTGTT | Annealing, GGA | / |
| **pMC0_1_12_5C3SR** | aiGEM1111_5'Con3_inv_fwd | CTCGAACACGTCTCGAAGCGGAG | aiGEM1112_5'Con3_inv_rev | CTCACTCCGCTTCGAGACGTGTT | Annealing, GGA | / |
| **pMC0_1_13_5C4LF#** | / | / | / | / | DNA Synthesis, GGA | / |
| **pMC0_1_14_5C4SF** | aiGEM1079_5'Con_4_Short_fwd | CTCGAACACGTCTCGGCTTGGAG | aiGEM1080_5'Con_4_Short_rev | CTCACTCCAAGCCGAGACGTGTT | Annealing, GGA | / |
| **pMC0_1_15_5C4SR** | aiGEM1113_5'Con4_inv_fwd | CTCGAACACGTCTCGTACCGGAG | aiGEM1114_5'Con4_inv_rev | CTCACTCCGGTACGAGACGTGTT | Annealing, GGA | / |
| **pMC0_1_16_5C5LF#** | / | / | / | / | DNA Synthesis, GGA | / |
| **pMC0_1_17_5C5SF** | aiGEM1081_5'Con_5_Short_fwd | CTCGAACACGTCTCGGGTAGGAG | aiGEM1082_5'Con_5_Short_rev | CTCACTCCTACCCGAGACGTGTT | Annealing, GGA | / |
| **pMC0_1_18_5C5CSR** | aiGEM1115_5'Con5_inv_fwd | CTCGAACACGTCTCGAGCGGGAG | aiGEM1116_5'Con5_inv_rev | CTCACTCCCGCTCGAGACGTGTT | Annealing, GGA | / |
| **pMC0_1_19_5C5OSR** | / | / | / | / | DNA Synthesis, GGA | / |
| **pMC0_1_20_5C0SF** | aiGEM1017_5'Conn._Dummy_fwd | CTCGAACAGAATTCGCGGCCGCTTCTAGAGGGAG | aiGEM1018_5'Conn._Dummy_rev | CTCACTCCCTCTAGAAGCGGCCGCGAATTCTGTT | Annealing, GGA | / |
| **pMC0_1_21_1 Dropout mScarlet-I (Vn)** | oiGEM1128_GA_Entry_Vec_fwd | CCAGGCATCAAATAAAACGAAAGG | oiGEM1127_GA_Entry_Vec_rev | CTAGTATTTCTCCTCTTTCTCTAGTAG | PCR, Gib | pMC0_1_01_1 Dropout |
| | oiGEM1194_GA_Scarlet_fwd | ACTAGAGAAAGAGGAGAAATACTAGATGGTTTCTAAAGGTGAAGCAG | oiGEM1195_GA_Scarlet_rev | GCCTTTCGTTTTATTTGATGCCTGGTTATTTGTACAGTTCATCCATACC | | pMC0_4_12_CDSmScarlet-I (Vn) |
| **pMC0_2-5_01_2-5 Dropout** | oiGEM1133_Dropout_2_fwd | AACGTCTCGCTCGGGAGAGAGACCGAAAGTGAAACGTGATTTCATGCG | oiGEM1140_Dropout_5_rev | TTCGTCTCCCTCAAGCGAGAGACCTATAAACGCAGAAAGGCCCACC | PCR, GGA | pMC_V_06_Part Entry sfGFP (Vn) Ori Muta |
| **pMC0_2-5_02_2-5 Dropout mScarlet-I (Vn)** | oiGEM1128_GA_Entry_Vec_fwd | CCAGGCATCAAATAAAACGAAAGG | oiGEM1127_GA_Entry_Vec_rev | CTAGTATTTCTCCTCTTTCTCTAGTAG | PCR, Gib | pMC0_2-5_01_2-5 Dropout |
| | oiGEM1194_GA_Scarlet_fwd | ACTAGAGAAAGAGGAGAAATACTAGATGGTTTCTAAAGGTGAAGCAG | oiGEM1195_GA_Scarlet_rev | GCCTTTCGTTTTATTTGATGCCTGGTTATTTGTACAGTTCATCCATACC | | pMC0_4_12_CDSmScarlet-I (Vn) |
| **pMC0_2_01_2 Dropout** | oiGEM1133_Dropout_2_fwd | AACGTCTCGCTCGGGAGAGAGACCGAAAGTGAAACGTGATTTCATGCG | oiGEM1134_Dropout_2_rev | TTCGTCTCCCTCAAGTAAGAGACCTATAAACGCAGAAAGGCCCACC | PCR, GGA | pMC_V_06_Part Entry sfGFP (Vn) Ori Muta |
| **pMC0_2_02_PJ23100** | aiGEM1009_J23100_fwd | CTCGGGAGTTGACGGCTAGCTCAGTCCTAGGTACAGTGCTAGCTACT | aiGEM1010_J23100_rev | CTCAAGTAGCTAGCACTGTACCTAGGACTGAGCTAGCCGTCAACTCC | Annealing, GGA | / |
| **pMC0_2_03_PJ23101** | aiGEM1021_J23101_fwd | CTCGGGAGTTTACAGCTAGCTCAGTCCTAGGTATTATGCTAGCTACT | aiGEM1022_J23101_rev | CTCAAGTAGCTAGCATAATACCTAGGACTGAGCTAGCTGTAAACTCC | Annealing, GGA | / |
| **pMC0_2_04_PJ23102** | aiGEM1023_J23102_fwd | CTCGGGAGTTGACAGCTAGCTCAGTCCTAGGTACTGTGCTAGCTACT | aiGEM1024_J23102_rev | CTCAAGTAGCTAGCACAGTACCTAGGACTGAGCTAGCTGTCAACTCC | Annealing, GGA | / |

| | | | | | | |
|---|---|---|---|---|---|---|
| **pMC0_2_05_PJ23103** | aiGEM1025_J23103_fwd | CTCGGGAGCTGATAGCTAGCTCAGTCCTAGGGATTATGCTAGCTACT | aiGEM1026_J23103_rev | CTCAAGTAGCTAGCATAATCCCTAGGACTGAGCTAGCTATCAGCTCC | Annealing, GGA | / |
| **pMC0_2_06_PJ23104** | aiGEM1011_J23104_fwd | CTCGGGAGTTGACAGCTAGCTCAGTCCTAGGTATTGTGCTAGCTACT | aiGEM1012_J23104_rev | CTCAAGTAGCTAGCACAATACCTAGGACTGAGCTAGCTGTCAACTCC | Annealing, GGA | / |
| **pMC0_2_07_PJ23105** | aiGEM1027_J23105_fwd | CTCGGGAGTTTACGGCTAGCTCAGTCCTAGGTACTATGCTAGCTACT | aiGEM1028_J23105_rev | CTCAAGTAGCTAGCATAGTACCTAGGACTGAGCTAGCCGTAAACTCC | Annealing, GGA | / |
| **pMC0_2_08_PJ23106** | aiGEM1013_J23106_fwd | CTCGGGAGTTTACGGCTAGCTCAGTCCTAGGTATAGTGCTAGCTACT | aiGEM1014_J23106_rev | CTCAAGTAGCTAGCACTATACCTAGGACTGAGCTAGCCGTAAACTCC | Annealing, GGA | / |
| **pMC0_2_09_PJ23107** | aiGEM1029_J23107_fwd | CTCGGGAGTTTACGGCTAGCTCAGCCCTAGGTATTATGCTAGCTACT | aiGEM1030_J23107_rev | CTCAAGTAGCTAGCATAATACCTAGGGCTGAGCTAGCCGTAAACTCC | Annealing, GGA | / |
| **pMC0_2_10_PJ23108** | aiGEM1031_J23108_fwd | CTCGGGAGCTGACAGCTAGCTCAGTCCTAGGTATAATGCTAGCTACT | aiGEM1032_J23108_rev | CTCAAGTAGCTAGCATTATACCTAGGACTGAGCTAGCTGTCAGCTCC | Annealing, GGA | / |
| **pMC0_2_11_PJ23109** | aiGEM1033_J23109_fwd | CTCGGGAGTTTACAGCTAGCTCAGTCCTAGGGACTGTGCTAGCTACT | aiGEM1034_J23109_rev | CTCAAGTACTAGCACAGTCCCTAGGACTGAGCTAGCTGTAAACTCC | Annealing, GGA | / |
| **pMC0_2_12_PJ23110** | aiGEM1035_J23110_fwd | CTCGGGAGTTTACGGCTAGCTCAGTCCTAGGTACAATGCTAGCTACT | aiGEM1036_J23110_rev | CTCAAGTAGCTAGCATTGTACCTAGGACTGAGCTAGCCGTAAACTCC | Annealing, GGA | / |
| **pMC0_2_13_PJ23111** | aiGEM1037_J23111_fwd | CTCGGGAGTTGACGGCTAGCTCAGTCCTAGGTATAGTGCTAGCTACT | aiGEM1038_J23111_rev | CTCAAGTAGCTAGCACTATACCTAGGACTGAGCTAGCCGTCAACTCC | Annealing, GGA | / |
| **pMC0_2_14_PJ23113** | aiGEM1039_J23113_fwd | CTCGGGAGCTGATGGCTAGCTCAGTCCTAGGGATTATGCTAGCTACT | aiGEM1040_J23113_rev | CTCAAGTAGCTAGCATAATCCCTAGGACTGAGCTAGCCATCAGCTCC | Annealing, GGA | / |
| **pMC0_2_15_PJ23114** | aiGEM1041_J23114_fwd | CTCGGGAGTTTATGGCTAGCTCAGTCCTAGGTACAATGCTAGCTACT | aiGEM1042_J23114_rev | CTCAAGTAGCTAGCATTGTACCTAGGACTGAGCTAGCCATAAACTCC | Annealing, GGA | / |
| **pMC0_2_16_PJ23115** | aiGEM1015_J23115_fwd | CTCGGGAGTTTATAGCTAGCTCAGCCCTTGGTACAATGCTAGCTACT | aiGEM1016_J23115_rev | CTCAAGTAGCTAGCATTGTACCAAGGGCTGAGCTAGCTATAAACTCC | Annealing, GGA | / |
| **pMC0_2_17_PJ23116** | aiGEM1043_J23116_fwd | CTCGGGAGTTGACAGCTAGCTCAGTCCTAGGGACTATGCTAGCTACT | aiGEM1044_J23116_rev | CTCAAGTAGCTAGCATAGTCCCTAGGACTGAGCTAGCTGTCAACTCC | Annealing, GGA | / |
| **pMC0_2_18_PJ23117** | aiGEM1045_J23117_fwd | CTCGGGAGTTGACAGCTAGCTCAGTCCTAGGGATTGTGCTAGCTACT | aiGEM1046_J23117_rev | CTCAAGTAGCTAGCACAATCCCTAGGACTGAGCTAGCTGTCAACTCC | Annealing, GGA | / |
| **pMC0_2_19_PJ23118** | aiGEM1047_J23118_fwd | CTCGGGAGTTGACGGCTAGCTCAGTCCTAGGTATTGTGCTAGCTACT | aiGEM1048_J23118_rev | CTCAAGTAGCTAGCACAATACCTAGGACTGAGCTAGCCGTCAACTCC | Annealing, GGA | / |
| **pMC0_2_20_PJ23119** | aiGEM1049_J23119_fwd | CTCGGGAGTTGACAGCTAGCTCAGTCCTAGGTATAATGCTAGCTACT | aiGEM1050_J23119_rev | CTCAAGTAGCTAGCATTATACCTAGGACTGAGCTAGCTGTCAACTCC | Annealing, GGA | / |
| **pMC0_2_21_PDummy** | aiGEM1139_PromDummy_fwd | CTCGGGAGCCCCTGGCGCCCCTTTACT | aiGEM1140_PromDummy_rev | CTCAAGTAAAGGGGCGCCAGGGGCTCC | Annealing, GGA | / |
| **pMC0_2_22_PTrc** | oiGEM1112_pLac_fwd | AACGTCTCGCTCGGGAGGTCTAGGGCGGCGGATTTG | oiGEM1113_pLac_rev_Muta | AACGTCTCGTGTCACTGGTGAAAAGAAAAACCAC | PCR, GGA | pSV0-1_004 (provided by Stefano Vecchione) |
| | oiGEM1114_pLac_fwd_Muta | AACGTCTCGGACACGGGCAACAGCTG | oiGEM1115_pLac_rev | TTCGTCTCCCTCAAGTAGGTCAGTGCGTCCTGCTG | | pSV0-1_004 (provided by Stefano Vecchione) |
| **pMC0_2_23_PTet** | oiGEM1110_pTet_fwd | AACGTCTCGCTCGGGAGTTTTGTTATCAATAAAAAAGGCCCCC | TTCGTCTCCCTCAAGTATTCACTTTTCTCTATCACTGATAGG | TTCGTCTCCCTCAAGTATTCACTTTTCTCTATCACTGATAGG | PCR, GGA | pSV0-1_005 (provided by Stefano Vecchione) |
| **pMC0_2_29_PBDA (Vn)** | oiGEM1153_pBDA(Vn)_fwd | AACGTCTCGCTCGGGAGTGTTATCCATCCACTGGTAGAGG | oiGEM1154_pBDA(Vn)_rev | TTCGTCTCCCTCAAGTATTTTGGTCAGCAAACTAAATCTACTTG | PCR, GGA | V. natriegens Genome |

| | | | | | | |
|---|---|---|---|---|---|---|
| **pMC0_2_33_PRham (Vn)** | oiGEM1159_pRhamnose(Vn)_fwd | AACGTCTCGCTCGGGAGGACACACTCTAATAACCAAGCC | oiGEM1160_pRhamnose(Vn)_rev | TTCGTCTCCCTCAAGTAGATTAACACCTCTTGTTCTTTAGAG | PCR, GGA | V. natriegens Genome |
| **pMC0_2_37_2 Dropout mScarlet-I (Vn)** | oiGEM1128_GA_Entry_Vec_fwd | CCAGGCATCAAATAAAACGAAAGG | oiGEM1127_GA_Entry_Vec_rev | CTAGTATTTCTCCTCTTTCTCTAGTAG | PCR, Gib | pMC0_2_01_2 Dropout |
| | oiGEM1194_GA_Scarlet_fwd | ACTAGAGAAAGAGGAGAAATACTAGATGGTTTCTAAAGGTGAAGCAG | oiGEM1195_GA_Scarlet_rev | GCCTTTCGTTTTATTTGATGCCTGGTTATTTGTACAGTTCATCCATACC | | pMC0_4_12_CDSmScarlet-I (Vn) |
| **pMC0_3_01_3 Dropout** | oiGEM1135_Dropout_3_fwd | AACGTCTCGCTCGTACTAGAGACCGAAAGTGAAACGTGATTTCATGCG | oiGEM1136_Dropout_3_rev | TTCGTCTCCCTCACATTAGAGACCTATAAACGCAGAAAGGCCCACC | PCR, GGA | pMC_V_06_Part Entry sfGFP (Vn) Ori Muta |
| **pMC0_3_02_RB0029** | aiGEM1141_B0029_fwd | CTCGTACTAGAGTTCACACAGGAAACCTAATCAATG | aiGEM1142_B0029_rev | CTCACATTGATTAGGTTTCCTGTGTGAACTCTAGTA | Annealing, GGA | / |
| **pMC0_3_03_RB0030** | aiGEM1001_B0030_fwd | CTCGTACTAGAGATTAAAGAGGAGAAATAATCAATG | aiGEM1002_B0030_rev | CTCACATTGATTATTTCTCCTCTTTAATCTCTAGTA | Annealing, GGA | / |
| **pMC0_3_04_RB0031** | aiGEM1003_B0031_fwd | CTCGTACTAGAGTCACACAGGAAACCTAATCAATG | aiGEM1004_B0031_rev | CTCACATTGATTAGGTTTCCTGTGTGACTCTAGTA | Annealing, GGA | / |
| **pMC0_3_05_RB0032** | aiGEM1005_B0032_fwd | CTCGTACTAGAGTCACACAGGAAAGTAATCAATG | aiGEM1006_B0032_rev | CTCACATTGATTACTTTCCTGTGTGACTCTAGTA | Annealing, GGA | / |
| **pMC0_3_06_RB0033** | aiGEM1143_B0033_fwd | CTCGTACTAGAGTCACACAGGACTAATCAATG | aiGEM1144_B0033_rev | CTCACATTGATTAGTCCTGTGTGACTCTAGTA | Annealing, GGA | / |
| **pMC0_3_07_RB0034** | aiGEM1007_B0034_fwd | CTCGTACTAGAGAAAGAGGAGAAATAATCAATG | aiGEM1008_B0034_rev | CTCACATTGATTATTTCTCCTCTTTCTCTAGTA | Annealing, GGA | / |
| **pMC0_3_08_RB0035** | aiGEM1145_B0035_fwd | CTCGTACTAGAGATTAAAGAGGAGAATAATCAATG | aiGEM1146_B0035_rev | CTCACATTGATTATTCTCCTCTTTAATCTCTAGTA | Annealing, GGA | / |
| **pMC0_3_09_RB0064** | aiGEM1147_B0064_fwd | CTCGTACTAGAGAAAGAGGGGAAATAATCAATG | aiGEM1148_B0064_rev | CTCACATTGATTATTCCCCTCTTTCTCTAGTA | Annealing, GGA | / |
| **pMC0_3_10_RDummy** | aiGEM1069_RBSDummy_fwd | CTCGTACTAGAGTGTCAGGATACCCGATAATCAATG | aiGEM1070_RBSDummy_rev | CTCACATTGATTATCGGGTATCCTGACACTCTAGTA | Annealing, GGA | / |
| **pMC0_3_11_3 Dropout mScarlet-I (Vn)** | oiGEM1128_GA_Entry_Vec_fwd | CCAGGCATCAAATAAAACGAAAGG | oiGEM1127_GA_Entry_Vec_rev | CTAGTATTTCTCCTCTTTCTCTAGTAG | PCR, Gib | pMC0_3_01_3 Dropout |
| | oiGEM1194_GA_Scarlet_fwd | ACTAGAGAAAGAGGAGAAATACTAGATGGTTTCTAAAGGTGAAGCAG | oiGEM1195_GA_Scarlet_rev | GCCTTTCGTTTTATTTGATGCCTGGTTATTTGTACAGTTCATCCATACC | | pMC0_4_12_CDSmScarlet-I (Vn) |
| **pMC0_4_01_4 Dropout** | oiGEM1137_Dropout_4_fwd | AACGTCTCGCTCGAATGAGAGACCGAAAGTGAAACGTGATTTCATGCG | oiGEM1138_Dropout_4_rev | TTCGTCTCCCTCAAAGCAGAGACCTATAAACGCAGAAAGGCCCACC | PCR, GGA | pMC_V_06_Part Entry sfGFP (Vn) Ori Muta |
| **pMC0_4_02_CDSlux Operon** | oiGEM1036_fwd_Lux Operon | AACGTCTCGCTCGAATGACTAAAAAAATTTCATTCATTATTAACGGCCAGG | oiGEM1037_rev_Lux Operon Muta | AACGTCTCGACGAGTCTCTTGTTTAGCTTTATTACTATCTTCG | PCR, GGA | pSV0-15_024 |
| | oiGEM1038_fwd_Lux Operon_Muta | AACGTCTCGTCGTGCATTTATTAGTGATTATGTTCTTGAAATGC | oiGEM1039_rev_Lux Operon | TTCGTCTCCCTCAAAGCACTATCAAACGCTTCGGTTAAGCTC | | pSV0-15_024 |
| **pMC0_4_09_CDSAzurite (Vn)** | / | / | / | / | DNA Synthesis, GGA | / |
| **pMC0_4_10_CDSmTurquoise (Vn)** | / | / | / | / | DNA Synthesis, GGA | / |
| **pMC0_4_11_CDSmVenus (Vn)** | / | / | / | / | DNA Synthesis, GGA | / |
| **pMC0_4_12_CDSmScarlet-I (Vn)** | / | / | / | / | DNA Synthesis, GGA | / |
| **pMC0_4_13_CDSmCherry (Vn)** | / | / | / | / | DNA Synthesis, GGA | / |
| **pMC0_4_14_CDSmKate-2 (Vn)** | / | / | / | / | DNA Synthesis, GGA | / |
| **pMC0_4_16_CDSVioA [#]** | oiGEM1181_VioA_fwd | AACGTCTCGCTCGAATGAGCACGTATTCTGACATTTGC | oiGEM1182_VioA_rev | TTCGTCTCCCTCAAAGCTGCGCGCTCGGTCGAGGAAG | PCR, GGA | pSwap (provided by Patrick Sobetzko) |
| **pMC0_4_17_CDSVioB [#]** | oiGEM1183_VioB_fwd | AACGTCTCGCTCGAATGAGCCTACTTGACTTCCCC | oiGEM1184_VioB_rev | TTCGTCTCCCTCAAAGCAGCCTCTCTTGACATCTTTCCC | PCR, GGA | pSwap (provided by Patrick Sobetzko) |
| **pMC0_4_18_CDSVioC [#]** | oiGEM1185_VioC_fwd | AACGTCTCGCTCGAATGCATAAAATCATTATCGTCGGC | oiGEM1186_VioC_rev | TTCGTCTCCCTCAAAGCATTTACCCTTCCAAGTTTGTACC | PCR, GGA | pSwap (provided by Patrick Sobetzko) |

| | | | | | | |
|---|---|---|---|---|---|---|
| **pMC0_4_20_CDSVioE [#]** | oiGEM1189_VioE_fwd | AACGTCTCGCTCGAATGCCGACACACGTCAGC | oiGEM1190_VioE_rev | TTCGTCTCCCTCAAAGCGGTGTTGCAAGACGTAAAGAC | PCR, GGA | pSwap (provided by Patrick Sobetzko) |
| **pMC0_4_21_4 Dropout mScarlet-I (Vn)** | oiGEM1128_GA_Entry_Vec_fwd | CCAGGCATCAAATAAAACGAAAGG | oiGEM1127_GA_Entry_Vec_rev | CTAGTATTTCTCCTCTTTCTCTAGTAG | PCR, Gib | pMC0_4_01_4 Dropout |
| | oiGEM1194_GA_Scarlet_fwd | ACTAGAGAAAGAGGAGAAATACTAGATGGTTTCTAAAGGTGAAGCAG | oiGEM1195_GA_Scarlet_rev | GCCTTTCGTTTTATTTGATGCCTGGTTATTTGTACAGTTCATCCATACC | | pMC0_4_12_CDSmScarlet-I (Vn) |
| **pMC0_4_22_ftsZ [#]** | oiGEM1214_ftsZ_4_fwd | AACGTCTCGCTCGAATGTTTGAACCGATGATGGAAATGTC | oiGEM1215_ftsZ_4_rev | TTCGTCTCCCTCAAAGCATCAGCCTGGCGACGCAAG | PCR, GGA | V. natriegens Genome |
| **pMC0_4_23_CDSsfgfp (Vn)** | / | / | / | / | DNA Synthesis, GGA | / |
| **pMC0_4a_04_N3xFLAG** | oiGEM1222_3xFLAG 4a_fwd | AACGTCTCGCTCGAATGGATTATAAGGATCATGATGGTGATTATAAGGATCATGATATCGACTAC | oiGEM1223_3xFLAG 4a_rev | TTCGTCTCCCTCACATCCCCTTGTCGTCATCGTCTTTGTAGTCGATATCATGATCCTTATAATC | Primer Extension, GGA | / |
| **pMC0_4a_05_N6xHis** | aiGEM1085_His_4x_fwd | CTCGAATGCACCATCACCACCATCATGGGATG | aiGEM1086_His_4x_rev | CTCACATCCCATGATGGTGGTGATGGTGCATT | Annealing, GGA | / |
| **pMC0_4a_06_NAzurite (Vn)** | oiGEM1198_Azurite_4a_fwd | AACGTCTCGCTCGAATGTCTAAAGGTGAAGAACTGTTTAC | oiGEM1202_Fluo_4a_rev | TTCGTCTCCCTCACATCCCTTTGTACAGTTCATCCATACC | PCR, GGA | pMC0_4_09_CDSAzurite (Vn) |
| **pMC0_4a_07_NmTurquoise 2 (Vn)** | oiGEM1199_Tur+Ven_4a_fwd | AACGTCTCGCTCGAATGGTTTCTAAAGGTGAAGAACTG | oiGEM1202_Fluo_4a_rev | TTCGTCTCCCTCACATCCCTTTGTACAGTTCATCCATACC | PCR, GGA | pMC0_4_10_CDSmTurquoise (Vn) |
| **pMC0_4a_09_NmVenus (Vn)** | oiGEM1199_Tur+Ven_4a_fwd | AACGTCTCGCTCGAATGGTTTCTAAAGGTGAAGAACTG | oiGEM1202_Fluo_4a_rev | TTCGTCTCCCTCACATCCCTTTGTACAGTTCATCCATACC | PCR, GGA | pMC0_4_11_CDSmVenus (Vn) |
| **pMC0_4a_10_NmScarlet-I (Vn)** | oiGEM1200_Scarlet_4a_fwd | AACGTCTCGCTCGAATGGTTTCTAAAGGTGAAGCAGTG | oiGEM1202_Fluo_4a_rev | TTCGTCTCCCTCACATCCCTTTGTACAGTTCATCCATACC | PCR, GGA | pMC0_4_12_CDSmScarlet-I (Vn) |
| **pMC0_4a_11_NmCherry (Vn)** | oiGEM1201_Cherry_4a_fwd | AACGTCTCGCTCGAATGGTTTCTAAAGGTGAAGAGGATAAC | oiGEM1202_Fluo_4a_rev | TTCGTCTCCCTCACATCCCTTTGTACAGTTCATCCATACC | PCR, GGA | pMC0_4_13_CDSmCherry (Vn) |
| **pMC0_4a_12_NmKate (Vn)** | oiGEM1203_Kate_4a_fwd | AACGTCTCGCTCGAATGGTTTCTGAACTGATTAAAGAAAACATG | oiGEM1204_Kate_4a_rev | TTCGTCTCCCTCACATCCCACGGTGACCCAGTTTAGATGG | PCR, GGA | pMC0_4_14_CDSmKate-2 (Vn) |
| **pMC0_4a_13_Nsfgfp(Vn)** | oiGEM1246_GFP(Vn)_new_4a_fwd | AACGTCTCGCTCGAATGCGTAAAGGTGAAGAACTGTTTACC | oiGEM1247_GFP(Vn)_new_4a_rev | TTCGTCTCCCTCACATCCCTGCTTTGTACAGTTCATCCATACC | PCR, GGA | pMC0_4_23_CDSsfgfp (Vn) |
| **pMC0_4b_01_ftsA [#]** | oiGEM1216_ftsA_4b_fwd | AACGTCTCGCTCGGATGACTAAGGCCGCAGACGAC | oiGEM1217_ftsA_4b_Muta_rev | TTCGTCTCCATAGACCAAGCGGGTTTTTAATCC | PCR, GGA | V. natriegens Genome |
| | oiGEM1218_ftsA_4b_Muta_fwd | TTCGTCTCCCTATCTGGGGTAAGAATGGAAG | oiGEM1219_ftsA_4b_rev | TTCGTCTCCCTCAAAGCAAACTCTTTTTGTATCCAGTTACGC | PCR, GGA | V. natriegens Genome |
| **pMC0_5_01_5 Dropout** | oiGEM1139_Dropout_5_fwd | AACGTCTCGCTCGGCTTAGAGACCGAAAGTGAAACGTGATTTCATGCG | oiGEM1140_Dropout_5_rev | TTCGTCTCCCTCAAGCGAGAGACCTATAAACGCAGAAAGGCCCACC | PCR, GGA | pMC_V_06_Part Entry sfGFP (Vn) Ori Muta |
| **pMC0_5_02_TB0010** | aiGEM1051_B0010a_fwd | CTCGGCTTAACCAGGCATCAAATAAAACGAAAGGCTCAGTCGAAA | aiGEM1052_B0010a_rev | AGTCTTTCGACTGAGCCTTTCGTTTTATTTGATGCCTGGTTAAGC | Annealing, GGA | / |
| | aiGEM1053_B0010b_fwd | GACTGGGCCTTTCGTTTTATCTGTTGTTTGTCGGTGAACGCTCTCCGCT | aiGEM1054_B0010b_rev | CTCAAGCGGAGAGCGTTCACCGACAAACAACAGATAAAACGAAAGGCCC | Annealing, GGA | / |
| **pMC0_5_03_TB0015** | oiGEM1017_fwd_B0015 | AACGTCTCGCTCGGCTTAACCAGGCATCAAATAAAACGAAAGGC | oiGEM1018_rev_B0015 | TTCGTCTCCCTCAAGCGTATAAACGCAGAAAGGCCCACCC | Primer Extension, GGA | / |
| **pMC0_5_04_TB1002** | aiGEM1093_B1002_5_fwd | CTCGGCTTAACGCAAAAAACCCCGCTTCGGCGGGGTTTTTTCGCCGCT | aiGEM1094_B1002_5_rev | CTCAAGCGGCGAAAAAACCCCGCCGAAGCGGGGTTTTTTGCGTTAAGC | Annealing, GGA | / |
| **pMC0_5_05_TB1003** | aiGEM1097_B1003_5_fwd | CTCGGCTTAACGCCAAAAACCCCGCTTCGGCGGGGTTTTTCCGCCGCT | aiGEM1098_B1003_5_rev | CTCAAGCGGCGGAAAAACCCCGCCGAAGCGGGGTTTTTGGCGTTAAGC | Annealing, GGA | / |
| **pMC0_5_06_TB1004** | aiGEM1149_B1004_fwd | CTCGGCTTAACGCCGAAAACCCCGCTTCGGCGGGGTTTTGCCGCCGCT | aiGEM1150_B1004_rev | CTCAAGCGGCGGCAAAACCCCGCCGAAGCGGGGTTTTCGGCGTTAAGC | Annealing, GGA | / |
| **pMC0_5_07_TB1005** | aiGEM1151_B1005_fwd | CTCGGCTTAACGCCGCAAACCCCGCTTCGGCGGGGTTTCGCCGCCGCT | aiGEM1152_B1005_rev | CTCAAGCGGCGGCGAAACCCCGCCGAAGCGGGGTTTGCGGCGTTAAGC | Annealing, GGA | / |

| | | | | | | |
|---|---|---|---|---|---|---|
| **pMC0_5_08_TB1006** | aiGEM1101_B1006_5_fwd | CTCGGCTTAAAAAAAAAACCCCGCCCCTGACAGGGCGGGGTTTTTTTTCGCT | aiGEM1102_B1006_5_rev | CTCAAGCGAAAAAAAACCCCGCCCTGTCAGGGGCGGGGTTTTTTTTTTAAGC | Annealing, GGA | / |
| **pMC0_5_09_TB1007** | aiGEM1153_B1007_fwd | CTCGGCTTAACGCAAAAAACCCCGCCCCTGACAGGGCGGGGTTTTTTCGCCGCT | aiGEM1154_B1007_rev | CTCAAGCGGCGAAAAAACCCCGCCCTGTCAGGGGCGGGGTTTTTTGCGTTAAGC | Annealing, GGA | / |
| **pMC0_5_11_TB1009** | aiGEM1157_B1009_fwd | CTCGGCTTAACGCCGAAAACCCCGCCCCTGACAGGGCGGGGTTTTGCCGCCGCT | aiGEM1158_B1009_rev | CTCAAGCGGCGGCAAAACCCCGCCCTGTCAGGGGCGGGGTTTTCGGCGTTAAGC | Annealing, GGA | / |
| **pMC0_5_12_TB1010** | aiGEM1159_B1010_fwd | CTCGGCTTAACGCCGCAAACCCCGCCCCTGACAGGGCGGGGTTTCGCCGCCGCT | aiGEM1160_B1010_rev | CTCAAGCGGCGGCGAAACCCCGCCCTGTCAGGGGCGGGGTTTGCGGCGTTAAGC | Annealing, GGA | / |
| **pMC0_5_13_TDummy**[#] | / | / | / | / | DNA Synthesis, GGA | / |
| **pMC0_5_14_5 Dropout mScarlet-I (Vn)** | oiGEM1128_GA_Entry_Vec_fwd | CCAGGCATCAAATAAAACGAAAGG | oiGEM1127_GA_Entry_Vec_rev | CTAGTATTTCTCCTCTTTCTCTAGTAG | PCR, Gib | pMC0_5_01_5 Dropout |
| | oiGEM1194_GA_Scarlet_fwd | ACTAGAGAAAGAGGAGAAATACTAGATGGTTTCTAAAGGTGAAGCAG | oiGEM1195_GA_Scarlet_rev | GCCTTTCGTTTTATTTGATGCCTGGTTATTTGTACAGTTCATCCATACC | | pMC0_4_12_CDSmScarlet-I (Vn) |
| **pMC0_5a_04_C3xFLAG** | oiGEM1224_3xFLAG 5a_fwd | AACGTCTCGCTCGGCTTTAGATTATAAGGATCATGATGGTGATTATAAGGATCATGATATCGACTAC | oiGEM1225_3xFLAG 5a_rev | TTCGTCTCCCTCATACCCCTTGTCGTCATCGTCTTTGTAGTCGATATCATGATCCTTATAATC | Primer Extension, GG | / |
| **pMC0_5a_05_C6xHis** | aiGEM1087_His_5a_fwd | CTCGGCTTTACACCATCACCACCATCATGGGTA | aiGEM1088_His_5a_rev | CTCATACCCATGATGGTGGTGATGGTGTAAAGC | Annealing, GGA | / |
| **pMC0_5a_06_CI11012** | aiGEM1131_I11012_fwd | CTCGGCTTTAGCAGCAAACGACGAAAACTACGCTGCTGCTGTTTAGGGGTA | aiGEM1132_I11012_rev | CTCATACCCCTAAACAGCAGCAGCGTAGTTTTCGTCGTTTGCTGCTAAAGC | Annealing, GGA | / |
| **pMC0_5a_07_CM0050** | aiGEM1133_M0050_fwd | CTCGGCTTTAGCTGCTAACGACGAAAACTACGCTCTGGCTGCTTAGGGGTA | aiGEM1134_M0050_rev | CCTCATACCCCTAAGCAGCCAGAGCGTAGTTTTCGTCGTTAGCAGCTAAAGC | Annealing, GGA | / |
| **pMC0_5a_08_CM0051** | aiGEM1135_M0051_fwd | CTCGGCTTTAGCTGCTAACGACGAAAACTACAACTACGCTGACGCTTCTTAGGGGTA | aiGEM1136_M0051_rev | CCTCATACCCCTAAGAAGCGTCAGCGTAGTTGTAGTTTTCGTCGTTAGCAGCTAAAGC | Annealing, GGA | / |
| **pMC0_5a_09_CM0052** | aiGEM1177_M0052_fwd | CTCGGCTTTAGCTGCTAACGACGAAAACTACGCTGACGCTTCTTAGGGGTA | aiGEM1183_M0052_rev | CTCATACCCCTAAGAAGCGTCAGCGTAGTTTTCGTCGTTAGCAGCTAAAGC | Annealing, GGA | / |
| **pMC0_5a_10_CAzurite (Vn)** | oiGEM1207_Azurite_5a_fwd | AACGTCTCGCTCGGCTTTATCTAAAGGTGAAGAACTGTTTAC | oiGEM1211_Fluo_5a_rev | TTCGTCTCCCTCATACCCTTTGTACAGTTCATCCATACC | PCR, GGA | pMC0_4_09_CDSAzurite (Vn) |
| **pMC0_5a_11_CmTurquoise (Vn)** | oiGEM1208_Tur+Ven_5a_fwd | AACGTCTCGCTCGGCTTTAGTTTCTAAAGGTGAAGAACTG | oiGEM1211_Fluo_5a_rev | TTCGTCTCCCTCATACCCTTTGTACAGTTCATCCATACC | PCR, GGA | pMC0_4_10_CDSmTurquoise (Vn) |
| **pMC0_5a_13_CmVenus (Vn)** | oiGEM1208_Tur+Ven_5a_fwd | AACGTCTCGCTCGGCTTTAGTTTCTAAAGGTGAAGAACTG | oiGEM1211_Fluo_5a_rev | TTCGTCTCCCTCATACCCTTTGTACAGTTCATCCATACC | PCR, GGA | pMC0_4_11_CDSmVenus (Vn) |
| **pMC0_5a_14_CmScarlet-I (Vn)** | oiGEM1209_Scarlet_5a_fwd | AACGTCTCGCTCGGCTTTAGTTTCTAAAGGTGAAGCAGTG | oiGEM1211_Fluo_5a_rev | TTCGTCTCCCTCATACCCTTTGTACAGTTCATCCATACC | PCR, GGA | pMC0_4_12_CDSmScarlet-I (Vn) |
| **pMC0_5a_15_CmCherry (Vn)** | oiGEM1210_Cherry_5a_fwd | AACGTCTCGCTCGGCTTTAGTTTCTAAAGGTGAAGAGGATAAC | oiGEM1211_Fluo_5a_rev | TTCGTCTCCCTCATACCCTTTGTACAGTTCATCCATACC | PCR, GGA | pMC0_4_13_CDSmCherry (Vn) |
| **pMC0_5a_16_CmKate-2 (Vn)** | oiGEM1212_Kate_5a_fwd | AACGTCTCGCTCGGCTTTAGTTTCTGAACTGATTAAAGAAAACATG | oiGEM1213_Kate_5a_rev | TTCGTCTCCCTCATACCCACGGTGACCCAGTTTAGATGG | PCR, GGA | pMC0_4_14_CDSmKate-2 (Vn) |
| **pMC0_5a_17_CsfGFP(Vn)** | oiGEM1248_GFP(Vn)_new_5a_fwd | AACGTCTCGCTCGGCTTTACGTAAAGGTGAAGAACTGTTTACC | oiGEM1249_GFP(Vn)_new_5a_rev | TTCGTCTCCCTCATACCCTGCTTTGTACAGTTCATCCATACC | PCR, GGA | pMC0_4_23_CDSsfgfp (Vn) |
| **pMC0_5b_01_TB0010** | oiGEM1143_B0010_5b_fwd | AACGTCTCGCTCGGGTAACCAGGCATCAAATAAAACGAAAGGCTCAGTCGAAAGACTGGGCCTTTC | oiGEM1144_B0010_5b_rev | TTCGTCTCCCTCAAGCGGAGAGCGTTCACCGACAAACAACAGATAAAACGAAAGGCCCAGTCTTTCGAC | Primer Extension, GG | / |
| **pMC0_5b_02_TB0015** | oiGEM1058_fwd_B0015_5b | AACGTCTCGCTCGGGTAACCAGGCATCA | oiGEM1018_rev_B0015 | TTCGTCTCCCTCAAGCGTATAAACGCAGAAAGGCCCACCC | Primer Extension, GG | / |

| | | | | | | |
|---|---|---|---|---|---|---|
| | | AATAAAACGAAAGGCTC | | | | |
| **pMC0_5b_03_TB1002** | aiGEM1095_B1002_5b_fwd | CTCGGGTAACGCAAAAAACCCCGCTTCGGCGGGGTTTTTTCGCCGCT | aiGEM1096_B1002_5b_rev | CTCAAGCGGCGAAAAAACCCCGCCGAAGCGGGGTTTTTTGCGTTACC | Annealing, GGA | / |
| **pMC0_5b_04_TB1003** | aiGEM1099_B1003_5b_fwd | CTCGGGTAACGCCAAAAACCCCGCTTCGGCGGGGTTTTTCCGCCGCT | aiGEM1100_B1003_5b_rev | CTCAAGCGGCGGAAAAACCCCGCCGAAGCGGGGTTTTTGGCGTTACC | Annealing, GGA | / |
| **pMC0_5b_05_TB1004** | aiGEM1161_B1004_5b_fwd | CTCGGGTAACGCCGAAAACCCCGCTTCGGCGGGGTTTTGCCGCCGCT | aiGEM1162_B1004_5b_rev | CTCAAGCGGCGGCAAAACCCCGCCGAAGCGGGGTTTTCGGCGTTACC | Annealing, GGA | / |
| **pMC0_5b_06_TB1005** | aiGEM1163_B1005_5b_fwd | CTCGGGTAACGCCGCAAACCCCGCTTCGGCGGGGTTTCGCCGCCGCT | aiGEM1164_B1005_5b_rev | CTCAAGCGGCGGCGAAACCCCGCCGAAGCGGGGTTTGCGGCGTTACC | Annealing, GGA | / |
| **pMC0_5b_07_TB1006** | aiGEM1103_B1006_5b_fwd | CTCGGGTAAAAAAAAAACCCCGCCCCTGACAGGGCGGGGTTTTTTTTCGCT | aiGEM1104_B1006_5b_rev | CTCAAGCGAAAAAAAACCCCGCCCTGTCAGGGGCGGGGTTTTTTTTTTACC | Annealing, GGA | / |
| **pMC0_5b_08_TB1007** | aiGEM1165_B1007_5b_fwd | CTCGGGTAACGCAAAAAACCCCGCCCCTGACAGGGCGGGGTTTTTTCGCCGCT | aiGEM1166_B1007_5b_rev | CTCAAGCGGCGAAAAAACCCCGCCCTGTCAGGGGCGGGGTTTTTTGCGTTACC | Annealing, GGA | / |
| **pMC0_5b_10_TB1009** | aiGEM1169_B1009_5b_fwd | CTCGGGTAACGCCGAAAACCCCGCCCCTGACAGGGCGGGGTTTTGCCGCCGCT | aiGEM1170_B1009_5b_rev | CTCAAGCGGCGGCAAAACCCCGCCCTGTCAGGGGCGGGGTTTTCGGCGTTACC | Annealing, GGA | / |
| **pMC0_5b_11_TB1010** | aiGEM1171_B1010_5b_fwd | CTCGGGTAACGCCGCAAACCCCGCCCCTGACAGGGCGGGGTTTCGCCGCCGCT | aiGEM1172_B1010_5b_rev | CTCAAGCGGCGGCGAAACCCCGCCCTGTCAGGGGCGGGGTTTGCGGCGTTACC | Annealing, GGA | / |
| **pMC0_6_01_6 Dropout** | oiGEM1141_Dropout_6_fwd | AACGTCTCGCTCGCGCTAGAGACCGAAAGTGAAACGTGATTTCATGCG | oiGEM1142_Dropout_6_rev | TTCGTCTCCCTCAAGCTAGAGACCTATAAACGCAGAAAGGCCCACC | PCR, GGA | pMC_V_06_Part Entry sfGFP (Vn) Ori Muta |
| **pMC0_6_02_3C1SF** | aiGEM1057_3'Con1_fwd | CTCGCGCTTACTGGAGACGAGCT | aiGEM1058_3'Con1_rev | CTCAAGCTCGTCTCCAGTAAGCG | Annealing, GGA | / |
| **pMC0_6_03_3C1CLR**[#] | oiGEM1100_Aqua_Inver_Conn_fwd | GGAGACGAGCTTGAGACCACTCATCGCGACTAG | oiGEM1099_Aqua_Invert_Conn_rev | AGCGCGAGACCACTCATCGCTCTAG | PCR, Gib | pMC0_1_03_5C1RLN |
| | oiGEM1101_Aqua_3Con1_inv_fwd | GCCGCTTCTAGAGCGATGAGTGGTCTCGCGCTGGACCAAAACGAAAAAAGACGC | oiGEM1102_Aqua_3Con1_inv_rev | TAGTCGCGATGAGTGGTCTCAAGCTCGTCTCCGGTGTTTTGTTATCAATAAAAAAGGCCCCC | | pMC0_1_03_5C1RLN |
| | oiGEM1250_GA_6_03_6_05_fwd | CTCCGGAGACGAGCTTGAGAC | oiGEM1251_GA_6_03_rev | AGTGGTCTCAAGCTCGTCTCCGGAGTTTTGTTATCAATAAAAAAGGC | PCR, Gib | Mutation in previous part was corrected |
| **pMC0_6_04_3C1RLR**[#] | oiGEM1100_Aqua_Inver_Conn_fwd | GGAGACGAGCTTGAGACCACTCATCGCGACTAG | oiGEM1099_Aqua_Invert_Conn_rev | AGCGCGAGACCACTCATCGCTCTAG | PCR, Gib | pMC0_1_03_5C1RLN |
| | oiGEM1101_Aqua_3Con1_inv_fwd | GCCGCTTCTAGAGCGATGAGTGGTCTCGCGCTGGACCAAAACGAAAAAAGACGC | oiGEM1103_Aqua_3Con1_inv_Res_rev | TAGTCGCGATGAGTGGTCTCAAGCTCGTCTCCAACATTTTGTTATCAATAAAAAAGGCCCCC | PCR, Gib | pMC0_1_03_5C1RLN |
| **pMC0_6_05_3C1CSR** | aiGEM1119_3'Con1_inv_Short_fwd | CTCGCGCTCACCGGAGACGAGCT | aiGEM1120_3'Con1_inv_Short_rev | CTCAAGCTCGTCTCCGGTGAGCG | Annealing, GGA | / |
| | oiGEM1250_GA_6_03_6_05_fwd | CTCCGGAGACGAGCTTGAGAC | oiGEM1252_GA_6_05_rev | AGTGGTCTCAAGCTCGTCTCCGGAGAGCGCGAGACCACTCATCGC | PCR, Gib | Mutation in previous part was corrected |
| **pMC0_6_06_3C1RSR** | aiGEM1129_3'Con1_inv_Short_Res_fwd | CTCGCGCTTGTTGGAGACGAGCT | aiGEM1130_3'Con1_inv_Short_Res_rev | CTCAAGCTCGTCTCCAACAAGCG | Annealing, GGA | / |
| **pMC0_6_07_3C2SF** | aiGEM1059_3'Con2_fwd | CTCGCGCTAATGGGAGACGAGCT | aiGEM1060_3'Con2_rev | CTCAAGCTCGTCTCCCATTAGCG | Annealing, GGA | / |
| **pMC0_6_08_3C2LR**[#] | oiGEM1100_Aqua_Inver_Conn_fwd | GGAGACGAGCTTGAGACCACTCATCGCGACTAG | oiGEM1099_Aqua_Invert_Conn_rev | AGCGCGAGACCACTCATCGCTCTAG | PCR, Gib | pMC0_1_03_5C1RLN |
| | oiGEM1104_Aqua_3Con2_inv_fwd | GCCGCTTCTAGAGCGATGAGTGGTCTCGCGCTGGACCAAAACGAAAAAAGGCCG | oiGEM1105_Aqua_3Con2_inv_rev | TAGTCGCGATGAGTGGTCTCAAGCTCGTCTCCTACTGGACCAAAACGAAAAAAGACGC | PCR, Gib | pMC0_1_07_5C2LN |
| **pMC0_6_09_3C2SR** | aiGEM1121_3'Con2_inv_Short_fwd | CTCGCGCTAGTAGGAGACGAGCT | aiGEM1122_3'Con2_inv_Short_rev | CTCAAGCTCGTCTCCTACTAGCG | Annealing, GGA | / |

| | | | | | | |
|---|---|---|---|---|---|---|
| **pMC0_6_10_3C3SF** | aiGEM1061_3'Con3_fwd | CTCGCGCTGCTTGGAGACGAGCT | aiGEM1062_3'Con3_rev | CTCAAGCTCGTCTCCAAGCAGCG | Annealing, GGA | / |
| **pMC0_6_11_3C3LR** # | oiGEM1100_Aqua_Inver_Conn_fwd | GGAGACGAGCTTGAGACCACTCATCGCGACTAG | oiGEM1099_Aqua_Invert_Conn_rev | AGCGCGAGACCACTCATCGCTCTAG | PCR, Gib | pMC0_1_03_5C1RLN |
| | oiGEM1101_Aqua_3Con1_inv_fwd | GCCGCTTCTAGAGCGATGAGTGGTCTCGCGCTGGACCAAAACGAAAAAAGACGC | oiGEM1106_Aqua_3Con3_inv_rev | TAGTCGCGATGAGTGGTCTCAAGCTCGTCTCCAATGTTTTGTTATCAATAAAAAAGGCCCCCC | PCR, Gib | pMC0_1_10_5C3LN |
| **pMC0_6_12_3C3SR** | aiGEM1123_3'Con3_inv_Short_fwd | CTCGCGCTCATTGGAGACGAGCT | aiGEM1124_3'Con3_inv_Short_rev | CTCAAGCTCGTCTCCAATGAGCG | Annealing, GGA | / |
| **pMC0_6_13_3C4SF** | aiGEM1063_3'Con4_fwd | CTCGCGCTGGTAGGAGACGAGCT | aiGEM1064_3'Con4_rev | CTCAAGCTCGTCTCCTACCAGCG | Annealing, GGA | / |
| **pMC0_6_14_3C4LR** # | oiGEM1100_Aqua_Inver_Conn_fwd | GGAGACGAGCTTGAGACCACTCATCGCGACTAG | oiGEM1099_Aqua_Invert_Conn_rev | AGCGCGAGACCACTCATCGCTCTAG | PCR, Gib | pMC0_1_03_5C1RLN |
| | oiGEM1101_Aqua_3Con1_inv_fwd | GCCGCTTCTAGAGCGATGAGTGGTCTCGCGCTGGACCAAAACGAAAAAAGACGC | oiGEM1107_Aqua_3Con4_inv_rev | TAGTCGCGATGAGTGGTCTCAAGCTCGTCTCCGCTTTTTTGTTATCAATAAAAAAGGCCCCCC | PCR, Gib | pMC0_1_13_5C4LN |
| **pMC0_6_15_3C4SR** | aiGEM1125_3'Con4_inv_Short_fwd | CTCGCGCTAAGCGGAGACGAGCT | aiGEM1126_3'Con4_inv_Short_rev | CTCAAGCTCGTCTCCGCTTAGCG | Annealing, GGA | / |
| **pMC0_6_16_3C5CSF** | aiGEM1065_3'Con5_fwd | CTCGCGCTCGCTGGAGACGAGCT | aiGEM1066_3'Con5_rev | CTCAAGCTCGTCTCCAGCGAGCG | Annealing, GGA | / |
| **pMC0_6_17_3C5OSF** | aiGEM1083_3'Con_5_Ori_fwd | CTCGCGCTAGCTGGAGACGAGCT | aiGEM1084_3'Con_5_Ori_rev | CTCAAGCTCGTCTCCAGCTAGCG | Annealing, GGA | / |
| **pMC0_6_18_3C5LR** # | oiGEM1100_Aqua_Inver_Conn_fwd | GGAGACGAGCTTGAGACCACTCATCGCGACTAG | oiGEM1099_Aqua_Invert_Conn_rev | AGCGCGAGACCACTCATCGCTCTAG | PCR, Gib | pMC0_1_03_5C1RLN |
| | oiGEM1108_Aqua_3Con5_inv_fwd | GCCGCTTCTAGAGCGATGAGTGGTCTCGCGCTTTTTGTTATCAATAAAAAAGGCCCCCC | oiGEM1109_Aqua_3Con5_inv_rev | TAGTCGCGATGAGTGGTCTCAAGCTCGTCTCCGGTATTTTGTTATCAATAAAAAAGGCCCCCC | PCR, Gib | pMC0_1_16_5C5LN |
| **pMC0_6_19_3C5SR** | aiGEM1127_3'Con5_inv_Short_fwd | CTCGCGCTTACCGGAGACGAGCT | aiGEM1128_3'Con5_inv_Short_rev | CTCAAGCTCGTCTCCGGTAAGCG | Annealing, GGA | / |
| **pMC0_6_20_3C0SF** | aiGEM1019_3'Conn._Dummy_fwd | CTCGCGCTTACTAGTAGCGGCCGCTGCAGAGCT | aiGEM1020_3'Conn._Dummy_rev | CTCAAGCTCTGCAGCGGCCGCTACTAGTAAGCG | Annealing, GGA | / |
| **pMC0_6_21_6 Dropout mScarlet-I (Vn)** | oiGEM1128_GA_Entry_Vec_fwd | CCAGGCATCAAATAAAACGAAAGG | oiGEM1127_GA_Entry_Vec_rev | CTAGTATTTCTCCTCTTTCTCTAGTAG | PCR, Gib | pMC0_5_01_5 Dropout |
| | oiGEM1194_GA_Scarlet_fwd | ACTAGAGAAAGAGGAGAAATACTAGATGGTTTCTAAAGGTGAAGCAG | oiGEM1195_GA_Scarlet_rev | GCCTTTCGTTTTATTTGATGCCTGGTTATTTGTACAGTTCATCCATACC | | pMC0_4_12_CDSmScarlet-I (Vn) |
| **pMC0_7_01_OColE1** | oiGEM1025_fwd_ColE1 | AACGTCTCGCTCGAGCTGAACCGTAAAAAGGCCGCGTTGC | oiGEM1026_rev_ColE1 | TTCGTCTCCCTCAAGCAGTGAAGATCCTTTTTGATAATCTCATGACC | PCR, GGA | pSB1C3 |
| **pMC0_7_02_Op15A** | oiGEM1050_fwd_p15A | AACGTCTCGCTCGAGCTGCGCTAGCGGAGTGTATAC | oiGEM1051_rev_p15A | TTCGTCTCCCTCAAGCAAATTTAAAAGGATCTAGGTGAAGATCC | PCR, GGA | pSB3C5 |
| **pMC0_7_04_OpMB1-M** | oiGEM1169_7_04_OpMB1-M_fwd | AACGTCTCGCTCGAGCTACGGTTATCCACAGAATCAGGG | oiGEM1170_7_04_OpMB1-M_rev | TTCGTCTCCCTCAAGCATCATGACCAAAATCCCTTAACGTG | PCR, GGA | pMC_V_06 |
| **pMC0_7_05_ORSF1010** | oiGEM1192_pMMB Ori_fwd | AACGTCTCGCTCGAGCTTTCTGAAAGCGACCAGGTGC | oiGEM1193_pMMB Ori_rev | TTCGTCTCCCTCAAGCAGCGCAACGCAATTAATGTAAGTTAG | PCR, GGA | pMMB-Tfox (provided by Ankur Dalia) |
| **pMC0_8_09_Akan (sfgfp)(Vn)** | oiGEM1046_fwd_Kan_Res | AACGTCTCGCTCGTGCTAAACGGGCAAGGTGTCACC | oiGEM1047_rev_KanRes_Muta | AACGTCTCGCTGGCTCAGGCGCAATCACG | PCR, GGA | pSB1K3 |
| | oiGEM1048_fwd_Kan_Res_Muta | AACGTCTCGCCAGACGAAATACGCGATCG | oiGEM1049_rev_Kan_Res | TTCGTCTCCCTCATGTTAAGATGCGTGATCTGATCCTTC | PCR, GGA | pSB1K3 |
| **pMC0_8_10_Akan (mScarlet-I)(Vn)** | oiGEM1046_fwd_Kan_Res | AACGTCTCGCTCGTGCTAAACGGGCAAGGTGTCACC | oiGEM1047_rev_KanRes_Muta | AACGTCTCGCTGGCTCAGGCGCAATCACG | PCR, GGA | pSB1K3 |
| | oiGEM1048_fwd_Kan_Res_Muta | AACGTCTCGCCAGACGAAATACGCGATCG | oiGEM1049_rev_Kan_Res | TTCGTCTCCCTCATGTTAAGATGCGTGATCTGATCCTTC | PCR, GGA | pSB1K3 |
| **pMC0_8_11_Atet (sfgfp)(Vn)** | oiGEM1049_rev_Kan_Res | TTCGTCTCCCTCATGTTAAGATGCGTGATCTGATCCTTC | oiGEM1054_rev_Tet from pSB3T5 | AACGTCTCGCTCGTGCTGAGTCACTAAGGGCTAACTAAC | PCR, GGA | pSB3T5 |
| **pMC0_8_12_Atet (mScarlet-I)(Vn)** | oiGEM1049_rev_Kan_Res | TTCGTCTCCCTCATGTTAAGATGCGTGATCTGATCCTTC | oiGEM1054_rev_Tet from pSB3T5 | AACGTCTCGCTCGTGCTGAGTCACTAAGGGCTAACTAAC | PCR, GGA | pSB3T5 |

| | | | | | | |
|---|---|---|---|---|---|---|
| **pMC0_8_13_Acam (sfgfp)(Vn)** | oiGEM1093_CamRes_new_fwd | AACGTCTCGCTCGTGCTACCAATAAAAAACGCCCGGCG | oiGEM1094_CamRes_new_rev | TTCGTCTCCCTCATGTTTTGATCGGGCACGTAAGAGG | PCR, GGA | pSB1C3 |
| **pMC0_8_14_Acam (mScarlet-I)(Vn)** | oiGEM1093_CamRes_new_fwd | AACGTCTCGCTCGTGCTACCAATAAAAAACGCCCGGCG | oiGEM1094_CamRes_new_rev | TTCGTCTCCCTCATGTTTTGATCGGGCACGTAAGAGG | PCR, GGA | pSB1C3 |
| **pMC0_8_15_Acarb (sfgfp)(Vn)** | oiGEM1027_fwd_Amp+Cam_Res | AACGTCTCGCTCGTGCTGATTATCAAAAAGGATCTTCACCTAGATCC | oiGEM1030_rev_Amp_Muta | AACGTCTCAGGGCTCGCGGTATCATTGCAGCACTGG | PCR, GGA | pSB1A2 |
| | oiGEM1029_fwd_Amp_Muta | AACGTCTCAGCCCCACGCTCACCGGCTCCAG | oiGEM1028_rev_Amp+Cam_Res | TTCGTCTCCCTCATGTTCAGAAATCATCCTTAGCGAAAGCTAAGG | | pSB1A2 |
| **pMC0_8_16_Acarb (mScarlet-I)(Vn)** | oiGEM1027_fwd_Amp+Cam_Res | AACGTCTCGCTCGTGCTGATTATCAAAAAGGATCTTCACCTAGATCC | oiGEM1030_rev_Amp_Muta | AACGTCTCAGGGCTCGCGGTATCATTGCAGCACTGG | PCR, GGA | pSB1A2 |
| | oiGEM1029_fwd_Amp_Muta | AACGTCTCAGCCCCACGCTCACCGGCTCCAG | oiGEM1028_rev_Amp+Cam_Res | TTCGTCTCCCTCATGTTCAGAAATCATCCTTAGCGAAAGCTAAGG | | pSB1A2 |
| **pMC0_8_18_Akan(Vn) (sfgfp)(Vn)** | oiGEM1232_GA_Kan_Codon_fwd | TGTGAGACATAACACCCCTTGTATTACTGTTTATG | oiGEM1233_GA_Kan_Codon_rev | GTTCTTTTAATCAGAATTGGTTAATTGGTTGTAACAC | PCR, DNA Synthesis, Gib | pMC0_8_09_Akan (sfgfp)(Vn) |
| **pMC0_8_19_Akan(Vn) (mScarlet)(Vn)** | oiGEM1232_GA_Kan_Codon_fwd | TGTGAGACATAACACCCCTTGTATTACTGTTTATG | oiGEM1233_GA_Kan_Codon_rev | GTTCTTTTAATCAGAATTGGTTAATTGGTTGTAACAC | PCR, DNA Synthesis, Gib | pMC0_8_10_Akan (sfgfp)(Vn) |
| **pMC0_8_21_Acarb J23106 (sfgfp)(Vn)** | oiGEM1244_GA_Carb_J23106_fwd | CTATACCTAGGACTGAGCTAGCCGTAAACTCCAACATGAGACC | oiGEM1245_GA_Carb_J23106_rev | CGGCTAGCTCAGTCCTAGGTATAGTGCTAGCTACTAGAGAAAGAGG | PCR, Gib | pMC0_8_15_Acarb (sfgfp)(Vn) |
| **pMC0_8_23_Acarb J23106 (mScarlet-I)(Vn)** | oiGEM1244_GA_Carb_J23106_fwd | CTATACCTAGGACTGAGCTAGCCGTAAACTCCAACATGAGACC | oiGEM1245_GA_Carb_J23106_rev | CGGCTAGCTCAGTCCTAGGTATAGTGCTAGCTACTAGAGAAAGAGG | PCR, Gib | pMC0_8_16_Acarb (mScarlet-I)(Vn) |
| **pMC0_1*_01_5C1C** | aiGEM1184_5C_01_5C1C_fwd | CTCGAACAGGTCTCGGGAGGGAG | aiGEM1185_5C_01_5C1C_rev | CTCACTCCCTCCCGAGACCTGTT | Annealing, GGA | / |
| **pMC0_1*_02_5C1R** | aiGEM1186_5C_02_5C1R_fwd | CTCGAACAGGTCTCGAACAGGAG | aiGEM1187_5C_02_5C1R_rev | CTCACTCCTGTTCGAGACCTGTT | Annealing, GGA | / |
| **pMC0_1*_03_5C2** | aiGEM1188_5C_03_5C2_fwd | CTCGAACAGGTCTCGTACTGGAG | aiGEM1189_5C_03_5C2_rev | CTCACTCCAGTACGAGACCTGTT | Annealing, GGA | / |
| **pMC0_1*_04_5C3** | aiGEM1190_5C_04_5C3_fwd | CTCGAACAGGTCTCGAATGGGAG | aiGEM1191_5C_04_5C3_rev | CTCACTCCCATTCGAGACCTGTT | Annealing, GGA | / |
| **pMC0_1*_05_5C4** | aiGEM1192_5C_05_5C4_fwd | CTCGAACAGGTCTCGGCTTGGAG | aiGEM1193_5C_05_5C4_rev | CTCACTCCAAGCCGAGACCTGTT | Annealing, GGA | / |
| **pMC0_1*_06_5C5** | aiGEM1194_3C_06_5C5_fwd | CTCGAACAGGTCTCGGGTAGGAG | aiGEM1195_3C_06_5C5_rev | CTCACTCCTACCCGAGACCTGTT | Annealing, GGA | / |
| **pMC0_6*_01_3C1** | aiGEM1196_3C_01_3C1_fwd | CTCGCGCTTACTGGAGACCAGCT | aiGEM1197_3C_01_3C1_rev | CTCAAGCTGGTCTCCAGTAAGCG | Annealing, GGA | / |
| **pMC0_6*_02_3C2** | aiGEM1198_3C_02_3C2_fwd | CTCGCGCTAATGGGAGACCAGCT | aiGEM1199_3C_02_3C2_rev | CTCAAGCTGGTCTCCCATTAGCG | Annealing, GGA | / |
| **pMC0_6*_03_3C3** | aiGEM1200_3C_03_3C3_fwd | CTCGCGCTGCTTGGAGACCAGCT | aiGEM1201_3C_03_3C3_rev | CTCAAGCTGGTCTCCAAGCAGCG | Annealing, GGA | / |
| **pMC0_6*_04_3C4** | aiGEM1202_3C_04_3C4_fwd | CTCGCGCTGGTAGGAGACCAGCT | aiGEM1203_3C_04_3C4_rev | CTCAAGCTGGTCTCCTACCAGCG | Annealing, GGA | / |
| **pMC0_6*_05_3C5C** | aiGEM1204_3C_05_3C5C_fwd | CTCGCGCTCGCTGGAGACCAGCT | aiGEM1205_3C_05_3C5C_rev | CTCAAGCTGGTCTCCAGCGAGCG | Annealing, GGA | / |
| **pMC0_6*_06_3C5O** | aiGEM1206_3C_06_3C5O_fwd | CTCGCGCTAGCTGGAGACCAGCT | aiGEM1207_3C_06_3C5O_rev | CTCAAGCTGGTCTCCAGCTAGCG | Annealing, GGA | / |
| **pMC0_7*_01_OColE1** | oiGEM1025_fwd_ColE1 | AACGTCTCGCTCGAGCTGAACCGTAAAAAGGCCGCGTTGC | oiGEM1026_rev_ColE1 | TTCGTCTCCCTCAAGCAGTGAAGATCCTTTTTGATAATCTCATGACC | PCR, Digest, GGA | pMC0_7_01_OColE1 |
| **pMC0_7*_02_Op15A** | oiGEM1050_fwd_p15A | AACGTCTCGCTCGAGCTGCGCTAGCGGAGTGTATAC | oiGEM1051_rev_p15A | TTCGTCTCCCTCAAGCAAATTTAAAAGGATCTAGGTGAAGATCC | PCR, Digest, GGA | pMC0_7_02_Op15A |
| **pMC0_7*_04_OpMB1-M** | oiGEM1169_7_04_OpMB1-M_fwd | AACGTCTCGCTCGAGCTACGGTTATCCACAGAATCAGGG | oiGEM1170_7_04_OpMB1-M_rev | TTCGTCTCCCTCAAGCATCATGACCAAAATCCCTTAACGTG | PCR, Digest, GGA | pMC0_7_04_OpMB1-M |
| **pMC0_7*_05_RSF1010** | oiGEM1192_pMMB Ori_fwd | AACGTCTCGCTCGAGCTTTCTGAAAGCGACCAGGTGC | oiGEM1193_pMMB Ori_rev | TTCGTCTCCCTCAAGCAGCGCAACGCAATTAATGTAAGTTAG | PCR, Digest, GGA | pMC0_7_05_OpMMB |
| **pMC0_8*_03_ATet (sfGFP)** | oiGEM1049_rev_Kan_Res | TTCGTCTCCCTCATGTTAAGATGCGTGATCTGATCCTTC | oiGEM1054_rev_Tet from pSB3T5 | AACGTCTCGCTCGTGCTGAGTCACTAAGGGCTAACTAAC | PCR, Digest, GGA | pMC0_8_11_Atet (sfgfp)(Vn) |
| **pMC0_8*_04_ATet (mScarlet-I)** | oiGEM1049_rev_Kan_Res | TTCGTCTCCCTCATGTTAAGATGCGTGATCTGATCCTTC | oiGEM1054_rev_Tet from pSB3T5 | AACGTCTCGCTCGTGCTGAGTCACTAAGGGCTAACTAAC | PCR, Digest, GGA | pMC0_8_12_Atet (mScarlet-I)(Vn) |

| | | | | | | |
|---|---|---|---|---|---|---|
| **pMC0_8*_05_ACam (sfGFP)** | oiGEM1093_CamRes_new_fwd | AACGTCTCGCTCGTGCTACCAATAAAAAACGCCCGGCG | oiGEM1094_CamRes_new_rev | TTCGTCTCCCTCATGTTTTGATCGGGCACGTAAGAGG | PCR, Digest, GGA | pMC0_8_13_Acam (sfgfp)(Vn) |
| **pMC0_8*_06_ACam (mScarlet-I)** | oiGEM1093_CamRes_new_fwd | AACGTCTCGCTCGTGCTACCAATAAAAAACGCCCGGCG | oiGEM1094_CamRes_new_rev | TTCGTCTCCCTCATGTTTTGATCGGGCACGTAAGAGG | PCR, Digest, GGA | pMC0_8_14_Acam (mScarlet-I)(Vn) |
| **pMC0_8*_09_AKan (Vn)(sfGFP)** | oiGEM1232_GA_Kan_Codon_fwd | TGTGAGACATAACACCCCTTGTATTACTGTTTATG | oiGEM1233_GA_Kan_Codon_rev | GTTCTTTTAATCAGAATTGGTTAATTGGTTGTAACAC | PCR, Digest, GGA | pMC0_8_18_Akan(Vn) (sfgfp)(Vn) |
| **pMC0_8*_10_AKan (Vn)(mScarlet-I)** | oiGEM1232_GA_Kan_Codon_fwd | TGTGAGACATAACACCCCTTGTATTACTGTTTATG | oiGEM1233_GA_Kan_Codon_rev | GTTCTTTTAATCAGAATTGGTTAATTGGTTGTAACAC | PCR, Digest, GGA | pMC0_8_19_Akan(Vn) (mScarlet)(Vn) |
| **pMC0_8*_12_ACarb J23106 (sfGFP)** | oiGEM1244_GA_Carb_J23106_fwd | CTATACCTAGGACTGAGCTAGCCGTAAACTCCAACATGAGACC | oiGEM1245_GA_Carb_J23106_rev | CGGCTAGCTCAGTCCTAGGTATAGTGCTAGCTACTAGAGAAAGAGG | PCR, Digest, GGA | pMC0_8_15_Acarb (sfgfp)(Vn) |
| **pMC0_8*_14_ACarb J23106 (mScarlet-I)** | oiGEM1244_GA_Carb_J23106_fwd | CTATACCTAGGACTGAGCTAGCCGTAAACTCCAACATGAGACC | oiGEM1245_GA_Carb_J23106_rev | CGGCTAGCTCAGTCCTAGGTATAGTGCTAGCTACTAGAGAAAGAGG | PCR, Digest, GGA | pMC0_8_16_Acarb (mScarlet-I)(Vn) |
| **pMC0_TU2-6*_EL_01** | aiGEM1208_EL_01_EL5C2-3C5O_fwd | CTCGTACTGCTGCGATTAGACAGCCAGCAGCT | aiGEM1209_EL_01_EL5C2-3C5O_rev | CTCAAGCTGCTGGCTGTCTAATCGCAGCAGTA | Annealing, GGA | / |
| **pMC0_TU3-6*_EL_02** | aiGEM1210_EL_02_EL5C3-3C5O_fwd | CTCGAATGGCTGCGATTAGACAGCCAGCAGCT | aiGEM1211_EL_02_EL5C3-3C5O_rev | CTCAAGCTGCTGGCTGTCTAATCGCAGCCATT | Annealing, GGA | / |
| **pMC0_TU4-6*_EL_03** | aiGEM1212_EL_03_EL5C4-3C5O_fwd | CTCGGCTTGCTGCGATTAGACAGCCAGCAGCT | aiGEM1213_EL_03_EL5C4-3C5O_rev | CTCAAGCTGCTGGCTGTCTAATCGCAGCAAGC | Annealing, GGA | / |
| **pMC0_TU5-6*_EL_04** | aiGEM1214_EL_04_EL5C5-3C5O_fwd | CTCGGGTAGCTGCGATTAGACAGCCAGCAGCT | aiGEM1215_EL_04_EL5C5-3C5O_rev | CTCAAGCTGCTGGCTGTCTAATCGCAGCTACC | Annealing, GGA | / |
| **pMC0_1*-6*_01_Dropout sfGFP** | oiGEM1179_Dropout_5C1R_fwd | AAGGTCTCGCTCGAACAAGAGACGGAAAGTGAAACGTGATTTCATGCG | oiGEM1180_Dropout_5C1O_rev | TTGGTCTCCCTCAAGCTAGAGACGTATAAACGCAGAAAGGCCCACC | PCR, GGA | pMC_V_06_Part Entry sfGFP (Vn) Ori Muta |
| **pMC0_1*-6*_02_Dropout mScarlet-I** | oiGEM1179_Dropout_5C1R_fwd | AAGGTCTCGCTCGAACAAGAGACGGAAAGTGAAACGTGATTTCATGCG | oiGEM1180_Dropout_5C1O_rev | TTGGTCTCCCTCAAGCTAGAGACGTATAAACGCAGAAAGGCCCACC | PCR, GGA | pMC_V_07_Part Entry mScarlet-I (Vn) Ori Muta |
| **pMC0*_TU1-TU5_01_Dropout sfGFP** | oiGEM1177_Dropout_5C1C_fwd | AAGGTCTCGCTCGGGAGAGAGACGGAAAGTGAAACGTGATTTCATGCG | oiGEM1178_Dropout_3C5C_rev | TTGGTCTCCCTCAAGCGAGAGACGTATAAACGCAGAAAGGCCCACC | PCR, GGA | pMC_V_06_Part Entry sfGFP (Vn) Ori Muta |
| **pMC0*_TU1-TU5_02_Dropout mScarlet-I** | oiGEM1177_Dropout_5C1C_fwd | AAGGTCTCGCTCGGGAGAGAGACGGAAAGTGAAACGTGATTTCATGCG | oiGEM1178_Dropout_3C5C_rev | TTGGTCTCCCTCAAGCGAGAGACGTATAAACGCAGAAAGGCCCACC | PCR, GGA | pMC_V_07_Part Entry mScarlet-I (Vn) Ori Muta |

**Table S5. Construction of plasmids used in this project.** Short nomenclature indicates the level 0 parts used in the assembly (e.g., 1_03 represents pMC0_**1_03**_5C1RLF).

| Plasmid | Used parts | Description |
|---|---|---|
| **pDS_107_1R1 kan** | 1_03, 2_23, 3_07, 4_01, 5_02, 6_02, 7_05, 8_19 | Dropout plasmid for CDS parts |
| **pDS_108_1R1 kan** | 1_03, 2_23, 3_01, 4_12, 5_02, 6_02, 7_05, 8_19 | Dropout plasmid for RBS parts |
| **pDS_109_1R1 kan** | 1_03, 2_23, 3_07, 4_12, 5_01, 6_02, 7_05, 8_19 | Dropout plasmid for terminator and degradation tag parts |
| **pDS_114_1R1 kan** | 1_03, 2_01, 3_05, 4_12, 5_02, 6_02, 7_05, 8_19 | Dropout plasmid for promoter parts |
| **pDS_115 kan** | 1-6_02, 7_05, 8_19 | Empty control plasmid |
| **pDS_119_LVL2 kan** | 1_03,1_08, 2_21,2_23, 3_07, 4_02, 4_12, 5_01, 5_02, 6_02, 7_05, 8_19 | LVL2 dropout plasmid for terminator parts, mScarlet-I and *lux* reporter |
| **pDS_120_1R1 kan** | 1_03, 2-5_01, 6_02, 7_05, 8_19 | LVL1 backbone, position 1, sfGFP dropout, 5' insulator connector |
| **pDS_185_1R1 kan** | 1_05, 2-5_01, 6_02, 7_05, 8_19 | LVL1 backbone, position 1, sfGFP dropout |
| **pDS_186_22 kan** | 1_08, 2-5_01, 6_07, 7_05, 8_19 | LVL1 backbone, position 2, sfGFP dropout |
| **pDS_187_33 kan** | 1_11, 2-5_01, 6_10, 7_05, 8_19 | LVL1 backbone, position 3, sfGFP dropout |
| **pDS_188_44 kan** | 1_14, 2-5_01, 6_13, 7_05, 8_19 | LVL1 backbone, position 4, sfGFP dropout |
| **pDS_189_55O kan** | 1_17, 2-5_01, 6_17, 7_05, 8_19 | LVL1 backbone, position 5, sfGFP dropout |
| **pDS_190_11R (inv) kan** | 1_06, 2-5_01, 6_06, 7_05, 8_19 | LVL1 backbone, position 1, for inversion, sfGFP dropout |
| **pDS_191_22 (inv) kan** | 1_09, 2-5_01, 6_09, 7_05, 8_19 | LVL1 backbone, position 2, for inversion, sfGFP dropout |
| **pDS_192_33 (inv) kan** | 1_12, 2-5_01, 6_12, 7_05, 8_19 | LVL1 backbone, position 3, for inversion, sfGFP dropout |
| **pDS_193_44 (inv) kan** | 1_15, 2-5_01, 6_15, 7_05, 8_19 | LVL1 backbone, position 4, for inversion, sfGFP dropout |
| **pDS_194_5O4 (inv) kan** | 1_19, 2-5_01, 6_19, 7_05, 8_19 | LVL1 backbone, position 5, for inversion, sfGFP dropout |
